# In vivo photocontrol of microtubule dynamics and integrity, migration and mitosis, by the potent GFP-imaging-compatible photoswitchable reagents SBTubA4P and SBTub2M

**DOI:** 10.1101/2021.03.26.437160

**Authors:** Li Gao, Joyce C.M. Meiring, Adam Varady, Iris E. Ruider, Constanze Heise, Maximilian Wranik, Cecilia D. Velasco, Jennifer A. Taylor, Beatrice Terni, Jörg Standfuss, Clemens C. Cabernard, Artur Llobet, Michel O. Steinmetz, Andreas R. Bausch, Martin Distel, Julia Thorn-Seshold, Anna Akhmanova, Oliver Thorn-Seshold

## Abstract

Photoswitchable reagents to modulate microtubule stability and dynamics are an exciting tool approach towards micron- and millisecond-scale control over endogenous cytoskeleton-dependent processes. When these reagents are globally administered yet locally photoactivated in 2D cell culture, they can exert precise biological control that would have great potential for *in vivo* translation across a variety of research fields and for all eukaryotes. However, photopharmacology’s reliance on the azobenzene photoswitch scaffold has been accompanied by a failure to translate this temporally- and cellularly-resolved control to 3D models or to *in vivo* applications in multi-organ animals, which we attribute substantially to the metabolic liabilities of azobenzenes.

Here, we optimised the potency and solubility of metabolically stable, druglike colchicinoid microtubule inhibitors based instead on the styrylbenzothiazole (SBT) photoswitch scaffold, that are non-responsive to the major fluorescent protein imaging channels and so enable multiplexed imaging studies. We applied these reagents to 3D systems (organoids, tissue explants) and classic model organisms (zebrafish, clawed frog) with one- and two-protein imaging experiments. We successfully used systemic treatment plus spatiotemporally-localised illuminations *in vivo* to photocontrol microtubule dynamics, network architecture, and microtubule-dependent processes in these systems with cellular precision and second-level resolution. These nanomolar, *in vivo*-capable photoswitchable reagents can prove a game-changer for high-precision cytoskeleton research in cargo transport, cell motility, cell division and development. More broadly, their straightforward design can also inspire the development of similarly capable optical reagents for a range of protein targets, so bringing general *in vivo* photopharmacology one step closer to productive realisation.

## 1. Introduction

Biological instrumentation, methods, and technologies now allow us to observe cellular structures and dynamics on the micron scale, with dynamics resolved to the scale of milliseconds, in settings from *in vitro* cell culture^1^ to *in vivo* (multi-organ) animal models^2^. The development of tools to manipulate these processes with matching micrometre spatial precision and millisecond temporal precision has lagged far behind, despite its clear value for research.^3,4^ One biological system with particularly urgent need of such tools is the microtubule (MT) cytoskeleton.^5,6^ MTs are giant tube-like noncovalent polymers of α/β-tubulin protein heterodimers, that are centrally structured and rapidly remodelled to support hundreds of spatiotemporally-regulated functions in the life of a cell. The most visible roles of MTs include force generation to maintain and change cell shape and position, and in scaffolding the transport of cargos by motor proteins, including chromosomes during cell division.

Tools to noninvasively manipulate MT network structure and remodelling dynamics with high spatiotemporal precision, have great potential to drive biological research. However, to fulfil this potential, tools must succeed in settings from 2D cell culture (e.g. cell migration and division) through to *in vivo*, 3D systems (embryonic development and neuroscience). It is also important that tools be easily transferrable across different models and organisms.^7,8^

Optogenetics has been extremely successful in patterning ion currents and cell signalling with high spatiotemporal precision. However, in MT cytoskeleton studies it has not yet succeeded, with the exception of one photo-inactivated fusion protein that stops promoting MT polymerisation under illumination.^9^ Optogenetics also faces difficulties for *in vivo* use that are common to all genetic approaches, including the costs of translating tools across species (time for breeding and validating transgenic lines; optimisation of expression systems; etc).

By comparison, drugs to modulate MT structure and dynamics reliably across all eukaryotes are very well-studied. Taxanes, epothilones, colchicine analogues, and vinca alkaloids are all used for nonspecific suppression of MT-dependent cellular processes, in settings from single-cell studies through to *in vivo* therapeutic use in humans. Photouncaging approaches have for years been applied to these drugs to improve the spatiotemporal precision with which their activity can be applied in cellular research (ideally towards the scale of μm and ms).^10^

Photoisomerisation-based drug analogues or “photopharmaceuticals” elegantly avoid many of the drawbacks that limited the *in vivo* use of typical photouncaging methods:^11^ such as toxic and/or phototoxic byproducts, requirements for <360 nm wavelengths and high light intensities, slow post-illumination fragmentation, irreversibly photosensitive stock solutions, and non-optical drug release mechanisms (e.g. enzymatic hydrolysis of cages). In the MT field, recent photoswitchable analogues of taxane^12^, epothilone^13^ and colchicinoid^5,14–19^ MT inhibitors have all been applied to cellular studies. These photopharmaceuticals have enabled noninvasive, reversible optical control over MT dynamics and MT-associated downstream effects, with cell-specific spatial precision and sub-second-scale temporal precision, and have been brought to bear on research in embryology^20^, neuroscience^21^, and cytoskeleton^5^.

However, the typical photoswitch scaffolds introduce problems particularly for *in vivo* application, three of which this paper will focus on: (1) *Metabolic stability*: azobenzenes, particularly as their *Z*-isomers, are reductively degraded by cellular glutathione (GSH).^22–24^ This reduces photoswitchability as well as potency, and on timescales typical for *in vivo* studies (> hours) the degradation byproducts are likely to give off-target effects, particularly in metabolically active multiorgan systems (tested below, and also discussed elsewhere^16^). (2) *Orthogonal photocontrol and imaging*: azobenzenes, and the less common hemithioindigos, are isomerised by excitation of the typical imaging labels used in biological studies (GFP/fluorescein at 490 nm, YFP at 514 nm); some also isomerise with RFP/rhodamine imaging at 561 nm (at focussed laser intensities).^16,25^ Typically this leaves only the 647 nm laser channel available for orthogonal imaging during photoswitching, yet this laser typically only addresses small molecule probes. These scaffolds’ inability to allow orthogonal fluorescent-protein-based imaging during photocontrol is particularly problematic for *in vivo* research that extensively relies on two- or three-channel imaging with fluorescent protein fusions to resolve processes with biological specificity. (3) *Practical applicability issues*: *in vivo* studies must ensure sufficient delivery of photopharmaceuticals to target tissues without organic cosolvents, as these are far more toxic to animals than they are in cell culture. This either requires high-potency compounds or chemically robust solubilising strategies: neither of which are often seen with typically hydrophobic azobenzene or hemithioindigo drug analogues. To minimise illumination while maximising bioactivity explicitly requires tuning the photoswitch scaffold’s photoresponse *at those laser wavelength/s* that are in practice used for photoconversion in the biological study^16^ (aim: high efficiency of isomerisation E(λ)^5^ and high photostationary state): since photoswitch performance at optimal but unavailable wavelengths (e.g. 260-380 nm) is irrelevant.

Novel photoswitches that improve practical performance are therefore gaining attention to expand the biological scope of photopharmacology.^26–28^ To tackle these three issues in the context of microtubule photocontrol, we introduced styrylbenzothiazoles **SBTub2/3**(**Fig 1a**) as highly metabolically stable, fully GFP/YFP/RFP-orthogonal photoswitchable tubulin inhibitors (**Fig 1b-c**), with excellent photoresponse to the 405 nm laser line that is standard in confocal microscopy. **SBTub3** could photocontrol microtubule dynamics, organization, and MT-dependent processes in live cells with reversible temporal patterning, and with cellular and even sub-cellular spatial precision.^16^

**Figure 1:**
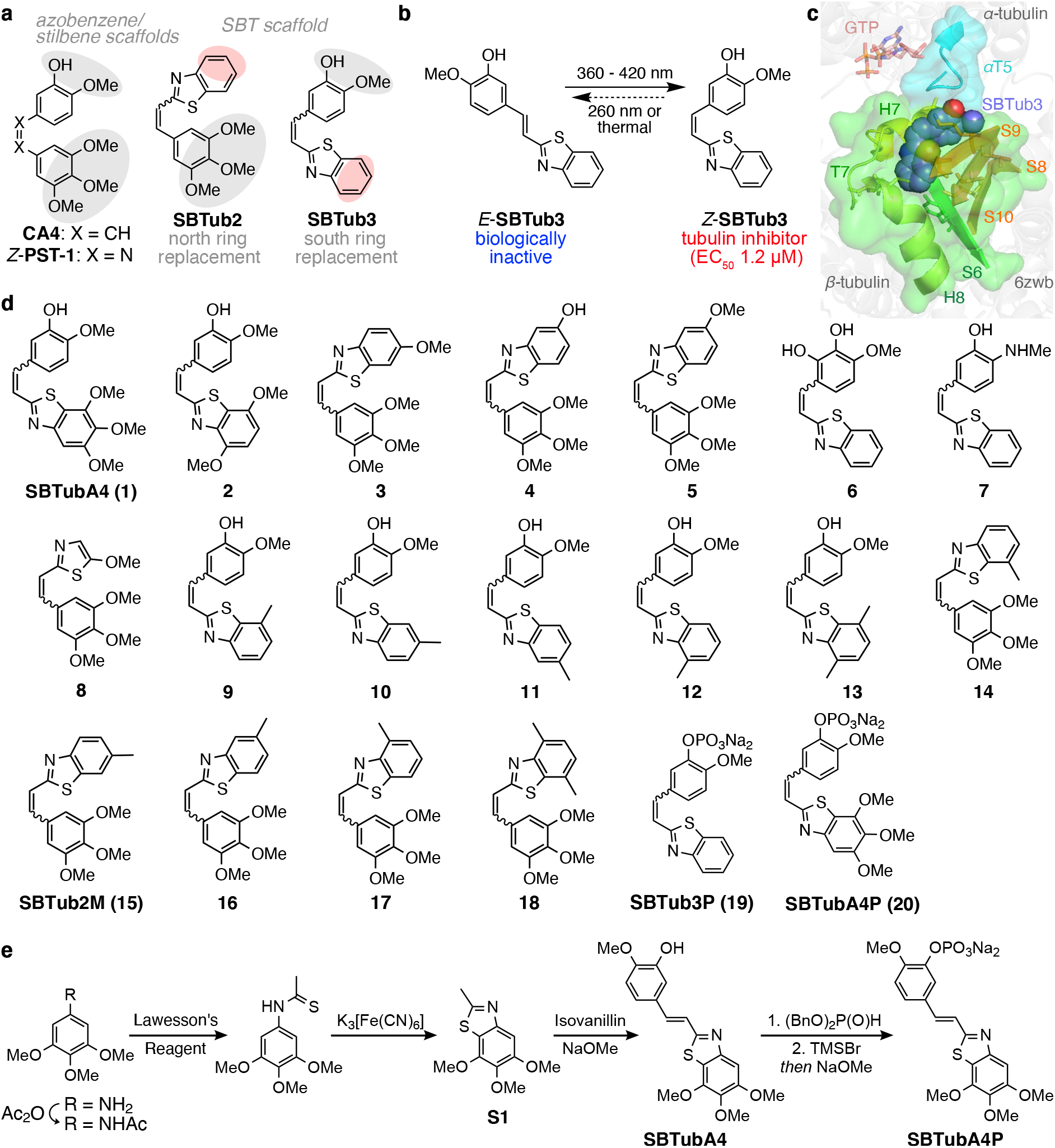
Design and synthesis. **(a)** The colchicinoid pharmacophore (grey shaded trimethoxyphenyl south ring and isovanillyl north ring) can be applied to various scaffolds, giving photoswitchable azobenzene-based **PST** and SBT-based **SBTub** antimitotics. Previously published **SBTub2/3** lacked key interaction residues (red shaded sites). **(b)** *Z-***SBTub3** inhibits tubulin polymerisation and MT-dependent processes. **(c)** X-ray structure of tubulin:*Z-* **SBTub3** complex (carbons as purple spheres). The south ring is buried in β-tubulin (green); only the north ring interacts with α-tubulin at the α-T5 loop (cyan). **(d)** Evolved **SBTub** compound library used in this paper. **(e)** Typical synthesis of **SBTub**s proceeds by acetanilide sulfurisation, Jacobson cyclisation, and basic condensation. Phosphate prodrug **SBTubA4P** is further accessed by oxidative phosphoester formation and deprotection.

The metabolic stability and imaging-orthogonal photocontrol of **SBTub3** were excellent features towards *in vivo* use, but practical limitations remained, which we determined to address in this study. Primarily, while their parent colchicinoid inhibitor combretastatin A-4 (**CA4**) has ca. 20 nM cellular potency, the slightly larger *Z-***SBTub2/3** gave only micromolar bioactivity (though note that the **CA4**-isosteric azobenzene analogue *Z-***PST-1** has only ca. 500 nM potency; **Fig 1a**). The lower the potency, the more compound and the more cosolvent would be needed, which we found blocked animal model applications of both **SBTub2/3** and **PST-1**(see below). Thus we prioritised a structure-activity relationship study to improve the potency of **SBTub**s. Secondly, the **SBTub**s are hydrophobic, so we prioritised fully water-soluble prodrugs for *in vivo* use without cosolvents. Thirdly, while the SBT scaffold is essentially a unidirectional (*E→Z*) photoswitch in the biological range because its isomers’ absorption bands overlap, we were curious if tautomerisable electron-donating substituents could accelerate its spontaneous thermal *Z→E* relaxation, as they do for azobenzenes.^25^

Our goal was then to test whether suitably potent and soluble SBT derivatives could be used for photocontrol not only in cell culture settings where many photoswitches succeed, but over a range of *in vivo* multi-organ animal models, with temporally-specific and cell-precise *in situ* photoswitching after systemic administration: settings in which no other photoswitchable reagents have so far succeeded. We now report the development of these potent, soluble, metabolically stable, GFP-orthogonal SBT-based photopharmaceuticals and characterise the optimal ligand **SBTubA4** with a tubulin:SBT X-ray crystal structure. We then showcase the unprecedented success of its fully water-soluble prodrug **SBTubA4P** in allowing systemic administration but local photoactivations to achieve (i) spatiotemporally-specific control over MT dynamics in cell culture, (ii) long-term spatially-specific control over development and migration in 3D organoid culture, (iii) short-term temporally-resolved photocontrol of neural development in fly brain explant, (iv) photocontrol of embryonic development in the clawed frog, and (v) temporally reversible photocontrol of microtubule dynamics in zebrafish.

## 2. Results

### 2.1 Cellular structure-activity optimisation of SBTubs

For a colchicinoid **SBTub** to be cellularly effective, its *Z-*isomer should bind tubulin,^5,29^ halting cell proliferation and inducing apoptosis^30^; while the corresponding *E-*isomer should have negligible tubulin binding and no other significant toxicity mechanisms over a wide concentration range where the *Z-*isomer is bioactive, resulting in a high lit/dark bioactivity ratio (“photoswitchability of bioactivity”).^16^ Thus we performed cellular structure-activity optimisation by comparing the proliferation of cells treated with **SBTub**s under *in situ* pulsed illuminations with near-UV light (“lit”: mostly-*Z*-isomer; aim: nanomolar potency), to their proliferation without illumination (“dark”: all-*E*-isomer; aim: no antiproliferative effects up to 100 μM).

Our first priority for *in vivo* use was to increase *Z-***SBTub** potency. We first tested whether restoring the methoxy groups of the **CA4** pharmacophore would increase potency despite the extra size, with **SBTubA4**(**1**). This, like most **SBTub**s, was synthesised in a short sequence (**Fig 1e**) via basic aldol condensation of the derivatised 2-methylbenzothiazole with the corresponding benzaldehyde, with the 2-methylbenzothiazole obtained *via* Jacobson cyclization^31^ of the thioacetanilide using potassium ferricyanide (exceptions were *para-*aniline **7** which was obtained by Horner-Wadsworth-Emmons olefination; and **9** and **11** where the 2,5- and 2,7-dimethylbenzothiazoles were synthesised by Ullmann-type coupling according to Ma^32^ since Jacobson cyclization of 3-methyl-thioacetoanilide gave an inseparable mixture; see Suppporting Information for details).

**SBTubA4** was an early hit, with ca. 10-fold more potent *Z-*isomer binding than previous best **SBTub3** and with an excellent dark/lit ratio of 30 (**Table 1**; **Fig 2c**. We then tested if the *Z-***SBTub**s obey similar structure-activity relationships as known for **CA4**. First, we rearranged the methoxy groups in **2**, reducing the *Z-*potency, which matched literature expectations^33^ as the middle methoxy group on the south ring otherwise accepts a hydrogen bond from β-Cys239. We also flipped the SBT scaffold orientation in analogues **3-5**. This was not convenient for installing the hydroxy/methoxy north ring substituent pair, so in **3** and **4** we retained only one of these groups and the compounds suffered a predictable^33^ though small loss of potency. The seven-fold potency loss when moving the methoxy group from the spacefilling 6-position to the 5-position which is best occupied by a small polar substituent^33^ (compound **5**) was striking and expected. We concluded that indeed *Z-***SBTub**s obey similar structure-activity relationships as **CA4**, which could guide further development.

**Table 1:** cytotoxicity EC_50_ values (mostly-*Z* lit (360-450 nm) and all-*E* dark) and dark/lit EC_50_ ratio; and standard photoactivation working concentration [I]_WC_ (defined later), all in HeLa cell line.

| compound | EC<sub>50</sub> (μM) |  | EC<sub>50</sub> ratio<br>dark / lit | [I]<sub>WC</sub><br>(μM) |
| --- | --- | --- | --- | --- |
|  | lit | dark |  |  |
| **SBTub2** | 1.4 | >25 | >18 | 6.0 |
| **SBTub3** | 1.2 | >25 | >21 | 3.2 |
| **SBTubA4** | 0.12 | 3.8 | 31 | 0.21 |
| **2** | 0.52 | >100 | >190 | 1.6 |
| **3** | 0.20 | 15 | 75 | 0.65 |
| **4** | 0.67 | 9.3 | 14 | 1.5 |
| **5** | 1.4 | 18 | 13 | 3.2 |
| **6** | 23 | 27 | n.d. | n.d. |
| **7** | 2.6 | >35 | >13 | n.d. |
| **8** | 1.2 | >100 | >83 | 3.0 |
| **9** | 1.0 | 48 | 48 | 2.6 |
| **10** | 3.0 | >100 | >33 | 10 |
| **11** | 1.7 | >100 | >58 | 4.3 |
| **12** | 0.77 | 64 | 83 | 1.6 |
| **13** | 0.42 | 8.8 | 21 | 1.8 |
| **14** | 1.0 | >15 | >14 | 2.7 |
| **SBTub2M** | 0.035 | 7.0 | 200 | 0.06 |
| **16** | 0.52 | 48 | 92 | 1.1 |
| **17** | 0.23 | 40 | 174 | 0.65 |
| **18** | 3.0 | >15 | 5 | 10 |
| **SBTub3P** | 3.4 | >20 | >5 | 7.5 |
| **SBTubA4P** | 0.052 | 0.45 | 8.7 | 0.10 |

**Figure 2:**
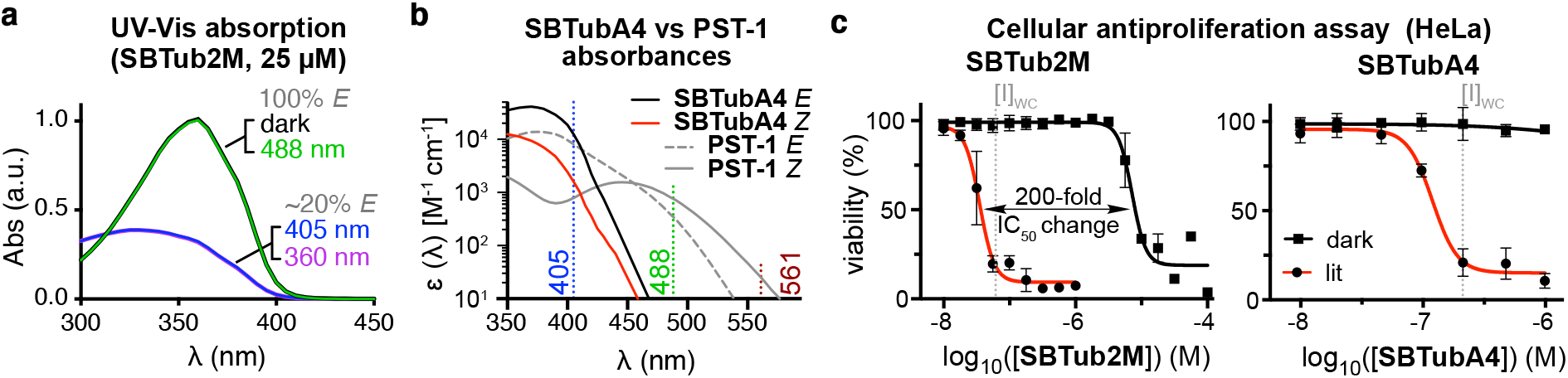
Photocharacterisation. **(a) SBTub2M** is not isomerised from its all-*E* “dark” state by 488 nm illumination, but is photoswitched to majority-*Z* “lit” states by UV/violet light (78% *Z* at 405 nm by NMR; 50:50 PBS:DMSO). **(b)** Comparison of absorbance spectra of **SBTubA4** and azobenzene **PST-1** illustrates the SBT’s ideal match to 405 nm photoactivation, combining stronger 405 nm absorption, with sharper absorption cut-off above 405 nm which makes it orthogonal to GFP (488 nm), YFP (514 nm) and RFP (561 nm) imaging. **(c)** Leads **SBTub2M** and **SBTubA4** have highly nonlinear dose-response profiles, high lit/dark ratio of bioactivity, and mid-nanomolar [I]_WC_ values.

At this stage we wished to test if common strategies to accelerate thermal *Z→E* relaxation and redshift spectral response in azobenzenes could also be applied to **SBTubs**. We created close steric matches of bioactive **SBTub3** but with tautomerisable *ortho*-catechol **6** and *para-* amino **7** groups. Since *o-*hydroxy derivatives of **CA4** have entered clinical trials^33^, we were surprised that **6** gave negligible potency under lit (and dark) conditions; however **7** retained similar light-specific bioactivity as its isostere **SBTub3** (discussed below in section Photoswitch Performance).

Continuing potency optimisation, we also tested whether retaining the **CA4** methoxy groups on a reduced-size photoswitch would be beneficial. We created styrylthiazole (ST) **8** which can be considered a near-perfect isostere of **CA4**/**PST-1** minus the small polar hydroxyl group, or a shrunken analogue of **3**. As far as we know, STs have never been used in photopharmacology before. We considered them interesting as we expected them to retain the metabolic stability and GFP-orthogonality of SBT, while additionally being isosteric to the better-known azobenzene: which offers attractive possibilities for adapting known azobenzenes into potentially biologically more applicable ST-based photopharmaceuticals. **8** was accessed by closing the thiazole, starting from a methyl cinnamoylglycinate (see Supporting Information). Pleasingly, **8** gave strong *Z*-specific cellular bioactivity with ca. 100-fold photoswitchability (**Table 1**), although its 10-fold loss of potency compared to **1** suggested that we should explore **SBTubs** intermediate in size between **SBTub3** and **1**.

Therefore, for potency optimisation, we tested both adding methyl groups to **SBTub3** and “walking” them around the scaffold south ring (**9-13**); and also deleting the north ring hydroxy/methoxy substituent pair from **1** and replacing them with smaller methyls (**14-18**). Note that the SBT *Z-*isomers orient their sulfur towards the inner face of the molecule^16^ so we did not rely on rotations around the alkene—benzothiazole single bond to ‘reflect’ substituents into similar positions (e.g. **9** vs **12**, or **10** vs **11**).

South ring derivatives **9-13** did not approach the potency of **1-5**, although **10-11** still have excellent performance (~1 μM *Z-*potency, ~100-fold photoswitchability of bioactivity). However, north ring derivative **SBTub2M**(**15**) in which the methyl group replaces the spacefilling OMe group, was six times as potent as its methoxy homologue **3** (*Z-*IC_50_ 35 nM) while having excellent photoswitchability of bioactivity (ca. 200-fold; **Fig 2c**). This makes it the most potent *and* the most photoswitchable of the photoswitchable antimitotics known. **16**(methyl at the small polar position) and **14** and **18** (methyl projecting to inner face, clashes with protein) were predictably weaker than **SBTub2M**(though **16** still has submicromolar *Z* activity and ca. 100-fold switching), while **17** that projects the methyl to the outer face (towards the exit tunnel) was well-tolerated (0.2 μM) and retained nearly 200-fold switching. It was highly satisfying that this methyl walk SAR also followed the SAR known for **CA4**, as this supports that their target and binding site are conserved.

Our second priority was to develop soluble **SBTubs** for *in vivo* use in multiorgan animal models, which do not easily tolerate even low amounts of organic cosolvents. We aimed to formulate water-soluble **SBTub** prodrugs by phenol phosphorylation – an approach that has been generally successful for colchicinoid inhibitors (including clinically-advanced **CA4P**, **BNC105P** etc)^34^. We had initially produced **SBTub3P**, the phosphate prodrug of previous lead compound **SBTub3**, however its aqueous solubility was only moderate (<2 mg/mL) potentially due to aggregate formation by π-stacking of the planar compound. We synthesised **SBTubA4P** hoping that the out-of-plane central methoxy group would reduce π-stacking, which combined with the hydrophilicity of the three extra methoxy groups would give better solubility, as we have seen in other contexts^6,19^. Indeed, **SBTA4P** dissolved to at least 10 mM in water, which was to prove important in later assays. Both prodrugs **SBTubA4P** and **SBTub3P** had similar cellular *Z* and *E* potencies as their drug forms.

In summary, we had developed **SBTubA4** and **SBTub2M** as mid-nanomolar antimitotics with 30-200-fold photoswitchability of bioactivity for cell culture studies (both cross-validated on A549 lung carcinoma cell line, **Fig S5**); and we had further developed **SBTubA4P** as a convenient fully water-soluble prodrug of **SBTubA4** for *in vivo* applications. These became the focus of our further biological evaluations.

### 2.2 SBTub photoswitch performance studies

The photoresponses of most **SBTub**s (**1-5** and **9-20**) were similar to previously reported SBTs **SBTub2/3**,^16^ with absorption maxima and absorption cut-offs being excellently balanced for both efficient *E→Z* photoswitching with the common 405 nm microscopy laser, and for full orthogonality to GFP imaging (ca. 488 nm). The separated π→π* absorption maxima for *E*-(~360 nm) and *Z*-isomers (~330 nm) enable efficient directional *E→Z* photoisomerisation at 360-420 nm reaching ca. 80% *Z* (**Fig 2a-b**), and extinction coefficients at 405 nm were up to twice those of similar azobenzenes (**Fig 2b**), promising high-efficiency photoactivation on the confocal microscope. Importantly, *E-* and *Z*-SBT absorptions drop sharply towards zero above 410 nm (**Fig 2b**, **Fig S1-S2**) which is crucial for avoiding photoresponse to 488 nm GFP imaging under high-intensity focussed lasers, as well as with broader filtered excitation sources e.g. 490±25 nm: since absorption “tails” extending far beyond band maxima can otherwise cause substantial photoswitching. (For example, 561 nm RFP imaging on the confocal microscope photoisomerises azobenzene **PST-1** despite its extinction coefficients being < 30 cm^−1^M^−1^.^21^) However, the **SBTubs**’ cutoff suggested they would indeed be GFP-orthogonal: which was later confirmed and found to be crucial for *in vivo* use (see below).

In biological settings (>360 nm) and on the population level, these SBTs act similarly to photoactivation probes: (i) illuminations all give similar majority-*Z* equilibrium photostationary states (PSSs) in the photoresponsive spectral region (360 – ca. <440 nm), and (ii) they show no significant (<2%) *Z→E* thermal relaxation after hours in physiological buffers at pH~7 at 25°C (**Fig S3**-**S4**). However, they have all the other practical advantages of photoswitches that are relevant to most research uses^11^ (no toxic/phototoxic byproducts, fast illumination response, no non-optical drug activation mechanisms, and stocks can be quantitatively relaxed to *E* by warming to 60°C which is advantageous for stock handling over sequential assays). Their photostability to continuous illumination at >380 nm was excellent. These features are shared with previously-reported **SBTub2/3**.^16^

We used *ortho*-hydroxy **6** and *para-*amino **7** to test if strong electron donor groups that can tautomerise to freely-rotatable quinoids (C-C single bond instead of C=C), can accelerate thermal *Z→E* relaxation to make better-reversible **SBTub**s; and if these would induce spectral redshifting. However, in cuvette, *Z-***6/7** thermally relaxed only slowly (halflives ≫ hours), and since *Z-***7** gave similar cytotoxicity in the longterm cellular assay as its isostere **SBTub3**, we concluded that its cellular relaxation was not fast on a biological timescale either. The spectra of **7** were redshifted by nearly 60 nm compared to typical SBTs putting the *E-***7** absorption maximum exactly at the 405 nm laser line (**Fig S1**), however, as neither **6/7** brought relaxation rate benefits, we did not pursue them.

Styrylthiazoles as in **8** have not yet been studied as scaffolds for photopharmaceuticals^13^. We were pleased that its isomers’ spectra were only ca. 25 nm blueshifted compared to the SBTs (giving less efficient isomerisation at 405 nm), and all other properties, such as its completeness of *E→Z* photoswitching, were similar to the SBTs. We believe this opens up new possibilities for GFP-orthogonal, metabolically-resistant photoswitches that are isosteric to azobenzenes, and so can replace them for applications where these biologically advantageous properties are required.

Given that each SBTub reached a PSS that was nearly invariant with excitation wavelengths in the range 360-440 nm, we defined a single “typically useful working concentration” **[I]_WC_** for cell culture use in HeLa cell line by closer examination of the dose-response curves (**Fig 2c**, **Table 1**, **Fig S5**). At [I]_WC_, illuminated cells should experience strong inhibition while non-illuminated cells experience insignificant inhibition. We systematically define [I]_WC_ as *either* the lowest concentration where the lit efficacy reaches >80% of its plateau, *or* the highest concentration where treatment in the dark causes <10% difference of biological effect as compared to the untreated control: whichever is the lower. Szymanski has previously proposed an alternative working concentration metric [I]_opt_, to maximise illuminated efficacy, which is based on idealised highly nonlinear dose-response curves^4^ that are very rarely obtained in practice. Our empirical [I]_WC_ instead emphasises that baseline/background inhibition of the system must be minimal for photopharmaceutical control to be biologically useful. For example, even when IC_50_ values are well-separated, [I]_WC_ values are undefined if background bioactivity is too strong for the reagent to be useful (e.g. **7**, **Table 1**). We believe that ranking by [I]_WC_ is the most useful systematic single-value method for early-stage optimisation of photopharmaceuticals’ general performance, and that [I]_WC_ will come to find traction in the community.

For all further details, including full discussion of photoswitch performance of **6**/**7** and of *E-***SBTub** toxicity, see Supporting Information including **Fig S1-S3**.

### 2.3. SBTubs isomer-dependently target tubulin in cells

To test their cellular mechanism of bioactivity, we first examined the **SBTubs’** isomer-dependent inhibition of polymerisation of purified tubulin protein in a cell-free assay. Both leads **SBTubA4** and **SBTub2M** were non-inhibiting in the all-*E* dark state, permitting tubulin to polymerise as in the cosolvent control (**Fig 3a**). However, in the majority-*Z* illuminated state they were potent inhibitors that suppressed polymerisation entirely to baseline readout levels, comparable to the archetypical colchicinoid nocodazole. Matching the antiproliferation assays, they were significantly stronger inhibitors than first generation **SBTub3**(**Fig 3a**).

**Figure 3:**
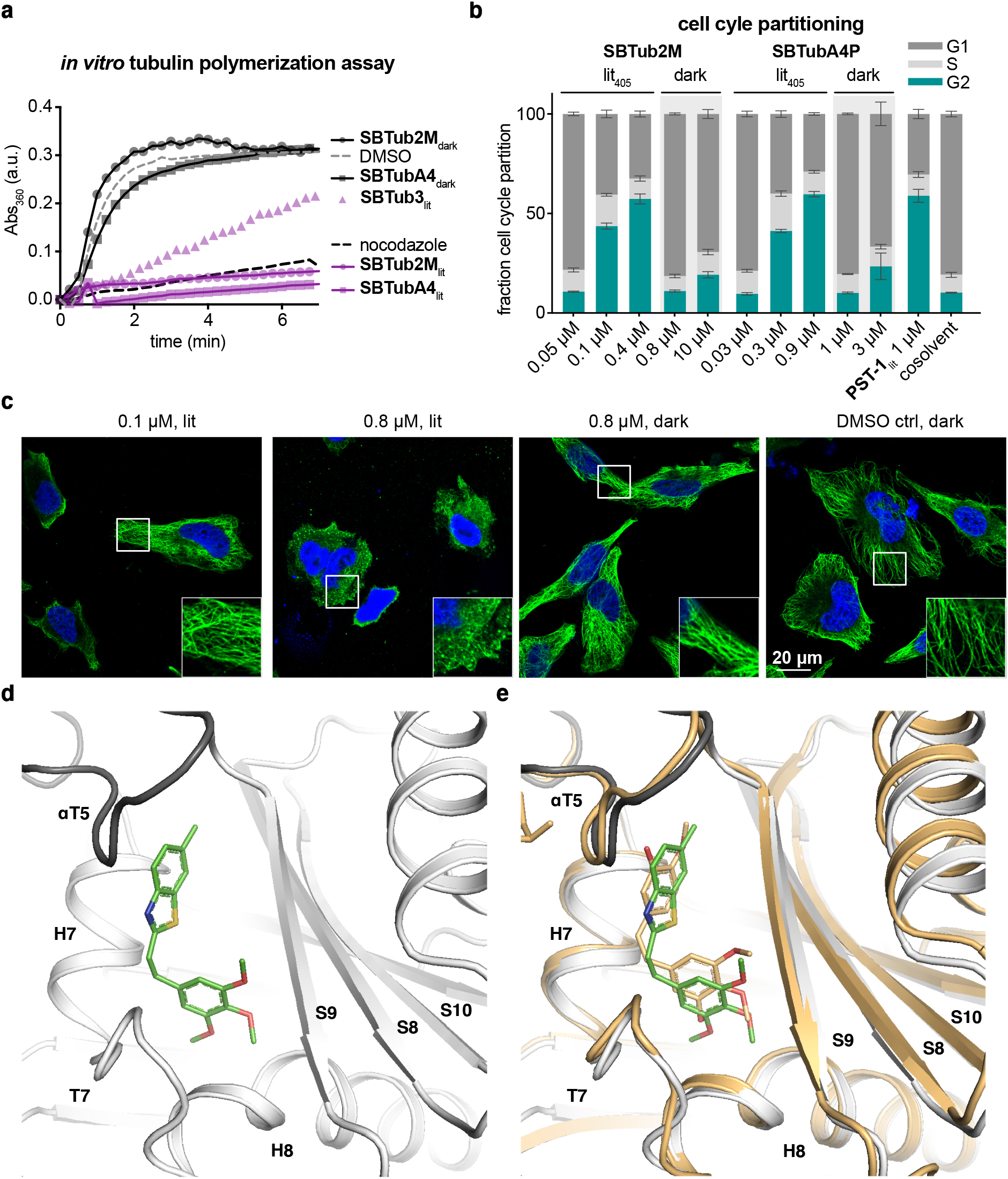
Tubulin-specific cellular mechanism. **(a) SBTubs** light-dependently inhibit tubulin polymerisation (turbidimetric cell-free assay; absorbance mirrors the degree of polymerisation; “lit” denotes use of **SBTubs** (20 μM) pre-isomerised to majority-*Z* PSS with 405 nm, nocodazole control at 10 μM. **(b)** Cell-cycle analysis of Jurkat cells treated with **SBTub2M** and **SBTubA4P** matches photoswitchable reference **PST-1**: with significant G_2_/M arrest under 405 nm pulsing (‘‘lit’’), but without cell cycle effects in the dark (matching cosolvent controls). **(c)** Immunofluorescence imaging of cells treated with **SBTubA4P** under pulsed 405 nm illuminations (“lit”, mostly-*Z*) and in the dark (all-*E*), compared to cosolvent control (HeLa cells, 20 h incubation; α-tubulin in green, DNA stained with DAPI in blue). **(d)** Close-up views at the colchicine-binding site of X-ray crystal structure of the Darpin D1:tubulin:*Z-***SBTub2M** complex (dark grey α-tubulin, light grey β-tubulin in cartoon representation; *Z-***SBTub2M** in stick representation with carbons green, oxygens red, nitrogen blue, and sulfur yellow. **(e)** Superimposition of tubulin:**CA4**(orange carbons; PDB 5LYJ) and TD1:*Z-***SBTub2M**(green carbons) structures (see also **Fig S11a-f**).

If microtubule inhibition is also the main cellular mechanism of action of *Z-***SBTubs**, we would expect *Z-***SBTub**-treated cells to show G_2_/M-phase cell cycle arrest due to mitotic checkpoint failure.^12^ We therefore used flow cytometry-based analysis to quantify cell cycle repartition. *E*-**SBTubs** caused either no change or small changes compared to controls, whereas G_2_/M arrest was strongly induced by *in situ-*lit (mostly-*Z*) **SBTubA4P** and **SBTub2M**(**Fig 3b**, **Fig S6**).

As G_2_/M-phase arrest is necessary but insufficient to conclude on cellular tubulin inhibition being their major mechanism of action, we next performed confocal microscopy imaging of the MT network architecture in immunofluorescently stained cells, to obtain a direct readout of tubulin-inhibiting effects. *In situ-*illuminated **SBTubA4P** treatment caused microtubules to curve and MT architecture to break down,^35^ but *E-***SBTubs** caused no disorganization at corresponding concentrations in the dark, matching cosolvent controls (**Fig 3c**). This matches the assumption that their photoswitchable cytotoxicity arises from their *Z-*isomers potently inhibiting MT dynamics and stability in cells. While microscopy is a qualitative method that can misrepresent population-level statistics, the quantitative cell cycle data from flow cytometry (10^4^ cells per condition) as well as the qualitative match to previous SBTub work^16^ give confidence to this result.

Lastly, to check our assumption that *Z-*SBTubs act as colchicinoids, we crystallised the *Z-***SBTub2M**:tubulin DARPin D1 (TD1) complex. X-ray diffraction studies showed that *Z-* **SBTub2M** indeed directly binds tubulin at the colchicine site (**Fig 3d**) very similarly to **CA4**, even with the same orientation of its substituents (**Fig 3e; Fig S11a-f**). This explains why the **SBTub** SAR determined in this study (**Table 1**) matched to that known for **CA4**. It also highlights the plasticity of the binding site, which accepts such differently-sized inhibitors.

Taken together, these cell-free and cellular assays support that the **SBTubs** act as light-dependent tubulin inhibitors in cells, with their *Z-*isomers binding potently at the colchicine binding site and their *E-*isomers having no effects at the corresponding concentrations.

### 2.4. SBTub photocontrol enables cell-precise, temporally reversible MT inhibition in 2D cell culture

Aiming later to apply **SBTubs** to photocontrol MT dynamics in complex models and *in vivo*, we now switched to using the fully water-soluble prodrug **SBTubA4P**. This is an important step: (i) it avoids cosolvents that can be problematic for *in vivo* toxicity; and (ii) it prevents hydrophobic adsorption onto the matrix materials (PDMS, collagen, agarose) that are used in 2D structured surfaces, 3D cell culture/organoid models, and for embedding motile animals during longterm imaging. Avoiding adsorption is important in our experience, as hydrophobic compounds can exhibit irreproducible apparent potencies or effects in these settings, which is timewise- and ethically prohibitive for resource-intensive low-throughput animal studies.

Before performing animal work, we probed the spatiotemporal resolution that **SBTubs** could achieve for *in situ-*photoswitching-based control over MT dynamics, in 2D cell culture. Using spinning disc confocal live cell microscopy we imaged **SBTubA4P**-treated cells transfected with a fluorescently-labelled fusion of the MT end binding protein EB3, to directly monitor MT polymerisation dynamics during photoswitching.^16^ This is possible since EB3 labels the GTP-cap of MTs, so EB3-tdTomato acts as a fluorescent marker revealing the tips of polymerising MT plus ends in cells as hundreds of dynamic “comets” that cascade through the cell at significant velocities (tdTomato excitation at 561 nm).^36^ Our protocol for cell-precise photoswitching of MT dynamics, with internal controls for compound application and for photobleaching, was as follows. We imaged transfected cells before **SBTubA4P** application to establish untreated MT dynamics baselines, simultaneously controlling for effects of 405 nm laser pulses on single selected cells (targeted by region of interest (ROI) illumination); then we added *E-***SBTubA4P** to these same cells, and continued imaging the whole field of view while applying targeted pulses of the 405 nm laser to a single ROI-selected cell.

**SBTubA4P** enabled repeatable cycles of temporally reversible, photoswitching-induced inhibition of MT dynamics in live cells, with single-cell spatial targeting precision and second-scale onset time precision (**Fig 4a-b**, **Movie S1**). In **SBTubA4P**-treated ROI cells, within seconds upon each single frame 405 nm pulse, polymerising MT tips stop moving and disappear, then more slowly reappear and resume movement (best seen in **Movie S1**). Statistics collected over multiple independent experiments showed these inhibition spikes are highly reproducible; recovery towards uninhibited baseline has a halflife of ca. 25 s, which we attribute to the diffusion of *Z-***SBTub** out of the ROI cell (**Fig 4a**). There were minor effects on MT dynamics in treated non-ROI neighbour cells as compared to pre-treatment controls; and the 405 nm pulsing protocol alone did not cause any readout changes (**Fig 4a**).

**Figure 4:**
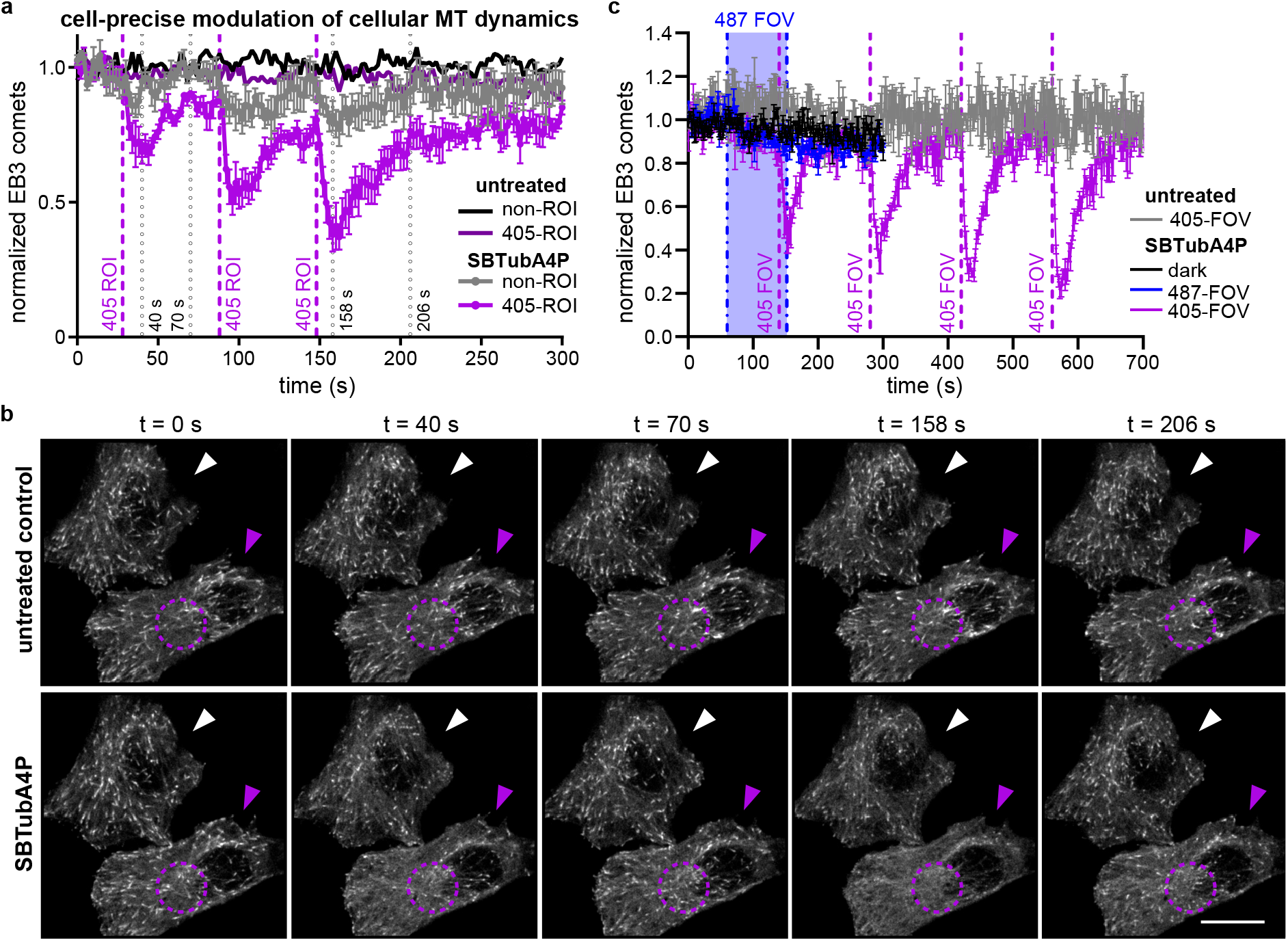
Spatiotemporal control over MT dynamics in 2D-cultured HeLa cells. **(a-b)** MT inhibition in **SBTubA4P**-treated cells is initiated only upon 405 nm illumination pulses and only in ROI-targeted cells (data related to **Movie S1**; live cell EB3-tdTomato “comets” quantify polymerising MTs). **(a)** Comet count statistics are similar to cosolvent-only baseline in both ROI-pulsed-cosolvent and non-ROI-**SBTubA4P** conditions; ROI-**SBTubA4P** statistics show inhibition spikes. **(b)** Stills from **Movie S1** at the times indicated in **(a)**, initially during the untreated timecourse, then during the **SBTubA4P**-treated timecourse on the same cells. Purple arrowhead indicates the ROI cell, purple dotted circle indicates where the 405 nm ROI is applied at times 26 s, 88 s and 148 s, white arrowhead indicates the non-ROI cell quantified as internal control (scale bar 15 μm). **(c)** EB3 comet counts of cells imaged at 561 nm only (“dark,” grey), with 47 frames at 487 nm applied to full field of view during the time span indicated with dashed lines (“487,” cyan) and **SBTubA4P**(6 μM), or with single-frame 405 nm pulses, **SBTubA4P**(0.6 μM), applied to full field of view at times indicated with dashed lines (“405,” violet) (n = 3 cells). Temporally precise onset and full field diffusional reversibility are shown (data related to **Movies S2-S3**). (a,c: mean ± SEM EB3 comet counts as normalized to the means of the first 5 timepoints; 405 nm ROIs applied at indicated times; for further details see Supporting Information).

We also modified this protocol to apply single frame 405 nm pulses to the whole field of view instead (**Fig 4c**, **Movie S2**). As expected, this confirmed the temporal precision of onset and the temporal reversibility of the spiking seen with the single-cell-resolved studies; though the video data are easier to interpret as they are even more visually impressive (**Movie S2**). Finally, we performed full-field-of-view imaging while applying a train of 47 frames of 487 nm pulses, to test whether **SBTubA4P** can be used orthogonally to GFP imaging wavelengths. Indeed, even at this high concentration (6 μM, 40×[I]_WC_), there was no induction of MT inhibition, so we concluded that **SBTubA4P** is indeed GFP-orthogonal (unlike azobenzene-based photopharmaceuticals)^16^, matching our design and expectations for GFP orthogonality by absorption cut-off (**Movie S3**; **Fig 4c**).

#### From cell culture, to MT photocontrol in 3D models, tissue explants, and animals

By now we had optimised the potency of metabolically stable, GFP-orthogonal photoswitchable **SBTub2M** and fully water-soluble analogue prodrug **SBTubA4P**, clarified the **SBTubs’** tubulin-specific cellular mechanism of action, and shown high-spatiotemporal-resolution photocontrol of MT dynamics in 2D cell culture. We were now primed to tackle the central photopharmacology research challenge, which has so far frustrated essentially all prior approaches: *in vivo* translation using systemic administration but local photocontrol, that clearly and usefully retains a defined cellular mechanism of action. We set out to test if **SBTubs’** performance features would permit this operation across a *range* of complex models from 3D culture, to 3D tissue explant, to two *in vivo* animal models: with spatiotemporally-localised illuminations photocontrolling the full sequence of their bioactivity (from suppressing MT polymerisation, to altering/depolymerising MT network structures, to reducing/stopping microtubule-dependent downstream processes), where appropriate with second-level resolution, but all with cellular precision.

### 2.5. SBTub photocontrol in 3D organoids enables spatially-targeted blockade of migration and mitosis in the long-term

We first tested **SBTubA4P** in 3D human mammary gland organoids grown from isolated patient tissue embedded in collagen gels. These resemble miniaturized and simplified organs with realistic micro-anatomy, and feature collective motility/invasion behaviour directing cells to migrate and proliferate to form ordered, branched structures.^37,38^ Controlling organoid morphology is a sought-after goal, which has been mostly interpreted as requiring spatiotemporal control of gene expression, for which optogenetic approaches have been suggested.^39^ Yet, the spatiotemporally-localized application of photochemical compounds offers an alternative, in which the possibility of *instantaneous* cellular response to stimulus is highly attractive for temporally-precise control. Based on the good performance of **SBTubA4P** in 2D cell culture, we assessed whether **SBTubA4P** can be controlled similarly precisely in a 3D organoid model, aiming to manipulate organoid development by locally interfering with the invasion of individual branches during the elongation phase^38^.

Though the actin cytoskeleton is the major driver of cell migration, MTs are integral to directional migration and leading edge stabilisation^21,40,41^ (as well as to proliferation): so we expected the single-cell motility as well as cell division rates could be locally affected. Therefore, we wished to test if repeated localized photoactivations of **SBTubA4P**(every 7 min) could be used over long timescales (>24 h) to inhibit outgrowth of light-targeted organoid branches, while leaving other branches of the same organoids to develop: so shaping and modulating cell migration and invasion with spatiotemporal control. This aim brings conceptual challenges for photopharmaceuticals: since all cells are exposed to the same drug concentration, and over long timescales the cumulative impacts of scattered photoactivation light, of diffused isomerised compound, and of imaging light itself, may build a spatially-nonspecific background pattern of bioactivity. Organoid morphology does not tolerate >0.1% organic cosolvent either, making full solubility important.

We first determined a suitable working concentration for **SBTubA4P** without spatially resolved activation, by monitoring organoid areas for morphological disruption under lit/dark conditions (**Fig S7a**). We determined an [I]_WC_ of 200 nM for preventing organoid growth over 24 h with UV-lit **SBTubA4P**, whereas organoids looked healthy with no antiproliferative / branch-retracting effects at 500 nM in the dark (**Fig 5a, Fig S7a-b**).

**Figure 5:**
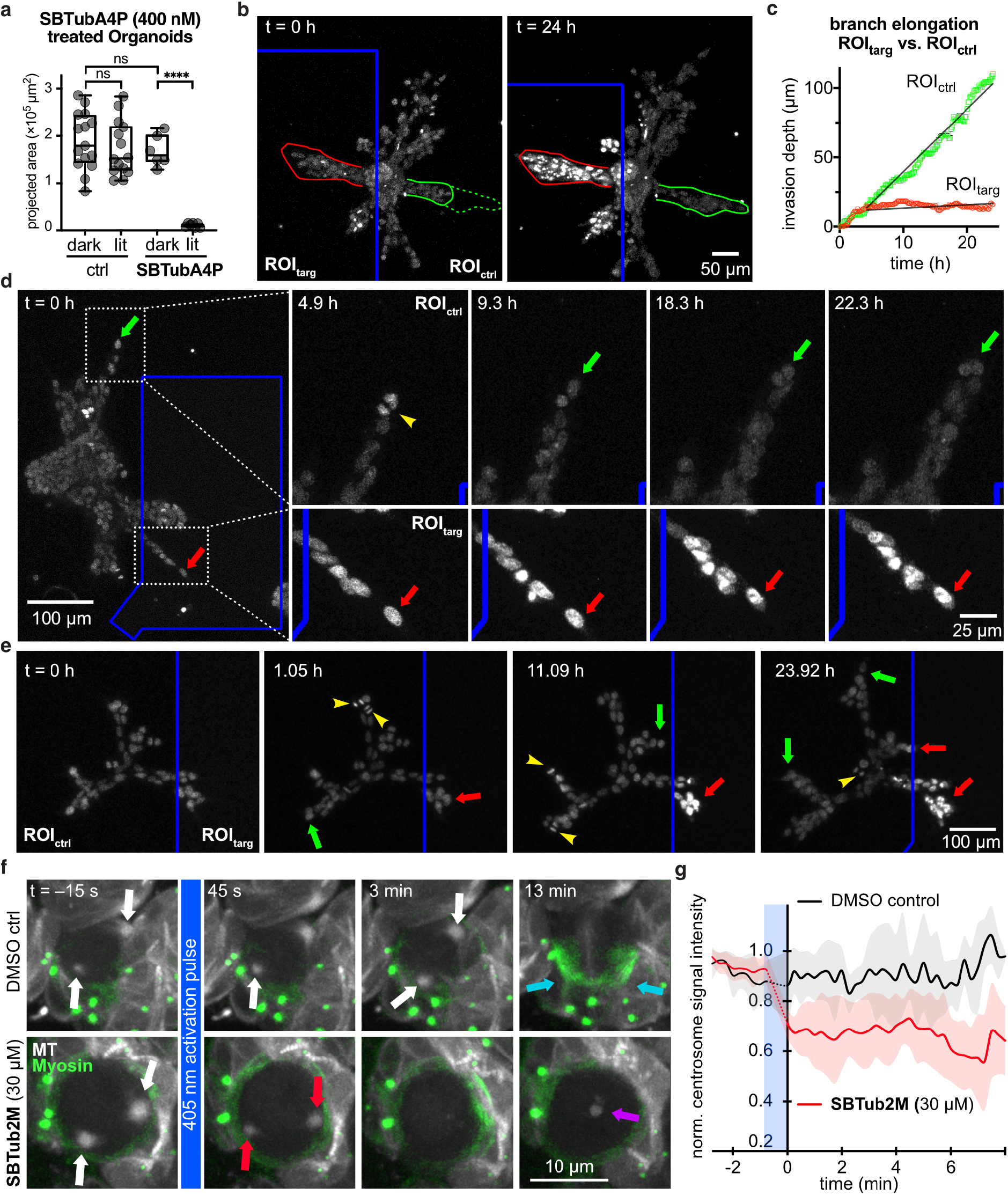
Spatiotemporal control over MT architecture, migration and mitosis in 3D culture and tissue explant. **(a)** 3D human mammary gland organoids embedded in collagen gels only have inhibited branch outgrowth when treated with both **SBTubA4P** and UV pulses. **(b)** Local applications of UV light to ROI_targ_ regions of **SBTubA4P**-treated organoids (blue boxed zone, one ca. 450 ms pulse per 7 min per z-stack) stops branch proliferation and outgrowth (red outline), while branches in untargeted ROI_ctrl_ regions develop dramatically (start: solid green line, final: dotted green line) (related to **Movie S4**). **(c)** Radial progress of branch tip fronts (directed and collective behaviour) in ROI_targ_ and ROI_ctrl_ regions. **(d)** Still image timecourse, zoomed on a branch tip in the ROI_ctrl_ (blue outline) region, showing cell proliferation (yellow arrowhead) and matrix invasion (one representative of the migrating cells is tracked over time with green arrows); while branch tip of ROI_targ_ region has static non-proliferating cells and even slight branch retraction (red arrows) (data related to **Movie S5**). **(e)** branch progression and proliferation are unimpeded and continuous in ROI_ctrl_ regions, while ROI_targ_ regions are static and branches growing into the ROI_targ_ stop their growth (colour code as in **(e)**, data related to **Movie S6**). (**a-e**: cell location in organoids tracked with siRDNA nuclear stain imaged at 647 nm). **(f)** Whole field of view 405 nm photoactivation of **SBTub2M**-treated intact 3D brain explants of larval *D. melanogaster* (bottom row) causes neuroblast centrosomes (red arrows) to rapidly shrink in size and signal intensity (45 s and 3 min) and prevents the cell from progressing through division (13 min). Some MT signal accumulates at mid-cell at later time points (purple arrow) (data related to **Movie S8**). In DMSO-only controls (top row), centrosome integrity (white arrows, 45 s and 3 min) and progression through the cell cycle (13 min) are unaffected, indicated by myosin accumulation at the cleavage furrow (cyan arrows) (data related to **Movie S9**). (MTs in white (Jupiter::mCherry imaged at 561 nm), myosin in green (Squash::GFP imaged at 488 nm.) **(g)** Relative mCherry fluorescence intensity of centrosomal ROIs in **SBTub2M**-treated prophase neuroblasts (red) after activation at 405 nm drops notably during the approx. 45 s activation period (blue box) as compared to the DMSO control prophase neuroblasts (black). Signal intensities are shown as the proportion of the per-cell maximum pre-activation signal intensity. (Shading indicates ±1 standard deviation, 1-2 centrosomes quantified from a total of 5 (DMSO) or 5 (**SBTub2M**) neuroblasts from 3 different animals). For details see Supporting Information.

Then, we applied localised UV ROI illuminations to selected organoid branches (ROI_targ_) every 7 min, comparing to non-UV-illuminated internal control branches (ROI_ctrl_). This allowed us to noninvasively block cell migration/invasion and proliferation with striking spatial resolution and longterm persistance. With **SBTubA4P**, development of ROI_targ_ branches was totally blocked over > 1 day (**Fig 5b-d**, **Movie S4-S5**); while no-compound controls showed no photo-inhibition of branch development (**Movie S7**, **Fig S7c**). Control branches displayed high motility and matrix invasion (green outline, **Fig 5b**) and their cells freely proliferated (yellow arrowhead, **Fig 5d**), resulting in considerable branch development. Out-directed motion of the collective branch fronts was continuous in ROI_ctrl_ regions (ca. 4.5 μm/h), but was almost completely stopped in ROI_targ_ (ca. 0.2 μm/h; **Fig 5c**). Overall development of branches, only entering ROI_ctrl_ areas, was however clearly visible without any statistical analysis (**Fig 5e, Movie S6**). Thus **SBTubA4P** can be used in 3D matrix cell culture settings to noninvasively control cell motility, invasion and proliferation, allowing photopatterning of branch growth and organoid development down to the spatial scale of individual cells.

### 2.6. SBTub photocontrol in intact 3D tissue explants allows temporally-precise MT depolymerisation and mitotic control in the short-term

We next tested **SBTub** performance and tubulin-specific mechanism of action when directed against the more complex 3D environment of live intact brain lobes of early third instar larval *Drosophila melanogaster* (fruit fly). Larvae are too motile for long-term imaging and the larval cuticle is largely impermeable, so neurodevelopment studies explant the whole brain. As the explant tolerates cosolvent, we took the opportunity to use **SBTub2M** in these assays to test the broader applicability of the **SBTub** design. Both the whole-organ and explant aspects bring significant challenges. (1) Compounds must permeate through two glia cell layers to reach mitotically active neural stem cells (neuroblasts). This forces the use of high bath concentrations and potent compounds: however, since surface and surrounding cells are exposed to far higher concentrations than central cells, only potent compounds with *extremely high photoswitchability of bioactivity* (“FDR”; discussed in^19^) can be used: otherwise outer cells die, and morphology and physiology are lost. Indeed, we could use high concentrations of high-FDR **SBTub2M**(30 μM) in brain explants without noticeable toxicity. (2) Using multiple fluorescent protein labels for multiplexed imaging is a typical requirement to achieve useful readouts in biology, but can block chemical photoswitch applications. The most common long-wavelength fluorescent proteins for animal work are excited at 561 nm, which forces the use of GFP (488 nm) or YFP (514 nm) fusions as the next-longest-wavelength markers. For example, to image both MTs and cellular structural elements, we used animals expressing the microtubule-binding protein mCherry::jupiter^42,43^, and cortical structure marker sqh::GFP (spaghetti squash, the regulatory light chain of the non-muscle type 2 myosin, fused to eGFP^44^). With hemithioindigo or azobenzene reagents, such two-channel FP imaging would isomerise the photoswitch throughout the sample due to photoresponse at ≤530 nm (Fig 2b), so destroying spatiotemporal specificity in the study zone. In contrast, the non-response of the GFP-orthogonal SBT to 488 nm imaging avoids any photoisomerisation during typical two-channel FP imaging, allowing precise temporal control of activation in our experiments.

We transferred freshly-dissected brain explants into 30 μM **SBTub2M** and started imaging after 30 minutes loading (**Fig S8**). We imaged in both mCherry (ex 561 nm) “red channel” and GFP (ex 488 nm) “green channel” for 15 minutes to establish baseline, then photoactivated **SBTub2M** throughout the imaging stack volume with 405 nm. Photoactivated **SBTub2M** depolymerised centrosome microtubules within 60 s (**Fig 5f** and **Movie S8-9**, MTs shown in white). To control for target specificity, we also used mCherry::tubulin^45^ for imaging MTs, and observed similar behavior (**Movie S10-11**). As microtubules are rapidly nucleated in prophase centrosomes, we quantified the loss of centrosomal fluorescence signal after activation as a highly conservative estimate of centrosome MT depolymerisation. We saw dramatic, temporally-resolved signal reduction at the approx. 45 s activation period, while **SBTub** controls were unaffected (**Fig 5g**). We used the second FP channel to image Sqh::GFP, a marker of the cell actomyosin cortex, which plays a key role in neuroblast asymmetric division^46,47^. Normally dividing neuroblasts accumulate Sqh::GFP at the cell cleavage furrow during anaphase. Neuroblasts in which **SBTub2M** was photoactivated retain uniform cortical Myosin, indicating mitotic arrest in the absence of mitotic spindles^46^ (cyan arrows, **Fig 5f**).

Previous short-term results imaging EBs at low **SBTub** concentrations in 2D cell culture had illustrated only its capacity to spatiotemporally block MT polymerisation (**Fig 4**). Now, these useful results in the intact brain underlined that **SBTub2M** maintains its *Z*-isomer-specific, MT-depolymerising mechanism of action in live tissue explant, casting **SBTubs** as flexible and powerful tools for cytoskeleton photomanipulation in complex 3D settings.

### 2.7. SBTub photocontrol in live animals enables targeted blockade of embryonic development and cell-precise temporally-reversible inhibition of MT dynamics

Encouraged by performance in 3D models, we evaluated using **SBTubs** for *in vivo* photocontrol. First we studied the effects of *in situ* photoactivations of **SBTubA4P** on the development of *Xenopus tropicalis* clawed frog embryos. During the initial 48 h of development, embryos normally transition over many division cycles from cell spheres through the blastula stage through to multiorgan tadpoles (**Fig S9a**). Initially we tested the effects of **SBTubA4P** in the earliest stages of development, just after embryonic divisions had started, by treating 2-cell stage embryos with *E*-**SBTubA4P**(1 hour loading, then medium exchange and optional *in situ* embryo-localised 410 nm photoactivation pulse during washout; note that this *transient* exposure to the **SBTub** parallels what could be expected for e.g. systemic *i.v.* administration in mammalian models – further discussion in Supporting Information). Embryos only failed to develop morphologically over the subsequent 2 days if they had received the 410 nm photoactivation; embryos without photoactivation developed normally (**Fig 6a-b**). We also tested interfering with development at the later blastula stage (>64 cells) by a similar protocol. While subsequent morphological development was normal under lit and dark conditions (**Fig S9b-c**), sensorimotor responses to mechanical stimulation^48^ were suppressed by lit **SBTubA4P** only (**Fig S9e**, **Movie S12**). Interestingly, the tubulin-inhibiting azobenzene photoswitch **PST-1P**(**Fig S9e**) light-*independently* suppresses sensorimotor responses. We believe the SBT’s success may reflect its greater metabolic robustness (see Supporting Information); and at any rate, it indicated that **SBTubs** are suitable for *in vivo* use.

**Figure 6:**
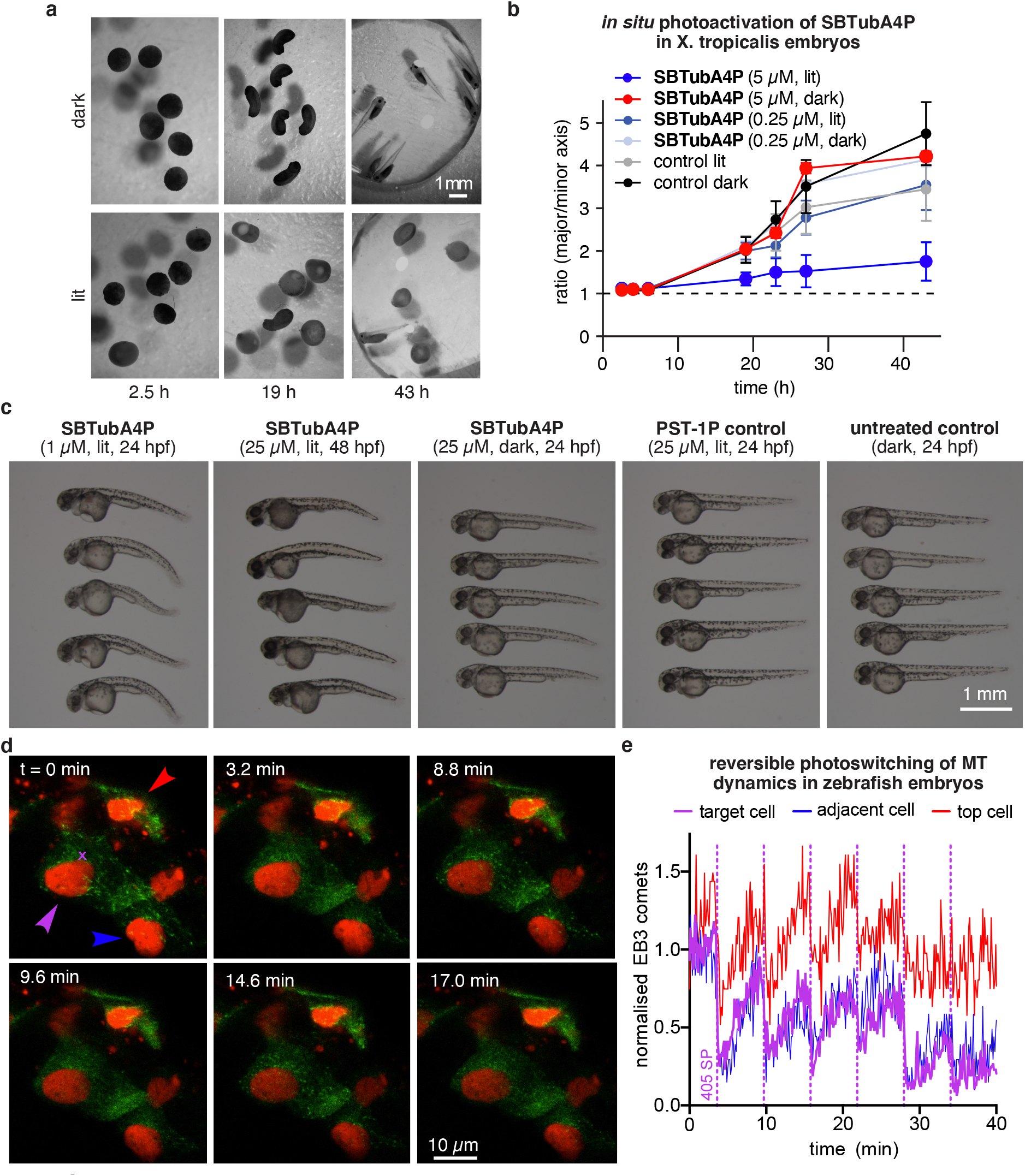
Photoinhibition of *X. tropicalis* **development, and** *in vivo* **photocontrol of MT dynamics in** *D. rerio*. **(a-b)** *Xenopus* embryos incubated with compounds for 1 h at the 2-cell-stage, before medium exchange optionally with 410 nm photoactivation. Embryos show irreversible development inhibition by *in situ-*formed *Z-***SBTubA4** in lit conditions, but had no effects in the dark or with low concentration of **SBTubA4P** (**a**: **SBTubA4P** at 5 μM; **b**: development quantified by the ratio of major to minor embryo axis lengths, 6 embryos per conditions, mean ± SEM). **(c)** Development of *D. rerio* treated at the indicated stages for 24 h with **SBTubA4P** or control compounds under pulsed lit conditions (1 s/5 min) or dark. **SBTubA4P** (1 or 25 μM) causes morphological abnormalities only in the lit state, showing it remains effective *in vivo*. **(d-e)** reversible modulation of MT dynamics in 48 hpf zebrafish embryo. (EB3-GFP in green, histone H2B in red). (data related to **Movies S12-16**; see Supporting Information).

Finally, we switched to highly spatiotemporally-resolved *in vivo* MT-imaging studies that would test **SBTubA4P**’s mechanism of action in the zebrafish *Danio rerio*, when systemically applied and maintained in the bath medium.

We first determined useful working concentrations in zebrafish, incubating 24 and 48 hpf (hours post fertilisation) embryos in **SBTubA4P** under lit and dark conditions. While zebrafish morphology remained unaltered in all dark **SBTub** treatments, 24 hpf embryos treated with lit **SBTub** showed major morphological changes even down to 1 μM, whereas more developed 48 hpf embryos showed similar morphological changes only at higher **SBTubA4P** concentrations e.g. 25 μM (**Fig 6c, Fig S10**). Again, we compared these effects to those of azobenzene reagent **PST-1P**, now observing a dramatic difference: even 25 μM lit **PST-1P** did not interfere with development at either the 24 hpf or 48 hpf stage (**Fig 6c, Fig S10**). This argues still more conclusively than **Fig 6b**, that the SBTub scaffold is uniquely suitable for light-controlled biological effects, compared to the previously known azobenzene scaffold. Lastly, we checked the lower-potency soluble prodrug **SBTub3P** in the same assay; matching expectations, it caused only weak changes at 24 hpf and no visible changes at 48 hpf (**Fig S10**), showing the necessity of the potency optimisations we performed in this study.

Aiming to test the MT-modulating effects of **SBTubs** in a challenging live animal system, we therefore decided to proceed with 48 hpf zebrafish embryos, and an **SBTubA4P** working concentration of 25 μM. We took 48 hpf embryos coexpressing EB3-GFP and histone H2B-mRFP as a nuclear marker,^49,50^ loaded them with 25 μM *E-***SBTubA4P** for 4 h, then washed and embedded them in agarose. Imaging at 488 nm caused no suppression of EB3 comets, confirming **SBTubA4P**’s GFP orthogonality *in vivo*. However, photoactivation with the 405 nm laser at a single point caused EB3 comets to vanish rapidly in cells around the targeted region, recovering over ca. 10 minutes. Cells further from the targeted region were predictably less inhibited than those with direct contact to the photoactivation region. The photoactivation-recovery cycle could be repeated multiple times during imaging (**Fig 6d-e, Movie S13-S16**). Not only microtubule polymerisation dynamics, but also mitotic progression, could be stopped by spatiotemporally-localised **SBTubA4** photoactivation *in vivo* (**Movie S16**). These experiments confirm that the SBT scaffold in general is viable for light-triggered *in vivo* studies, and that **SBTubA4P** when applied *in vivo* retains its mechanism of action as a potent, light-dependent MT inhibitor with excellent spatial specificity and satisfying temporal reversibility.

## 3. Conclusions

Noninvasive optical tools to modulate microtubule dynamics, structure and function with high precision, offer unique potential in the many fields of biology impacted by the spatiotemporally-resolved processes that MTs support: such as cargo transport, cell motility, cell division, development and neuroscience. Photopharmaceutical chemical reagents are conceptually elegant tools, in that they can be rapidly transitioned across models and settings, and they can be rationally designed for photoresponse patterns that interfere minimally with imaging while maximising optical response to a chosen photoactivation wavelength. Locally-applied photopharmaceuticals, particularly intraocularly-applied reagents for action potential control in the retina, have made great progress in adult mammalian disease models.^51,52^ However, reaching the true potential of photopharmacology – achieving precisely-targeted *in vivo* applications by localised *in situ* photoactivations following systemic administration – remains an unsolved challenge. This would require combining high photoswitchability of bioactivity, high potency, metabolic robustness, aqueous solubility, and imaging orthogonality. Indeed, very few systemic *in vivo* applications of photopharmaceuticals have been made. As none of those have tested a defined mechanism of drug action, nor explored the optical-scale spatiotemporally-resolved targeting which is the only benefit of photopharmaceutical approaches as compared to conventional drugs,^53–55^ the ability of photopharmacology to contribute useful systemically-applicable reagents for organism studies has remained unclear.

In this work we develop highly light-specific tubulin polymerisation inhibitors with unprecedented applicability from 2D cell culture, to 3D culture and whole-organ explant, to systemic *in vivo* administration with local photoactivation. Realising that the metabolic robustness and imaging orthogonality of a photopharmaceutical largely depend on its photoswitch scaffold, we consciously avoided the typical azobenzene photoswitch scaffold. Using the recently-developed SBT photoswitch, we created a panel of twenty **SBTubs**. We optimised potency and photoswitchability of bioactivity in lead **SBTub2M**; and we solubilised another lead to create **SBTubA4P** which does not require organic cosolvents. **SBTubs** are efficiently photoactivated with the common 405 nm laser, but their sharp absorption cutoff leaves GFP, YFP, and RFP channels free for multiplexed imaging of fusion protein markers without risking compound photoactivation. This is a highly desirable feature for areas of research where photopharmacology’s optical precision can best contribute unique solutions on the cellular spatial scale (although we note that this runs against the goal of “photoswitch redshifting” that is near-universally cited by chemist photopharmaceutical designers). We cross-validated the **SBTubs’** molecular mechanism of action in cell-free, cellular and *in vivo* settings. **SBTubs** light-dependently interfere with mitosis in cell culture, and depolymerise mitotic spindles and ultimately block development *in vivo*; they can be optically patterned to control motility and branch development in 3D organoid cultures; and their photocontrol allows rapid-response, cell-specific inhibition of microtubule dynamics in cell culture and *in vivo*.

These consistent results across a range of models at different scales of time, length, and biological complexity, recommend the **SBTubs** as excellent and unique general-purpose tools for optically manipulating microtubule dynamics, microtubule structure, and microtubule-dependent processes with high spatiotemporal precision.

The proof-of-concept biological performance of the **SBTubs** has been very satisfying. We believe that the most valuable improvement to this system will now be to extend the temporal reversibility of inhibition (seen in 2D cell culture by diffusion to the medium with ca. 20 s halftime, **Fig 4**) to whole organ / whole animal settings. In these settings, diffusion is slower to achieve reversibility (ca. 10 min), so we seek techniques for *in situ* bidirectional isomerisation of **SBTubs** in our ongoing research. We believe that bidirectional photoswitching may be difficult within the biologically-compatible wavelength range^13^. However, accelerating thermal relaxation to the minute scale, which is probably the most appropriate scale for 3D / *in vivo* applications of interest, may be feasible, and efforts are underway.

In conclusion, the **SBTubs** are excellent photoswitchable microtubule-depolymerising reagents for use in cell culture, 3D culture, small explant, and early-stage animals. Their potency, flexibility and ease of use recommend them for high-spatiotemporal-precision research across cytoskeleton biology; particularly, we feel, for cell-specific applications to motility and development, but they will also be of great interest in cargo transport, biophysics, cell polarity, neurodegeneration, and cell division. Lastly, we expect that by supporting conceptual innovations in photoswitch scaffold chemistry and rational photopharmaceutical design, and particularly by starting to unlock the applications promise of photopharmacology for globally-administered, locally-targeted *in vivo* use, this **SBTub** research represents a promising advance for high-performance photopharmacology against other protein targets in general, beyond their immediate impact on microtubule biology.

## Funding Statement

This research was supported by funds from the German Research Foundation (DFG: Emmy Noether grant number 400324123 to O.T.-S.; SFB 1032 number 201269156 project B09 to O.T.-S. and project A10 to A.R.B.; SFB TRR 152 number 239283807 project P24 to O.T.-S.; and SPP 1926 number 426018126 project XVIII to O.T.-S.). J.C.M.M. acknowledges support from an EMBO Long Term Fellowship. J.T.-S. acknowledges support from a Joachim Herz Foundation Stipend. A.R.B. gratefully acknowledges the financial support of the European Research Council (ERC) through the funding of the grant Principles of Integrin Mechanics and Adhesion (PoINT). A.L. acknowledges funding from the Spanish government (Ministerio de Ciencia e Innovación), grant RTI2018-096948-B-100 (A.L.), co-funded by the European Regional Development Fund (ERDF). M.D.acknowledges funding from the Austrian Research Promotion Agency (FFG) project 7940628 (Danio4Can). C.C.C. is supported by NIH grant 1R01GM126029.

## Acknowledgement

We are grateful to Henrietta Lacks, now deceased, and to her surviving family members for their contributions to biomedical research. We thank Monique Preusse for early cell viability testing, and Rebekkah Hammar for performing the tubulin polymerisation assay. We thank Christian Gabka from the Nymphenburg Clinic for Plastic and Aesthetic Surgery, Munich 80637, Germany for providing primary human mammary gland tissue.

## Author contributions

L.G. performed synthesis, photocharacterisation, and long-term cellular studies, coordinated data assembly, and wrote the manuscript. J.C.M.M. performed live cell EB3 imaging during photoswitching, and assembled and quantified EB imaging data. A.V. performed zebrafish assays. I.E.R. performed organoid assays. C.H. performed cell viability assays, immunofluorescence staining, and cell cycle analysis. J.A.T. performed *Drosophila* experiments. M.W. performed structural biology. C.D.V. and B.T. performed clawed frog assays. J.S. and M.O.S. supervised structural biology. C.C.C. supervised *Drosophila* experiments. A.L. supervised clawed frog assays. A.R.B. supervised organoid assays. M.D. supervised zebrafish assays. A.A. supervised EB3 imaging. J.T.-S. performed cell cycle analysis, immunofluorescence microscopy, coordinated data assembly and supervised all other cell biology. O.T.-S. designed the concept and experiments, supervised all other experiments, coordinated data assembly and wrote the manuscript with input from all authors.

## Declaration of interests

The authors declare no competing interests.

## Supplementary Information

1. **PDF** containing (i) chemical synthesis and NMR spectra (ii) photocharacterisation in vitro (iii) biochemistry (iv) biological methods and data (v) X-Ray crystallography data.
2. Supplementary Movies

***Movies S1-S3**: Timestamps in mm:ss, HeLa cells transiently transfected with EB3-tdTomato imaged with 561 nm. Purple dots in **Movies S1-S2** indicate 405 nm illumination pulses; blue dots in **Movie S3** indicate 487 nm illumination pulses.*
**Movie S1, temporally precise and cell-precise inhibitions of cellular MT polymerisation dynamics by photoactivations of E-SBTubA4P at 405 nm, related to**
**Figure 4a-b:** no inhibition of MT polymerisation dynamics unless **SBTubA4P** is present (6 μM) and then 405 nm ROI illumination pulses (indicated by purple dot) are applied to the cell of interest (indicated by purple arrow) with minor impact on non-targeted neighbouring cell (indicated by white arrow) (EB3-tdTomato imaged at 561 nm).
**Movie S2, temporally precise full-field-of-view inhibitions of cellular MT polymerisation dynamics by photoactivations of***E-***SBTubA4P at 405 nm, related to Figure 4c:** no inhibition of MT polymerisation dynamics unless **SBTubA4P** is present (6 μM) and then 405 nm field of view (FOV) illumination pulses (indicated by purple dot) are applied (EB3-tdTomato imaged at 561 nm).
**Movie S3, no inhibition of cellular MT polymerisation dynamics by illumination of***E-***SBTubA4P at 487 nm, related to Figure 4c:** cells treated with *E-***SBTubA4P**(6 μM) then imaged while 487 nm field of view (FOV) illumination pulses (indicated by blue dot) are applied show no inhibition.
***Movies S4-S7**: primary human mammary gland organoids, cell nuclei stained with siRDNA and imaged at 674 nm. Blue rectangles in Movies **S4-S7** indicate the 405 nm ROI_targ_.*
**Movie S4-S6, SBTubA4P blocks branch development light-dependently and with spatiotemporal precision, related to Figure 5b-e:** organoids treated with **SBTubA4P**(200 nM) were imaged for 4 h without 405 nm photoactivation (countdown) showing normal proliferation and mitosis, then pulsed 405 nm illumination was begun within the targeted area ROI_targ_ at time t = 0 (1 pulse per 7 min), locally stopping motility and proliferation.
**Movie S7, no-compound control shows no photoinhibition of branch development in both ROI_targ_ and ROI_ctrl_, related to Supporting Figure S7:** organoid not treated with any compound was imaged for 4 h without 405 nm photoactivation pulses, then ROI_targ_ illumination with 405 nm pulses was performed as in **Movies S4-S6**. Note that the 405 nm-triggered increase in nuclear fluorescence intensity in the ROI_targ_ (seen in **Movies S4-S6**) is however not accompanied by stoppage of motility or proliferation (unlike in **Movies S4-S6**).
***Movies S8-S11:** Timestamps in mm:ss; explanted drosophila brain lobe with focus centred on prophase neuroblast; 2% DMSO final concentration; imaged at 561 nm (mCherry; microtubules; white) and 488 nm (EGFP; spaghetti squash; cortical structure); imaged for 15 min (countdown) prior to 405 nm activation (from t = 0; blue frames; 20 μm stack with 1 μm z-spacing, 2 s/slice at 40% laser power, approx. 45 s total) then for 30 min post-activation.*
**Movie S8, temporally precise depolymerization of the mitotic spindle in a prophase Drosophila neuroblast by photoactivation of***E-***SBTub2M (30 μM) at 405 nm, related to Figure 5f-g.** Field of view is centred on a prophase neuroblast. White = mCherry::Jupiter; green = Squash::EGFP.
**Movie S9, DMSO-only control to Movie S8 shows prophase Drosophila neuroblast undergoing normal mitosis after 405 nm illumination, related to figure 5f-g.** Field of view is centred on a prophase neuroblast. White = mCherry::Jupiter; green = Squash::EGFP.
**Movie S10, temporally precise depolymerization of the mitotic spindle in a prophase Drosophila neuroblast by photoactivation of***E-***SBTub2M (30 μM) at 405 nm.** Field of view is centred on a prophase neuroblast. Compared to Movies S9-S10, a different microtubule marker is used (White = mCherry::Tubulin).
**Movie S11, DMSO-only control to Movie S10 shows prophase Drosophila neuroblast undergoing normal mitosis after 405 nm illumination.** Field of view is centred on a prophase neuroblast. White = mCherry::Tubulin.
**Movie S12, related to Figure 6a-b and Figure S9: Reaction of hatchling embryos to mechanical stimulus depends on temporally precise application of***Z-***SBTubA4 during prior development.** Healthy larvae initiate a movement upon mechanical stimulation with the tip of a glass pipette. Comparison of control conditions (dark, water), to 25 μM **SBTubA4P** post-lit and dark.
***Movies S13-S16:** Timestamps in mm:ss; wt AB* zebrafish embryos microinjected with pSK_H2B-mRFP:5xUAS:EB3-GFP and pCS_KalTA4 plasmid DNA at the single-cell stage; embryos treated with **SBTubA4P** from 26-30 hpf then embedded and imaged at 561 nm (RFP; DNA; red) and 488 nm (GFP; microtubule dynamics; green). Photoactivations (405 nm, 7 s) as bleachpoints at the targeted area are indicated with a purple X at the time of application.*
**Movie S13-S14, Photoactivation of SBTubA4P (25 μM) in a live zebrafish embryo shows temporally reversible inhibitions of EB3 dynamics over several cycles, related to Figure 6d-e:** inhibition of MT polymerisation dynamics at each time when **SBTubA4P**(25 μM) is photoactivated, as repeated over several cycles. Movies show the same field of view, with Movie S13 showing DNA (histones; red) and microtubule plus tips (EB3; green), while Movie S14 shows only the EB3 channel (in white) as can be used for quantification of EB3 dynamics (photoactivations in this movie are temporally indicated by the purple circle).
**Movie S15, Inhibition of MT polymerisation dynamics when SBTubA4P (25 μM) is photoactivated.** Assay performed as in Movie S13-S14, showing similar effects.
**Movie S16, Photoactivation of SBTubA4P (25 μM) stops both EB3 dynamics and cell division in the developing embryo.** Assay performed as in Movie S13-S14, showing similar effects; a mitotic cell is also to be seen in the field of view (lower right); photoactivation of **SBTubA4P** stops its EB3 dynamics and after approx. 2.5 minutes the spindle starts to lose integrity; division is not productively continued during the rest of the movie.

## Notes

### Competing Interest Statement

The authors have declared no competing interest.

