## Supporting Information for "In vivo photocontrol of microtubule dynamics and integrity, migration and mitosis, by the potent GFP-imaging-compatible photoswitchable reagents SBTubA4P and SBTub2M"

[1] Department of Pharmacy, Ludwig-Maximilians University of Munich; Munich 81377, Germany
[2] Cell Biology, Department of Biology, Faculty of Science, Utrecht University, Utrecht 3584, Netherlands
[3] St. Anna Kinderkrebsforschung Children's Cancer Research Institute, Wien 1090, Austria
[4] Physics Department, Technical University of Munich and Center for Protein Assemblies; Garching 85747, Germany
[5] Laboratory of Biomolecular Research, Division of Biology and Chemistry, Paul Scherrer Institut, Villigen 5232, Switzerland
[6] Laboratory of Neurobiology, Department of Pathology and Experimental Therapy, Institute of Neurosciences, University of Barcelona; L'Hospitalet de Llobregat, Barcelona 08907, Spain
[7] Bellvitge Biomedical Research Institute (IDIBELL), L'Hospitalet de Llobregat, Barcelona 08907, Spain
[8] Department of Biology, University of Washington; Seattle 98195, USA
[9] Biozentrum, University of Basel, Basel 4056, Switzerland
[10] Zebrafish Platform Austria for Preclinical Drug Screening, Vienna 1090, Austria
[11] Lead Contact

ORCID: O.T.-S. 0000-0003-3981-651X

***Keywords**: microtubule dynamics; cytoskeleton; photopharmacology; tubulin polymerisation inhibitor; optical control; colchicine; antimitotic; cell division; cell migration; development; photoswitch.*

**Table of Contents**

**Part A: Chemical Synthesis ... 3**

Conventions ... 3

General procedures ... 4

Experimental data of SBTubs ... 6

Experimental data of synthetic building blocks ... 17

**Part B: Photocharacterisation *in vitro* ... 24**

Spectrophotometry equipment ... 24

Photoisomerisation: further discussion ... 26

Photostability ... 26

Spontaneous relaxation of SBTubA4 ... 28

**Part C: Biological Data ... 29**

Tubulin polymerisation ... 29

Cell assay methods ... 29

General cell culture ... 29

Resazurin antiproliferation assay ... 30

Cell cycle analysis ... 32

Immunofluorescence staining ... 33

Live cell imaging ... 33

Quantification and statistical analysis ... 34

3D cell culture ... 34

Tissue and Animal Assays ... 37

Drosophilia melanogaster ... 37

Xenopus tropicalis ... 38

Danio rerio ... 41

**Part D: Protein Crystallisation ... 43**

Protein crystallisation materials and methods ... 43

**Part E: NMR Spectra ... 46**

**Part F: Bibliography ... 78**

#### Part A: Chemical Synthesis

##### *Conventions*

Abbreviations: The following abbreviations are used: Hex – distilled isohexanes, EA – ethyl acetate, DCM – dichloromethane, Et – ethyl, Ac – acetyl, Me – methyl, MeCN – acetonitrile, DMSO – dimethylsulfoxide, PBS – phosphate buffered saline. TFA – trifluoroacetic acid, DMAP – 4-(dimethylamino)pyridine, TEA – triethylamine.

Safety Hazards: no unexpected or unusually high safety hazards were encountered.

Reagents and Conditions: Unless stated otherwise, (1) all reactions and characterisations were performed with unpurified, undried, non-degassed solvents and reagents, used as obtained, under closed air atmosphere without special precautions; (2) "hexane" used for chromatography was distilled from commercial crude isohexane fraction by rotary evaporation; (3) "column" and "chromatography" refer to manual flash column chromatography on Merck silica gel Si-60 (40–63 µm); (4) MPLC flash column chromatography refers to purification on a Biotage Isolera Spektra, using prepacked silica cartridges from Biotage; (5) procedures and yields are unoptimized; (6) yields refer to isolated chromatographically and spectroscopically pure materials, corrected for residual solvent content; (7) all eluent and solvent mixtures are given as volume ratios unless otherwise specified, thus "1:1 Hex:EA" indicates a 1:1 (v/v) mixture of hexanes and ethyl acetate; (8) chromatography eluents e.g. "0→25% EA:Hex" indicate a linear gradient of eluent composition.

Thin-layer chromatography (TLC) was run on 0.25 mm Merck silica gel plates (60, F-254), typically with Hex:EA eluents, except where indicated. UV light (254 nm) was used as a visualizing agent, with cross-checking by 365 nm UV lamp. TLC characterizations are abbreviated as $\mathbf{R}_f$ = 0.64 (EA:Hex = 1:1).

NMR: Standard NMR characterization was by $^{1}$H- and $^{13}$C-NMR spectra on a Bruker Ascend 400 (400 **MHz** & 100 **MHz** for $^{1}$H and $^{13}$C respectively) or a Bruker Ascend 500 (500 **MHz** & 125 **MHz** for $^{1}$H and $^{13}$C, respectively). Known compounds were checked against literature data and their spectral analysis is not detailed unless necessary. Chemical shifts (δ) are reported in ppm calibrated to residual non-perdeuterated solvent as an internal reference[1]. Peak descriptions singlet (s), doublet (d), triplet (t), quartet (q) and multiplet (m).

Analytical HPLC and Mass Spectra: Analytical HPLC-MS measurements were performed on an Agilent 1100 SL coupled HPLC-MS system with (a) a binary pump to deliver $H_2O$:MeCN eluent mixtures containing 0.1% formic acid at a 0.4 mL/min flow rate, (b) Thermo Scientific

Hypersil GOLD™ C18 column (1.9 µm; 3 × 50 mm) maintained at 25°C, whereby the solvent front eluted at $t_{ret}$ = 0.5 min, (c) an Agilent 1100 series diode array detector used to acquire peak spectra of separated compounds/isomers in the range 200-550 nm after manually baselining across each elution peak of interest to correct for eluent composition effects, (d) a Bruker Daltonics HCT-Ultra mass spectrometer used in ESI mode at unit mass resolution. Run conditions were a linear gradient of $H_2O$:MeCN eluent composition from the starting ratio through to 10:90, applied during the separation phase (first 5 min), then 0:100 maintained until all peaks of interest had been observed (typically 2 min more); the column was equilibrated with the $H_2O$:MeCN eluent mixture for 2 minutes before each run. Unless stated otherwise, all reported peaks in the positive mode were $[M+H]^+$ peaks, and all observed peaks in the negative mode were $[M-H]^-$ peaks. HRMS was carried out by the Zentrale Analytik of the LMU Munich using ESI or EI ionisation as specified. LRMS was carried out on an expression CMS by Advion with either APCI or ESI as ionization source.

#### *General procedures*

##### General procedure 1: Synthesis of SBT *via* Aldol Condensation

Equimolar amounts of the 2-methylbenzothiazole (1 eq) and the benzaldehyde (1 eq) is dissolved in DMSO (1 mL/0.1 mmol) and NaOMe (6 M in MeOH, 2 eq) is added. The reaction mixture is stirred at room temperature overnight. $H_2O$ (15 mL) and sat. aq. $NH_4Cl$ (15 mL) is added and the mixture is extracted with ethyl acetate (3 × 20 mL). The combined organic layers are dried over $MgSO_4$ and concentrated *in vacuo*. The crude product is purified by normal phase MPLC.

##### General procedure 2: Synthesis of 2-methylbenzothiazoles *via* Jacobsen cyclization

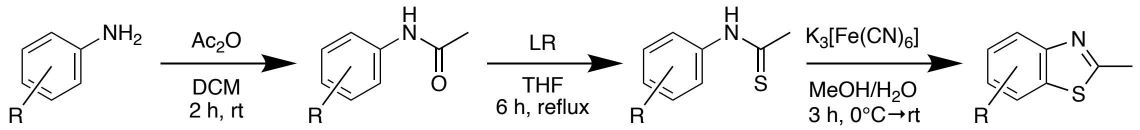

Acetic anhydride (1.2 eq) is slowly added to a solution of the aniline (1 eq) in dry DCM. The reaction mixture is stirred for 2 h at room temperature and reaction progress is monitored by TLC. After completion, the mixture is quenched with sat. aq. $NaHCO_3$ (25 mL) and extracted with DCM. The combined organic layers are dried over $MgSO_4$, filtrated and the solvent is removed under reduced pressure. The unpurified phenyl acetamide is dissolved in THF and Lawesson's reagent (0.7 eq) is added. The reaction mixture is stirred for 6 h at reflux. After the reaction is finished, the solution is cooled down to room temperature and water is added. The suspension is extracted with EA and washed with brine. The combined organic layers are dried over $Na_2SO_4$, filtrated and the solvent is removed under reduced pressure. The crude is filtrated through a silica plug and used without further purification. The crude phenyl thioamide is dissolved in MeOH and aq. NaOH (2 M, 2 eq.) is added. After stirring for 1 h at room temperature, MeOH is removed under reduced pressure and a solution of $K_3[Fe(CN)_6]$ (1.2 eq) in $H_2O$ is added at 0°C and warmed up to room temperature. Stirring is continued for another 3 h at room temperature. The reaction mixture is extracted with EA and washed with brine. The combined organic layers are dried over $Na_2SO_4$, filtrated and concentrated *in vacuo*. The crude is purified by Kugelrohr distillation or normal phase MPLC to give the desired 2-methylbenzothiazole.

**General procedure 3: Synthesis of 2-methylbenzothiazoles *via* Cu-catalysis**

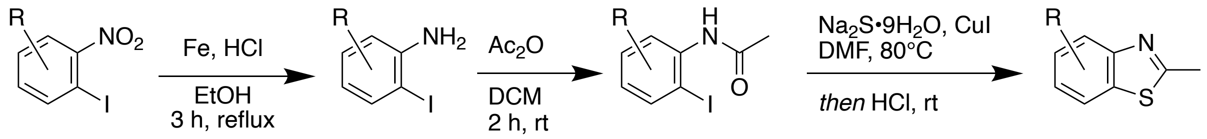

Iron powder (6.2 eq) and conc. HCl (1.2 eq) is added to a solution of *o*-iodo nitrotoluene in ethanol. After 3 h of reflux, the mixture is cooled to room temperature and $Na_2CO_3$ is added portionwise until gas evolution ceased. After filtration over Celite, the filtrate is extracted with EA and the combined organic fractions are washed with $H_2O$ and brine, dried over $MgSO_4$ and concentrated by rotary evaporation. The crude toluidine is redissolved in dry DCM and acetic anhydride (1.2 eq) is slowly added and stirred at room temperature for 2 h. The suspension is extracted with EA and washed with brine. The combined organic layers are dried over $Na_2SO_4$, filtrated and the solvent is removed under reduced pressure. Heterocyclic ring closure was performed following a procedure by Ma et al.[2] A mixture of CuI (0.1 mmol), *o*-iodobenzamide (1 mmol), and $Na_2S{\cdot}9H_2O$ (3 mmol) in DMF (2 mL) is stirred at 80°C for 12 h. The reaction mixture is cooled to room temperature before adding 0.8 mL of conc. HCl and the reaction mixture is stirred for another 5-10 h. After adding 10 mL sat. aq. $NaHCO_3$, it is extracted with EA and purified by normal phase MPLC to furnish the desired product.

##### *Experimental data of SBTubs*

**Synthesis of SBTubA4**

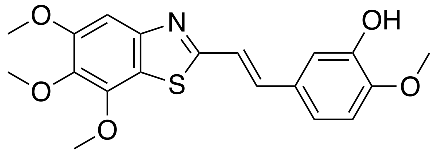

Chemical Formula: $C_{19}H_{19}NO_5S$
Molecular Weight: 373.42

**SBTubA4** was synthesized from **S1** (200 mg, 0.84 mmol) and isovanillin (127 mg, 0.84 mmol) following general procedure 1. The crude product was purified by normal phase MPLC (30→100% EA:Hex) to give **SBTubA4** (130 mg, 0.35 mmol, 42%) as yellow solid.

**$^1$H-NMR (400 MHz, $CDCl_3$):** δ = 7.40 (d, *J* = 16.2 Hz, 1H), 7.26 (s, 1H), 7.21 – 7.15 (m, 2H), 7.06 (dd, *J* = 8.4, 2.0 Hz, 1H), 6.87 (d, *J* = 8.3 Hz, 1H), 5.74 (s, 1H), 4.08 (s, 3H), 3.94 (s, 3H), 3.93 (s, 6H) ppm. **$^{13}$C-NMR (100 MHz, $CDCl_3$):** δ = 167.3, 154.1, 150.4, 147.9, 146.8, 146.1, 140.0, 136.7, 129.3, 120.8, 120.4, 119.6, 112.6, 110.8, 100.6, 61.6, 60.7, 56.4, 56.2 ppm. **$R_f$** = 0.19 (EA:Hex = 4:6). **HRMS (ESI, positive):** 374.10567 calculated for $C_{19}H_{20}NO_5S^+$ $[M+H]^+$, 374.10521 found.

**Synthesis of 2**

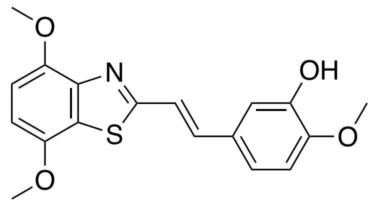

Chemical Formula: $C_{18}H_{17}NO_4S$
Molecular Weight: 343.40

**2** was synthesized from **S2** (175 mg, 0.84 mmol) and isovanillin (127 mg, 0.84 mmol) following general procedure 1. The crude product was purified by normal phase MPLC (30→80% EA:Hex) to give **2** (144 mg, 0.42 mmol, 50%) as orange solid.

**$^1$H-NMR (400 MHz, DMSO-*d*$_6$):** δ = 9.25 (s, 1H), 7.56 (d, *J* = 16.2 Hz, 1H), 7.39 (d, *J* = 16.2 Hz, 1H), 7.28 – 7.22 (m, 2H), 7.09 – 6.97 (m, 3H), 3.99 (s, 3H), 3.99 (s, 3H), 3.90 (s, 3H) ppm. **$^{13}$C-NMR (125 MHz, DMSO-*d*$_6$):** δ = 165.6, 149.3, 147.4, 147.4, 146.7, 144.5, 137.5, 128.1, 123.5, 120.3, 119.3, 113.7, 112.1, 108.3, 106.2, 56.1, 56.1, 55.6 ppm. **$R_f$** = 0.21 (EA:Hex = 4:6). **HRMS (ESI, positive):** 344.09511 calculated for $C_{18}H_{18}NO_4S^+$ $[M+H]^+$, 344.09487 found.

**Synthesis of 3**

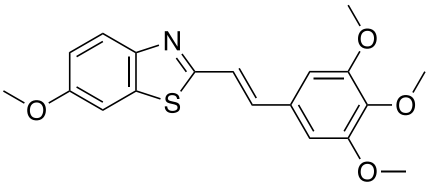

Chemical Formula: $C_{19}H_{19}NO_4S$
Molecular Weight: 357.42

**3** was synthesized from 6-methoxy-2-methylbenzothiazole (100 mg, 0.56 mmol) and 3,4,5-trimethoxybenzaldehyde (109 mg, 0.56 mmol) following general procedure 1. The crude product was purified by normal phase MPLC (10→30% EA:Hex) to give **3** (118 mg, 0.33 mmol, 59%) as yellow solid.

**$^1$H-NMR (500 MHz, CDCl$_3$):** δ = 7.86 (d, *J* = 8.9 Hz, 1H), 7.34 (d, *J* = 16.2 Hz, 1H), 7.32 (d, *J* = 2.5 Hz, 1H), 7.29 (d, *J* = 16.1 Hz, 1H), 7.07 (dd, *J* = 8.9, 2.5 Hz, 1H), 6.80 (s, 2H), 3.91 (s, 6H), 3.89 (s, 6H) ppm. **$^{13}$C-NMR (125 MHz, CDCl$_3$):** δ = 164.6, 158.1, 153.6, 148.5, 139.4, 136.6, 135.9, 131.3, 123.6, 121.9, 115.8, 104.4, 104.3, 61.1, 56.3, 56.0 ppm. **R*f*** = 0.34 (EA:Hex = 3:7). **HRMS (ESI, positive):** 358.11076 calculated for $C_{19}H_{20}NO_4S^+$ [M+H]$^+$, 358.11038 found.

**Synthesis of 4**

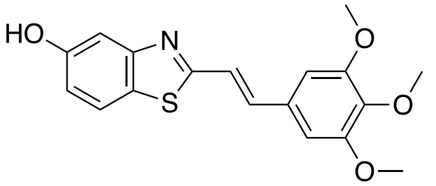

Chemical Formula: $C_{18}H_{17}NO_4S$
Molecular Weight: 343.40

**4** was synthesized from 5-hydroxy-2-methylbenzothiazole (50 mg, 0.30 mmol) and 3,4,5-trimethoxybenzaldehyde (59 mg, 0.30 mmol) following general procedure 1. The crude product was purified by normal phase MPLC (10→30% EA:Hex) to give **4** (30 mg, 0.087 mmol, 29%) as yellow solid.

**$^1$H-NMR (500 MHz, CDCl$_3$):** δ = 7.67 (d, *J* = 8.7 Hz, 1H), 7.47 (d, *J* = 2.4 Hz, 1H), 7.41 (d, *J* = 16.2 Hz, 1H), 7.29 (d, *J* = 16.2 Hz, 1H), 6.98 (dd, *J* = 8.6, 2.5 Hz, 1H), 6.78 (s, 2H), 6.49 (s, 1H), 3.89 (s, 6H), 3.89 (s, 3H) ppm. **$^{13}$C-NMR (125 MHz, CDCl$_3$):** δ = 168.7, 155.6, 154.8, 153.6, 139.6, 137.7, 131.1, 126.1, 122.2, 121.4, 115.6, 108.3, 104.7, 61.2, 56.3 ppm. **R*f*** =

0.44 (EA:Hex = 1:1). **HRMS (ESI, positive):** 344.09511 calculated for $C_{18}H_{18}NO_4S^+$ [M+H]+, 344.09479 found.

#### Synthesis of 5

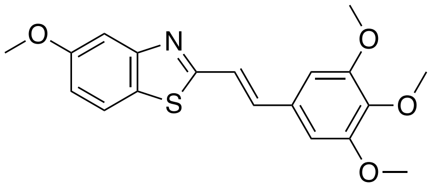

Chemical Formula: $C_{19}H_{19}NO_4S$
Molecular Weight: 357.42

To a solution of **4** (20 mg, 0.058 mmol) in acetone (1 mL) was added potassium carbonate (10 mg, 0.058 mmol, 1.24 eq) and dimethyl sulfate (5.5 $\mu$L, .0.059, 1 eq). The reaction mixture was stirred overnight at room temperature. The solvents were removed under reduced pressure. $H_2O$ (3 mL) and DCM (3 mL) was added to the residue. Extraction with DCM (3 × 3 mL). The combined organic layers were dried over $MgSO_4$, filtered and concentrated under reduced pressure. The crude product was purified by normal phase MPLC (10→30% EA:Hex) to give **5** (10 mg, 0.028 mmol, 48%) as yellow solid.

**$^1$H-NMR (500 MHz, $CDCl_3$):** δ = 7.71 (d, *J* = 8.8 Hz, 1H), 7.48 (d, *J* = 2.5 Hz, 1H), 7.45 (d, *J* = 16.1 Hz, 1H), 7.30 (d, *J* = 16.1 Hz, 1H), 7.03 (dd, *J* = 8.8, 2.5 Hz, 1H), 6.82 (s, 2H), 3.93 (s, 6H), 3.91 (s, 3H), 3.90 (s, 3H) ppm. **$^{13}$C-NMR (125 MHz, $CDCl_3$):** δ = 168.1, 159.3, 155.2, 153.7, 139.6, 137.2, 131.2, 126.3, 121.9, 121.7, 115.6, 105.4, 104.6, 61.2, 56.3, 55.8 ppm. **$R_f$** = 0.20 (EA:Hex = 2:8). **HRMS (ESI, positive):** 358.11076 calculated for $C_{19}H_{20}NO_4S^+$ [M+H]+, 358.11040 found.

#### Synthesis of 6

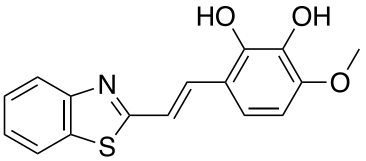

Chemical Formula: $C_{16}H_{13}NO_3S$
Molecular Weight: 299.34

2-methylbenzothiazole (0.07 mL, 0.55 mmol) and **S3** (140 mg, 0.55 mmol, 1 eq) was dissolved in DMSO (5 mL) and NaOMe (5.4 M in MeOH, 0.1 mL, 1 mmol, 1 eq) was added and the reaction mixture was stirred overnight. $H_2O$ (15 mL) was added and the mixture was extracted with ethyl acetate (3 × 10 mL). The combined organic layers were dried over $MgSO_4$ and concentrated *in vacuo*. The residue was redissolved in 3 M HCl/THF (1:1, 10 mL) and stirred at 40°C for 5 h. $H_2O$ (20 mL) was added and the reaction mixture was extracted with EA (3 × 15 mL) and dried over $MgSO_4$. The crude product was purified by normal phase MPLC (20 → 70% EA:Hex) to give **6** (135 mg, 0.45 mmol, 83%) as brown solids.

**$^{1}$H-NMR (400 MHz, DMSO-*d6*):** δ = 8.04 (dd, *J* = 8.0, 1.2 Hz, 1H), 7.92 (d, *J* = 8.0 Hz, 1H), 7.75 (d, *J* = 16.3 Hz, 1H), 7.52 – 7.43 (m, 2H), 7.39 (t, *J* = 8.1 Hz, 1H), 7.19 (d, *J* = 8.8 Hz, 1H), 6.58 (d, *J* = 8.8 Hz, 1H), 3.83 (s, 3H) ppm. **$^{13}$C-NMR (100 MHz, DMSO-*d6*):** δ = 167.8, 153.6, 149.7, 145.6, 134.0, 133.8, 133.7, 126.4, 125.1, 122.2, 122.1, 119.1, 118.6, 116.2, 103.7, 55.9 ppm. **$R_f$** = 0.34 (EA:Hex = 1:1). **HRMS (ESI, positive):** 300.06889 calculated for $C_{16}H_{14}NO_3S^+$ $[M+H]^+$, 300.06893 found.

<u>**Synthesis of 7**</u>

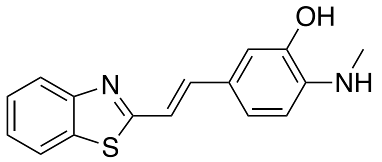

Chemical Formula: $C_{16}H_{14}N_2OS$
Molecular Weight: 282.36

To a cooled solution of **S4** (200 mg, 0.71 mmol) in 3 mL anhydrous THF was added sodium hydride (34 mg, 1.4 mmol, 2 eq) in small portions. After stirring for 10 min at room temperature, aldehyde **S6** (124 mg, 0.71 mmol) in anhydrous THF (3 mL) was added dropwise. The mixture was stirred overnight at room temperature before quenching with water (20 mL) and transferring into a separation funnel. The aqueous layer was extracted with EA (3 x 15 mL) and concentrated *in vacuo*. The crude carbamate was dissolved in 10% aq. NaOH solution (20 mL) and refluxed for 4 h. After cooling, the reaction mixture was transferred into a seperation funnel, EA (30 mL) was added and the organic layer was washed with sat. aq. $NH_4Cl$ (20 mL). The crude mixture was purified by normal phase MPLC (10→100% EA:Hex) and recrystallized from $H_2O$/MeOH to give compound **7** as brown microneedles (69 mg, 0.24 mmol, 35%).

**$^{1}$H-NMR (400 MHz, DMSO-*d6*):** δ = 9.57 (s, 1H), 8.02 (dd, *J* = 7.9, 0.7 Hz, 1H), 7.88 (d, *J* = 7.8 Hz, 1H), 7.50 – 7.39 (m, 2H), 7.36 (ddd, *J* = 8.3, 7.2, 1.2 Hz, 1H), 7.11 – 7.04 (m, 2H), 7.00 (d, *J* = 2.0 Hz, 1H), 6.45 (d, *J* = 8.2 Hz, 1H), 5.46 (q, *J* = 5.1 Hz, 1H), 2.77 (d, *J* = 5.0 Hz, 3H) ppm. **$^{13}$C-NMR (100 MHz, DMSO-*d6*):** δ = δ 167.5, 153.7, 144.1, 141.2, 139.2, 133.6, 126.3, 124.8, 122.6, 122.2, 121.9, 121.9, 115.4, 110.6, 108.3, 29.5 ppm. **$R_f$** = 0.62 (EA:Hex = 1:1). **HRMS (ESI, positive):** 283.08996 calculated for $C_{16}H_{15}N_2OS^+$ $[M+H]^+$, 283.08998 found.

#### Synthesis of 8

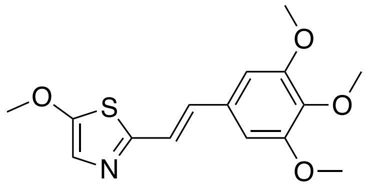

Chemical Formula: $C_{15}H_{17}NO_4S$
Molecular Weight: 307.36

Phosphorus pentasulfide (10 g, 52 mmol) was added portionwise to a solution of **S7** (315 mg, 1 mmol) in chloroform (8 mL), and the suspension was stirred at 80°C overnight. The reaction mixture was poured in 2 M NaOH (15 mL), and extracted with DCM (3 × 10 mL). The organic layer was dried over MgSO4 and concentrated under reduced pressure. The residue was purified by column chromatography (EA:Hex = 3:7) to afford compound **8** as a yellow solid (201 mg, 0.65 mmol, 64%).

**$^{1}$H-NMR (500 MHz, CDCl$_3$):** δ = 7.05 (s, 1H), 7.03 (d, *J* = 2.9 Hz, 2H), 6.71 (s, 2H), 3.95 (s, 3H), 3.89 (s, 6H), 3.87 (s, 3H) ppm. **$^{13}$C-NMR (125 MHz, CDCl$_3$):** δ = 162.3, 155.2, 153.6, 138.9, 132.3, 131.7, 122.3, 121.5, 104.0, 61.5, 61.1, 56.3 ppm. **$R_f$** = 0.42 (EA:Hex = 1:1). **HRMS (ESI, positive):** 308.09511 calculated for $C_{15}H_{18}NO_4S^+$ [M+H]$^+$, 308.09494 found.

#### Synthesis of 9

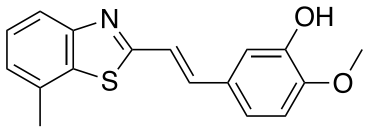

Chemical Formula: $C_{17}H_{15}NO_2S$
Molecular Weight: 297.37

**9** was synthesized from **S9** (82 mg, 0.5 mmol) and isovanillin (76 mg, 0.5 mmol) following general procedure 1. The crude product was purified by normal phase MPLC (20→70% EA:Hex) to give **9** (110 mg, 0.37 mmol, 74%) as slightly yellowish solid.

**$^{1}$H-NMR (400 MHz, CDCl$_3$):** δ = 7.83 (d, *J* = 8.1 Hz, 1H), 7.46 (d, *J* = 16.2 Hz, 1H), 7.38 (dd, *J* = 8.2, 7.3 Hz, 1H), 7.28 (s, 1H), 7.20 (d, *J* = 2.1 Hz, 1H), 7.16 (d, *J* = 7.3 Hz, 1H), 7.09 (dd, *J* = 8.3, 2.1 Hz, 1H), 6.88 (d, *J* = 8.3 Hz, 1H), 5.68 (s, 1H), 3.94 (s, 3H), 2.58 (s, 3H) ppm. **$^{13}$C-NMR (100 MHz, CDCl$_3$):** δ = 167.0, 153.9, 148.0, 146.1, 137.5, 134.9, 131.7, 129.3, 126.6, 125.5, 120.9, 120.7, 120.4, 112.7, 110.8, 56.2, 21.6 ppm. **$R_f$** = 0.31 (EA:Hex = 3:7). **HRMS (ESI, positive):** 298.08963 calculated for $C_{17}H_{16}NO_2S^+$ [M+H]$^+$, 298.08936 found.

#### Synthesis of 10

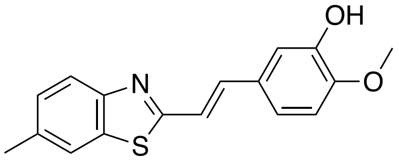

Chemical Formula: $C_{17}H_{15}NO_2S$
Molecular Weight: 297.37

**10** was synthesized from **S10** (163 mg, 1 mmol) and isovanillin (152 mg, 1 mmol) following general procedure 1. The crude product was purified by normal phase MPLC (20→70% EA:Hex) to give **10** (183 mg, 0.62 mmol, 62%) as orange solid.

**$^1$H-NMR (400 MHz, DMSO-*d6*):** δ = 9.17 (s, 1H), 7.85 (dt, *J* = 1.6, 0.7 Hz, 1H), 7.81 (d, *J* = 8.3 Hz, 1H), 7.46 (d, *J* = 16.2 Hz, 1H), 7.35 – 7.25 (m, 2H), 7.20 – 7.13 (m, 2H), 6.98 (d, *J* = 8.9 Hz, 1H), 3.82 (s, 3H), 2.44 (s, 3H) ppm. **$^{13}$C-NMR (100 MHz, DMSO-*d6*):** δ = 165.8, 151.7, 149.3, 146.7, 137.3, 135.0, 134.0, 128.2, 127.9, 121.9, 121.6, 120.3, 119.4, 113.6, 112.1, 55.7, 21.1. ppm. **R*f*** = 0.28 (EA:Hex = 3:7). **HRMS** (EI, positive): 296.0745 calculated for $C_{17}H_{14}NO_2S^{\cdot}$ [M–H]$^{\cdot}$, 296.0740 found.

#### Synthesis of 11

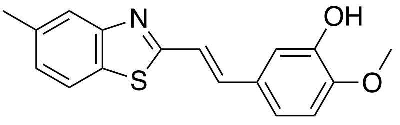

Chemical Formula: $C_{17}H_{15}NO_2S$
Molecular Weight: 297.37

**11** was synthesized from **S11** (82 mg, 0.5 mmol) and isovanillin (76 mg, 0.5 mmol) following general procedure 1. The crude product was purified by normal phase MPLC (20→70% EA:Hex) to give **11** (100 mg, 0.34 mmol, 67%) as slightly yellowish solid.

**$^1$H-NMR (400 MHz, CDCl$_3$):** δ = 7.82 (dt, *J* = 1.5, 0.8 Hz, 1H), 7.75 (d, *J* = 8.1 Hz, 1H), 7.45 (d, *J* = 16.1 Hz, 1H), 7.30 (d, *J* = 0.6 Hz, 1H), 7.25 – 7.20 (m, 2H), 7.11 (dd, *J* = 8.5, 2.1 Hz, 1H), 6.91 (d, *J* = 8.3 Hz, 1H), 5.78 (s, 1H), 3.97 (s, 3H), 2.53 (s, 3H) ppm. **$^{13}$C-NMR (100 MHz, CDCl$_3$):** δ = 167.6, 154.4, 148.0, 146.1, 137.3, 136.5, 131.3, 129.3, 126.9, 123.0, 121.1, 120.8, 120.7, 112.7, 110.8, 56.2, 21.6 ppm. **R*f*** = 0.33 (EA:Hex = 3:7). **HRMS (ESI, positive):** 298.08963 calculated for $C_{17}H_{16}NO_2S^+$ [M+H]$^+$, 298.08932 found.

#### Synthesis of 12

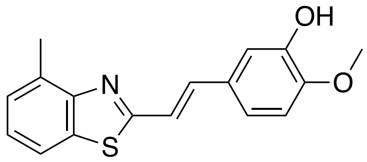

Chemical Formula: $C_{17}H_{15}NO_2S$
Molecular Weight: 297.37

**12** was synthesized from **S12** (230 mg, 1.41 mmol) and isovanillin (214 mg, 1.41 mmol) following general procedure 1. The crude product was purified by normal phase MPLC (20→70% EA:Hex) to give **12** (338 mg, 1.14 mmol, 81%) as yellow solid.

**$^1$H-NMR (400 MHz, DMSO-*d*$_6$):** δ = 9.16 (s, 1H), 7.86 (dd, *J* = 5.3, 3.9 Hz, 1H), 7.46 (d, *J* = 16.2 Hz, 1H), 7.36 (d, *J* = 16.2 Hz, 1H), 7.32 – 7.27 (m, 2H), 7.22 – 7.16 (m, 2H), 6.98 (d, *J* = 8.2 Hz, 1H), 3.82 (s, 3H), 2.66 (s, 3H) ppm. **$^{13}$C-NMR (100 MHz, DMSO-*d*$_6$):** δ = 165.9, 152.7, 149.3, 146.7, 137.7, 133.6, 131.9, 128.2, 126.8, 125.2, 120.2, 119.6, 119.4, 113.8, 112.1, 55.6, 18.1 ppm. **$R_f$** = 0.70 (EA:Hex = 1:1). **HRMS (ESI, positive):** 298.08963 calculated for $C_{17}H_{16}NO_2S^+$ [M+H]$^+$, 298.08949 found.

#### Synthesis of 13

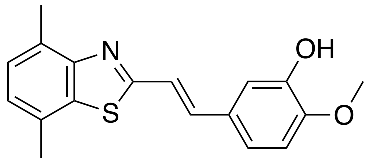

Chemical Formula: $C_{18}H_{17}NO_2S$
Molecular Weight: 311.40

**13** was synthesized from **S13** (200 mg, 1.13 mmol) and isovanillin (172 mg, 1.13 mmol) following general procedure 1. The crude product was purified by normal phase MPLC (20→70% EA:Hex) to give **13** (219 mg, 0.70 mmol, 62%) as orange solid.

**$^1$H-NMR (500 MHz, CDCl$_3$):** δ = 7.40 (d, *J* = 16.1 Hz, 1H), 7.33 (d, *J* = 16.1 Hz, 1H), 7.20 (d, *J* = 2.1 Hz, 1H), 7.18 (d, *J* = 7.4 Hz, 1H), 7.09 (dd, *J* = 8.3, 2.1 Hz, 1H), 7.06 (d, *J* = 7.4 Hz, 1H), 6.87 (d, *J* = 8.3 Hz, 1H), 5.70 (s, 1H), 3.93 (s, 3H), 2.72 (s, 3H), 2.52 (s, 3H). **$^{13}$C-NMR (125 MHz, CDCl$_3$):** δ = 166.2, 147.9, 146.1, 137.3, 134.5, 130.1, 129.5, 128.8, 127.2, 125.5, 121.2, 120.7, 112.8, 110.8, 56.2, 21.3, 18.4 ppm. **$R_f$** = 0.21 (EA:Hex = 2:8). **HRMS (ESI, positive):** 312.10528 calculated for $C_{18}H_{18}NO_2S^+$ [M+H]$^+$, 312.10510 found.

#### Synthesis of 14

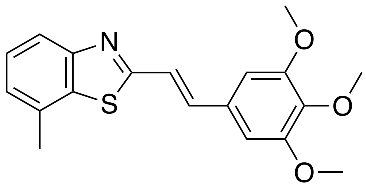

Chemical Formula: $C_{19}H_{19}NO_3S$
Molecular Weight: 341.43

**14** was synthesized from **S9** (57 mg, 0.35 mmol) and 3,4,5-trimethoxybenzaldehyde (69 mg, 0.35 mmol) following general procedure 1. The crude product was purified by normal phase MPLC (5→40% EA:Hex) to give **14** (56 mg, 0.16 mmol, 47%) as slightly yellowish solid.

**$^1$H-NMR (500 MHz, $CDCl_3$):** δ = 7.83 (dt, *J* = 8.0, 0.8 Hz, 1H), 7.46 (d, *J* = 16.1 Hz, 1H), 7.39 (dd, *J* = 8.1, 7.3 Hz, 1H), 7.34 (d, *J* = 16.1 Hz, 1H), 7.18 (dt, *J* = 7.2, 0.9 Hz, 1H), 6.82 (s, 2H), 3.92 (s, 6H), 3.90 (s, 3H), 2.58 (s, 3H) ppm. **$^{13}$C-NMR (125 MHz, $CDCl_3$):** δ = 166.5, 153.8, 153.7, 139.5, 137.5, 134.9, 131.7, 131.2, 126.7, 125.7, 121.9, 120.5, 104.6, 61.2, 56.3, 21.6 ppm. **$R_f$** = 0.19 (EA:Hex = 1:9). **HRMS (ESI, positive):** 342.11584 calculated for $C_{19}H_{20}NO_3S^+$ $[M+H]^+$, 342.11547 found.

#### Synthesis of SBTub2M

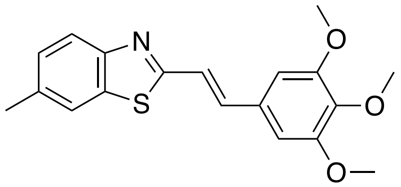

Chemical Formula: $C_{19}H_{19}NO_3S$
Molecular Weight: 341.43

**SBTub2M** was synthesized from **S10** (69 mg, 0.42 mmol) and 3,4,5-trimethoxybenzaldehyde (83 mg, 0.42 mmol) following general procedure 1. The crude product was purified by normal phase MPLC (5→40% EA:Hex) to give **GAO558** (83 mg, 0.24 mmol, 58%) as yellow solid.

**$^1$H-NMR (500 MHz, $CDCl_3$):** δ = 7.86 (d, *J* = 8.3 Hz, 1H), 7.65 (dt, *J* = 1.7, 0.8 Hz, 1H), 7.40 (d, *J* = 16.2 Hz, 1H), 7.32 (d, *J* = 16.2 Hz, 1H), 7.28 (ddd, *J* = 8.3, 1.7, 0.8 Hz, 1H), 6.81 (s, 2H), 3.92 (s, 6H), 3.89 (s, 3H), 2.49 (s, 3H) ppm. **$^{13}$C-NMR (125 MHz, $CDCl_3$):** δ = 165.9, 153.5, 151.9, 139.3, 137.0, 135.7, 134.4, 131.1, 128.0, 122.4, 121.8, 121.3, 104.4, 61.0, 56.2, 21.6 ppm. **$R_f$** = 0.20 (EA:Hex = 1:9). **HRMS (ESI, positive):** 342.11584 calculated for $C_{19}H_{20}NO_3S^+$ $[M+H]^+$, 342.11565 found.

#### Synthesis of 16

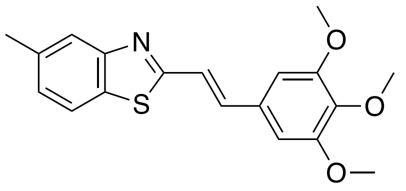

Chemical Formula: $C_{19}H_{19}NO_3S$
Molecular Weight: 341.43

**16** was synthesized from **S11** (69 mg, 0.32 mmol) and 3,4,5-trimethoxybenzaldehyde (83 mg, 0.32 mmol) following general procedure 1. The crude product was purified by normal phase MPLC (5→40% EA:Hex) to give **16** (59 mg, 0.17 mmol, 54%) as slightly yellowish solid.

**$^1$H-NMR (500 MHz, $CDCl_3$):** δ = 7.78 (p, *J* = 0.8 Hz, 1H), 7.73 (d, *J* = 8.1 Hz, 1H), 7.42 (d, *J* = 16.2 Hz, 1H), 7.32 (d, *J* = 16.1 Hz, 1H), 7.21 (ddd, *J* = 8.1, 1.7, 0.7 Hz, 1H), 6.81 (s, 2H), 3.92 (s, 6H), 3.89 (s, 3H), 2.50 (s, 3H) ppm. **$^{13}$C-NMR (125 MHz, $CDCl_3$):** δ = 167.1, 154.4, 153.7, 139.5, 137.3, 136.6, 131.4, 131.2, 127.2, 123.1, 121.9, 121.1, 104.6, 61.2, 56.3, 21.6 ppm. **$R_f$** = 0.25 (EA:Hex = 1:9). **HRMS (ESI, positive):** 342.11584 calculated for $C_{19}H_{20}NO_3S^+$ $[M+H]^+$, 342.11581 found.

#### Synthesis of 17

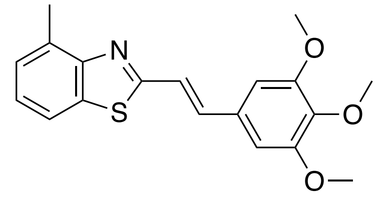

Chemical Formula: $C_{19}H_{19}NO_3S$
Molecular Weight: 341.43

**17** was synthesized from **S12** (66 mg, 0.40 mmol) and 3,4,5-trimethoxybenzaldehyde (79 mg, 0.40 mmol) following general procedure 1. The crude product was purified by normal phase MPLC (5→40% EA:Hex) to give **17** (35 mg, 0.10 mmol, 25%) as slightly yellowish solid.

**$^1$H-NMR** (500 MHz, $CDCl_3$): δ = 7.72 – 7.67 (m, 1H), 7.43 (d, *J* = 16.2 Hz, 1H), 7.38 (d, *J* = 16.1 Hz, 1H), 7.28 – 7.27 (m, 2H), 6.83 (s, 2H), 3.92 (s, 6H), 3.90 (s, 3H), 2.77 (s, 3H) ppm. **$^{13}$C-NMR (125 MHz, $CDCl_3$):** δ = 166.1, 153.7, 139.5, 137.5, 134.2, 133.0, 131.3, 127.1, 125.5, 122.3, 119.1, 106.3, 104.6, 61.2, 56.3, 18.7. ppm. **$R_f$** = 0.53 (EA:Hex = 3:7). **HRMS** (ESI, positive): 342.11584 calculated for $C_{19}H_{20}NO_3S^+$ $[M+H]^+$, 342.11552 found.

**Synthesis of 18**

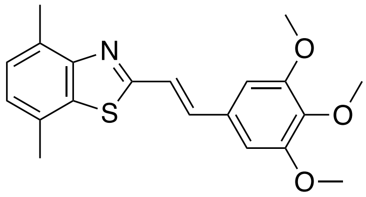

Chemical Formula: $C_{20}H_{21}NO_3S$
Molecular Weight: 355.45

**18** was synthesized from **S13** (188 mg, 1.1 mmol) and 3,4,5-trimethoxybenzaldehyde (208 mg, 1.1 mmol) following general procedure 1. The crude product was purified by normal phase MPLC (5→40% EA:Hex) to give **18** (60 mg, 0.17 mmol, 16%) as yellow solid.

**$^1$H-NMR (500 MHz, CDCl$_3$):** δ = 7.41 (s, 1H), 7.41 (s, 1H), 7.19 (dd, *J* = 7.3, 1.0 Hz, 1H), 7.08 (dd, *J* = 7.3, 1.0 Hz, 1H), 6.83 (s, 2H), 3.92 (s, 6H), 3.90 (s, 3H), 2.72 (s, 3H), 2.53 (s, 3H) ppm. **$^{13}$C-NMR (125 MHz, CDCl$_3$):** δ = 165.6, 153.5, 152.8, 139.3, 137.2, 134.5, 131.2, 130.1, 128.8, 127.1, 125.6, 122.4, 104.4, 61.0, 56.2, 21.2, 18.3 ppm. **$R_f$** = 0.36 (EA:Hex = 1:9). **HRMS (ESI, positive):** 356.13149 calculated for $C_{20}H_{22}NO_3S^+$ [M+H]$^+$, 356.13112 found.

**Synthesis of SBTub3P**

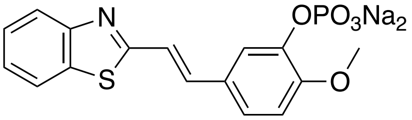

Chemical Formula: $C_{16}H_{12}NNa_2O_5PS$
Molecular Weight: 407.29

To a cooled solution of **SBTub3** (594 mg, 2.1 mmol), TEA (0.58 mL, 4.2 mmol, 2 eq), DMAP (51 mg, 0.42 mmol, 0.2 eq) and $CCl_4$ (1 mL) in MeCN (20 mL), was slowly added dibenzyl phosphite (770 mg, 2.9 mmol, 1.4 eq) and the resulting mixture was stirred at room temperature until phosphorylation was complete (~15 min). The volatiles were evaporated to give the phenolic (bis)benzyl ester, which was purified by flash column purification (EA:Hex = 4:6) to give 1.14 g of a colourless solid, that was directly redissolved in a mixture of DCM:TFA (14 mL, 1:1) and stirred for 6 h at 40°C. After removing all volatiles under reduced pressure, water (50 mL) was added and the suspension was basified with 2M NaOH until all solids fully dissolved. The clear solution was transferred into a separation funnel and washed with EA (3 × 30 mL). The aqueous layer was set to pH ~2 with 2M HCl and loaded on a C18 solid phase extraction cartridge, washed with water (3 × 10 mL), eluted with MeOH and concentrated *in vacuo*. The crude product was purified by RP-MPLC (0→25% MeCN:$H_2O$) to give **SBTub3P** (393 mg, 1.1 mmol, 52%) as slightly yellowish powder after lyophilizing.

**$^1$H-NMR (400 MHz, $D_2O$):** δ = 7.85 (d, *J* = 7.9 Hz, 1H), 7.79 (d, *J* = 8.0 Hz, 1H), 7.69 (s, 1H), 7.47 – 7.37 (m, 2H), 7.34 (t, *J* = 7.6 Hz, 1H), 7.22 (d, *J* = 16.2 Hz, 1H), 7.15 (dd, *J* = 8.4, 1.6 Hz, 1H), 6.90 (d, *J* = 8.5 Hz, 1H), 3.80 (s, 3H) ppm. **$^{13}$C-NMR (100 MHz, $D_2O$):** δ = 169.6, 152.1, 151.2, 143.2, 138.6, 133.5, 128.0, 126.5, 125.3, 123.0, 121.9, 121.3, 118.7, 118.3, 112.3, 55.6 ppm. **HRMS (ESI, negative):** 362.02575 calculated for $C_{16}H_{13}NO_5PS^-$ $[M{-}H]^-$,362.02566 found.

**<u>Synthesis of SBTubA4P</u>**

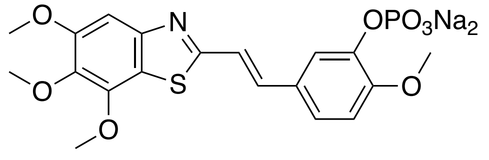

Chemical Formula: $C_{19}H_{18}NNa_2O_8PS$
Molecular Weight: 497.37

A solution of SBTubA4 (1.10 g, 2.95 mmol, 1 eq) in 30 mL MeCN was cooled to 0°C. Dibenzyl diphosphate (1.16 g, 4.42 mmol, 1.5 eq), DIPEA (1 mL, 6.19 mmol, 2.1 eq), DMAP (7 mg, 58.9 *µ*mol, 0.02 eq) and $CCl_4$ (1 mL). After stirring for 5 min the reaction mixture was allowed to warm to room temperature. After 4 h no further conversion was observed by TLC and HPLC-MS. Water (80 mL) was added and the reaction mixture is extracted with EA (3 x 50 mL) and washed with brine (1 x 50 mL). The combined organic layers were dried over $Na_2SO_4$, filtrated and concentrated *in vacuo*. The crude was purified by normal phase MPLC to give the bis(benzyl) phosphate of SBTubA4 (1.38 g, 2.17 mmol, 74%) as colourless solid.

**$^1$H-NMR (400 MHz, $CDCl_3$):** δ = 7.40 – 7.29 (m, 13H), 7.27 (s, 1H), 7.08 (d, *J* = 16.1 Hz, 1H), 6.92 (d, *J* = 8.2 Hz, 1H), 5.20 (dd, *J* = 8.2, 2.0 Hz, 4H), 4.10 (s, 3H), 3.95 (s, 3H), 3.94 (s, 3H), 3.83 (s, 3H) ppm. **$^{13}$C-NMR (100 MHz, $CDCl_3$):** δ = 166.8, 154.1, 151.8, 150.5, 146.8, 140.2, 135.8, 135.5, 128.8, 128.1, 125.6, 120.8, 120.1, 119.8, 112.8, 100.6, 70.2, 61.6, 60.7, 56.4, 56.1 ppm.

The bis(benzyl)phosphate (1.33 g, 2.1 mmol, 1 eq) was dissolved in dichloromethane (40 mL) at 0°C under nitrogen atmosphere. Bromotrimethylsilane (0.583 *µ*L, 4.42 mmol, 2.1 eq) was added was added dropwise upon which the phosphoric acid intermediate started to precipitate. Methanol (20 mL) was added and stirring was continued for 15 min before concentrating *in vacuo.* The crude mixture was resuspended in EtOH (50 mL) and sodium methoxide (0.77 mL 6M in MeOH, 4.63 mmol, 2.2 eq) was added at 0°C. Stirring was continued overnight at room temperature. The solvent was removed under reduced pressure and the crude product was purified by RP-MPLC ( 10→100% MeCN:$H_2O$) to give **SBTubA4P** (864 mg, 1.74 mmol, 83%) as yellow solid.

**$^1$H-NMR (400 MHz, MeOD$_4$):** δ = 7.99 (d, *J* = 2.1 Hz, 1H), 7.48 (d, *J* = 16.1 Hz, 1H), 7.34 (d, *J* = 16.1 Hz, 1H), 7.23 (s, 1H), 7.16 (dd, *J* = 8.5, 2.1 Hz, 1H), 6.95 (dd, *J* = 8.5, 1.0 Hz, 1H), 4.05 (s, 3H), 3.93 (s, 3H), 3.88 (s, 3H), 3.87 (s, 3H) ppm. **$^{13}$C-NMR (100 MHz, MeOD$_4$):** δ = 169.7, 155.6, 153.1, 151.2, 147.9, 145.8, 141.1, 139.3, 129.7, 123.2, 120.2, 120.1, 119.4, 112.9, 101.1, 61.8, 61.1, 56.8, 56.5 ppm. **HRMS (ESI, positive):** 498.0359 calculated for $C_{19}H_{19}NNa_2O_8PS^+$ [M+H]$^+$, 498.03618 found.

#### ***Experimental data of synthetic building blocks***

**Synthesis of S1**

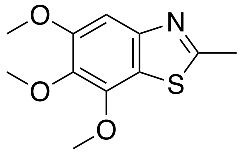

Chemical Formula: $C_{11}H_{13}NO_3S$
Molecular Weight: 239.29

**S1** was synthesized from 3,4,5-trimethoxyaniline (3.0 g, 16 mmol) following general procedure 2. The crude product was purified by normal phase MPLC (0→30% EA:Hex) to give **S1** (0.56 g, 2.3 mmol, 14%) as yellow oil.

**$^1$H-NMR (400 MHz, CDCl$_3$):** δ = 7.23 (s, 1H), 4.05 (s, 3H), 3.92 (s, 3H), 3.90 (s, 3H), 2.78 (s, 3H) ppm. **$^{13}$C-NMR (100 MHz, CDCl$_3$):** δ = 166.8, 153.7, 149.8, 146.7, 139.3, 120.5, 100.2, 61.4, 60.5, 56.2, 20.0 ppm. **$R_f$** = 0.23 (EA:Hex = 2:8). **HRMS (ESI, positive):** 240.06889 calculated for $C_{11}H_{14}NO_3S^+$ [M+H]$^+$, 240.06869 found.

**Synthesis of S2**

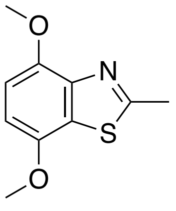

Chemical Formula: $C_{10}H_{11}NO_2S$
Molecular Weight: 209.26

**S2** was synthesized from 2,5-dimethoxyaniline (3.0 g, 20 mmol) following general procedure 2. The crude product was purified by normal phase MPLC (5→20% EA:Hex) to give **S2** (0.75 g, 3.6 mmol, 18%) as yellow oil.

**$^1$H-NMR (400 MHz, $CDCl_3$):** δ = 6.80 (d, $J$ = 8.6 Hz, 1H), 6.71 (d, $J$ = 8.6 Hz, 1H), 3.99 (s, 3H), 3.93 (s, 3H), 2.85 (s, 3H) ppm. **$^{13}$C-NMR (100 MHz, $CDCl_3$):** δ = 166.6, 148.1, 147.6, 144.6, 126.1, 106.6, 105.0, 56.3, 56.2, 20.2. ppm. **$R_f$** = 0.24 (EA:Hex = 2:8). **HRMS (ESI, positive):** 210.05833 calculated for $C_{10}H_{12}NO_2S^+$ [M+H]$^+$, 210.05818 found.

**<u>Synthesis of S3 MOM protected benzaldehyde</u>**

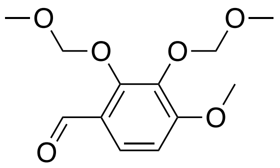

Chemical Formula: $C_{12}H_{16}O_6$
Molecular Weight: 256.25

2,3-dihydroxy-4-methoxybenzaldehyde (100 mg, 0.6 mmol) was dissolved in dry dichloromethane under nitrogen atmosphere and cooled to 0°C. Diisopropylethylamine (0.3 mL, 1.8 mmol, 3 eq) was added dropwise. MOM-Cl (0.09 mL, 1.2 mmol, 2 eq) was added dropwise. The reaction was stirred at 0°C for 2 h, and then warmed to room temperature overnight. Water (30 mL) was added and the aqueous layer was extracted with DCM (3 × 30 mL). The combined organic layers were washed with AcOH (2 × 15 mL, 10%), sat. aq. $NaHCO_3$ (15 mL) and brine (30 mL), dried over $MgSO_4$ and the solvent was removed *in vacuo*. The compound was used without further purification.

**$^1$H-NMR (400 MHz, $CDCl_3$):** δ = 10.28 (s, 1H), 7.65 (d, $J$ = 8.8 Hz, 1H), 6.81 (d, $J$ = 8.8 Hz, 1H), 5.26 (s, 2H), 5.13 (s, 2H), 3.93 (s, 3H), 3.60 (s, 3H), 3.57 (s, 3H) ppm. **$^{13}$C-NMR (100 MHz, $CDCl_3$):** δ = 189.2, 159.1, 154.3, 138.3, 125.0, 124.3, 108.1, 100.5, 98.8, 58.2, 57.7, 56.3 ppm. **$R_f$** = 0.53 (EA:Hex = 4:6). **LRMS (ESI, positive):** 257.10196 calculated for $C_{12}H_{17}O_6^+$ [M+H]$^+$, 257.1 found.

**<u>Synthesis of S4[3]</u>**

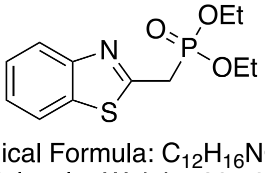

Chemical Formula: $C_{12}H_{16}NO_3PS$
Molecular Weight: 285.30

A mixture of **S5** (0.50 g, 2.19 mmol) in triethyl phosphite (1 mL) was refluxed overnight. The excess $P(OEt)_3$ was removed under a nitrogen flow and the residue was purified by silica gel column chromatography to afford product **S4** in quantitative yield.

**$^1$H-NMR (400 MHz, $CDCl_3$):** δ = 8.00 (ddt, *J* = 8.1, 1.2, 0.7 Hz, 1H), 7.86 (ddd, *J* = 8.0, 1.2, 0.6 Hz, 1H), 7.47 (ddd, *J* = 8.3, 7.2, 1.3 Hz, 1H), 7.38 (ddt, *J* = 8.2, 7.2, 1.0 Hz, 1H), 4.16 (p, *J* = 7.3 Hz, 4H), 3.76 (s, 1H), 3.71 (s, 1H), 1.32 (t, *J* = 7.1 Hz, 6H) ppm. **$R_f$** = 0.12 (EA:Hex = 1:1). **LRMS (ESI, positive):** 286.06613 calculated for $C_{12}H_{17}NO_3PS^+$ $[M+H]^+$, 286.2 found.

**Synthesis of S5[4]**

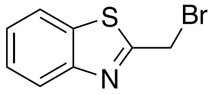

Chemical Formula: $C_8H_6BrNS$
Molecular Weight: 228.11

2-methylbenzothiazole (1.7 mL, 13.4 mmol), N-bromosuccinimide (2.39 g, 13.40 mmol), and AIBN (0.5 g, 3.04 mmol) was dissolved in $CCl_4$ (25 mL) and refluxed for overnight. The mixture was cooled and filtered. The filtrate was concentrated *in vacuo* and purified by normal phase MPLC (0→ 10% EA:Hex) to give **S5** (1.29 g, 42%) as a yellow oil.

**$^1$H-NMR (400 MHz, $CDCl_3$):** δ = 8.03 (dt, *J* = 8.3, 0.9 Hz, 1H), 7.88 (dt, *J* = 8.0, 1.0 Hz, 1H), 7.51 (ddd, *J* = 8.3, 7.2, 1.3 Hz, 1H), 7.43 (ddd, *J* = 8.3, 7.2, 1.3 Hz, 1H), 4.82 (s, 2H) ppm. **$R_f$** = 0.53 (EA:Hex = 1:9).

**Synthesis of S6[5]**

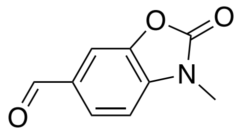

Chemical Formula: $C_9H_7NO_3$
Molecular Weight: 177.16

To 3-Methyl-2-benzoxazolone (0.50 g, 3.35 mmol) and hexamethylenetetramine (1.4 g, 10 mmol, 3 eq) was added TFA (10 mL). The resulting mixture was refluxed overnight at 80°C. The reaction mixture was cooled and poured into ice-water. The solution was neutralized with sat. aq. $NaHCO_3$ and extracted with ethyl acetate (3 × 30 mL). The combined organic layers were dried over $Na_2SO_4$ and concentrated under reduced pressure to obtain the desired aldehyde **S6** as colorless solid (0.53 mg, 3 mmol, 90%).

**$^1$H-NMR (400 MHz, $CDCl_3$):** δ = 9.95 (s, 1H), 7.79 (dd, *J* = 8.0, 1.4 Hz, 1H), 7.74 (d, *J* = 1.4 Hz, 1H), 7.11 (d, *J* = 8.0 Hz, 1H), 3.48 (s, 3H) ppm. **$R_f$** = 0.38 (EA:Hex = 1:1).

#### Synthesis of S7

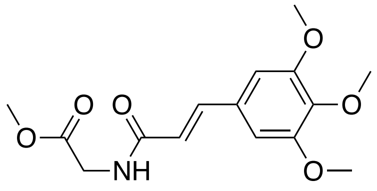

Chemical Formula: $C_{15}H_{19}NO_6$
Molecular Weight: 309.32

To a stirring suspension of cinnamic acid **S8** (0.5 g, 2.1 mmol) was added $SOCl_2$ (1.5 mL, 21 mmol, 10 eq) at 0°C. The mixture was stirred for 3 h at room temperature. The solvent was removed, the cinnamic acid chloride was redissolved in $CHCl_3$ (5 mL) and added to a stirred suspension of glycine methyl ester hydrochloride (290 mg, 2.31 mmol, 1.1 eq) and triethylamine (0.64 mL, 4.62 mmol, 2.2 eq) in chloroform (5 mL). Stirring was continued for 3 h, before the solvent was removed under reduced pressure to afford a yellowish crude, which was purified by flash column chromatography (EA:Hex = 3:7) to afford desired compound **S7** (0.55 g, 1.78 mmol, 85%) as colorless solid.

**$^1$H-NMR (500 MHz, $CDCl_3$):** δ = 7.56 (d, *J* = 15.5 Hz, 1H), 6.74 (s, 2H), 6.38 (d, *J* = 15.5 Hz, 1H), 6.15 (t, *J* = 4.6 Hz, 1H), 4.20 (d, *J* = 5.0 Hz, 2H), 3.89 (s, 6H), 3.87 (s, 3H), 3.80 (s, 3H) ppm. **$^{13}$C-NMR (125 MHz, $CDCl_3$):** δ = 170.7, 165.9, 153.6, 142.0, 139.9, 130.3, 119.2, 105.2, 61.1, 56.3, 52.6, 41.6 ppm. **$R_f$** = 0.42 (EA:Hex = 9:1). **LRMS (ESI, positive):** 310.13 calculated for $C_{15}H_{20}NO_6^+$ $[M+H]^+$, 310.1 found.

#### Synthesis of cinnamic acid S8

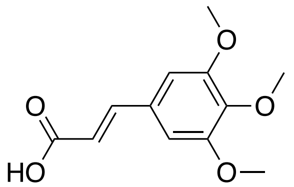

Chemical Formula: $C_{12}H_{14}O_5$
Molecular Weight: 238.24

To a solution of 3,4,5-trimethoxybenzaldehyde (5 g, 25.5 mmol), malonic acid (3.18 g, 30.6 mmol, 1.2 eq) in pyridine (10 mL) was added piperidine (0.25 mL, 2.55 mmol, 0.1 eq) and refluxed for 16 h. The solvent was removed under reduced pressure, redissolved in EA (20 mL) and washed with 2 M HCl (2 × 10 mL), water (10 mL) and brine (10 mL) and recrystallized to afford **S8** as colorless solid (5.19 g, 21.7 mmol, 85%).

**$^1$H-NMR (400 MHz, $CDCl_3$):** δ = 7.71 (d, *J* = 15.9 Hz, 1H), 6.78 (s, 2H), 6.36 (d, *J* = 15.9 Hz, 1H), 3.90 (s, 6H), 3.90 (s, 3H) ppm. **$^{13}$C-NMR (100 MHz, $CDCl_3$):** δ =171.4, 153.6, 147.2, 140.7, 129.6, 116.4, 105.7, 61.2, 56.3. ppm. **$R_f$** = 0.57 (EA:Hex + 0.1% Formic Acid = 1:1). **HRMS (EI, positive):** 238.0836 calculated for $C_{12}H_{14}O_5^+$ $[M]^+$, 238.0836 found.

**Synthesis of S9**

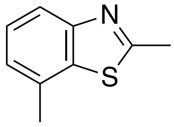

Chemical Formula: $C_9H_9NS$
Molecular Weight: 163.24

**S9** was synthesized following general procedure 3. Starting from 2-Iodo-3-nitrotoluene (1.5 g, 5.7 mmol), obtaining *N*-(2-iodo-3-methylphenyl)acetamide (1.5 g, 5.45 mmol, 96% over two steps). (550 mg, 2 mmol). The crude product from the copper catalysed heteroaromatic ring closure of the acetamide (550 mg, 2 mmol) was purified by normal phase MPLC (10→30% EA:Hex) to give **S9** (244 mg, 1.49 mmol, 75%) as colourless solid.

**$^1$H-NMR (500 MHz, $CDCl_3$):** δ = 7.80 (d, *J* = 8.1 Hz, 1H), 7.37 (dd, *J* = 8.1, 7.3 Hz, 1H), 7.15 (dt, *J* = 7.3, 0.9 Hz, 1H), 2.85 (s, 3H), 2.54 (s, 3H). **$^{13}$C-NMR (125 MHz, $CDCl_3$):** δ = 166.6, 153.4, 136.3, 131.7, 126.2, 125.0, 119.9, 21.6, 20.4 ppm. **$R_f$** = 0.52 (EA:Hex = 2:8). **LRMS** (APCI, positive): 164.05285 calculated for $C_9H_{10}NS^+$ $[M+H]^+$, 164.0 found.

**Synthesis of S10**[6]

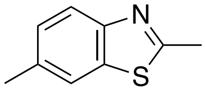

Chemical Formula: $C_9H_9NS$
Molecular Weight: 163.24

**S10** was synthesized from *p*-toluidine (3.1 mL, 28 mmol) following general procedure 2. The crude product was purified by Kugelrohr distillation (50 mbar, 200°C) to give **S10** (2.2 g, 13 mmol, 48%). **$^1$H-NMR (400 MHz, $CDCl_3$):** δ = 7.82 (d, *J* = 8.3 Hz, 1H), 7.60 (dt, *J* = 1.8, 0.8 Hz, 1H), 7.24 (dd, *J* = 8.3, 1.8 Hz, 1H), 2.81 (s, 3H), 2.46 (s, 3H) ppm. **$R_f$** = 0.20 (EA:Hex = 5:95).

#### Synthesis of S11

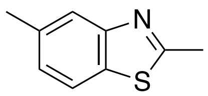

Chemical Formula: $C_9H_9NS$
Molecular Weight: 163.24

**S11** was synthesized following general procedure 3. Starting from 4-Iodo-3-nitrotoluene (1.5 g, 5.7 mmol), first obtaining *N*-(2-iodo-5-methylphenyl)acetamide (1.52 g, 5.5 mmol, 97% over two steps). The crude product from the copper catalysed heteroaromatic ring closure of the acetamide (550 mg, 2 mmol) was purified by normal phase MPLC (10→30% EA:Hex) to give **S11** (185 mg, 1.1 mmol, 57%) as colourless solid.

**$^1$H-NMR (500 MHz, CDCl$_3$):** δ = 7.75 (s, 1H), 7.68 (d, *J* = 8.2 Hz, 1H), 7.17 (dd, *J* = 8.2, 1.6 Hz, 1H), 2.82 (s, 3H), 2.49 (s, 3H) ppm. **$^{13}$C-NMR (125 MHz, CDCl$_3$):** δ = 167.2, 153.9, 136.1, 132.7, 126.4, 122.6, 121.0, 21.6, 20.3 ppm. **R$_f$** = 0.52 (EA:Hex = 2:8). **LRMS** (APCI, positive): 164.05285 calculated for $C_9H_{10}NS^+$ $[M+H]^+$, 164.0 found.

#### Synthesis of S12

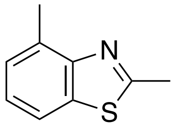

Chemical Formula: $C_9H_9NS$
Molecular Weight: 163.24

**S12** was synthesized from *o*-toluidine (3.6 mL, 34 mmol) following general procedure 2. The crude product was purified by normal phase MPLC (0→20% EA:Hex) to give **S12** (1.93 g, 12 mmol, 35%) as yellow oil.

**$^1$H-NMR (400 MHz, CDCl$_3$):** δ =7.68 (t, *J* = 4.6 Hz, 1H), 7.29 – 7.25 (m, 2H), 2.88 (s, 3H), 2.78 (s, 3H). ppm. **$^{13}$C-NMR (100 MHz, CDCl$_3$):** δ = 165.7, 152.8, 135.6, 132.4, 126.6, 124.6, 118.9, 20.3, 18.6 ppm. **R$_f$** = 0.38 (EA:Hex = 1:40). **HRMS (ESI, positive):** 164.05285 calculated for $C_9H_{10}NS$ $[M+H]^+$, 164.05281 found.

#### Synthesis of S13

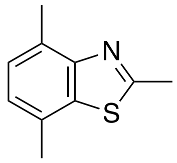

Chemical Formula: $C_{10}H_{11}NS$
Molecular Weight: 177.27

**S13** was synthesized from 2,5-dimethylaniline (5 mL, 40 mmol) following general procedure 2. The crude product was purified by Kugelrohr distillation (50 mbar, 200°C) to give **S13** (1.05 g, 6 mmol, 15%) as yellow oil.

**$^1$H-NMR (400 MHz, $CDCl_3$):** δ = 7.18 (d, $J$ = 7.4 Hz, 1H), 7.06 (d, $J$ = 7.4 Hz, 1H), 2.89 (s, 3H), 2.71 (s, 3H), 2.49 (s, 3H) ppm. **$^{13}$C-NMR (100 MHz, $CDCl_3$):** δ = 165.2, 152.1, 135.7, 129.2, 128.5, 126.5, 124.5, 21.1, 20.1, 18.1 ppm. **$R_f$** = 0.5 (EA:Hex = 1:20). **HRMS (EI, positive):** 177.0612 calculated for $C_{10}H_{11}NS$ $[M]^+$, 177.0603 found.

#### Part B: Photocharacterisation *in vitro*

##### *Spectrophotometry*

Absorption spectra in cuvette ("UV-Vis") were acquired on a Cary 60 UV-Vis Spectrophotometer from Agilent (1 cm pathlength). For photoisomerisation measurements, Hellma microcuvettes (108-002-10-40) taking 500 $\mu$L volume to top of optical window were used with the default test solution concentrations of 25 $\mu$M. Measurements were performed in PBS at pH~7.4 with 10% of DMSO or 50% DMSO indicated by asterisk (*). Prodrugs SBTubA4P and SBTub3P were measured in PBS only. Photoisomerisations were performed at room temperature. Medium-power LEDs (H2A1-models spanning 360–490 nm from Roithner Lasertechnik) were used to deliver high-intensity and relatively monochromatic light (FWHM ~25 nm) into the cuvette, for rapid PSS determinations that were also predictive of what would be obtained in LED-illuminated cell culture (**Fig S1**). Spectra of pure *E* and *Z* isomers were acquired from the inline Diode Array Detector during analytical separation on the HPLC (injection of 10 $\mu$L, 5→100% MeCN:$H_2O$ over 20 min), after injecting DMSO stocks (0.5 – 2.5 mM) that had been irradiated with a 420 nm LED (~ 5 min) (**Fig S2**).

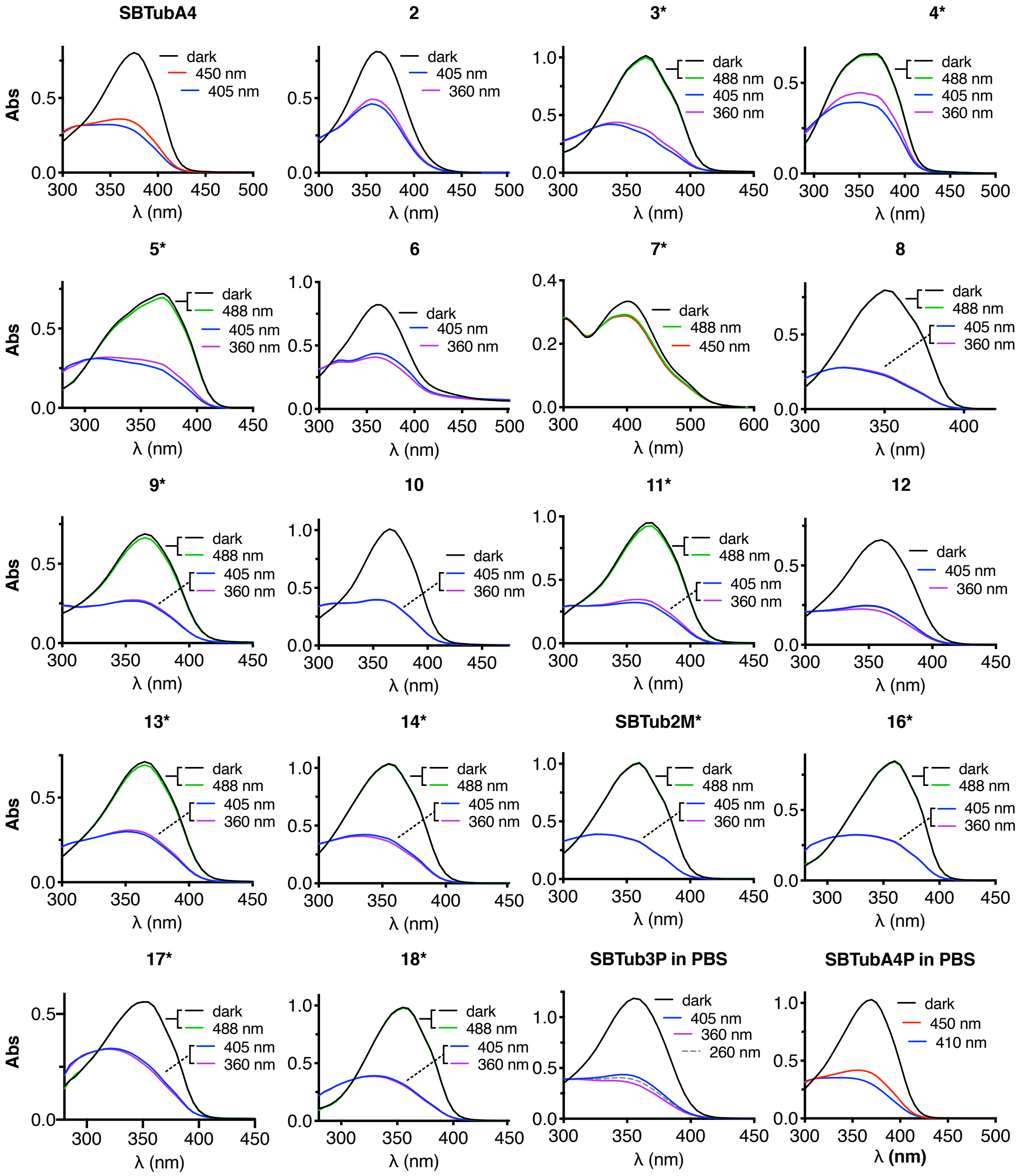

**Fig S1:** UV-Vis spectra of second generation SBTubs at various photostationary states, shows that the SBT scaffold is relatively unaffected by various functional residues, indicating that most of these newly synthesized SBTubs are also GFP-orthogonal. 25 μM in PBS, pH ~7.4, 10% DMSO unless indicated by asterisk* (50% DMSO) or "in PBS" (0% DMSO), room temperature, under closed air atmosphere.

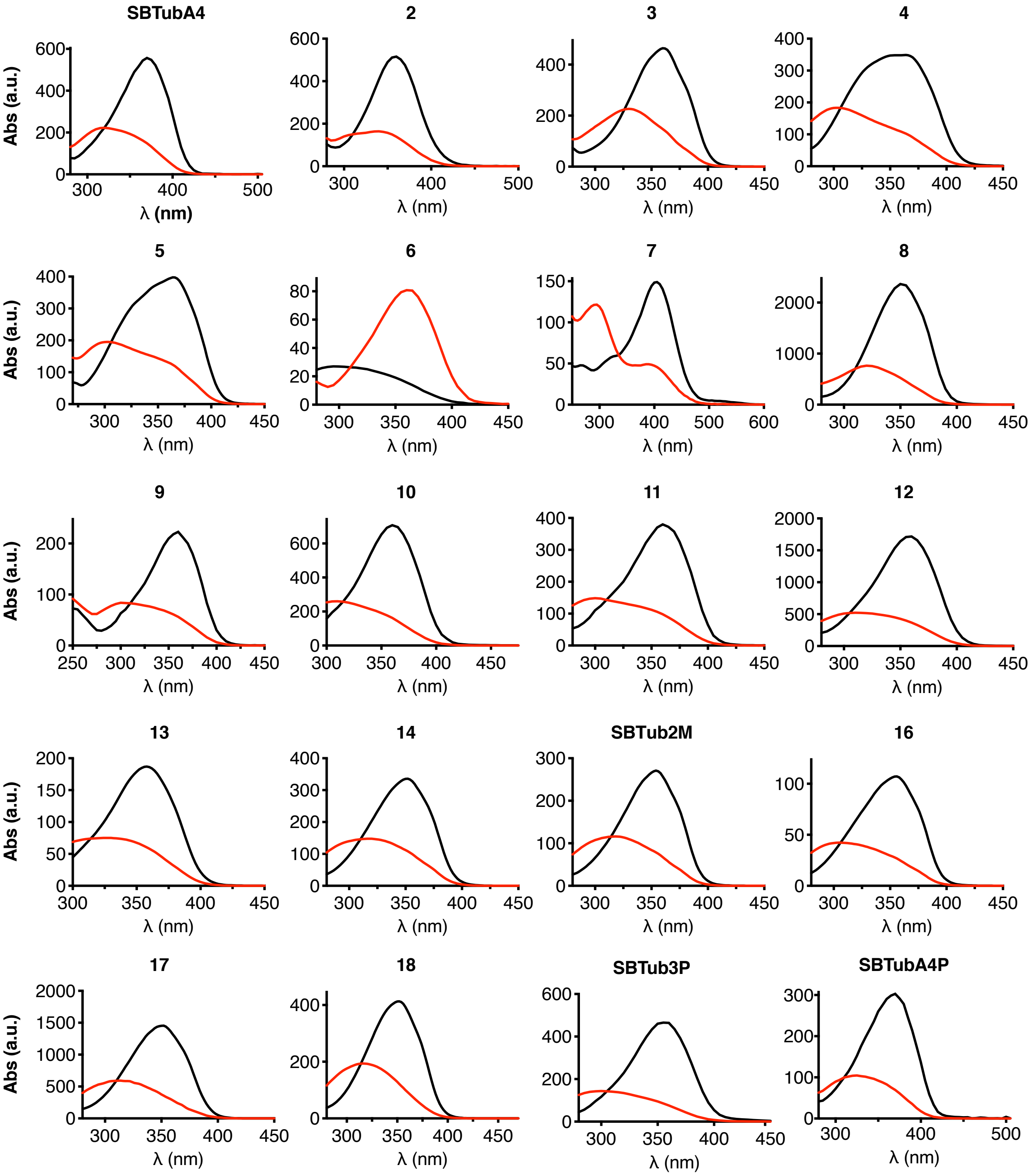

**Fig S2:** all-*E* and all-*Z* spectra of second generation SBTubs from inline HPLC-DAD

#### *Photoisomerisation: further discussion*

*Ortho*-hydroxy **6** and *para*-amino **7** were designed to test if these strong electron donors, that can also tautomerise to freely-rotatable quinoids (C-C single bond instead of C=C), can accelerate thermal *Z*→*E* relaxation to make better-reversible **SBTub**s, as well as inducing spectral redshifting. On azobenzenes and hemithioindigos, such strong *ortho*/*para*-electron-donating tautomerisable groups typically cause millisecond to second aqueous relaxation rates, often limiting their use on cytosolic targets. However, in cuvette, *Z*-**6/7** thermally relaxed only slowly compared to the timescales relevant to biological assays (halflives >> hours). *Z*-**7**'s stability was supported by lit-**7**'s appreciable cytotoxicity in cells, which is a remarkable result for the SBT scaffold as an alternative to azobenzenes and hemithioindigos. We had previously shown that a *p*-hydroxy SBT is photoswitchable in cuvette[7] but its design as non-binding control could not test whether its *Z*-isomer would also be stable enough for bioactivity in cells.[8,9] **7** showed that SBT photopharmaceuticals on cytosolic targets indeed tolerate *ortho/para* electron-donating tautomerisable groups. Furthermore, although the spectra of *E/Z*-**6** were similar to normal SBTs, **7** was redshifted by nearly 70 nm (**Fig S1-S2**). This may suit *para*-aminoSBTs as comparably absorbing alternatives to *p*-aminoazobenzenes that retain the often crucial *p*-amino function yet offer bistable isomerisation, which can be of interest in e.g. ion channel photopharmacology (where *p*-aminoazobenzenes are most widely employed). Since **6** was photoswitchable *in cuvette* yet showed no light-dependent cytotoxicity we suppose that it may suffer biochemical problems as known for catechols (e.g. metal complexation, oxidation to the *ortho*-quinone, etc[10]). In this study, as it was clear that neither **6/7** brought relaxation rate benefits, we did not pursue such derivatives further; but these results do showcase the unique compatibility of the SBT scaffold for bistable photopharmaceuticals where small polar hydroxy or amino groups (common groups for high-potency ligand-protein interactions) are needed in the phenyl *ortho* or *para* positions.

##### *Photostability*

The SBTubs' photostability to continuous UV illumination at >380 nm was typically excellent.

We also observed that primarily electron-rich SBTubs (**SBTubA4**, **Fig S3**) undergo unwanted photoreactions when short-wavelength irradiation (typically <350 nm) stimulates the S2←S0 transition, a band that is visible as a low-intensity shoulder to the principal S1←S0 band (**Fig S2-S3**; this band is particularly pronounced for aniline **7**). Therefore, for clean *E*→*Z* photoswitching, it is beneficial to use switching wavelengths $\lambda_{lit}$ that are usually 10-30 nm red-shifted from $\lambda_{max}(E)$. In contrast, continuous illumination with intense < 350 nm light caused slow degradation (ca. 10% lost during 10 min illumination with 360 nm band centre, ca. 25 nm FWHM, LED with nominal 200 mW/cm$^2$ optical output).

To characterise this photodegradation, electron-rich **SBTubA4** (25 $\mu$M in PBS:DMSO = 1:1) was irradiated with LEDs of emission maxima λ > 400 nm until corresponding PSSs were reached (reversible photoswitching; **Fig S3a**). Then applying 100 mW/cm$^2$ LED illumination with nominal band maximum 360 nm first gave rapid partial bulk *Z*→*E* photoisomerization to the ~360 nm PSS (**Fig S3b**: isosbestic point retained), then slower light-induced side reactions that destroy the ArC=CAr' chromophore (indicated by the drop of the isosbestic point, **Fig 3c**). Since this photodegradation does not occur at λ > 400 nm it should not pose a problem in biological assays when appropriate pure photoactivation wavelengths are used, such as the standard 405 nm laser (bandwidth < 1 nm) available on standard microscopy setups (see also previously performed photostability studies on SBTubs[7]).

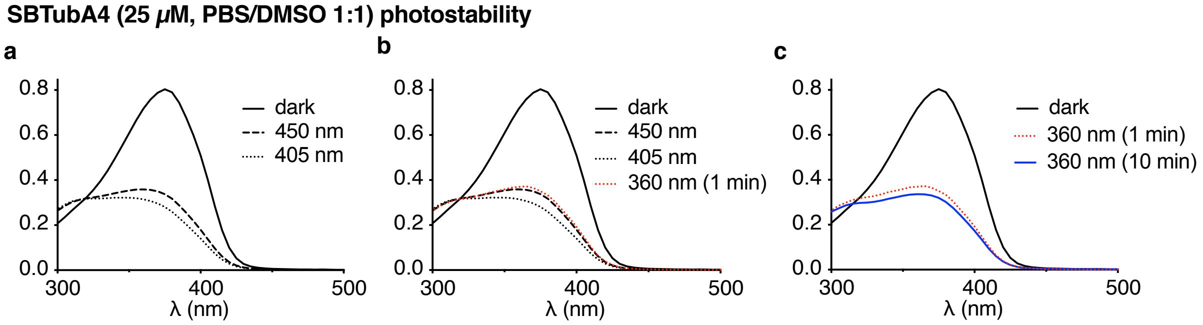

**Fig S3:** (a) Dark state spectrum, then PSS establishment with LED sources first at 450 nm then at 405 nm (note: LED bandwidth implies that wavelengths below 450 nm are substantially responsible for the photoswitching speed at nominal "450 nm"). (b) From 405 nm PSS, 360 nm LED was applied, reaching new PSS. (c) Under continued 360 nm illumination, the absorption profile drops, indicating side reactions that destroy the ArC=CAr' chromophore.

#### *Spontaneous relaxation of SBTubA4*

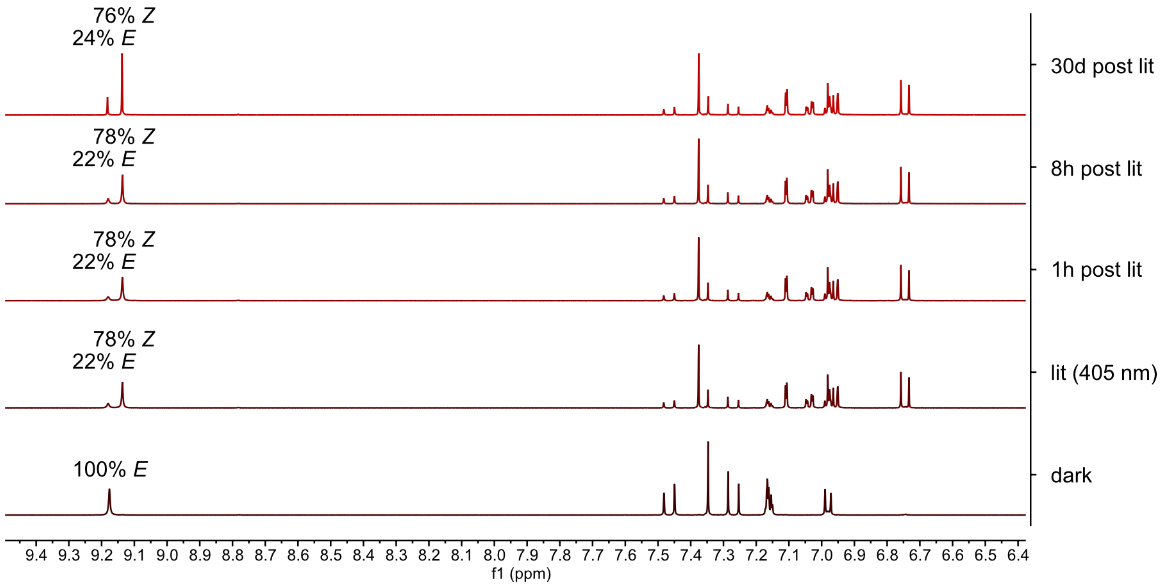

**Fig S4:** NMR of SBTubA4 showing the aromatic region. Illumination with 405 nm light reaches a PSS of ~80% *Z*-isomer, which is does not relax back over several days.

**SBTubA4** (6 mg) was dissolved in deuterated DMSO (0.6 mL) and photoisomerised with a 405 nm LED for 5 minutes to reach a PSS of ~80% *Z*-isomer as measured by NMR, subsequent measurements after storing the NMR tube in the dark for 1 h, 8 h (at 60°C) and 30 days (at room temperature) showed little to none spontaneous relaxation (**Fig S4**). (Side note: a pre-illuminated DMSO stock left in the dark in a fridge at 4°C was found >99% reverted back to the *E*-isomer after 9 months, giving an upper bound to the relaxation halflife in DMSO). Therefore, DMSO stocks of **SBTubA4** need more careful light-avoiding handling than **SBTub3** or **SBTub2M**, which both thermally revert back to 100% *E* when kept overnight at 60°C.

### Part C: Biological Data

##### *Tubulin polymerisation*

99% tubulin from porcine brain was obtained from Cytoskeleton Inc. (cat. #T240). The polymerisation reaction was performed at 5 mg/mL tubulin, in polymerisation buffer BRB80 (80 mM piperazine-N,N′-bis(2-ethanesulfonic acid) (PIPES) pH = 6.9; 0.5 mM EGTA; 2 mM $MgCl_2$), in a cuvette (120 µL final volume, 1 cm path length) in a Agilent CaryScan 60 with Peltier cell temperature control unit maintained at 37°C; with glycerol (10 µL). Tubulin was first incubated for 10 min at 37°C with "lit"- (405 nm-pre-illuminated; mostly-*Z*-) or dark- (all-*E*) **SBTub** (final concentration 20 µM) in buffer with 3% DMSO, without GTP. Then GTP was added to achieve final GTP concentration 1 mM (with mixing), and the change in absorbance at 340 nm was monitored for 15 min, scanning at 15 s intervals.

##### *Cell assay methods*

##### *General cell culture*

HeLa, Jurkat and A549 cells were maintained under standard cell culture conditions in Dulbecco's modified Eagle's medium (DMEM; PAN-Biotech: P04-035550) supplemented with 10% fetal calf serum (FCS), 100 U/mL penicillin and 100 U/mL streptomycin. Cells were grown and incubated at 37°C in a 5% $CO_2$ atmosphere. Cells were typically transferred to phenol red free medium prior to assays (DMEM; PAN-Biotech: P04-03591). Substrates and cosolvent (DMSO; 1% final concentration) were added *via* a D300e digital dispenser (Tecan). Cells were either incubated under "lit" or "dark" conditions; "lit" indicates a pulsed illumination protocol was applied by multi-LED arrays to create the *Z*-isomers of the compounds in *situ* in cells and then maintain the wavelength-dependent PSS isomer ratio throughout the experiment, as described previously.[11] Typical "lit" timing conditions were 75 ms pulses of ~1 mW/cm$^2$ applied every 15 s. "Dark" indicates that **SBTub** biostocks were used with an all-*trans* state, as determined by analytical HPLC, kept in the dark at 4°C, applied while working under red-light conditions, and cells were then incubated in light-proof boxes to shield from ambient light, thereby maintaining the all-*E*-isomer population throughout the experiment.

##### *Resazurin antiproliferation assay*

HeLa or A549 cells were seeded in 96-well plates at 5,000 cells/well and left to adhere for 24 h before treating with test compounds. *E*-SBTubs were added for 48 h (final well volume 100 µL, 1% DMSO; three technical replicates); the “cosolvent control” (“ctrl”) indicates treatment with DMSO only. Cells were then treated with Resazurin 150 mg/mL for 3 h. Fluorescence was measured at 590 nm (excitation 544 nm) using a FLUOstar Omega microplate reader (BMG Labtech). Absorbance data was averaged over the technical replicates, then normalized to viable cell count from the cosolvent control cells (%control) as 100%, where 0% viability was assumed to correspond to fluorescence signal in PBS only with no cells. Three independent experiments were performed. Data were plotted against the log of **SBTub** concentration ($log_{10}$([**SBTub**]) (M)) with mean and SD. For A549 cell line one experiment out of three is shown. With the exception of **6**, all SBTubs show light dependent cytotoxicity with $IC_{50}$ values ranging from mid nanomolar to mid micromolar concentrations in HeLa cells (**Fig S5**). Lead compounds **SBTub2M** and **SBTubA4** also show light dependent toxicity in A549 cells at low micromolar concentrations (**Fig S5**).

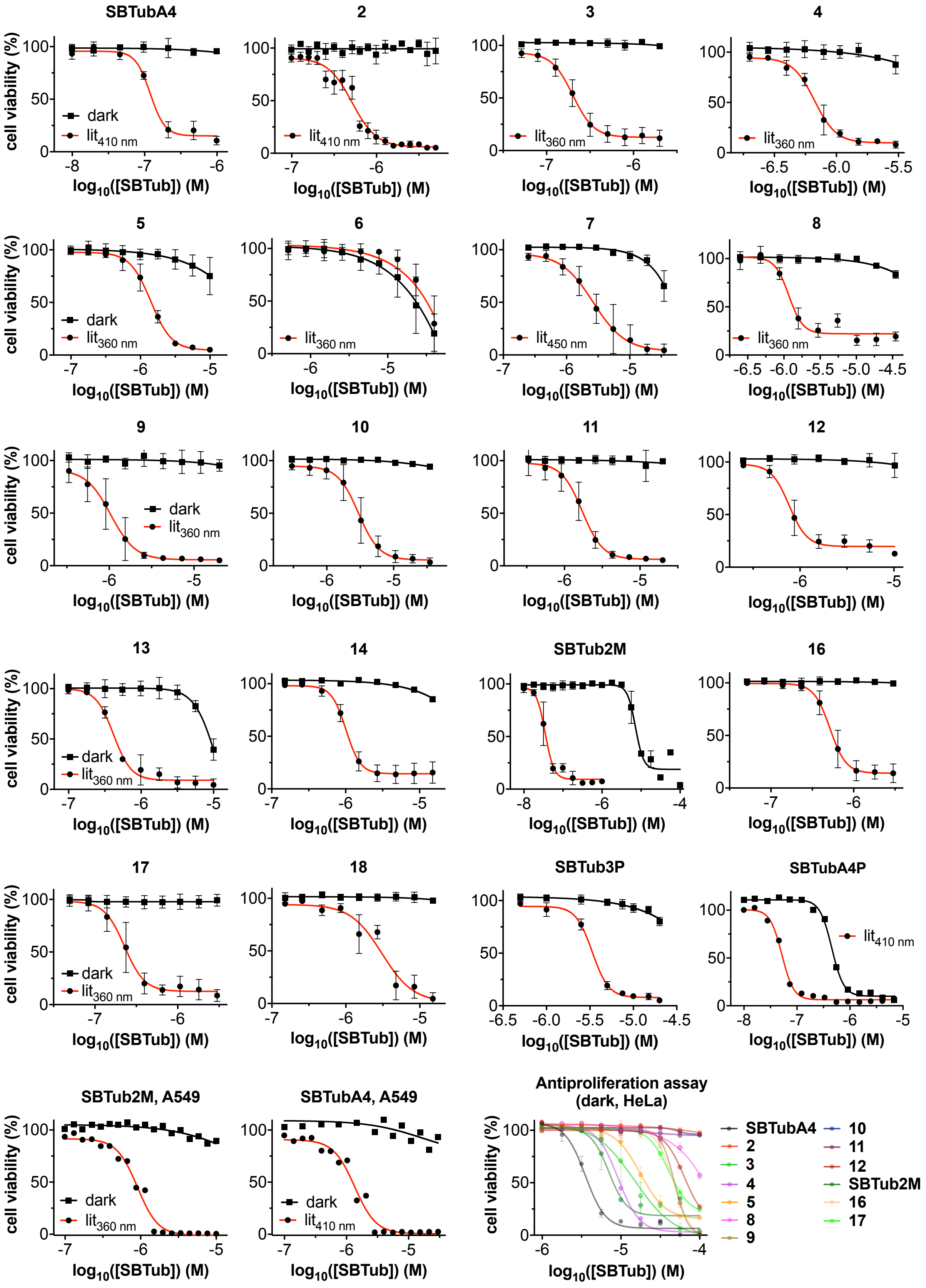

**Fig S5:** Antiproliferation assay of second generation SBTubs in HeLa cells and A549 cells for selected SBTubs. 40 h incubation; all-*E* dark conditions versus lit conditions with predominantly *Z*-isomer using low-power pulsed LED illuminations (75 ms per 15 s, <1 mW/cm$^2$); HeLa cells; N = 3; mean ±SD. A549 cells, one representative experiment out of three independent experiments shown. Resazurin antiproliferation assays were run of compounds in the dark up to 100 μM to determine the $IC_{50}$ of SBTubs in the dark.

#### *Cell cycle analysis*

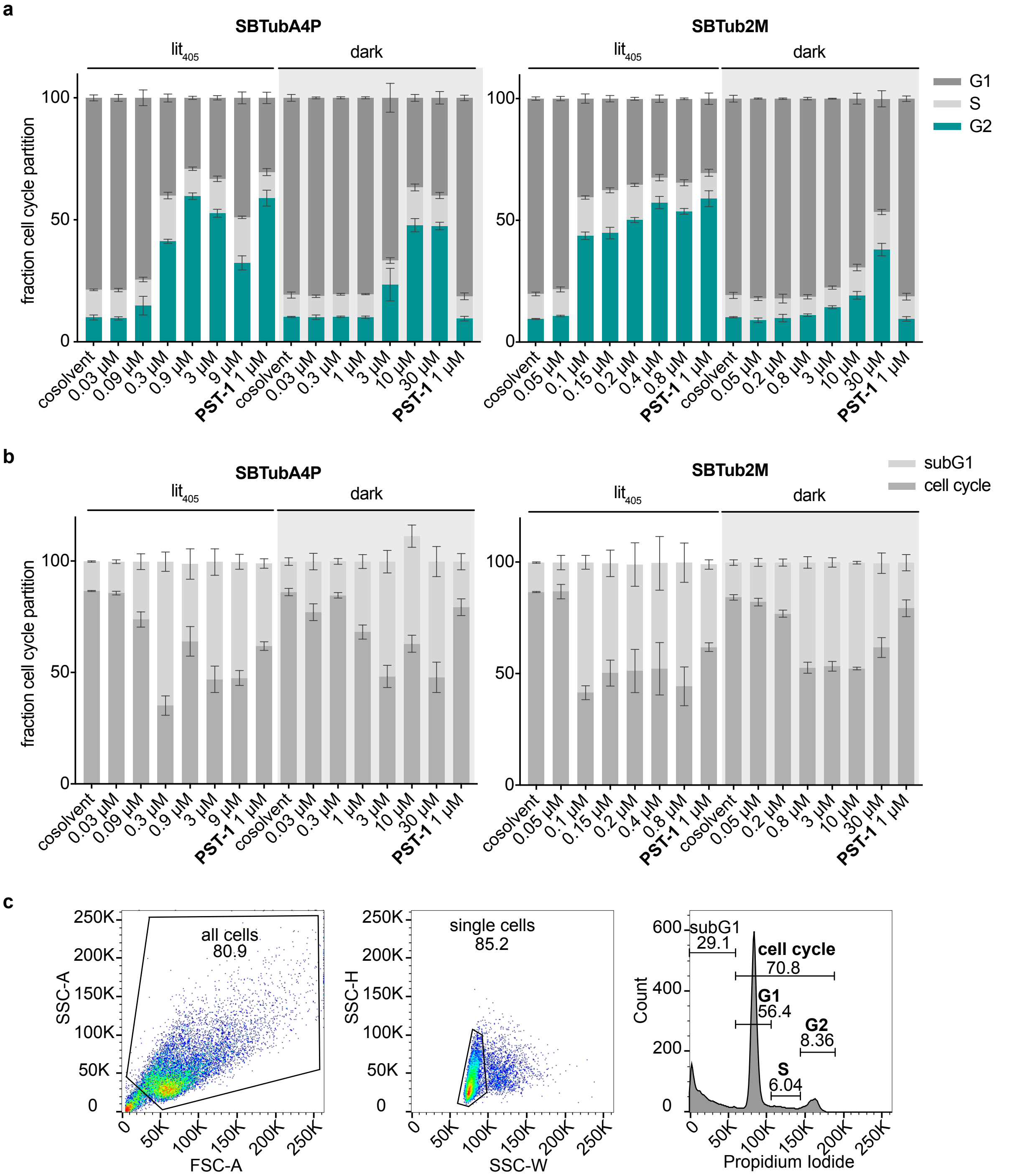

**Fig S6: a** Cell cycle partitioning of **SBTubA4P** and **SBTub2M** in Jurkat cells shows light dependent $G_2$/M arrest similar to photoswitchable positive control **PST-1** under lit conditions with 405 nm pulsing but no cell cycle effects when kept in the dark comparable to the DMSO only cosolvent controls. **b** Comparison of cells in subG1 (not shown in **Fig3b** and **Fig S6a**) compared to alive cells which were included in the cell cycle analysis. **c** gating and analysis strategy used in cell cycle quantification on a cosolvent treated sample.

*E*-**SBTub**s were added to Jurkat cells in 24-well plates (50,000 cells/well; three technical replicates, three biological replicates) and incubated under “dark” or “lit” conditions for 24 h. Cells were collected, permeabilized and stained with 2 $\mu$g/mL propidium iodide (PI) in HFS buffer (PBS, 0.1% Triton X-100, 0.1% sodium citrate) at 4°C for 30 min then analysed by flow cytometry using a LSR Fortessa flow cytometer (Becton Dickinson) run by BD FACSDiva software (version 8.0.1). 10,000 events per technical replicate were analysed; the PI signal per event (corresponding to cellular DNA content) was measured and cells were binned into sub-G1, G1, S and G2 phase according to DNA content using FlowJo software (version 10.7.1). Results (means of three technical replicates) from one experiment of three independent trials are shown (**Fig S6a-b**). Gating strategy shown in **Fig 6c**.

##### *Immunofluorescence staining*

HeLa cells were seeded on glass coverslips in 24-well plates (50,000 cells/well) and treated with SBTubs the next day under "dark" or "lit" conditions for 24 h. Cells were fixed and permeabilised in ice-cold methanol for 5 min, then washed and kept in PBS at 4°C until staining. Samples were equilibrated to room temperature and blocked with PBS + 1% BSA for 30 min. Cells were treated with primary antibody (1:200 rabbit alpha-tubulin; Abcam ab18251) in PBS/1% BSA/0.1% Triton X-100 overnight and with secondary antibody (1:500 goat-anti-rabbit Alexa Fluor 488; Abcam ab150077) in PBS/1% BSA/0.1% Triton X-100 for 1 h. Coverslips were mounted onto glass slides using Roti-Mount FluorCare DAPI (Roth) and imaged with a Zeiss LSM710 confocal microscope (CALM platform, LMU). Images were processed using the free Fiji software[12] and Affinity Designer (Serif) for clarification. Postprocessing was only performed to improve visibility.

##### *Live cell imaging*

HeLa cells were transfected with EB3-tdTomato using FuGENE 6 (Promega) according to manufacturer's instructions. Experiments were imaged on a Nikon Eclipse Ti microscope equipped with a perfect focus system (Nikon), a spinning disk-based confocal scanner unit (CSU-X1-A1, Yokogawa), an Evolve 512 EMCCD camera (Photometrics) attached to a 2.0× intermediate lens (Edmund Optics), a Roper Scientific custom-made set of Vortran Stradus 405 nm (100 mW), 488 nm (487 nm / 150 mW) and Cobolt Jive 561 nm (110 mW) lasers, a set of ET-DAPI, ET-GFP and ET-mCherry filters (Chroma), a motorized stage MS-2000-XYZ, a stage top incubator INUBG2E-ZILCS (Tokai Hit) and lens heating calibrated for incubation at 37°C with 5% $CO_2$. Microscope image acquisition was controlled using MetaMorph 7.7 and images were acquired using a Plan Apo VC 40× NA 1.3 oil objective. Imaging conditions were initially optimized to minimize tdTomato bleaching and phototoxicity for untreated cells. For compound treated acquisitions 6 $\mu$M **SBTubA4P** diluted in prewarmed cell medium was applied to cells in a dark room with only red ambient light, cells were protected from all ambient light after application of drug. Drug was incubated on cells for at least 1 min before commencing experiment. Comet count analysis was performed in ImageJ using the ComDet plugin (E. Katrukha, University of Utrecht, Netherlands, https://github.com/ekatrukha/ComDet).

##### *Quantification and statistical analysis.*

All relevant assays were done in independent biological replicates. All attempts at replication were successful and no data were excluded from analysis. Blinding was not performed as assay readout is mostly unbiased (plate reader, flow cytometry, Fiji/ImageJ plugins). Microscopic evaluation was performed independently by two separate scientists. Data were analysed using Prism 9 software (GraphPad). Two-tailed unpaired t tests were used in pairwise group comparisons; * was used for $P < 0.05$, ** for $P < 0.01$, *** for $P < 0.001$, **** for $P < 0.0001$.

##### *3D cell culture*

**Organoid growth conditions:**

Organoid culture followed a previously published protocol[13]. Freshly isolated human mammary gland epithelial cells from healthy women undergoing reduction mammoplasty were seeded in a collagen gel (collagen type I from rat tail, Cornings) of final concentration 1.3 mg/ml. The cells were cultured in mammary epithelial growth medium (PromoCell MECGM) with 3 $\mu$M Y-27632 (Biomol), 10 $\mu$M Forskolin (Biomol) and 0.5% FBS as additives for the first 5 days. Starting from day 5 the medium was replaced by MECGM medium with only 10 $\mu$M Forskolin as supplement. The medium was exchanged every second day. Experiments were carried out on organoids prepared from donor M21 (Age: 61 years, Parity: 1).

**Dose response curves:**

The effect **SBTubA4P** on organoid growth was first assessed in a dilution series (**Fig S7a-b**). Organoids were prepared and cultured based on the afore mentioned protocol up to day 5. On day 5 besides exchanging the medium **SBTubA4P** was added. For the lit culture conditions **SBTubA4P** was supplemented to achieve final concentrations ranging from 10 nM up to 600 nM. **SBTubA4P** was added to yield concentration between 125 nM and 400 nM for dark culture conditions. Medium was exchanged every second day. An additional washing step was carried out to completely remove the old compound from the collagen gel and avoid upconcentration. The organoids cultured under lit conditions were placed on LEDs emitting a 100 ms light pulse at 405 nm every 60 s. The dark control was shielded from light with aluminium foil. After 2 weeks the organoids were fixed with 4% PFA incubation for 15 min. Immunofluorescence staining was performed by permeabilizing cells using a 0.2% Triton X-100 solution followed by incubation with a blocking solution containing 10% donkey serum and 0.1 %BSA. Cells were stained for the presence of the proliferation marker Ki67 using primary monoclonal mouse antibody (ab238020, 1:200, Abcam) incubation at 4°C overnight succeeded by secondary antibody incubation with donkey anti-mouse IgG (A21202, 1:250, Invitrogen) at 4°C overnight. Cell nuclei were visualized using DAPI. Collagen gels were mounted on microscopy slides using mounting medium (18606-20, Polysciences).

The stained organoids were imaged using a Leica SP8 lightning confocal microscope and a HCX PL APO 10x/0.40 CS dry objective. Z-projections of the DAPI signal were generated to create a mask and determine the projected area of the organoids in the Z-projection (Fig 5a). Images were processed using Fiji software. Data Analysis was carried out using a Python Script. For the dose response curve, the area of the organoids was normalized by using the mean of the area from organoids cultured under normal culture conditions (**Fig S7a**).

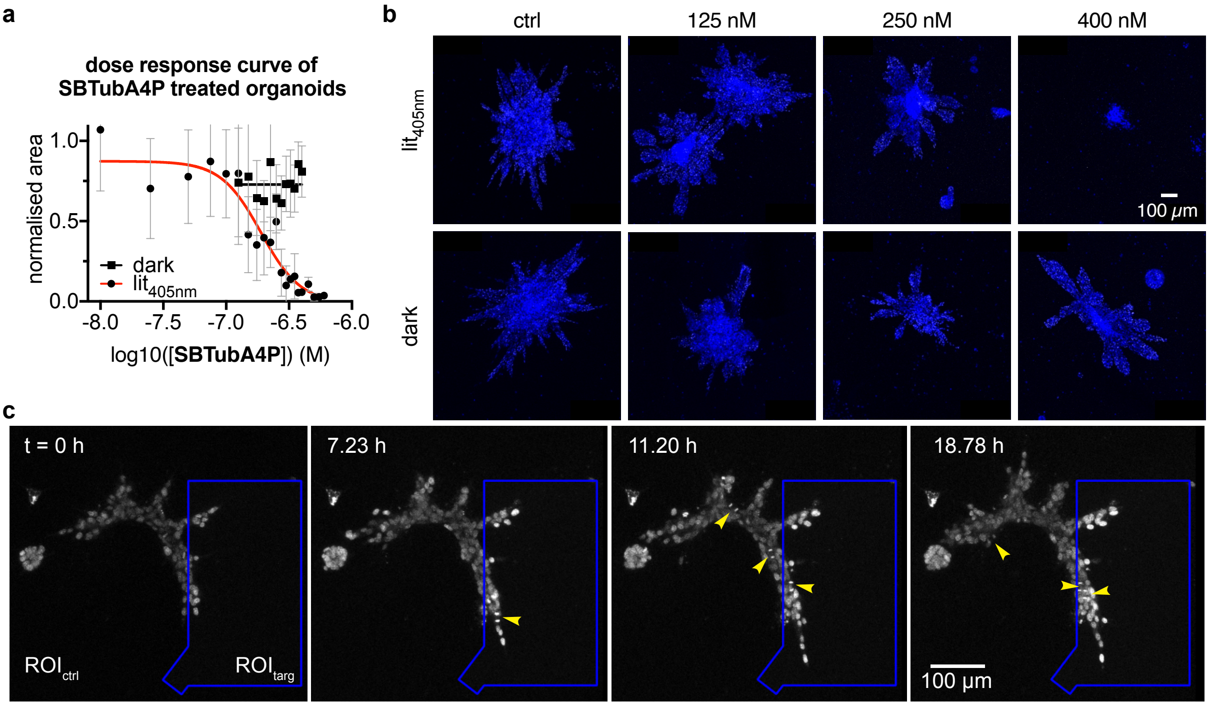

**Fig S7: a-b** Determination of **SBTubA4P** working concentration $[I]_{wc}$ in human mammary gland epithelial cells. **(c)** Imaging of organoids in the absence of **SBTubA4P** (no compound control) shows that branches are migrating and proliferating (yellow arrowheads) both within and outside the illuminated $ROI_{targ}$.

#### Imaging of organoid growth:

Live cell imaging was carried out on a Leica SP8 lightning confocal microscope using a HCX PL APO 10x/0.40 CS dry objective. Laser intensities at full power were as following: 405 nm (0.70 mW), 488 nm (4.50 mW) 552 nm (4.50 mW), 638 nm (0.22 mW). The $CO_2$ level and humidity were controlled using an ibidi gas incubation chamber. Organoids were imaged 8 to 10 days after seeding to interfere during the branch elongation phase. At this time point substantial branch invasion into the extracellular matrix was reported to take place[14]. Cell nuclei were labelled by addition of siRDNA (10 $\mu$M, Spirochrome AG) 3 h before starting the measurement. **SBTubA4P** was added to the medium to achieve a concentration of 200 nM. After addition of the compound the samples were protected from light to prevent activation of **SBTubA4P**. The excitation wavelength was set to 638 nm and filters were adjusted accordingly to record the emission light peaking at 674 nm for nuclei visualization. To fully capture the whole structure of the organoids several planes in the z-direction were selected. Organoid growth was monitored for 4 h to determine the region of interest in which the cells displayed the most invasive behaviour. Fiji software was used to further process the data. A python script was used to correct for translational drift of the organoids.

**Local inhibition of organoid growth:**

To locally inhibit organoid growth a region of interest was selected in an area where the cells displayed increased motility. The region of interest was set to be irradiated with light at 405 nm wavelength. The intensity for the 405 nm light was set to 20% (0.14 mW) in the region of interest. The size of the region of interest was varied depending on the cell motility in that area and the translational drift of the organoid observed during the preceding 4 h of measurement. 405 nm light irradiation in the region of interest was active throughout the whole z-stack of the organoids. The pixel dwell time was 2.09 $\mu$s/pixel. Therefore, scanning the designated regions of interest took approximately between 390 ms up to 520 ms for each frame of the z-stack. A complete z-stack of the organoids was acquired every 7 min. Cell motility was visualized by imaging the stained nuclei in and outside of the region of interest as described above. Fiji software was used to further process the data. A python script was used to correct for translational drift and to draw the outlines of the region of interest.

#### *Tissue and Animal Assays*

#### *Drosophilia melanogaster*

##### Fly strains and culturing

Flies were caged above solid fly nutrient plates and allowed to lay eggs for 2 h. Flies were then removed from the plates as larvae developed. Plates were incubated for a total of 72-96 h at 25°C before larvae were harvested for dissection.

The following transgenes and fluorescent markers were used:
*worGal4*, *UAS-mCherry*::*Jupiter*, *Sqh*::*GFP*[15]; *worGal4*, *UAS-mCherry*::*tubulin*[16]

##### Live cell imaging of drosophila neuroblasts and SBTub2M activation

Early third instar (72-96 h) larvae were dissected in Schneider's insect medium (Sigma-Aldrich S0146) supplemented with 1% Bovine Growth Serum (Cytivia SH3054102) and transferred to one well of a $\mu$-slide 8 well (Ibidi 80826) with 200 $\mu$L supplemented Schneider's plus 2% DMSO or 30 $\mu$M **SBTub2M** (2% final DMSO concentration). Brains were oriented with the lobes facing the coverslip. Once placed, brains were allowed to settle 15 min prior to transferring the slide to the microscope, where the brains were then allowed to settle an additional 15 min prior to the onset of imaging. Experiments with **SBTub2M** were performed in the dark or under red light.

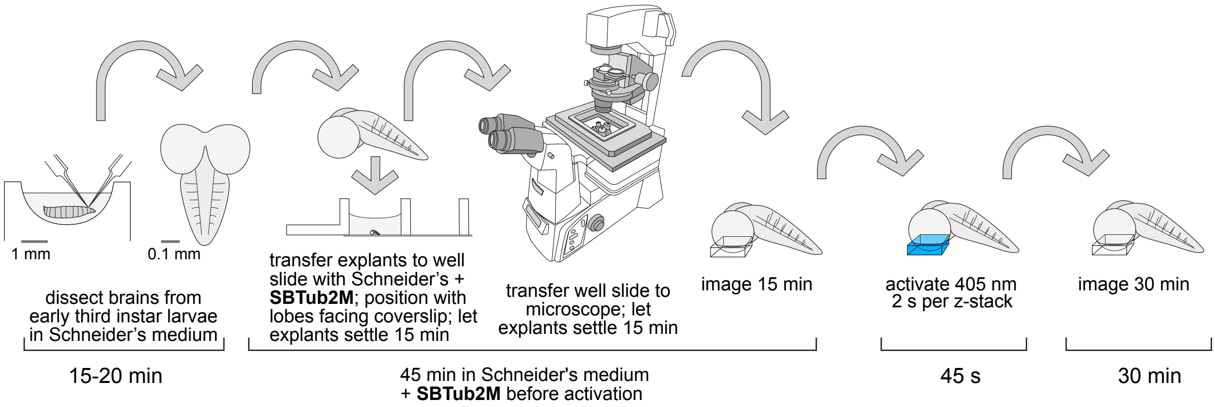

**Fig S8:** Experimental setup for photoswitching during live cell imaging of **SBTub2M** in fly brain explants.

Brain lobes were imaged with an Intelligent Imaging Innovations (3i) spinning disc confocal system, consisting of a Yokogawa CSU-W1 spinning disc unit and two Prime 95B Scientific CMOS cameras. A 60x/1.4NA oil immersion objective mounted on a Nikon Eclipse Ti microscope was used for imaging. 20 $\mu$m stacks with a 1 $\mu$m z-spacing at the top of the neural lobe were acquired every 15 s for 15 min before 405 nm activation and for 30 min following activation. mCherry::Jupiter and mCherry::tubulin were imaged for 100 ms at 561 nm (100 mW) with 10% laser power and Sqh::GFP was imaged at 488 nm (150 mW) for 100 ms with 20% laser power. Activation was performed by imaging the 20 μm stack for 2 s/stack with a 405 nm laser (100 mW) at 40% laser power.

Movies and extended z-stack projections were generated using Imaris v.9.5.1 and 9.2.1. Movies were annotated using FIJI.

##### Centrosome intensity measurements

Centrosome intensity was calculated with Imaris v.9.5.1 using the median intensity of centrosomes that were manually tracked for 2 min prior to activation and up to 8 min following activation using spots with a 2 μm diameter. Raw data for each track were exported to Microsoft Excel. For each centrosome, the signal intensity for each timepoint was normalized by dividing by the maximum observed signal intensity for each centrosome during the two minutes prior to activation. We used built-in functions in Excel to calculate the average and standard deviation of all of the normalized individual observations per timepoint. These data were plotted in Excel and edited for style in Adobe Illustrator.

#### *Xenopus tropicalis*

*X. tropicalis* embryos were generated according to standard methods by in vitro fertilization.[17] Briefly, a priming step is performed by injecting human chorionic gonadotropin (hCG) into the dorsal lymph sac of sexually mature *X.tropicalis* females (50 μL of a 200 U/mL solution). Frogs are then left for ~20 h in normal housing conditions separated from males. Boosting is completed by injecting 70 μL of a 2000 U/mL hGC solution in female frogs. ~3 h after injection females are gently squeezed to lay eggs, which are collected and fecundated in vitro using frozen sperm. After fecundation, embryos are dejellied with 2% L-cysteine prepared in 0.1x Marc's Modified Ringers (MMR). 1x MMR contains 100 mM NaCl, 2 mM KCl, 1 mM $MgSO_4$, 2 mM $CaCl_2$, 5 mM HEPES, pH 7.8. Embryos are kept in 0.1x MMR in Petri dishes surface-coated with 1% agarose. Temperature is kept constant at 23–25°C. Selection of embryos at 2 cell-stage or at the blastula stage was performed under the stereo microscope.

<u>SBTub4AP incubation/photoactivation:</u> Selected embryos either at the 2-cell or the blastula (>64-cell) stage (spherical shape ca. 1 mm diameter) were incubated with **SBTubA4P** diluted in 0.1x MMR for 1 h in the dark. The manipulation of the compound and embryos were always performed under green light to prevent photoactivation by ambient light. Embryos were selected with a glass pipette and transferred to new dishes (2 mL volume) containing 0.1x MMR without **SBTubA4P** as a washout procedure; the total residual volume of original medium transferred with the embryos was estimated at <50 μL per dish, 10 embryos were

transferred to an imaging RC-25 chamber (Warner Instruments, Hamden, CT) immediately after washout.

For photoactivation, the chamber was mounted on an inverted Olympus IX-50 microscope using a Hg arc lamp (100 W) as light source. Each embryo was illuminated individually *via* the epifluorescence port with 410 nm light (excitation filter D410/40, Chroma) for 5 s. Light was applied through a 20X objective (LCPlanFl 20X/0.4, Olympus, Japan). The diaphragm was fully open to illuminate a zone of diameter ca. 1 mm, i.e. one embryo was fully illuminated, but not the surrounding washout medium. A first round of illumination was followed by a second cycle, which was applied in a time interval of 5 min. Embryos were returned to dishes to monitor their development.

Micrographs of embryos were taken using an AxioCam ERc 5s (Carl Zeiss, Jena, Germany) mounted on a stereo microscope (Leica, Wetzlar, Germany). Movies to assess the movement of hatchling embryos were obtained with a webcam controlled by Image J Webcam Capture Plugin.

Embryonic development was quantified using a morphological (1) and a functional assay (2). (1) Long/short axis ratio: Morphology was evaluated by measuring the ratio between the longest and shortest axis of the embryos. During the initial 48 h of normal development embryos dramatically change their shape and transition from being cell spheres to tadpoles. This characteristic elongation can be quantified by calculating the ratio between the longest and shortest axis of the embryo (Feret's diameter in Image J) (**Fig 6b** from 2-cell-stage treatment, (**Fig S9b-d**) from blastula treatment). The suppression of this elongation reflects a drug's ability to interfere with normal cell divisions and to stop the growth of embryos.

(2) The functional assay was carried out by investigating sensorimotor behaviour. Hatchling tadpoles start to move in reaction to mechanical stimuli with a pipette tip.[18] A qualitative analysis was performed by calculating the proportion of the tadpoles that moved after mechanical stimuli and those that remained still. The ratio of the proportions of tadpoles incubated with the molecule, **SBTubA4P** or **PST-1P**, in lit or dark conditions was used to quantify the light-dependency effect of **SBTubA4P** and **PST-1P** in advanced stages of development (**Fig S9e**).

More than 80% of the tadpoles raised in control conditions or with 5 or 25 $\mu$M **SBTubA4P** in the dark consistently showed this behaviour 30 h post-fertilisation (**Movie SXX**). However, the proportion of tadpoles moving in response to mechanical stimulation decreased to about half if 5 or 25 $\mu$M **SBTubA4P** was activated at 410 nm light. The defects in the processing of sensorimotor information might reflect alterations in the nervous system. As this is generated by the ectoderm, the more external layer of the embryo, it is possible that the greater exposure of ectodermic cells to **SBTubA4P** causes defects in the nervous system. The action of **PST-1P** was compared to the effects of **SBTubA4P**. Application of **PST-1P** at 5 or 25 $\mu$M did not modify the long/short axis ratio of the embryos. With 25 $\mu$M the motion was suppressed both in the dark and after 410nm illumination, while with 5 $\mu$M neither lit nor dark suppressed motion.

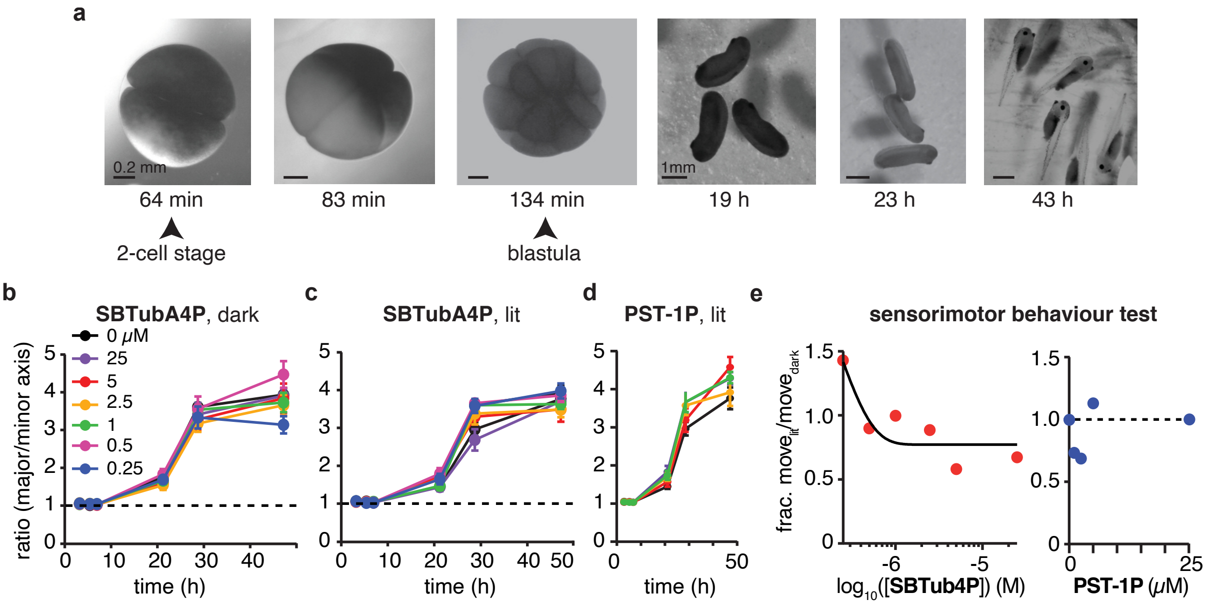

**Fig S9:** Effect of **SBTubA4P** on the development of *X. tropicalis* tadpoles. **a** Images of *X. tropicalis* embryos at the indicated developmental times post-fertilization. Embryos were exposed to varying **SBTubA4P** concentrations during 1 h, starting at the 2-cell stage (protocol 1) or at the blastula stage (protocol 2). **b-d** Plots evaluating the elongation of embryos during development show that neither **SBTubA4P** or **PST-1P** cause obvious morphological defects. Each dot is the mean of 10 embryos. Bars indicate s.e.m. on embryonic development using protocol 2. **e** The ratio between the proportion of embryos illuminated at 410 nm that moved after mechanical stimulation and the proportion of those kept in the dark is modified as a function of **SBTubA4P** and **PST-1P** concentration. Physiological responses are affected in SBTubA4P treated embryos. For **PST-1P** treated embryos no difference was observed between embryos kept in the dark or exposed to 410 nm light on the physiological level.

This suggests that **PST-1P** did not act light-dependently in the embryos, but can be light-independently toxic (**Fig S9d**). Note that 25 $\mu$M is ca. 16× the $[I]_{WC}$ of **PST-1P**[19], and 19× the $[I]_{WC}$ of **SBTubA4P**, so these concentrations could have been thought to deliver roughly equivalent biological effects. Analysing the reasons for azobenzene failure is outside the scope of this paper, but reductive *Z*-diazene degradation[20–22] in this highly metabolically active developing animal seems a likely culprit (which may have inspired the prior *in vivo* use of one azobenzene at 1000× its $[I]_{WC}$[23]); at any rate, the novel SBT photoswitch scaffold is far more robust against such degradation[21].

##### *Danio rerio*

##### Zebrafish strains and maintenance

Zebrafish (Danio rerio) were maintained according to the guidelines of the local authorities under licenses GZ:565304/2014/6 and GZ:534619/2014/4. Zebrafish WT AB* (ZFIN ID: ZDB-GENO-960809-7) embryos were used for all *in vivo* experiments. All embryos were kept in E3 medium (0.63 g/L KCl, 14.0 g/L NaCl, 1.83 g/l $CaCl_2 \cdot 2H_2O$, 1.99 g/l $MgSO_4 \cdot 7H_2O$, pH 7.4) and only for use in confocal microscopy experiments treated with 0.003% Phenylthiourea (PTU, Sigma-Aldrich, St. Louis, MO) before 24 hpf to inhibit melanogenesis. Zebrafish embryos used in all experiments were held at 28°C for upbringing and incubation times.

##### Determining compound toxicity in zebrafish

For determining toxicity and permeability of **SBTubA4P**, **SBTub3P** and **PST-1P**, zebrafish embryos were dechorionated after 24 hpf or 48 hpf and placed in 12 well tissue culture plates (VWR, Radnor, PA) in 2 mL E3 medium, and treated with either: 25 μM **SBTubA4P**, 1 μM **SBTubA4P**, 25 μM **SBTub3P**, 25 μM **PST-1P**, 0.25% DMSO only, or no treatment (vehicle control). Handling of the compounds was performed in a dark room under red lighting. All treatments were performed as both dark and as illuminated incubations, for 24 h. Global UV illumination was carried out by placing the tissue culture plates on a pulsed 24-LED array (395 nm; previously described[19]), emitting a 1 second UV pulse every five minutes in an otherwise dark incubator. For dark incubation the tissue culture plates containing treated embryos were wrapped in aluminum foil to shield them from light.

For endpoint imaging and evaluation (**Fig S10**), embryos were treated with 0.3 mg/mL Tricaine (ethyl 3-aminobenzoate methanesulfonate 98%, Sigma-Aldrich) for immobilization during the procedure. Imaging and counting of phenotypical abnormalities were performed between 52-54 and 76-78 hpf (for 24 hpf and 48 hpf embryo treatments, respectively). Representative images were recorded using a microscope camera (MC170 HD, Leica Microsystems, Wetzlar, Germany) mounted on a Leica M125 stereomicroscope and the LAS V4 software.

One-day old fish treated with **SBTubA4P** (25 and 1 μM) and **SBTub3P** (25 μM) showed light dependent phenotypical changes from the untreated and vehicle control. Two-day old fish did not show any morphological changes except when treated with **SBTubA4P** (25 μM) under lit conditions. Azobenzene-based tubulin inhibitor **PST-1P** did not show any phenotypical changes up to 25 μM under lit conditions. No phenotypical changes were observed for fish kept in the dark except for slightly curved tails in fish treated with **SBTubA4P** (25 μM).

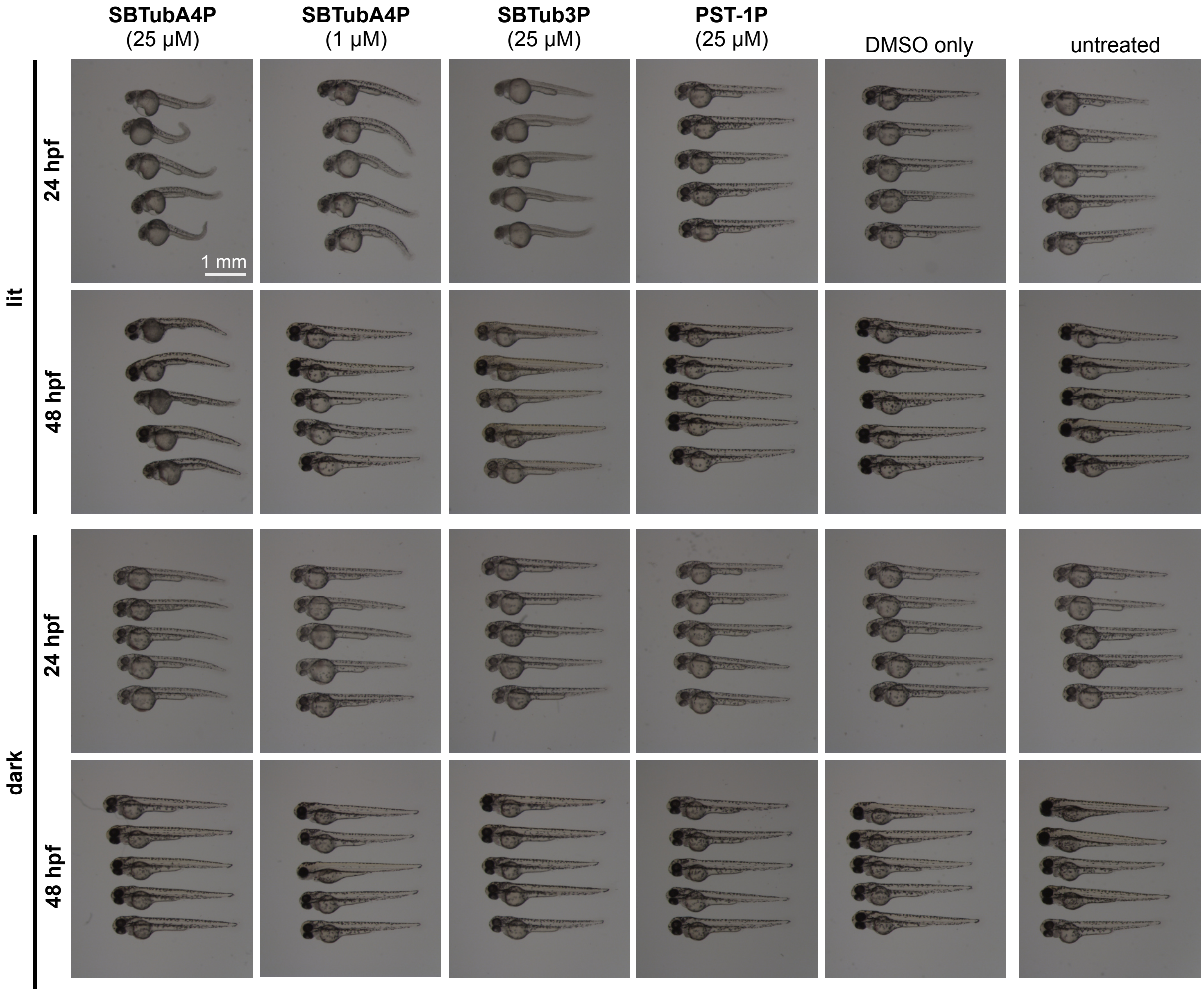

**Fig S10:** Determining toxicity in zebrafish embryos: 25 µM **SBTubA4P**, 1 µM **SBTubA4P**, 25 µM **SBTub3P**, 25 µM **PST-1P**, 0.25% DMSO (vehicle control) and no treatment.

#### Targeted inhibition of microtubule dynamics in vivo

For targeted activation of SBTubA4P in embryonic zebrafish to demonstrate inhibition of microtubule dynamics, WT AB* zebrafish embryos were microinjected with 20 pg each of pSK_H2B-mRFP:5xUAS:EB3-GFP[24] and pCS_KalTA4[25] plasmid DNA. Injections were performed with glass capillaries pulled with a needle puller (P97, Sutter Instruments, Novato, CA), mounted onto a micromanipulator (World Precision Instruments Inc., Berlin, Germany) and connected to a microinjector (FemtoJet i4, Eppendorf, Hamburg, Germany). Embryos were dechorionated and incubated with **SBTubA4P** or left untreated as control wrapped in aluminum foil for 4 h, starting from 26 hpf. After the incubation period, embryos were embedded in 1.2% low-melting agarose (Agarose Type IX-A, Ultra-low Gelling Temperature, Sigma-Aldrich) dissolved in E3 medium on #1.5 Glass Bottom Dishes (Cellvis, Mountain View, CA) under red light. Targeted activation of **SBTubA4P** *in situ* and imaging was performed using a SP8 confocal microscope (Leica Microsystems, Wetzlar, Germany). The bleachpoint function was used to target single EB3-GFP positive cells for photoactivation, while time-lapse imaging was continued to capture inhibition and recovery of microtubule dynamics over time. All photoactivation experiments at the confocal microscope were conducted at 28°C using a temperature control system (The Cube 2, Live Imaging Services, Basel, Switzerland). Images were recorded using the confocal microscope in conjunction with the LAS X software (Leica Microsystems). Videos were edited and labeled with ImageJ. Comet count analysis was performed by J.C.M.M. in ImageJ using the ComDet plugin (E. Katrukha, University of Utrecht, Netherlands, https://github.com/ekatrukha/ComDet).

##### Laser power and intensity calculations

Laser powers were measured with a 40x water objective (NA 1.1) using a Nova II power meter (Ophir, North Logan, USA). Using a 40x objective the beam diameter was calculated using the following formula: 1.22 x 405 nm/1.1 = 449 nm and the illuminated area of the bleachpoint is 158478 $nm^2$. Intensity for the bleachpoint experiments was calculated at 18900 $W/cm^2$.

#### Part D: Protein Crystallisation

##### *Protein crystallisation materials and methods*

##### Protein production, crystallization and soaking.

The DARPin D1 was prepared as previously described by Pecqueur et al.[26]. Bovine brain tubulin was purchased from the Centro de Investigaciones Biológicas (Microtubule Stabilizing Agents Group, CSIC, Madrid, Spain). The tubulin-DARPin D1 complex (TD1)[26–28] was formed by mixing the respective components in a 1:1.1 molar ratio.

The TD1 complex was crystallized overnight by the hanging drop vapor diffusion method (EasyXTal plates from QIAGEN, drop size 2 µL, drop ratio 1:1), at a complex concentration of 9.8 mg/mL and at 20°C with a precipitant solution containing 21% PEG 3000, 0.2 M ammonium sulfate and 0.1 M bis-tris methane buffer, pH 5.5. All crystallization drops were subsequently hair-seeded with crystalline material obtained from a previous PEG screen. Grown crystals were washed from the plate with precipitant solution and transferred into 0.6 mL centrifuge tubes in order to start batch-crystallization (overnight, 20°C). The average size of obtained batch-crystals (monoclinic needles) was about 120x10x5 µm.

Crystals were transferred into new precipitant solution drops containing 1.7 mM of **SBTub2M** compound (dissolved in DMSO) and 10% Glycerol. Crystals were soaked and illumined for 60 min at 385 nm to generate *cis*-**SBTub2M** *in situ* before being frozen in liquid nitrogen for X-ray diffraction data collection.

#### Data Collection, processing and refinement.

Data were collected at beamline X10SA (PXII) at the Swiss Light Source (Paul Scherrer Institut, Villigen, Switzerland). The X-ray beam spot was 30 x 15 µm², the flux was 7 x $10^{11}$ photons/s (100% beam intensity, 30 µm aperture). The data were collected with an exposure time of 0.05 s and an oscillation range of 0.1° per frame (see supplementary table S1). For TD1-SBTub2M, the data of three single crystals (210°, 180°, 180°) were merged. Data processing was done with XDS[29]. Due to anisotropy, the data were scaled, merged and corrected using the Staraniso server[30] (http:// staraniso.globalphasing.org/). The structures were solved by molecular replacement using PDB: 5NQT as a search model[27]. The ligands and restraints were generated with the grade server[31] (http://grade.globalphasing.org/) using their SMILES annotation. The structures were then refined iteratively in PHENIX[32] with manual editing cycles in COOT[33].

**Supplementary table S1:** Crystallographic statistics for *cis*-**SBTub2M** bound to tubulin-DARPin D1 (TD1) complex. Related to Methods X-ray diffraction data collection, processing and refinement.

| **Data statistics** | |
|---|---|
| Space group | $P2_1$ |
| Unit cell (a b c β) | 73.68 91.61 83.11 97.39 |
| Wavelength (Å) | 1.0 |
| Resolution (Å) | 45.81 - 2.36 |
| $R_{pim}$ (%) | 10.8 (49.1) |
| I/σI | 4.7 (1.6) |
| Spherical completeness (%) | 58.3 (8.7) |
| Ellipsoidal completeness (%) | 92.9 (66.1) |
| Ellipsoidal truncation resolution limits (Å for a*, b*, c*) | 2.36 3.15 2.81 |
| Multiplicity | 11.7 (10) |
| $CC_{1/2}$ | 0.974 (0.517) |
| **Refinement statistics** | |
| Resolution | 45.81 - 2.36 |
| Number of reflections | 27889 |
| $R_{work}$ / $R_{free}$ | 20.84% / 25.84% |
| Ramanchandran favoured | 93.90% |
| Ramanchandran outliers | 0.1% |
| R.m.s.d bond lenght (Å) | 0.0026 |
| R.m.s.d bond angles (°) | 0.572 |
| PDB code | |

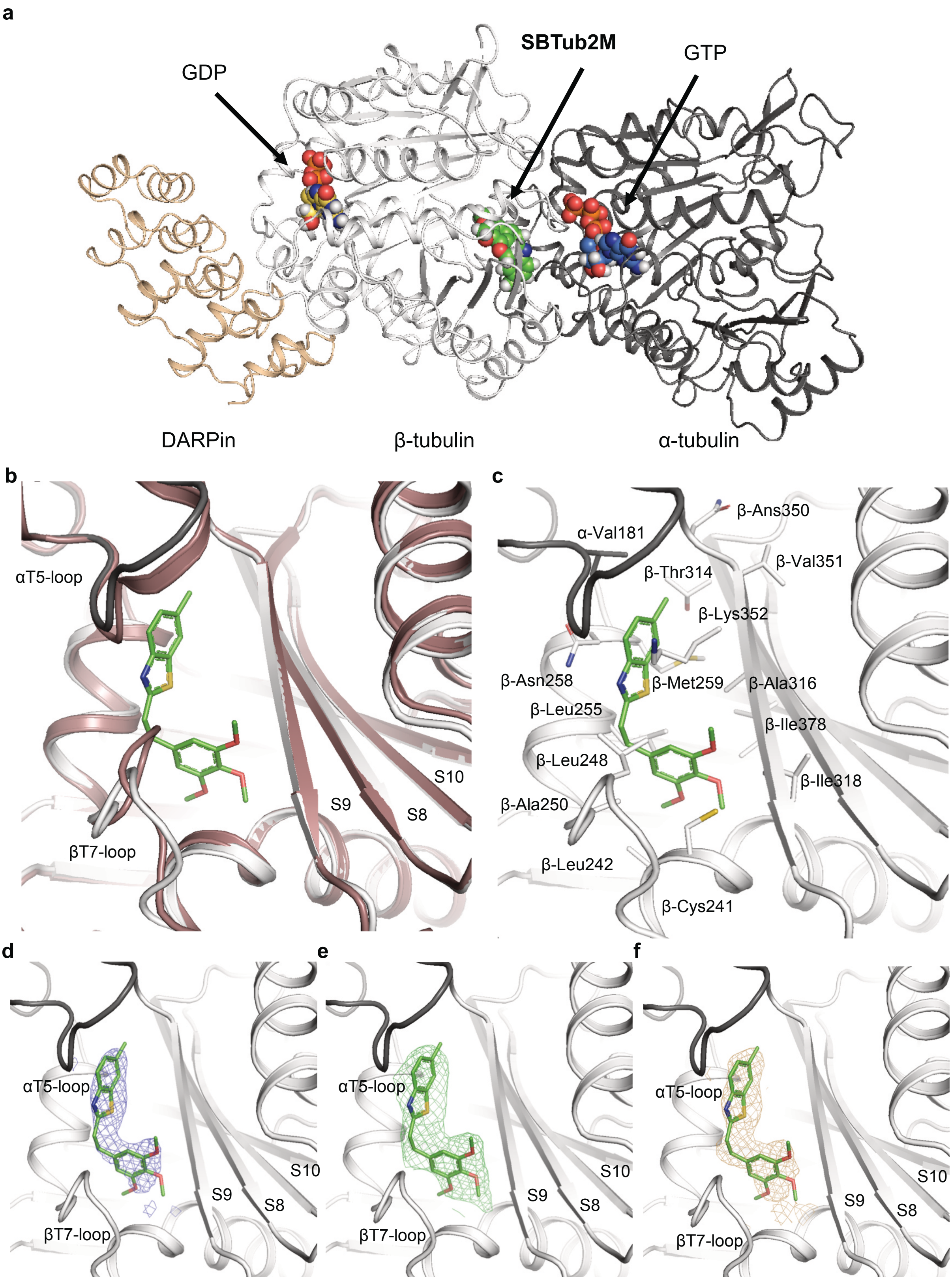

**Fig S11: a** Structure overview TD1:*Z*-**SBTub2M**. Structures are shown in cartoon representation, ligands shown as sphere. DARPin = wheat, β-tubulin = white, α-tubulin = grey. **b** Overlay of TD1:*Z*-**SBTub2M** and the ligand free structure (red, PDB 5NQU). **c** Close-up view of the colchicine-site showing *Z*-**SBTub2M**. Tubulin shown in cartoon representation; interacting residues are labeled and shown in stick representation. Oxygen = red, nitrogen = blue, sulfur = yellow. **d** *Z*-**SBTub2M** with 2FoFc map at 1.5σ shown in blue **e** *Z*-**SBTub2M** with simple omit map at 3.0σ shown in green **f** *Z*-**SBTub2M** compound with simulated annealing omit map at 2.5σ shown in yellow.

### Part E: NMR Spectra

#### 1H-NMR of SBTubA4:

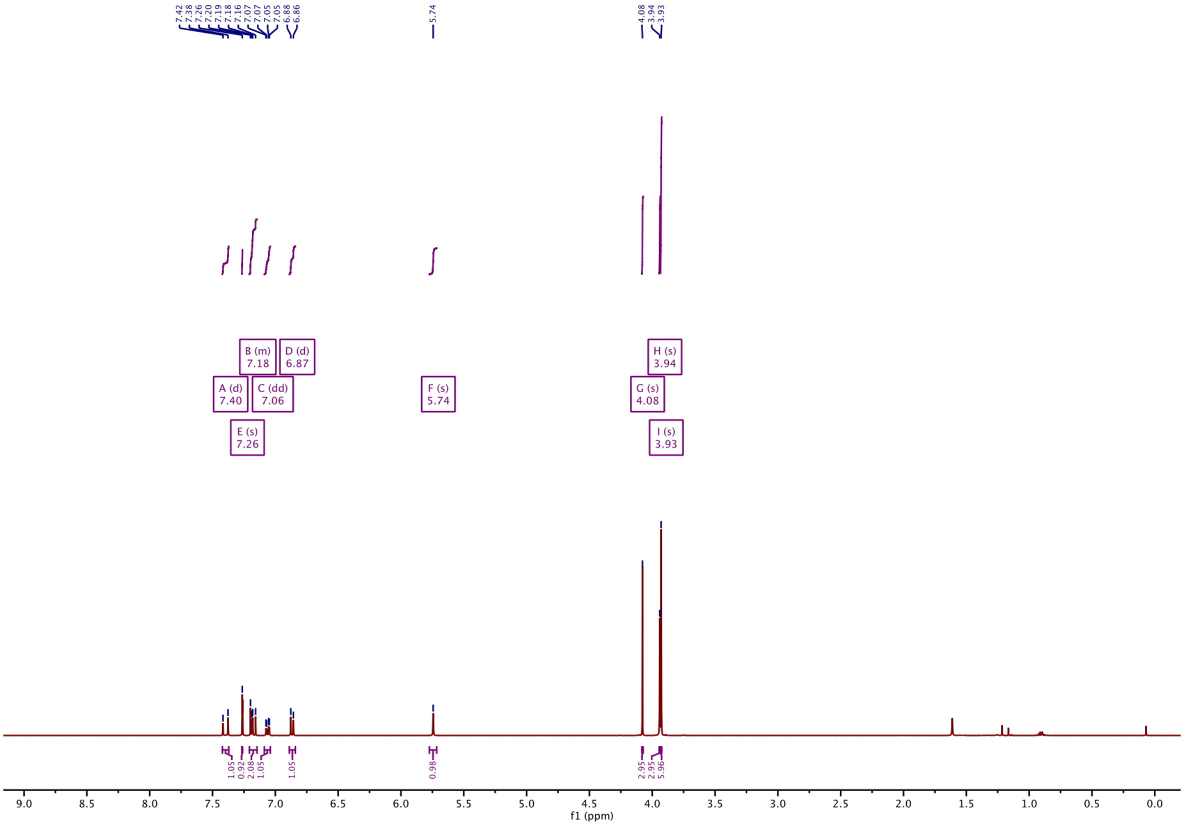

#### 13C-NMR of SBTubA4:

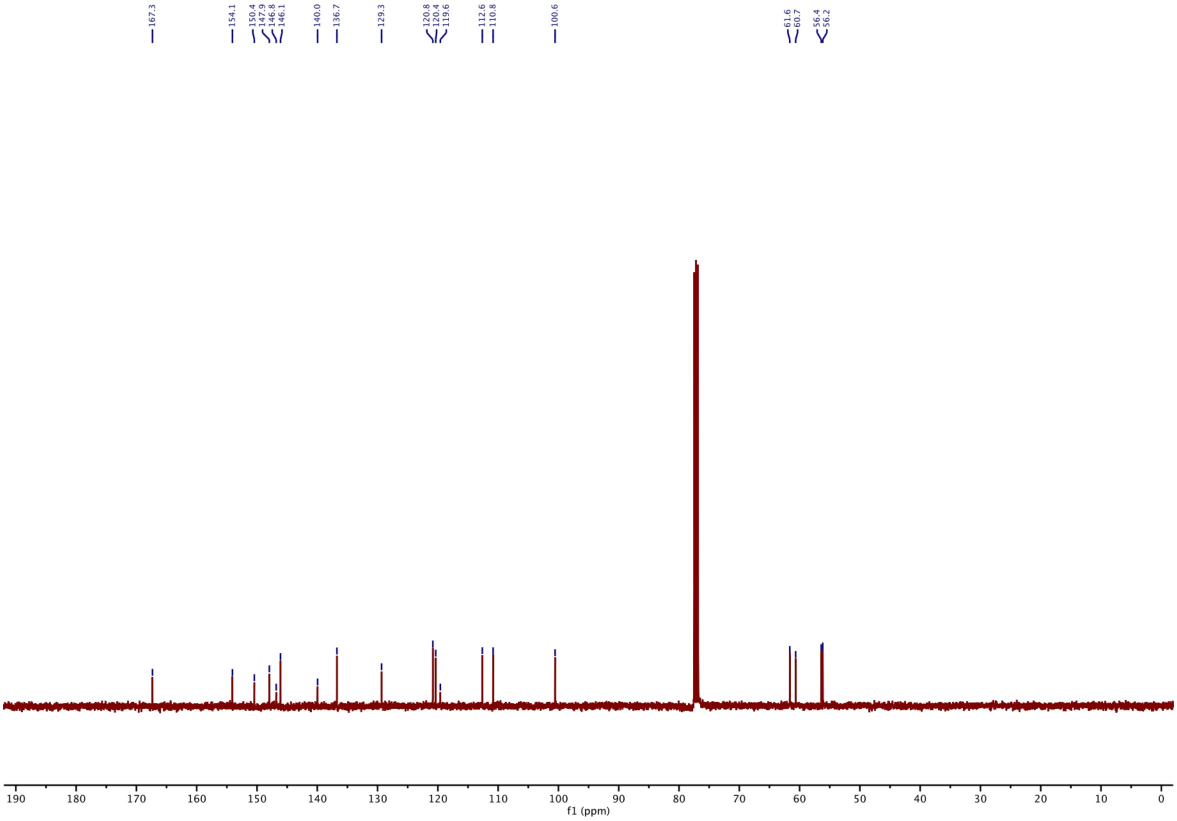

#### [1]H-NMR of 2:

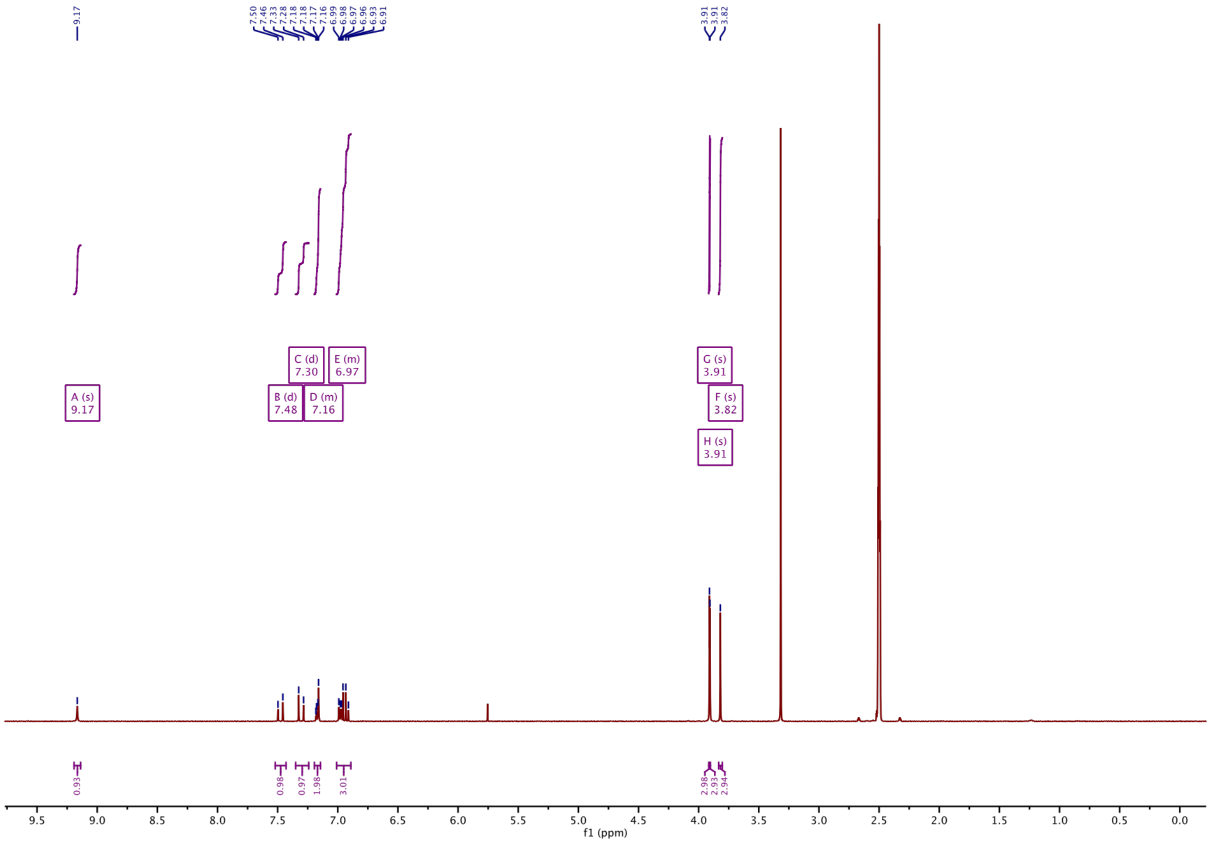

#### [13]C-NMR of 2:

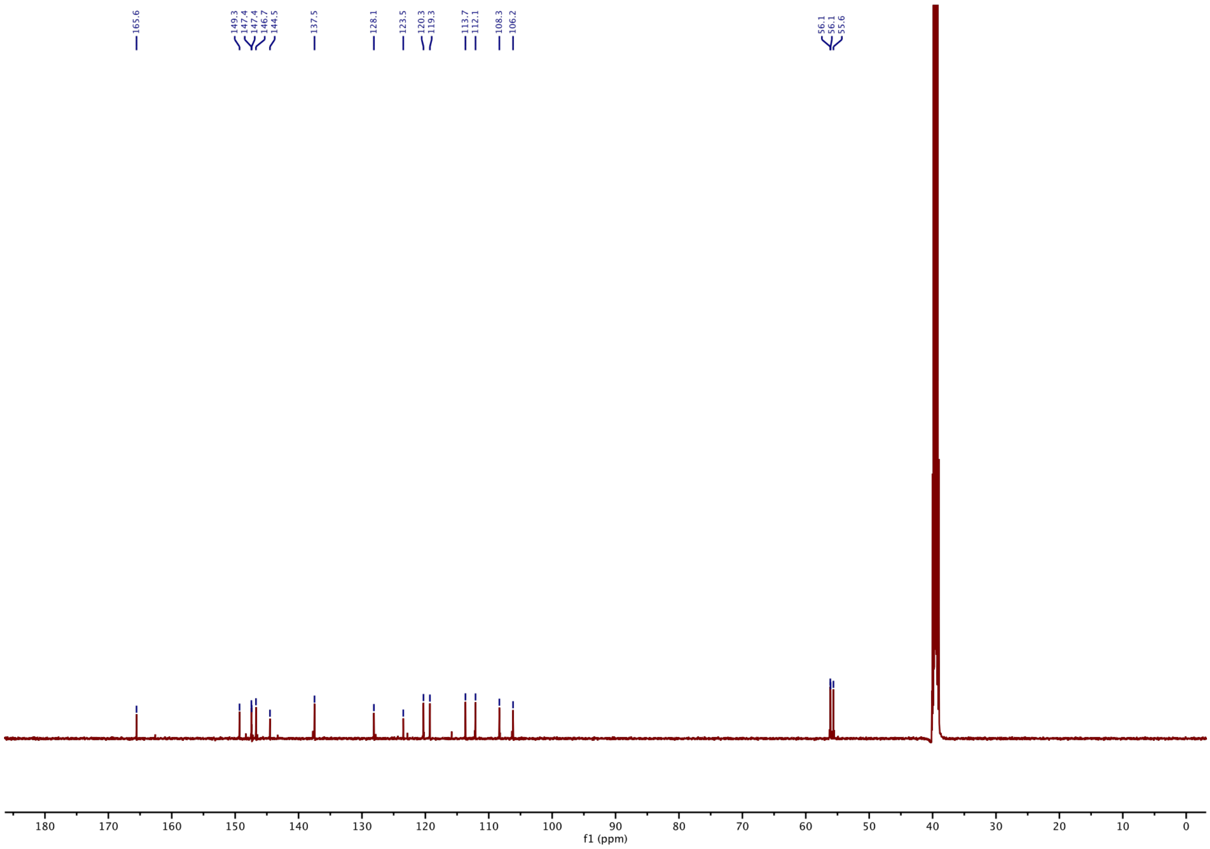

#### [1]H-NMR of 3:

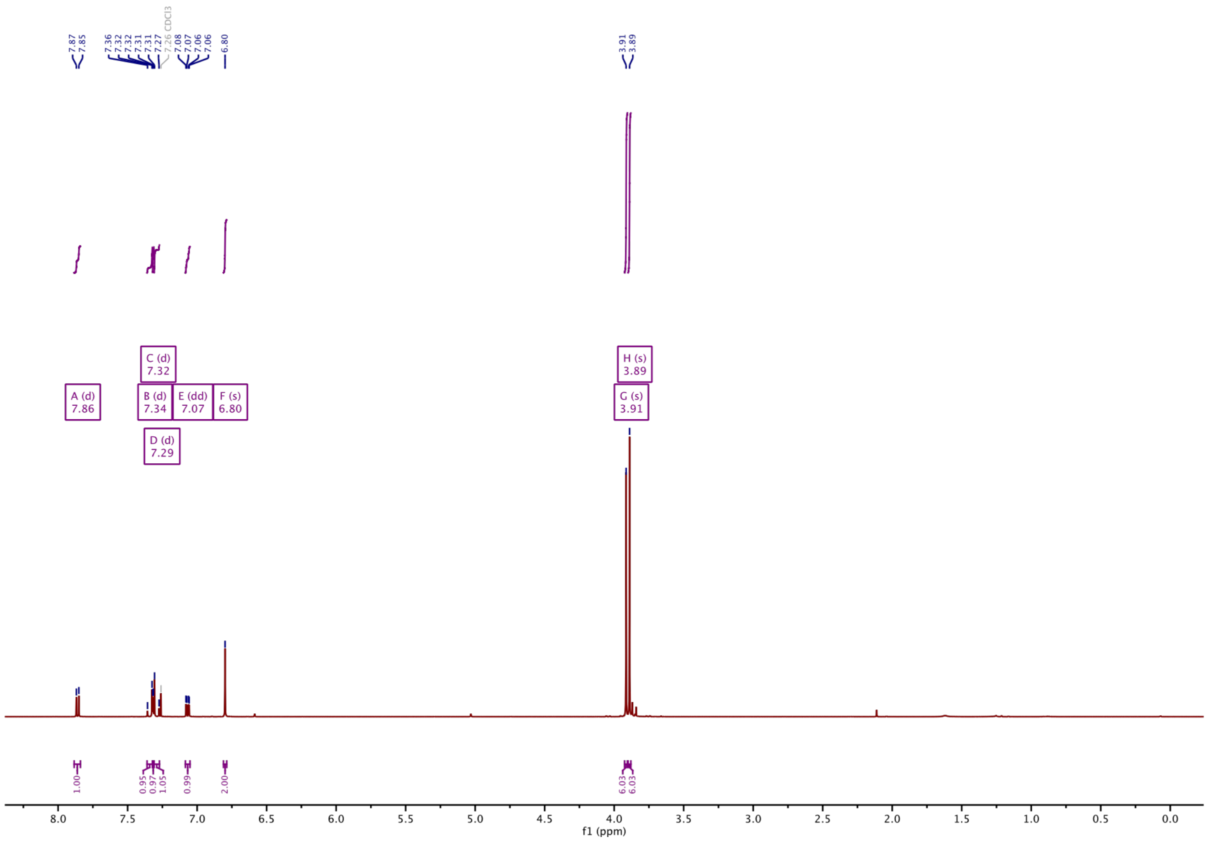

#### [13]C-NMR of 3:

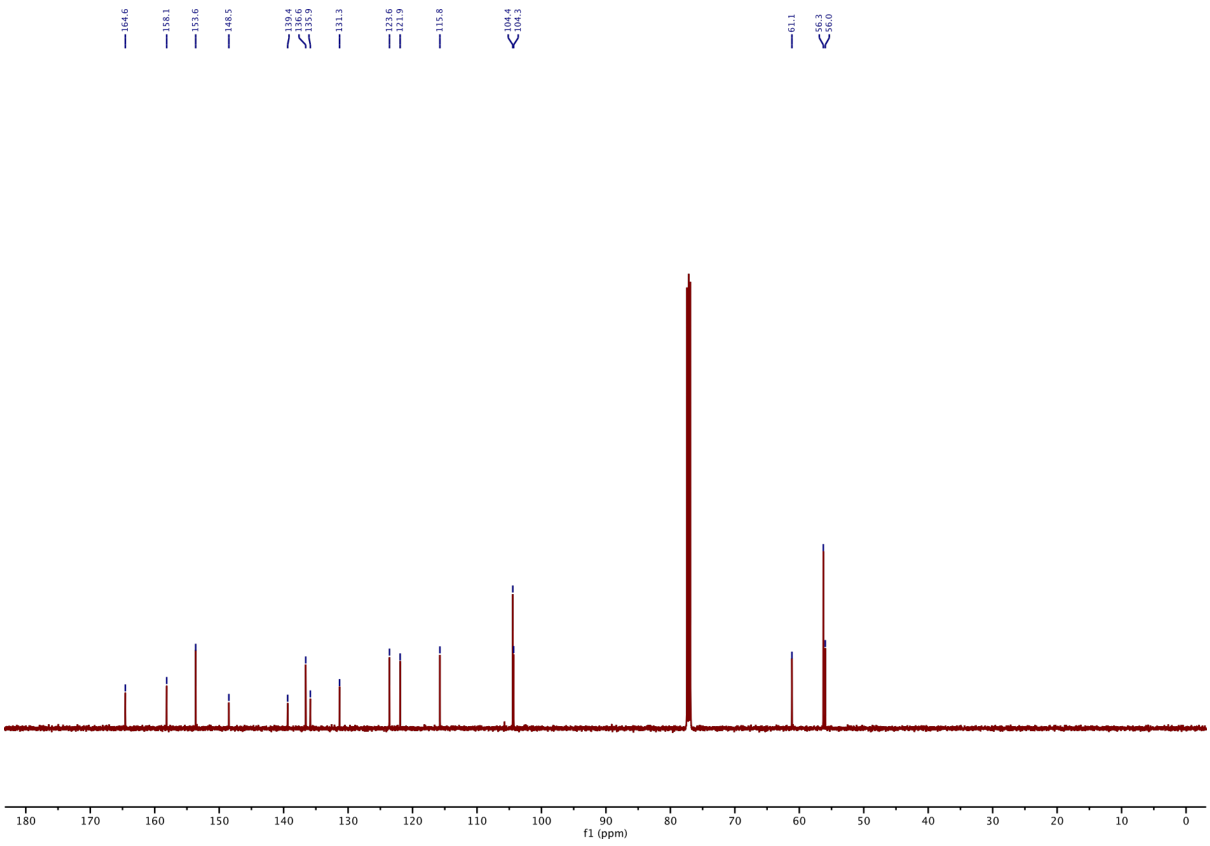

#### $^1$H-NMR of 4:

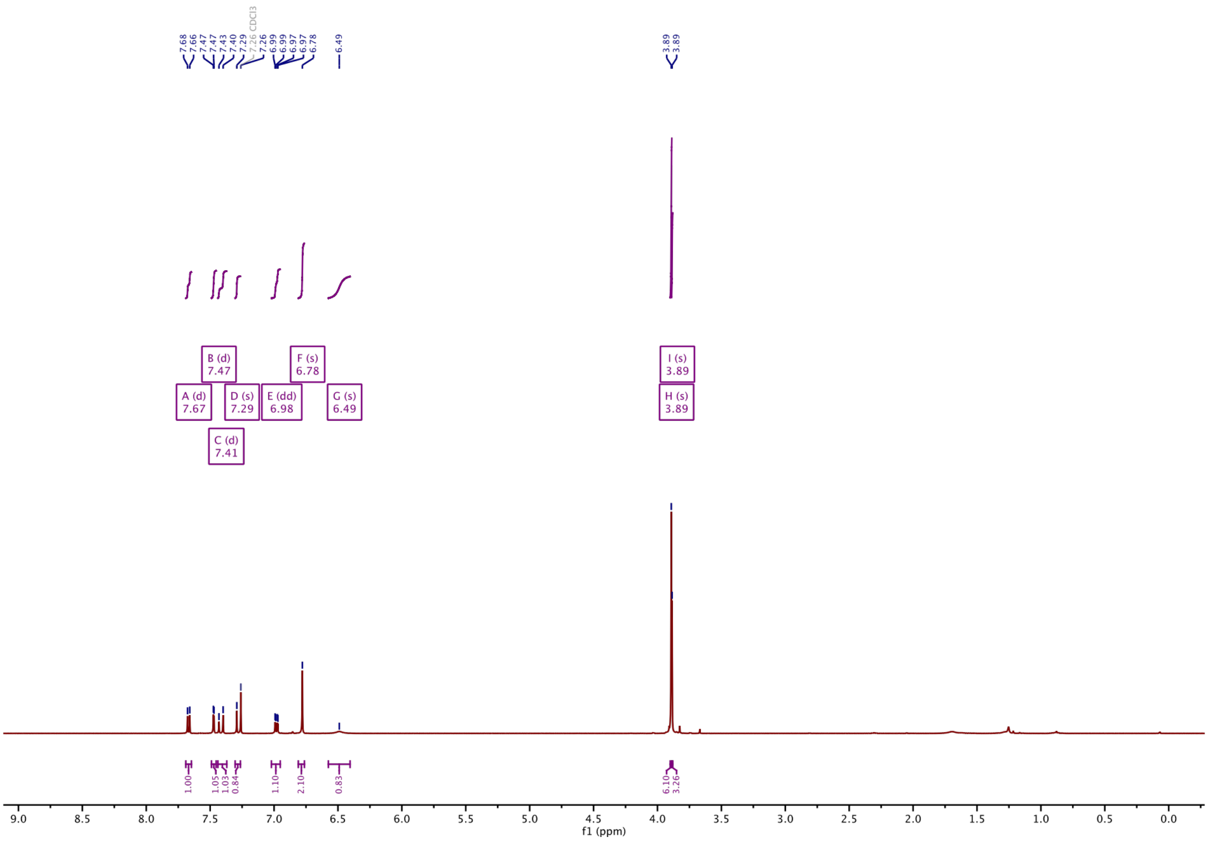

#### $^{13}$C-NMR of 4:

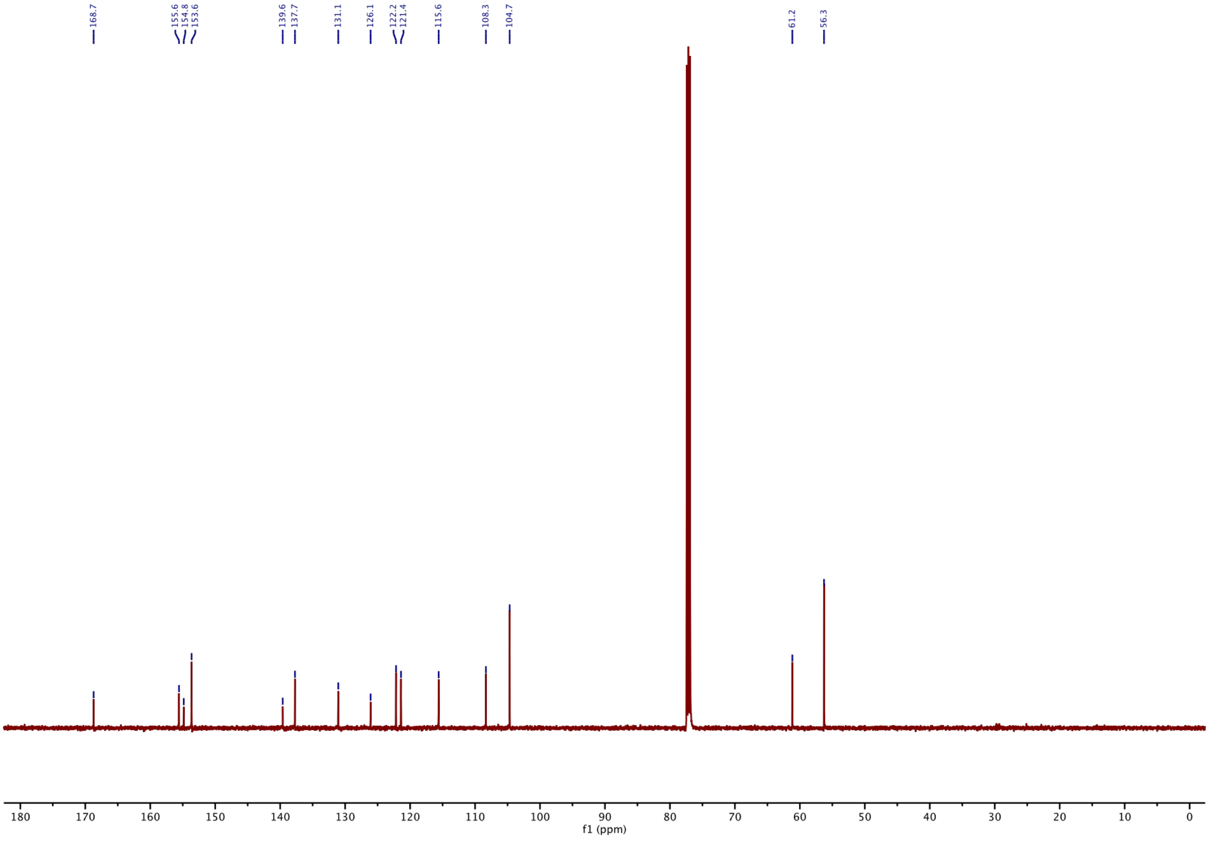

**[1]H-NMR of 5:**

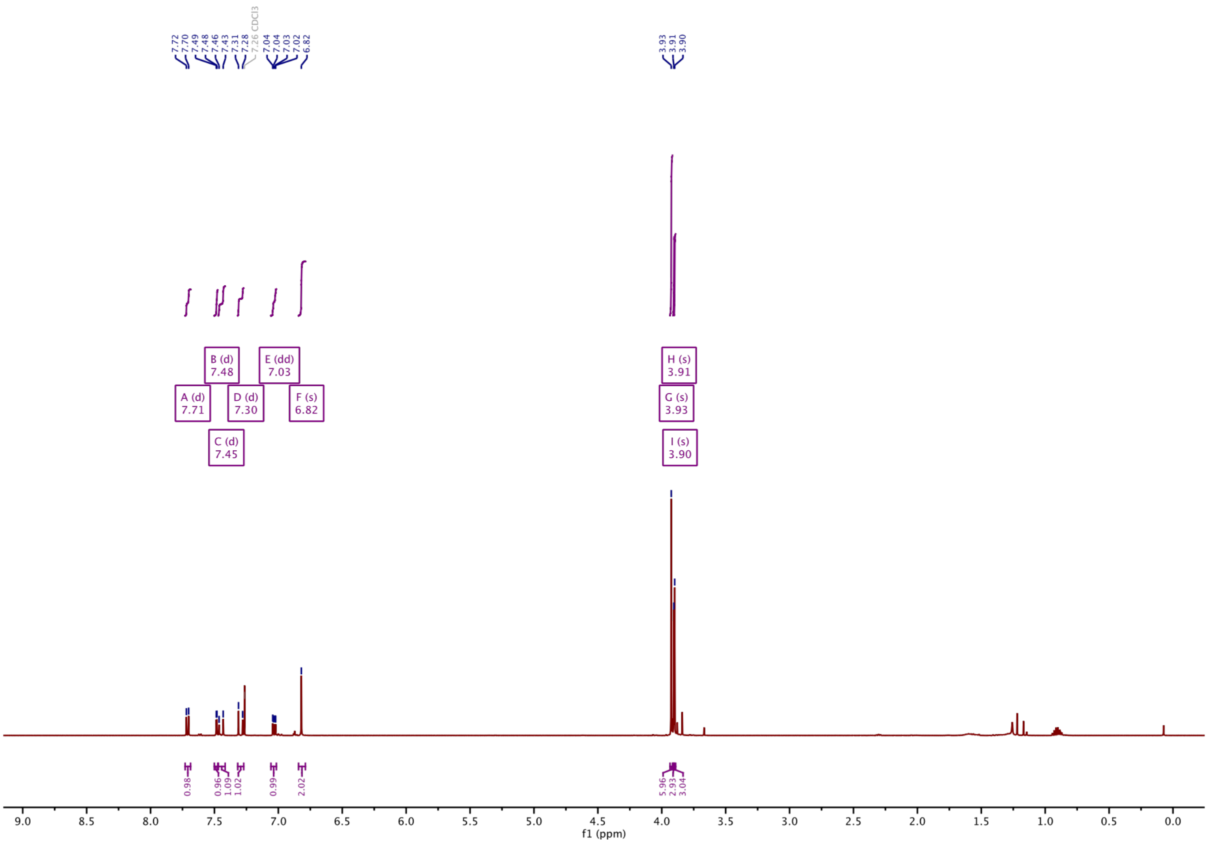

**[13]C-NMR of 5:**

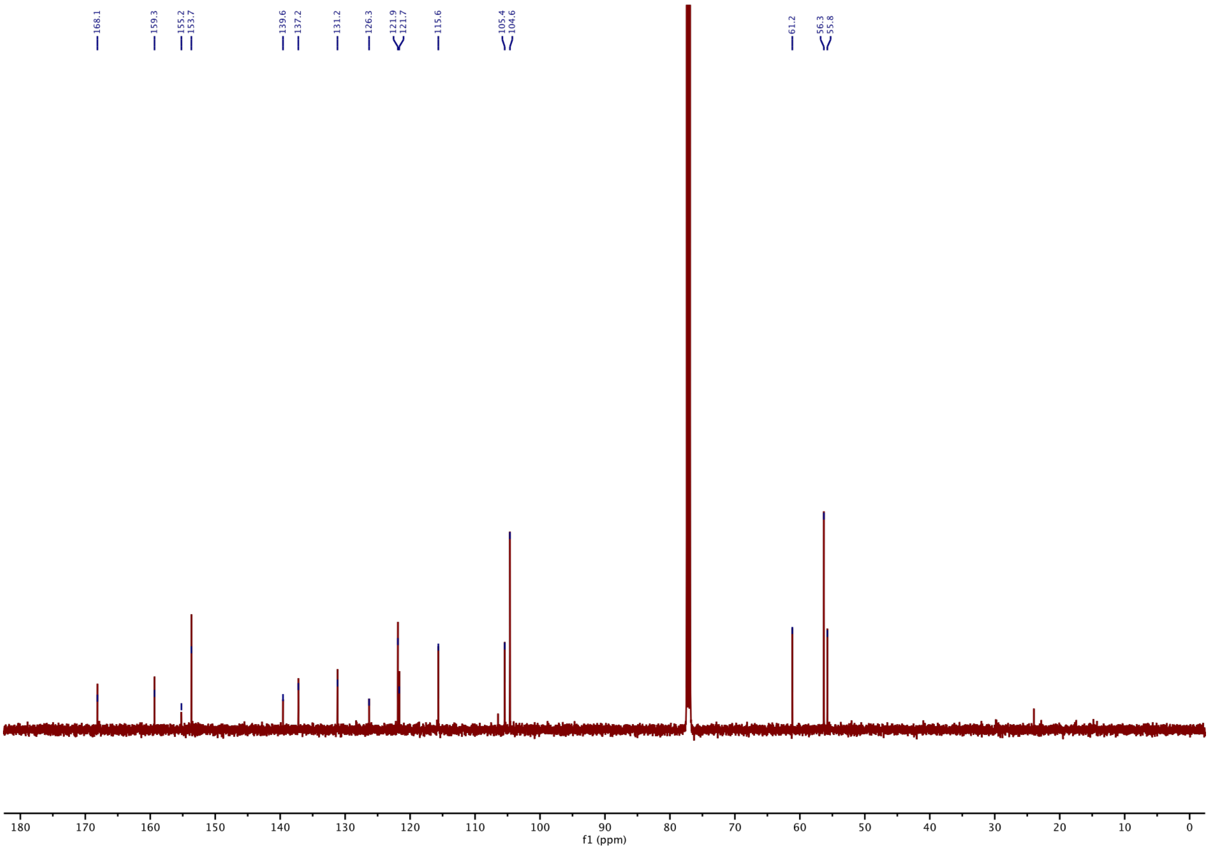

**[1]H-NMR of 6:**

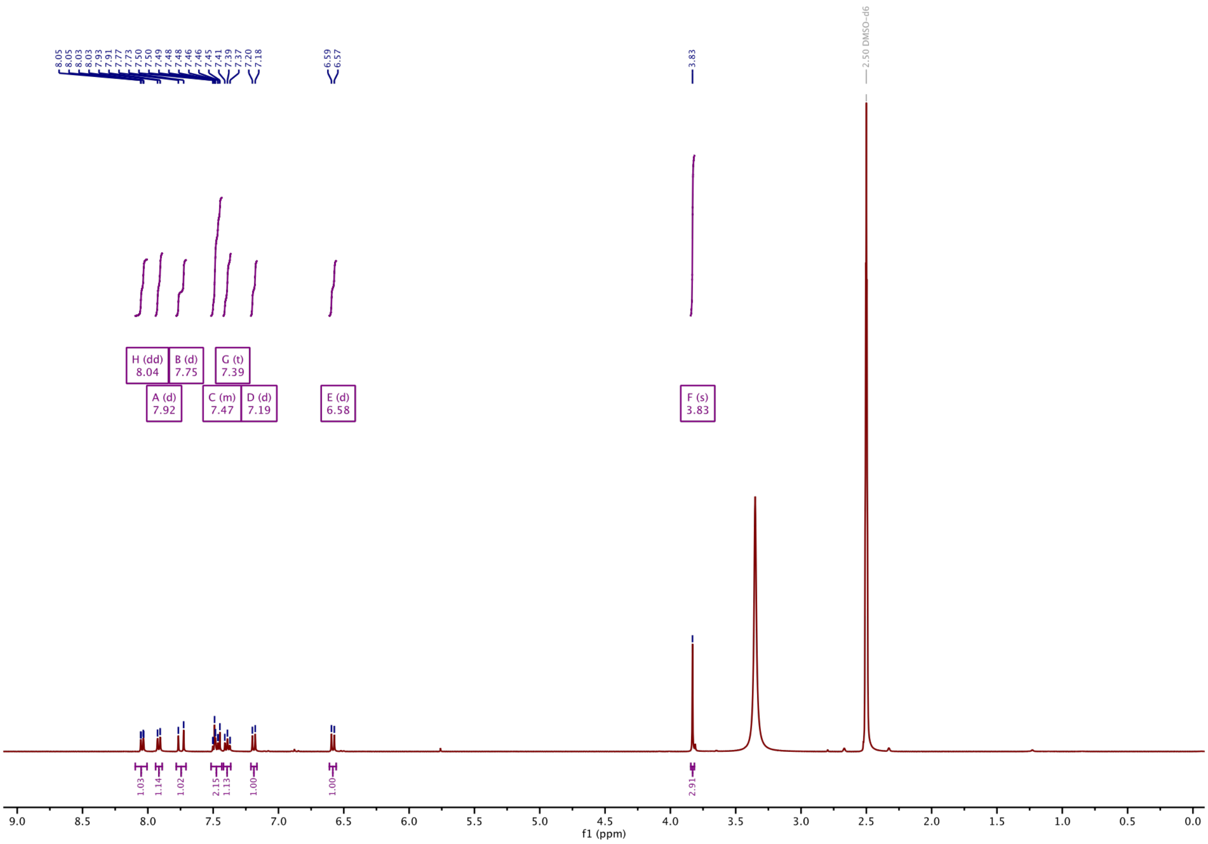

**[13]C-NMR of 6:**

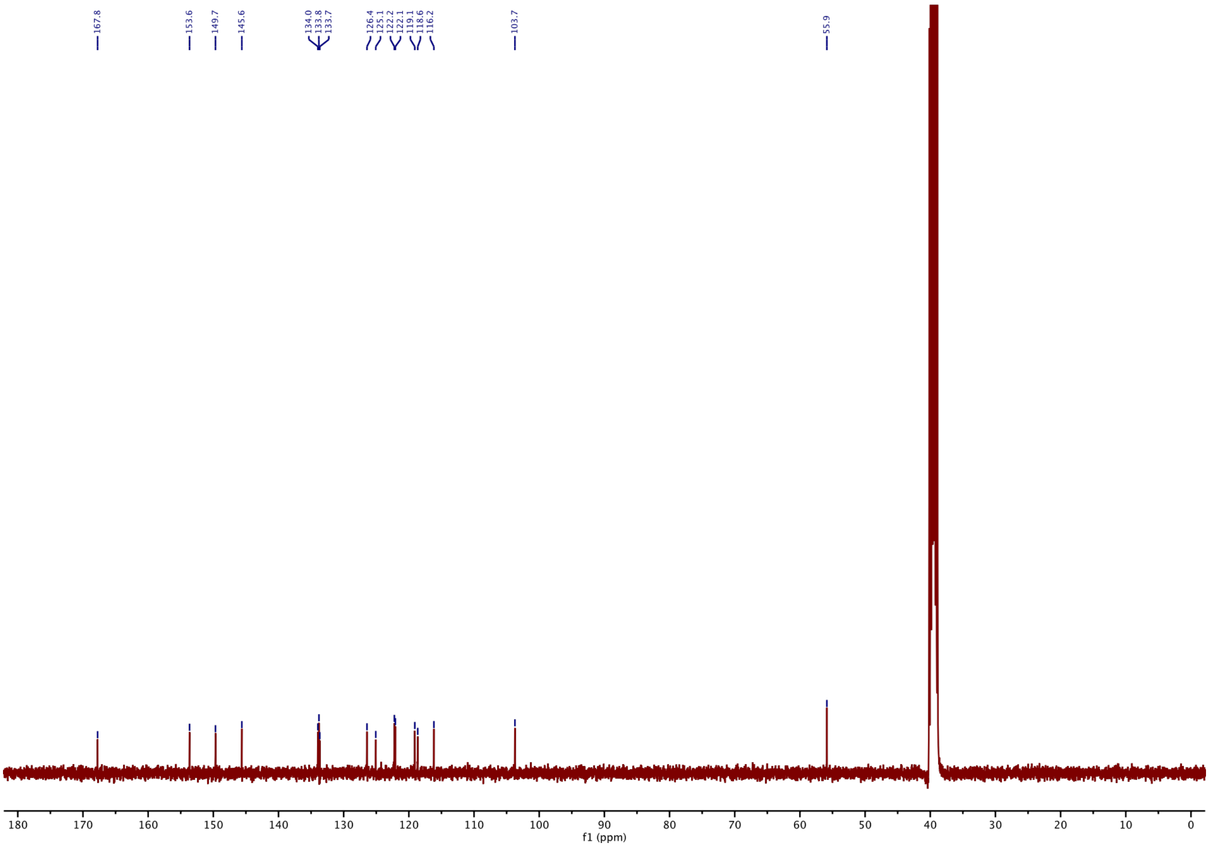

#### $^{1}$H-NMR of 7:

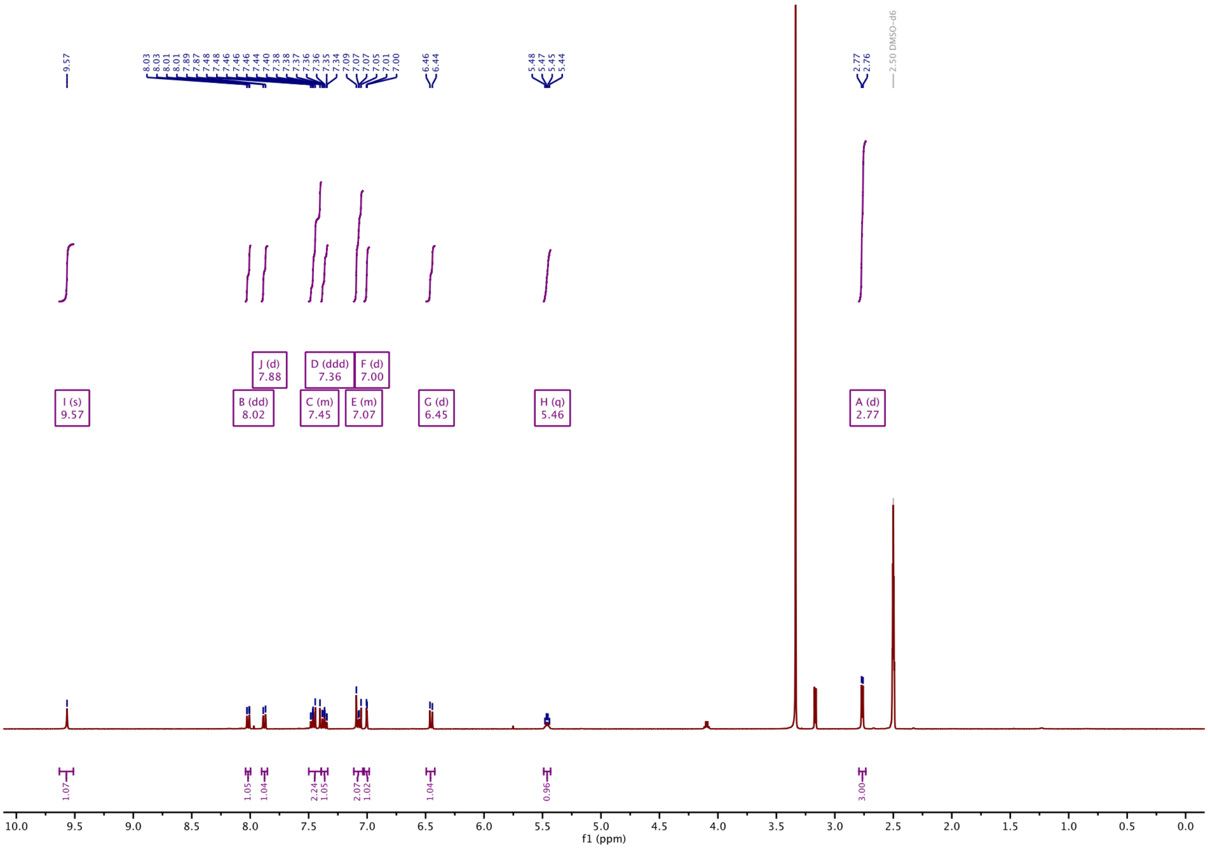

#### $^{13}$C-NMR of 7:

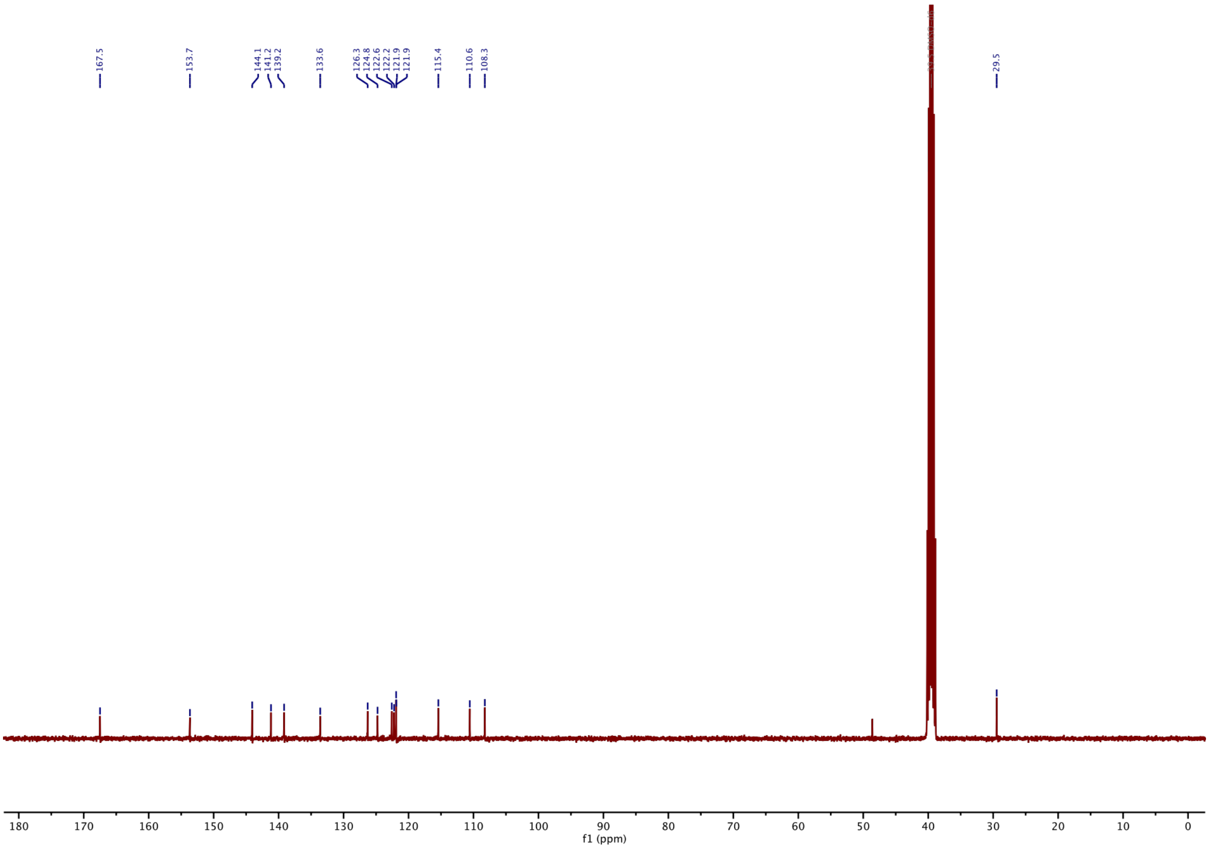

**$^1$H-NMR of 8:**

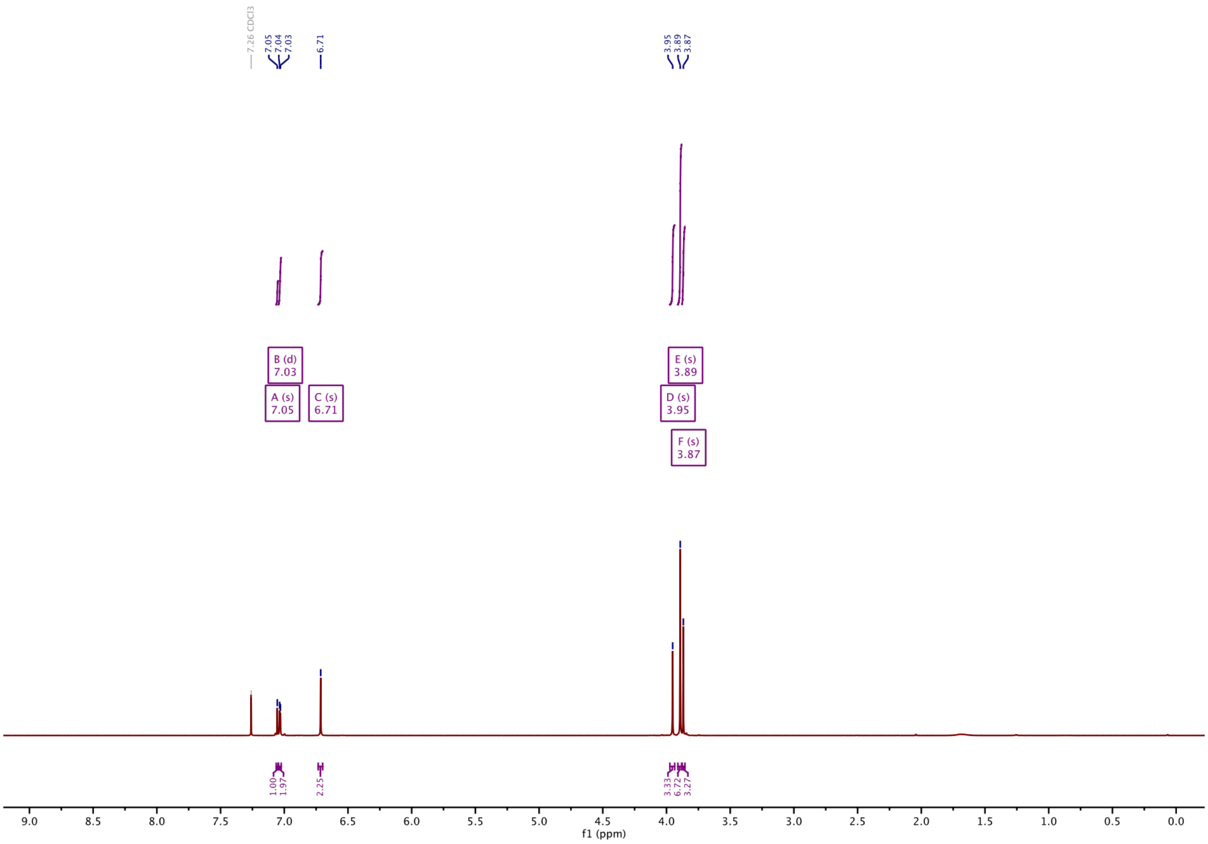

**$^{13}$C-NMR of 8:**

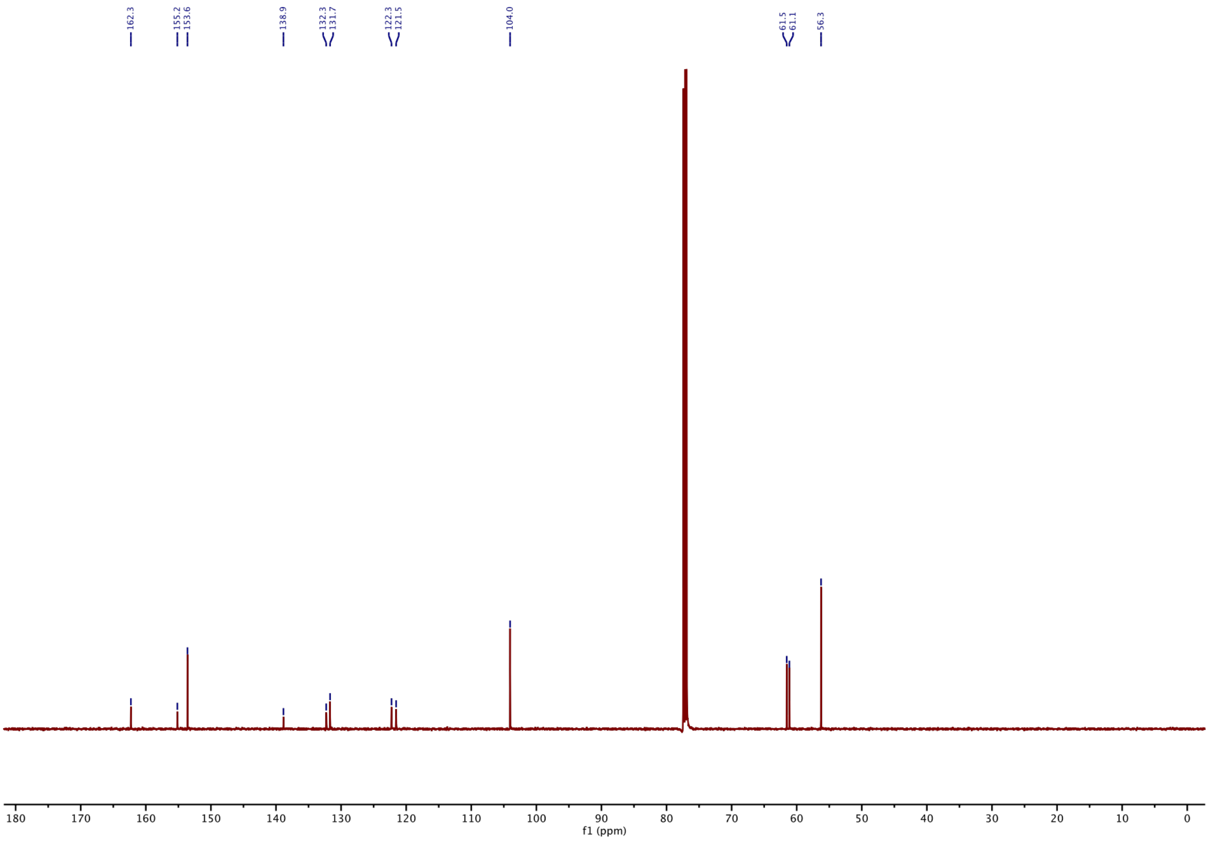

#### $^1$H-NMR of 9:

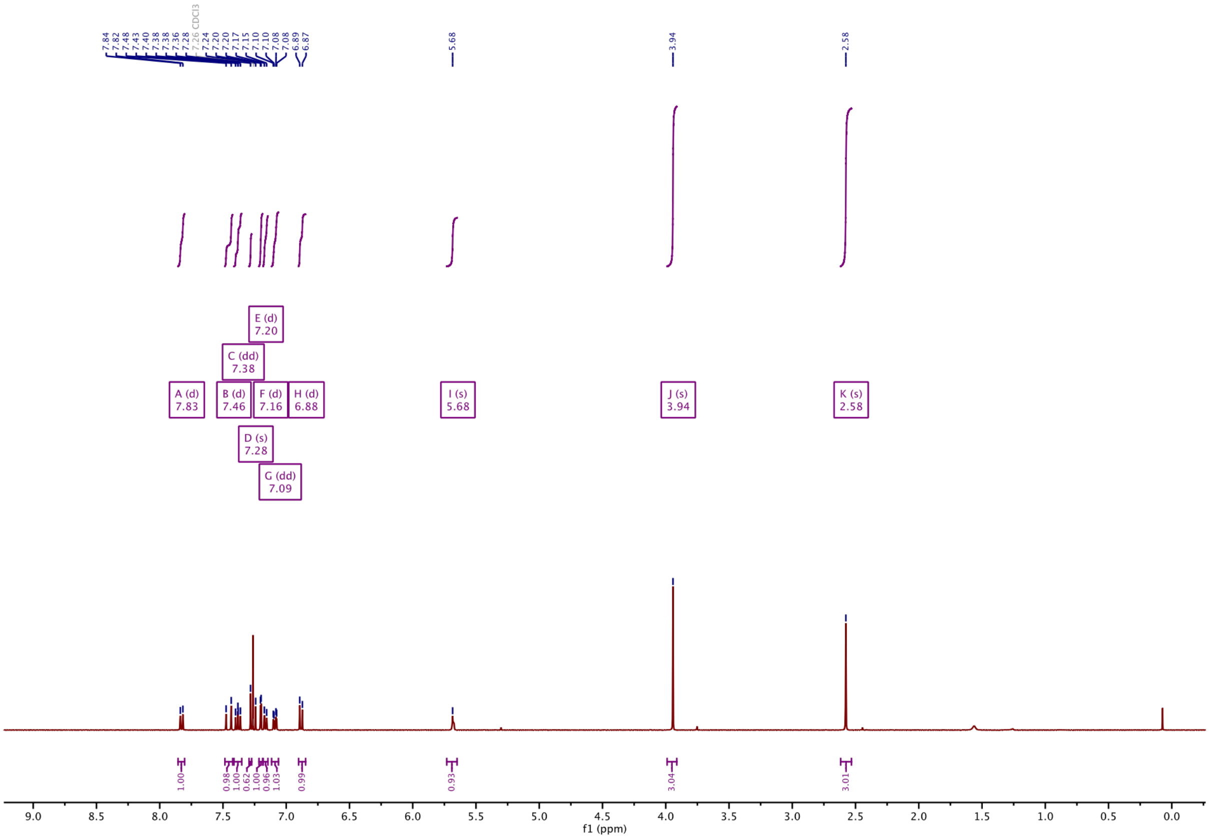

#### $^{13}$C-NMR of 9:

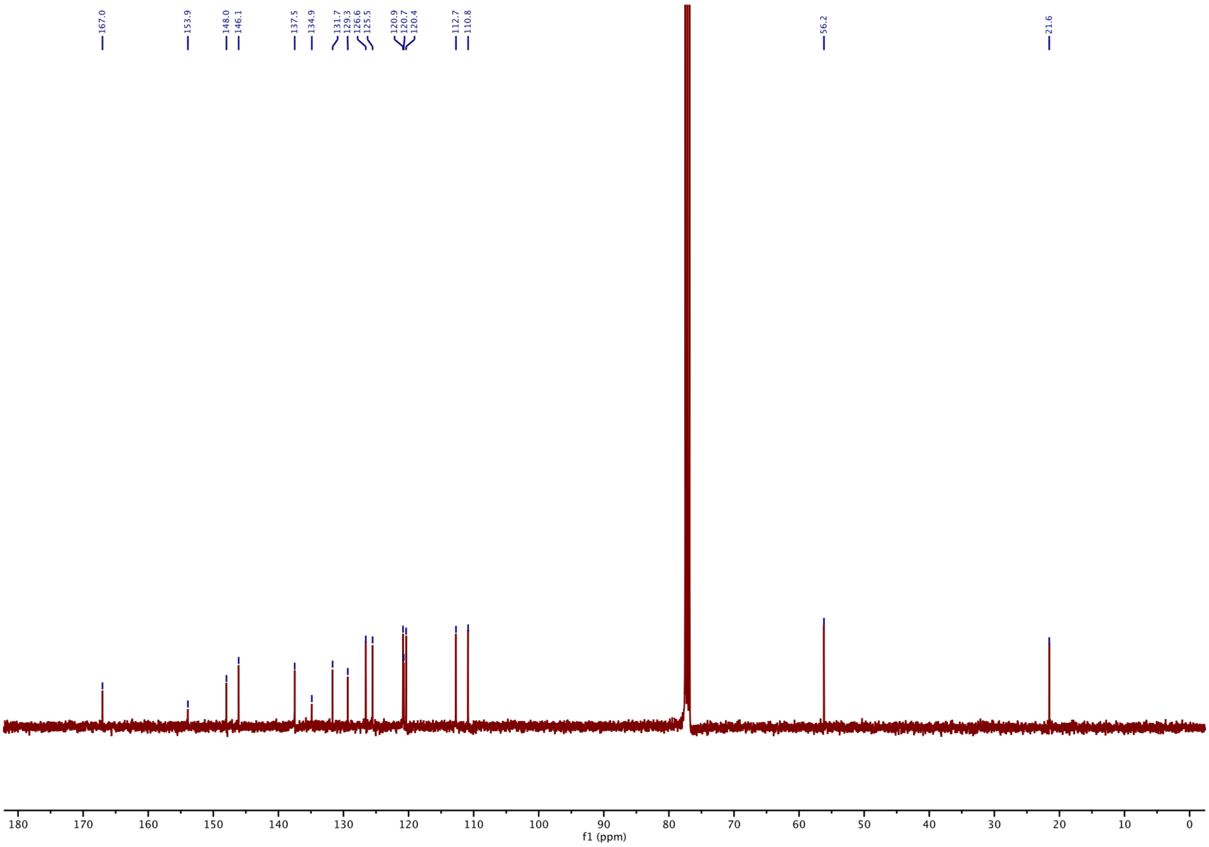

#### [1]H-NMR of 10:

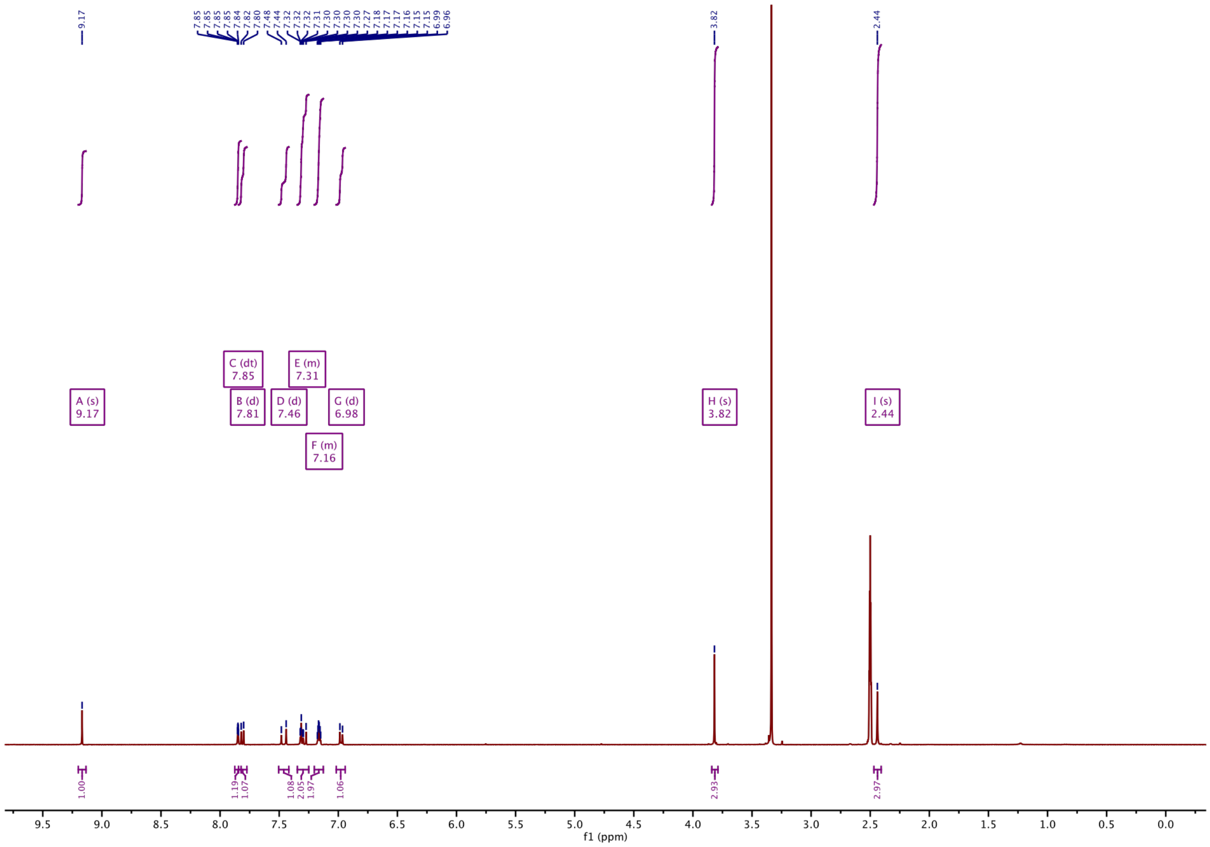

#### [13]C-NMR of 10:

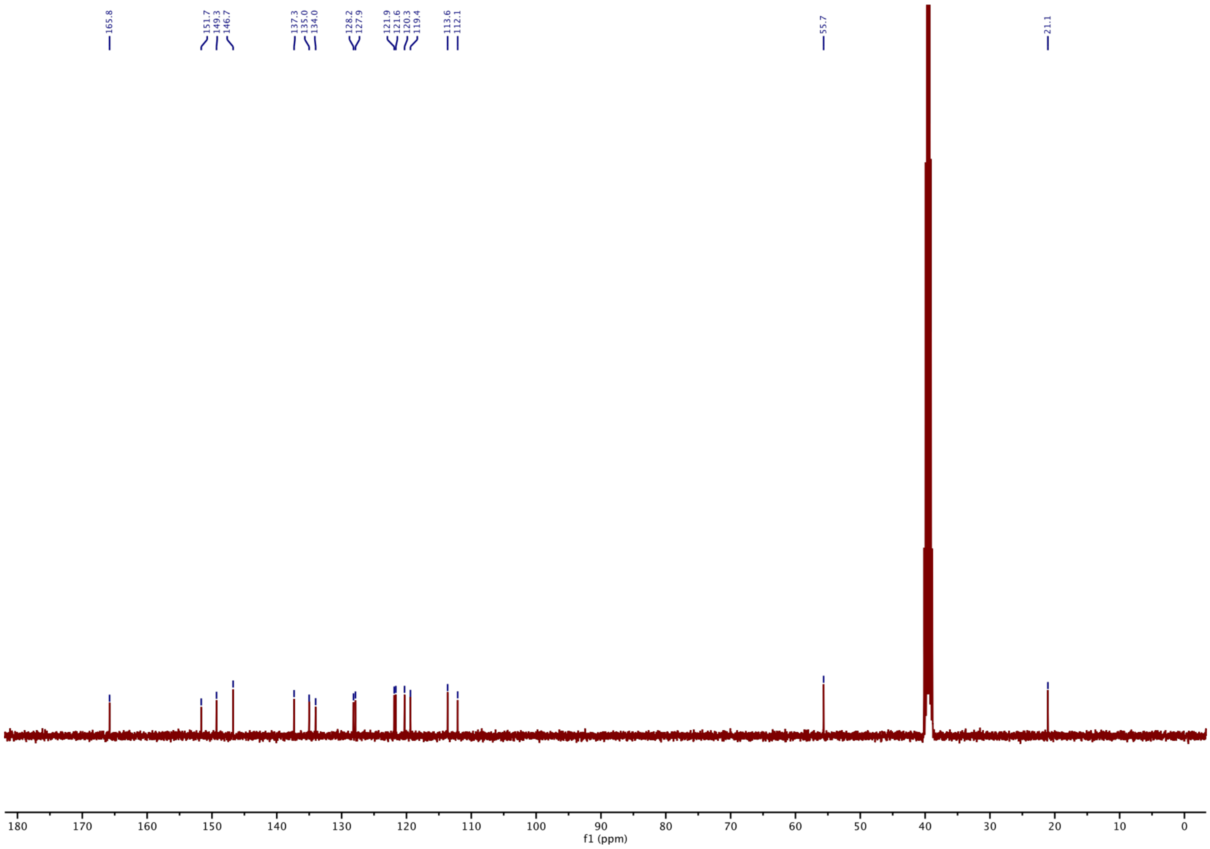

#### $^1$H-NMR of 11:

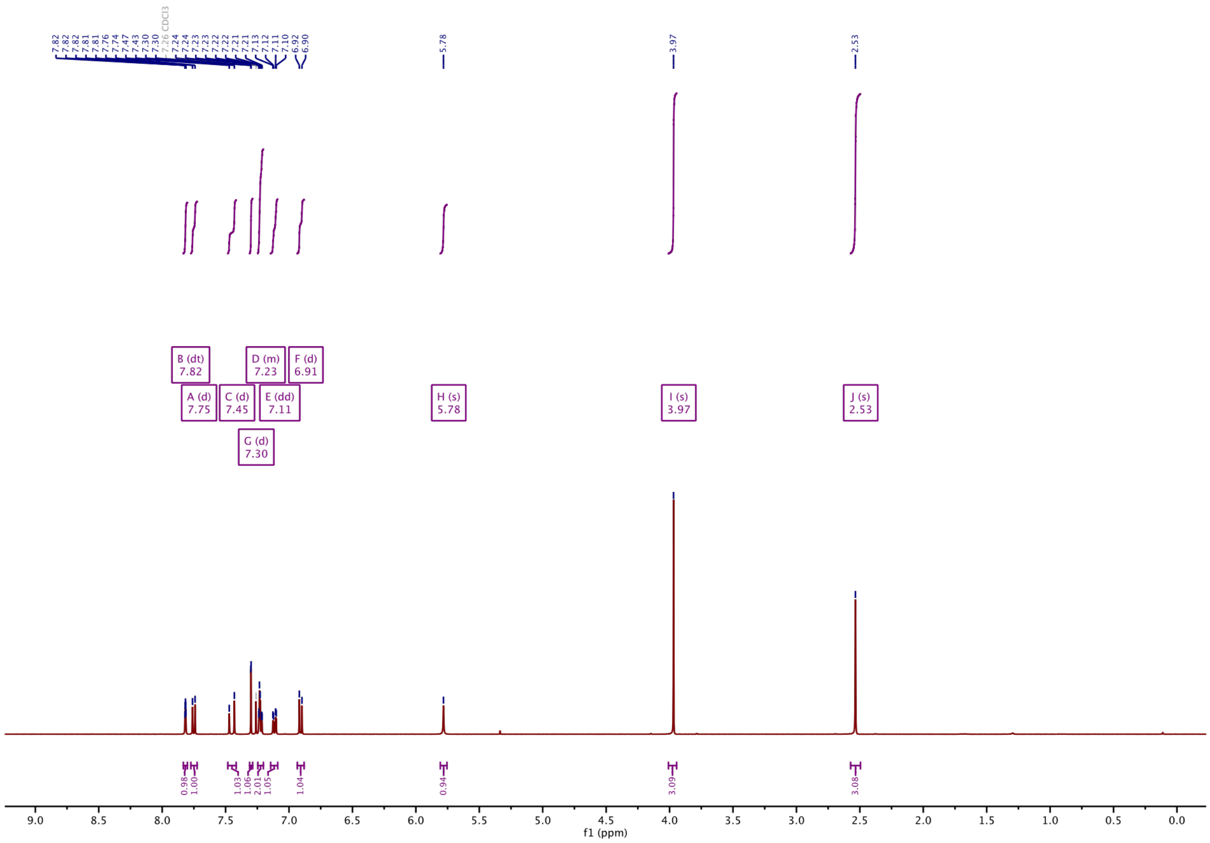

#### $^{13}$C-NMR of 11:

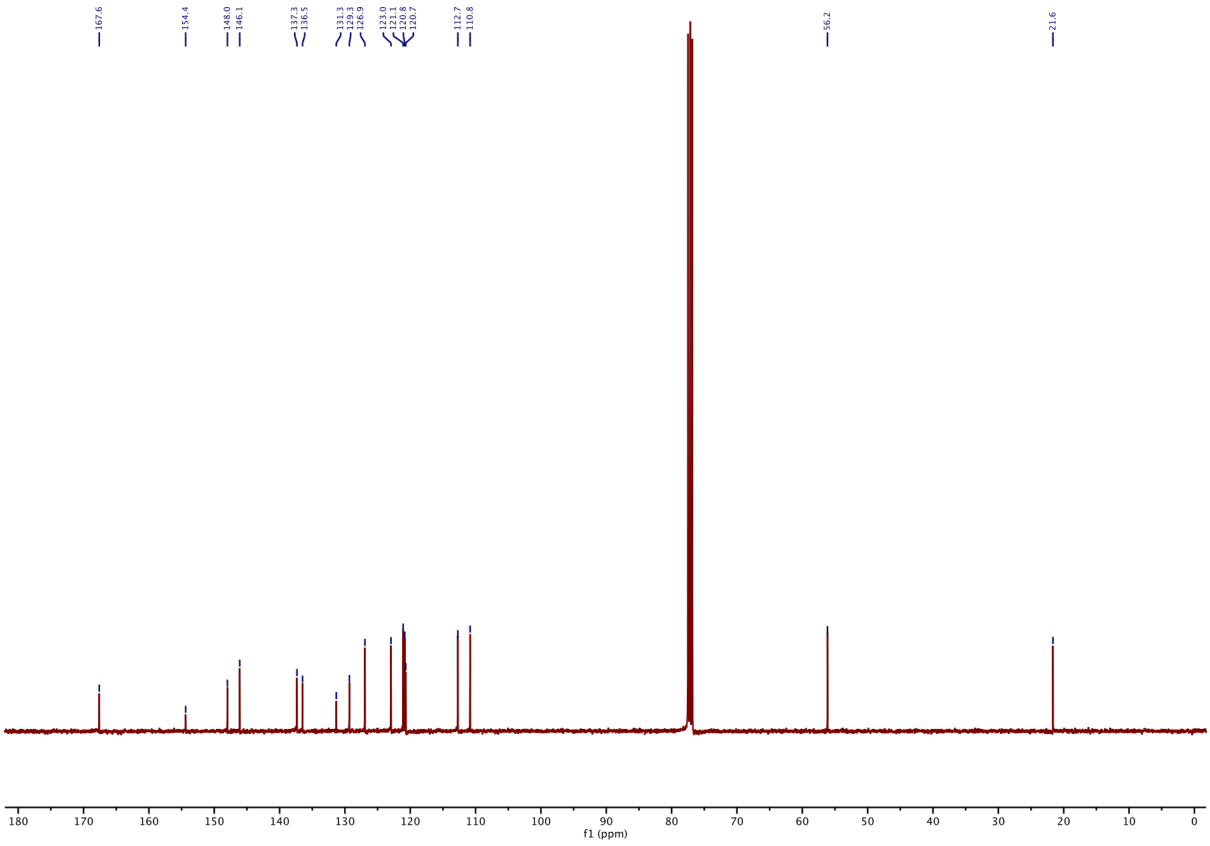

**[1]H-NMR of 12:**

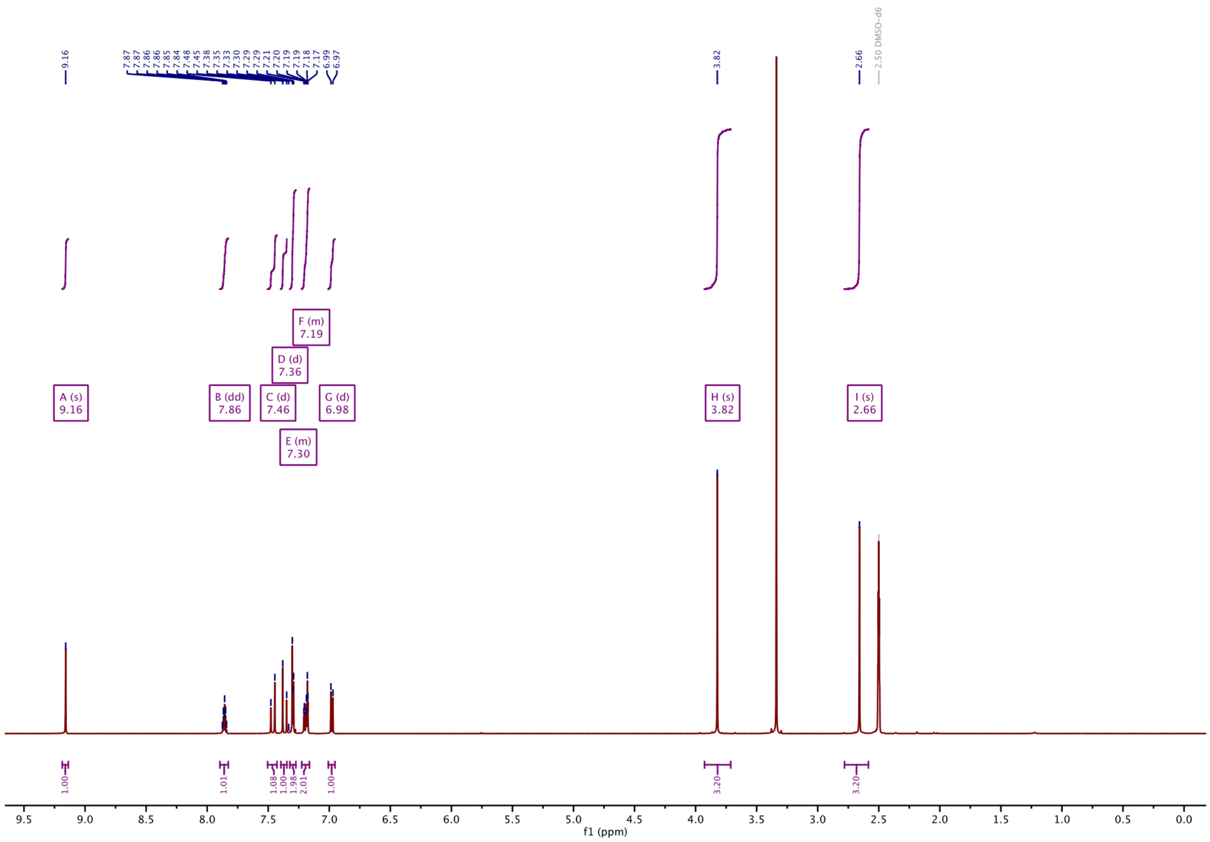

**[13]C-NMR of 12:**

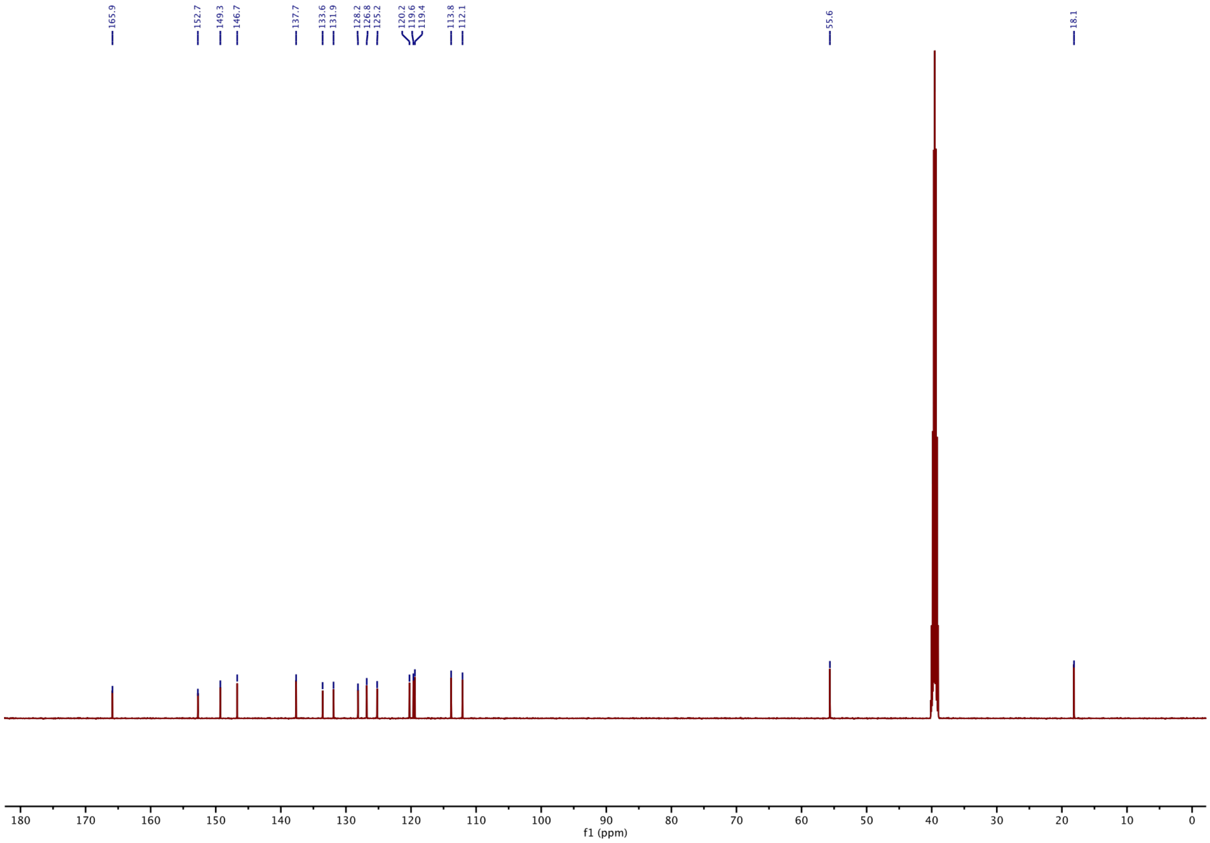

#### [1]H-NMR of 13:

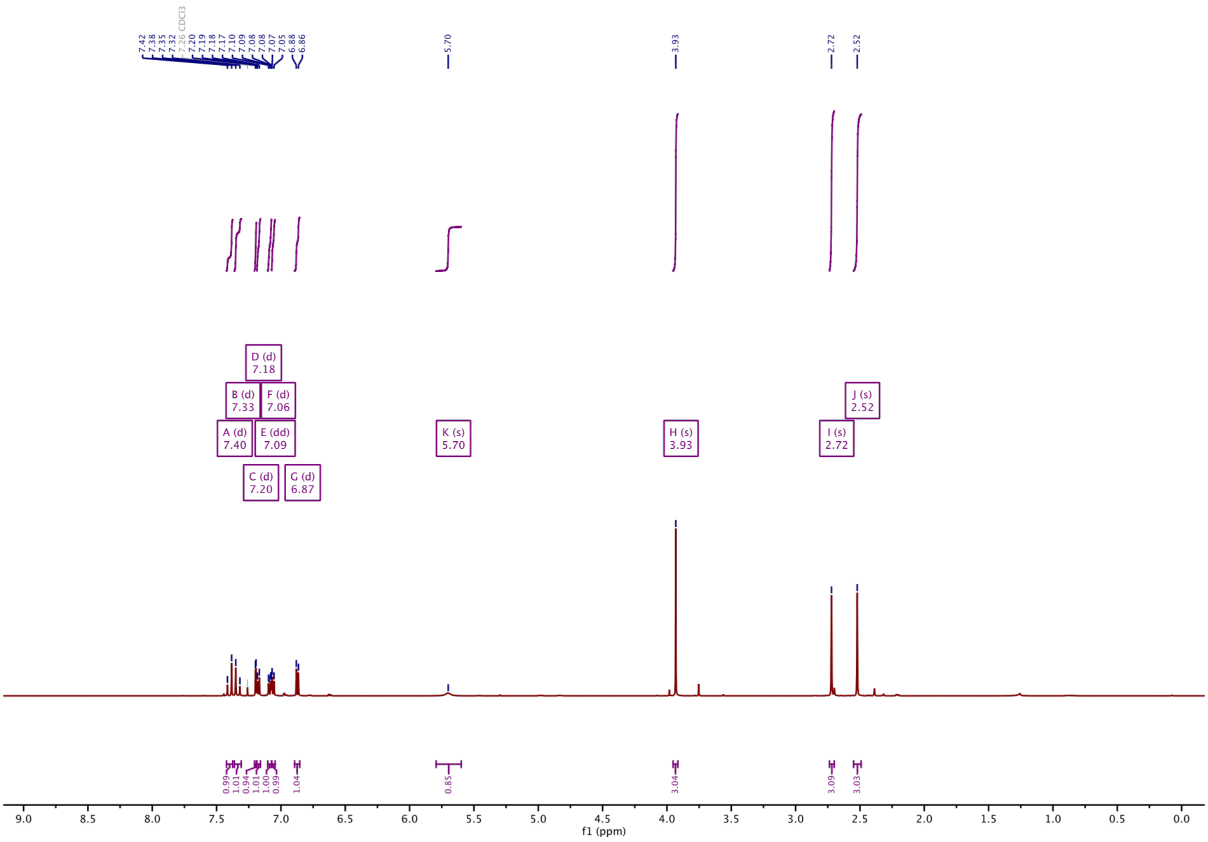

#### [13]C-NMR of 13:

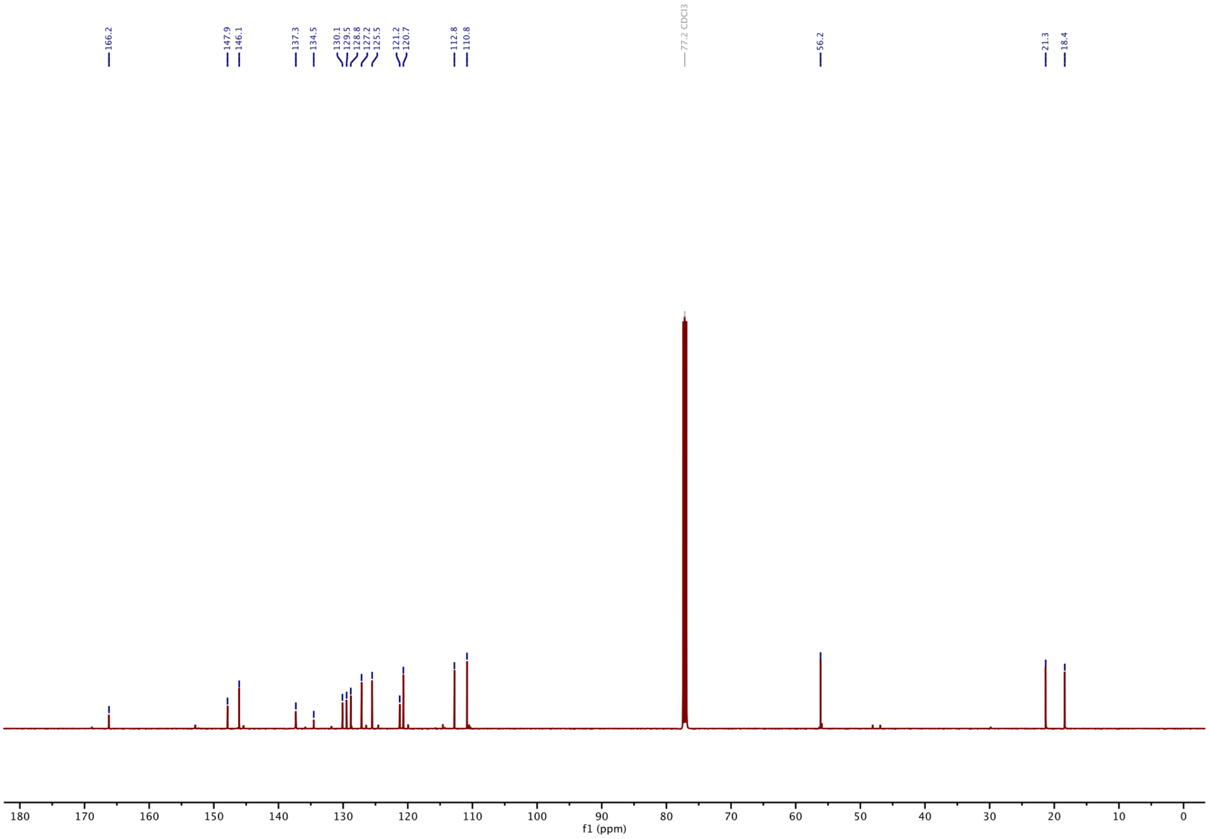

#### $^1$H-NMR of 14:

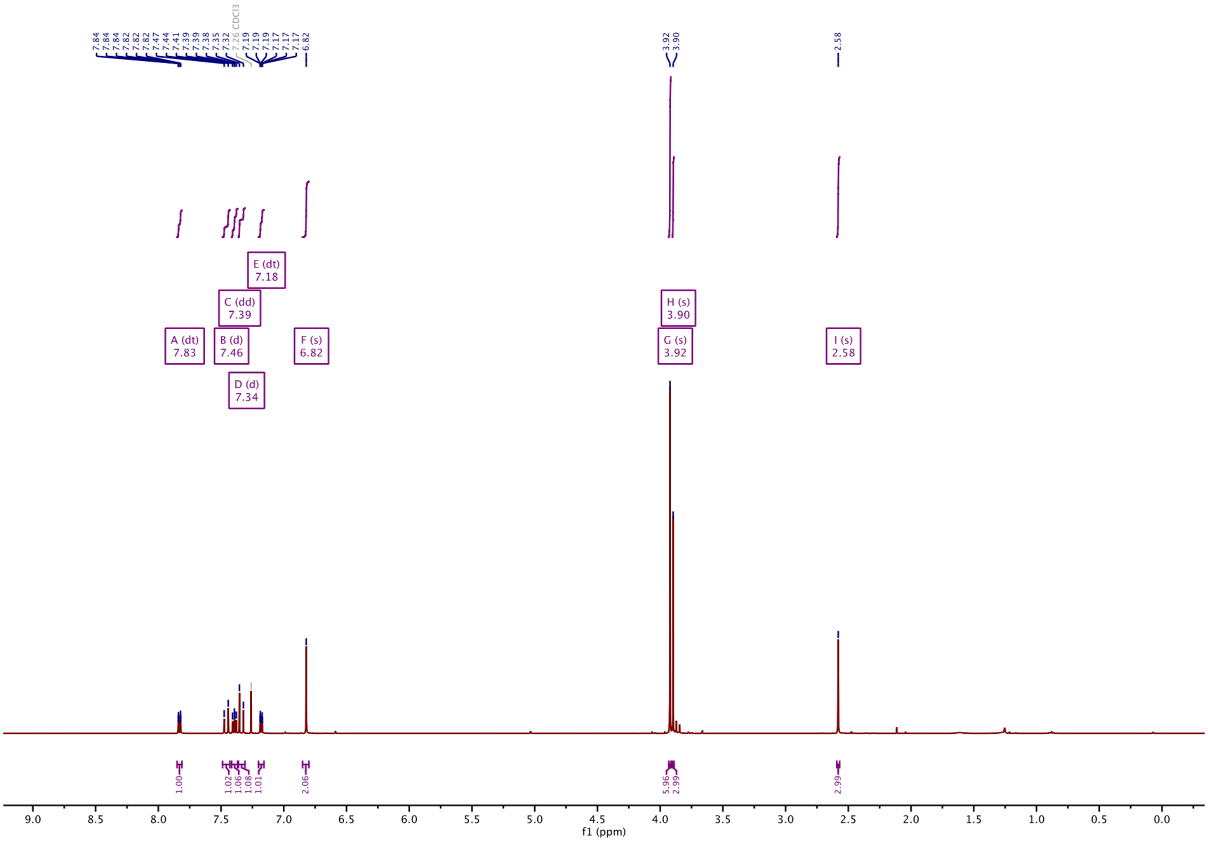

#### $^{13}$C-NMR of 14:

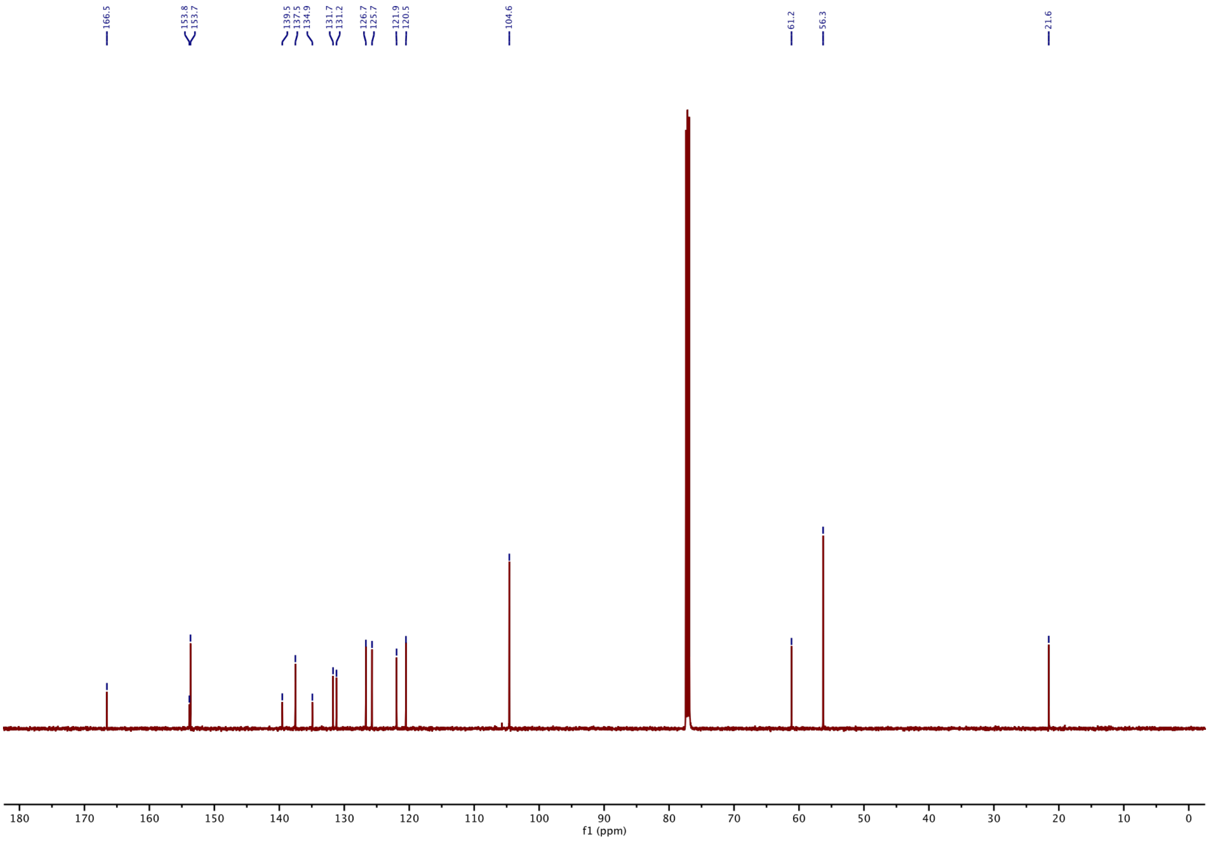

#### 1H-NMR of SBTub2M:

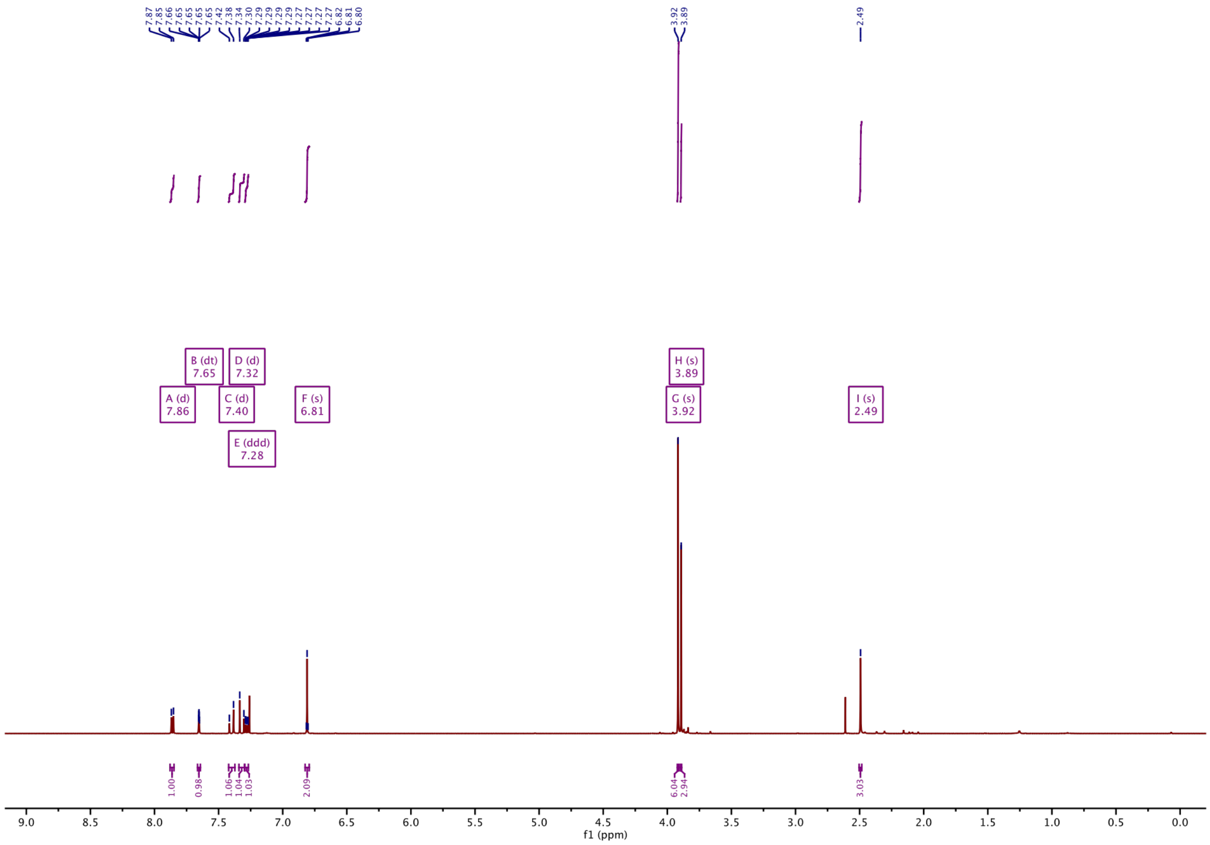

#### 13C-NMR of SBTub2M:

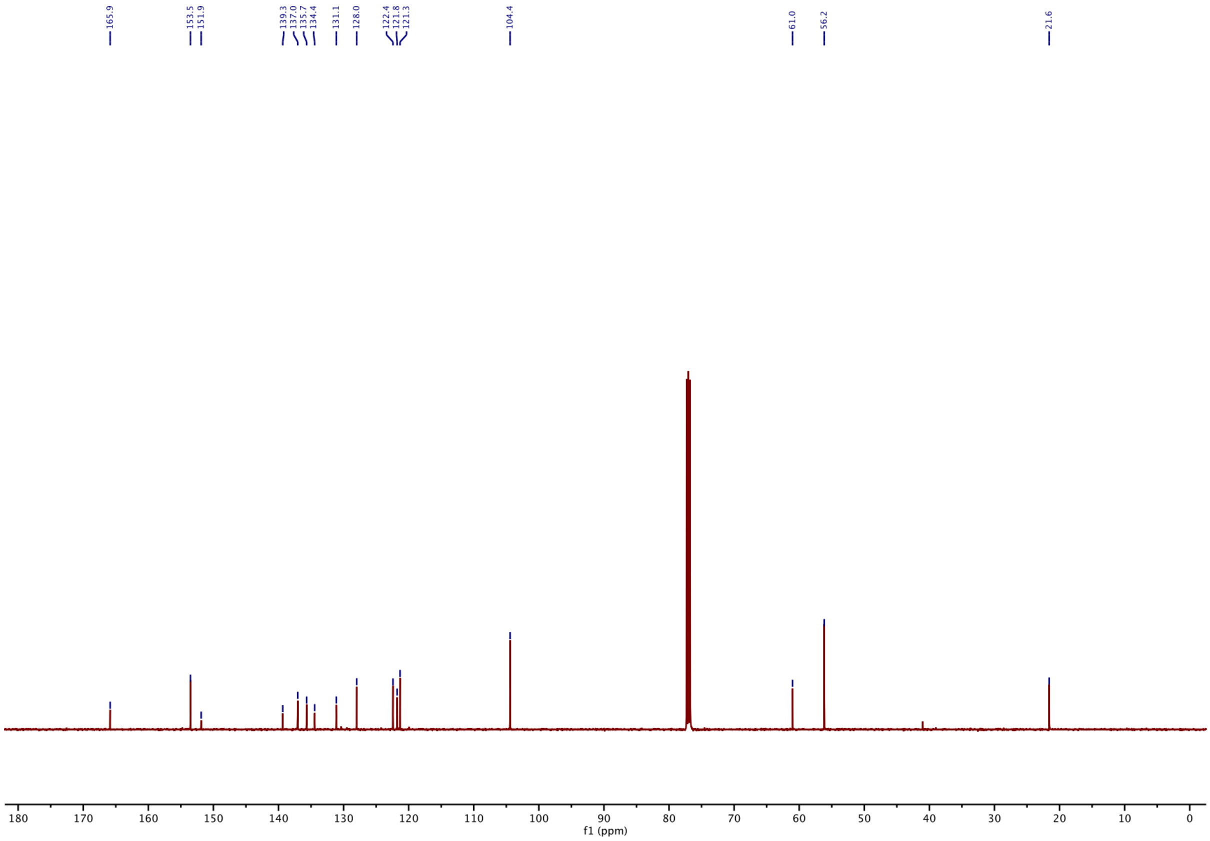

#### ¹H-NMR of 16:

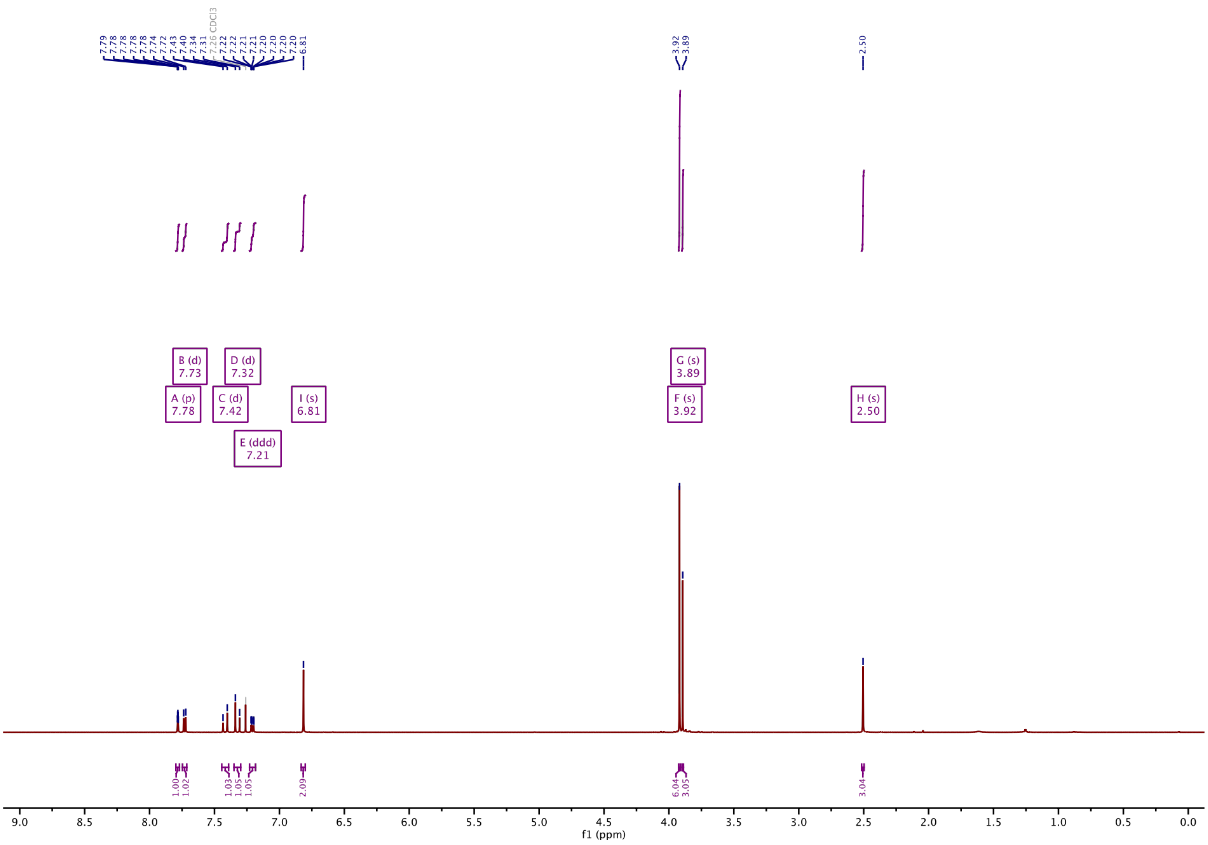

#### ¹³C-NMR of 16:

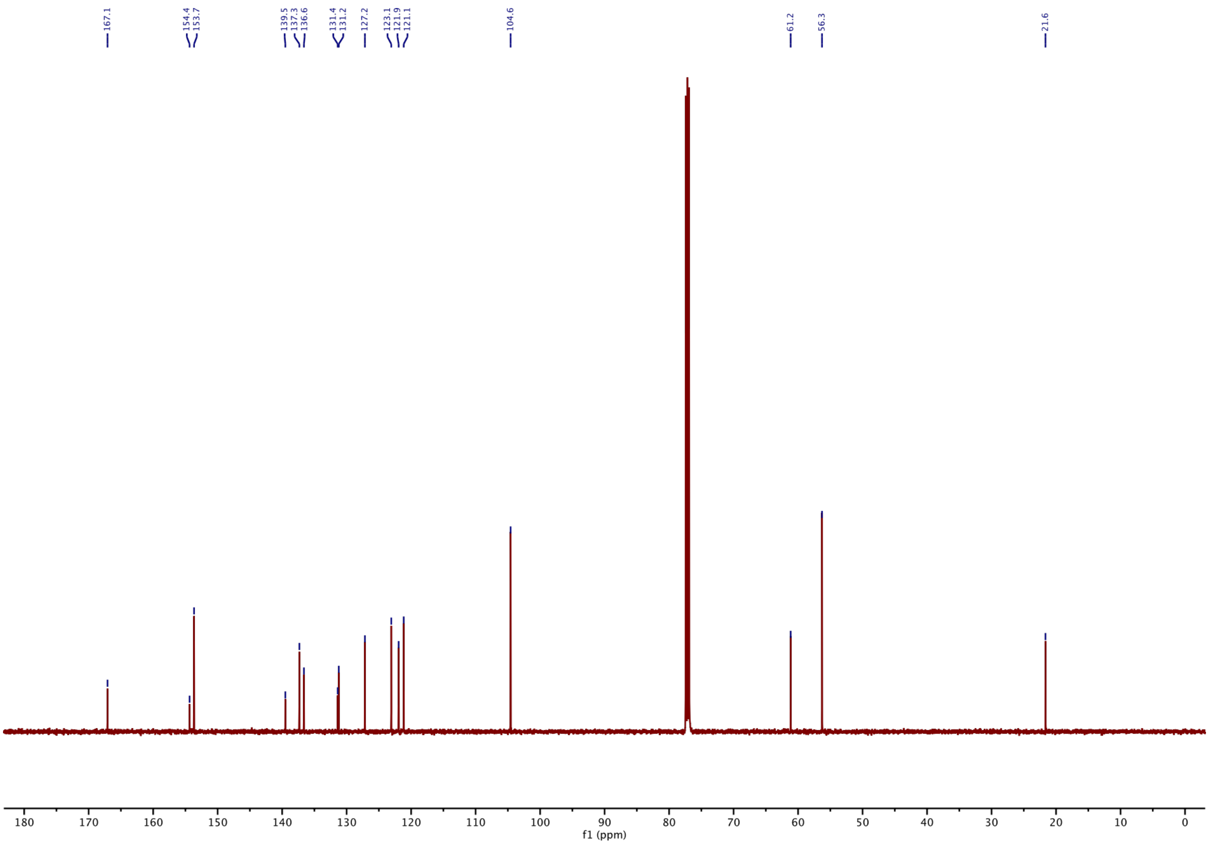

#### [1]H-NMR of 17:

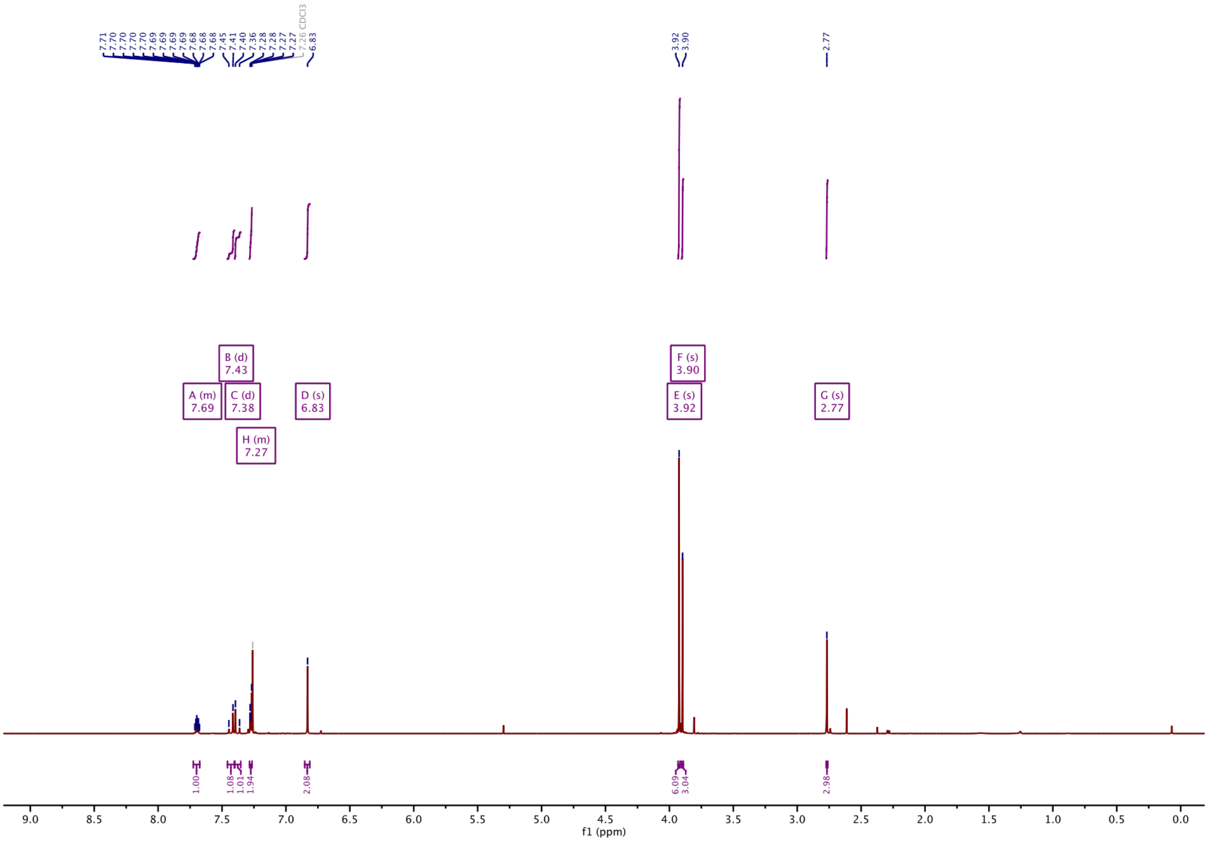

#### [13]C-NMR of 17:

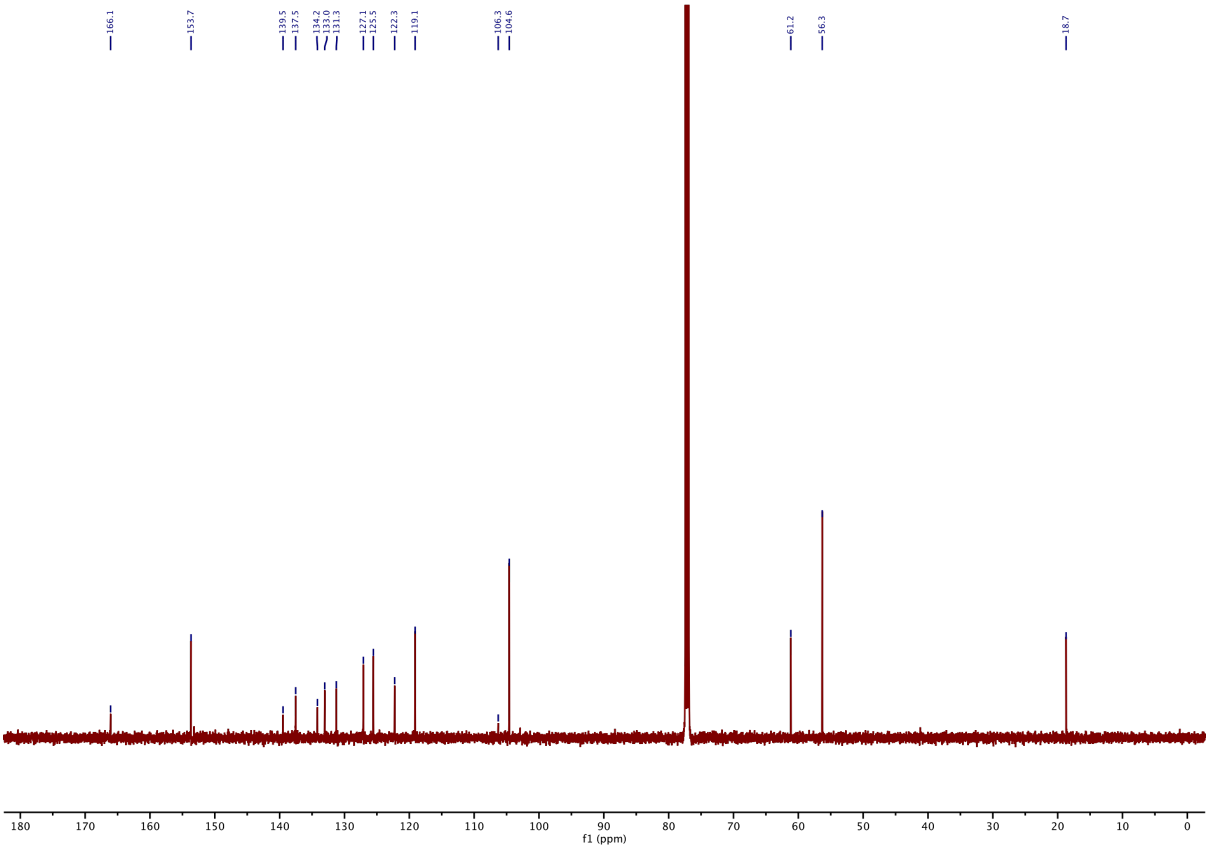

**$^{1}$H-NMR of 18:**

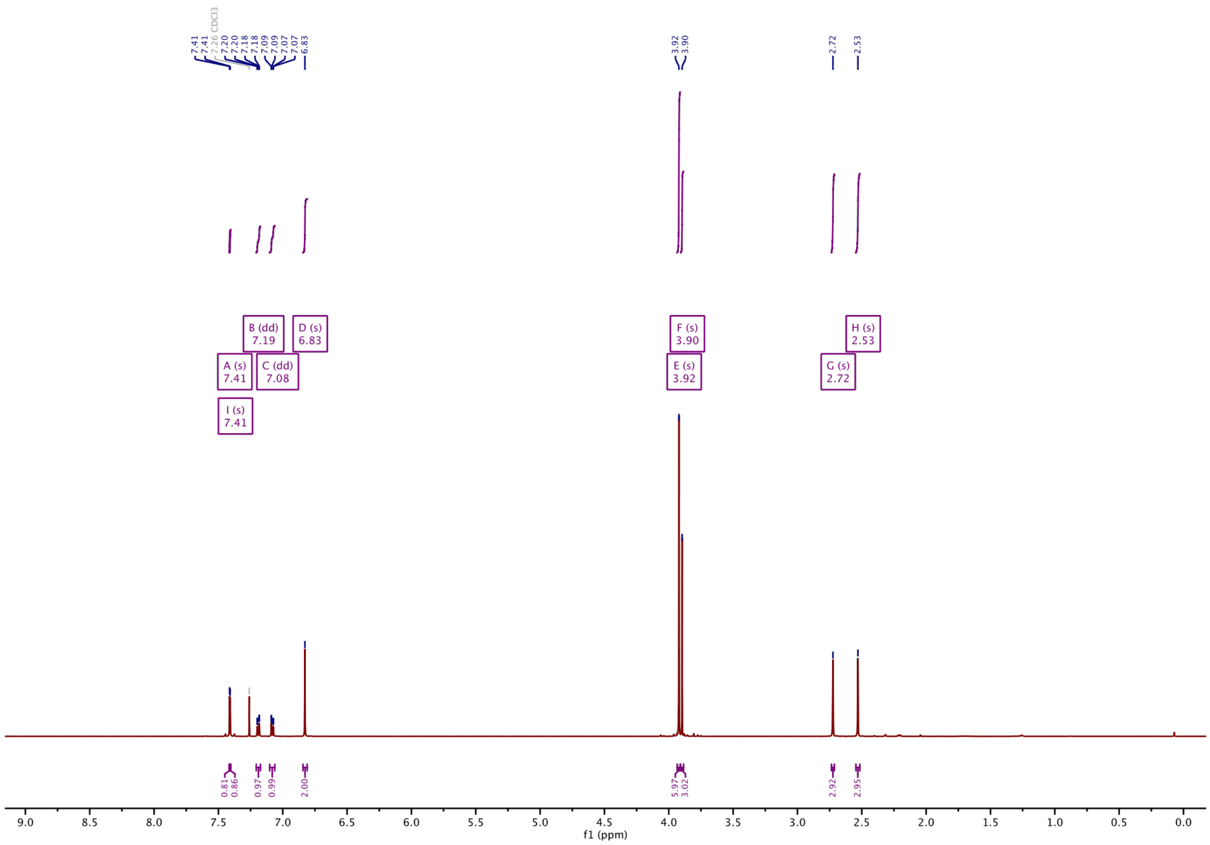

**$^{13}$C-NMR of 18:**

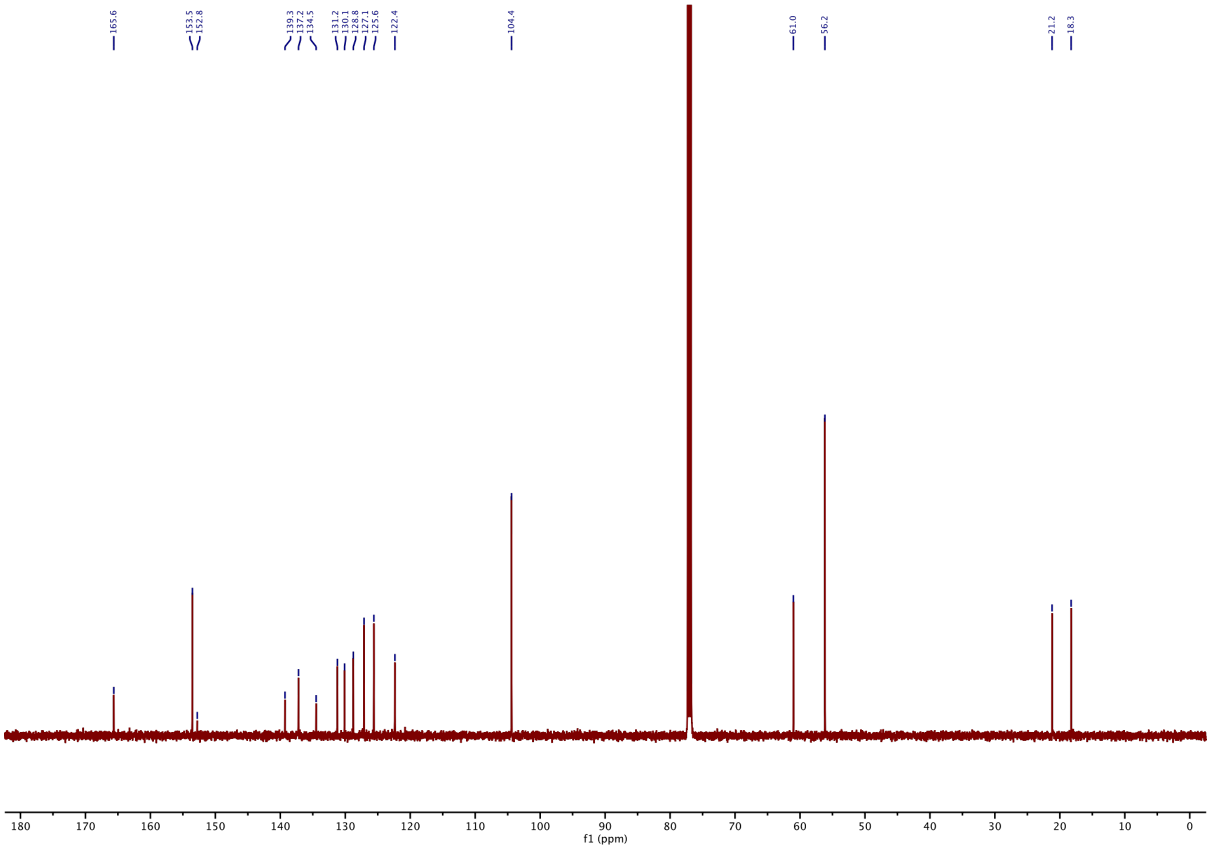

**[1]H-NMR of SBTub3P:**

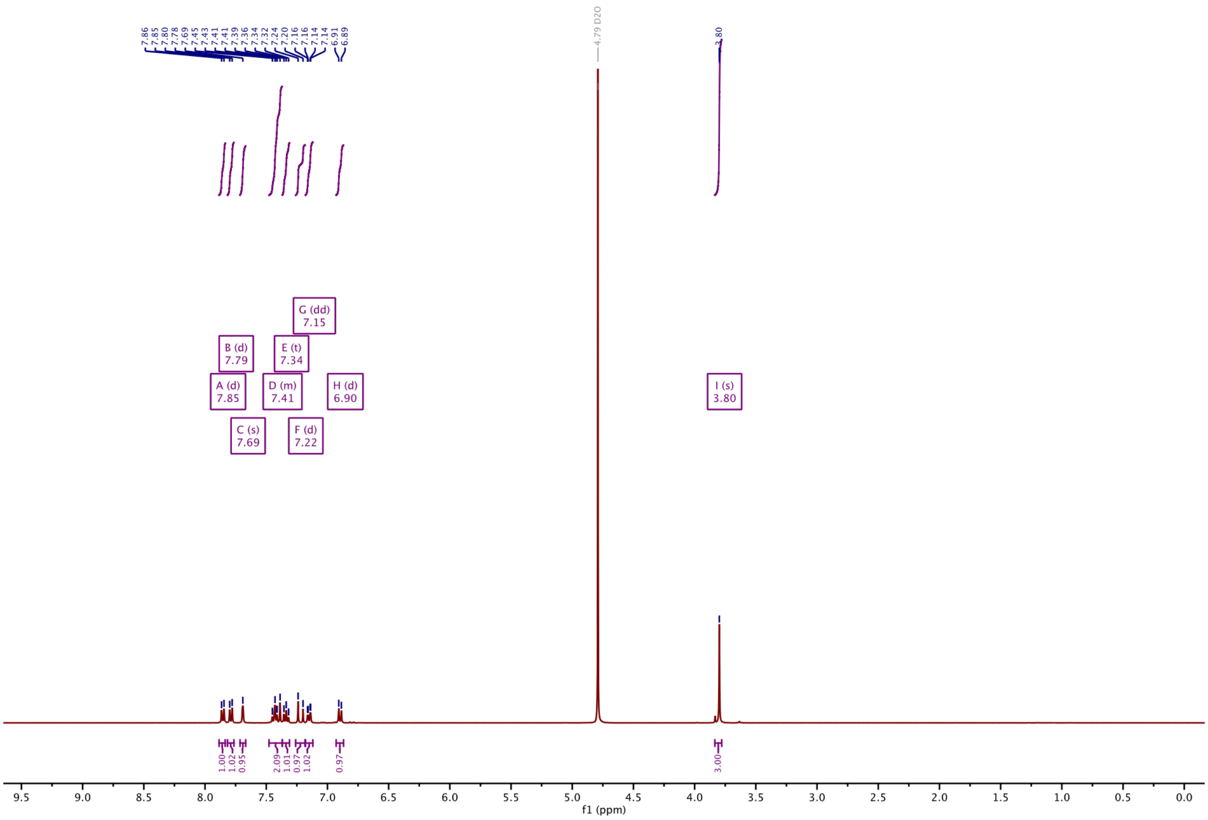

**[13]C-NMR of SBTub3P:**

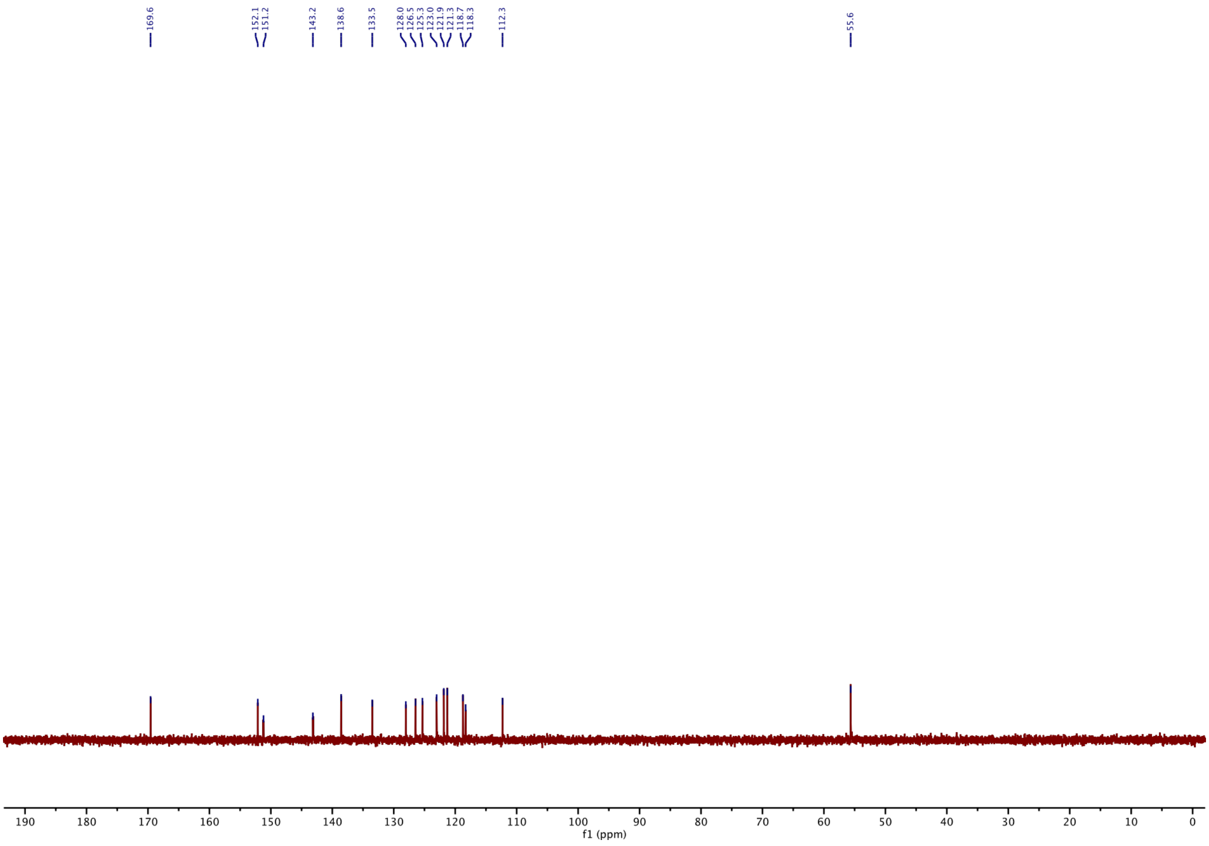

#### [1]H-NMR of Dibenzyl protected SBTubA4P:

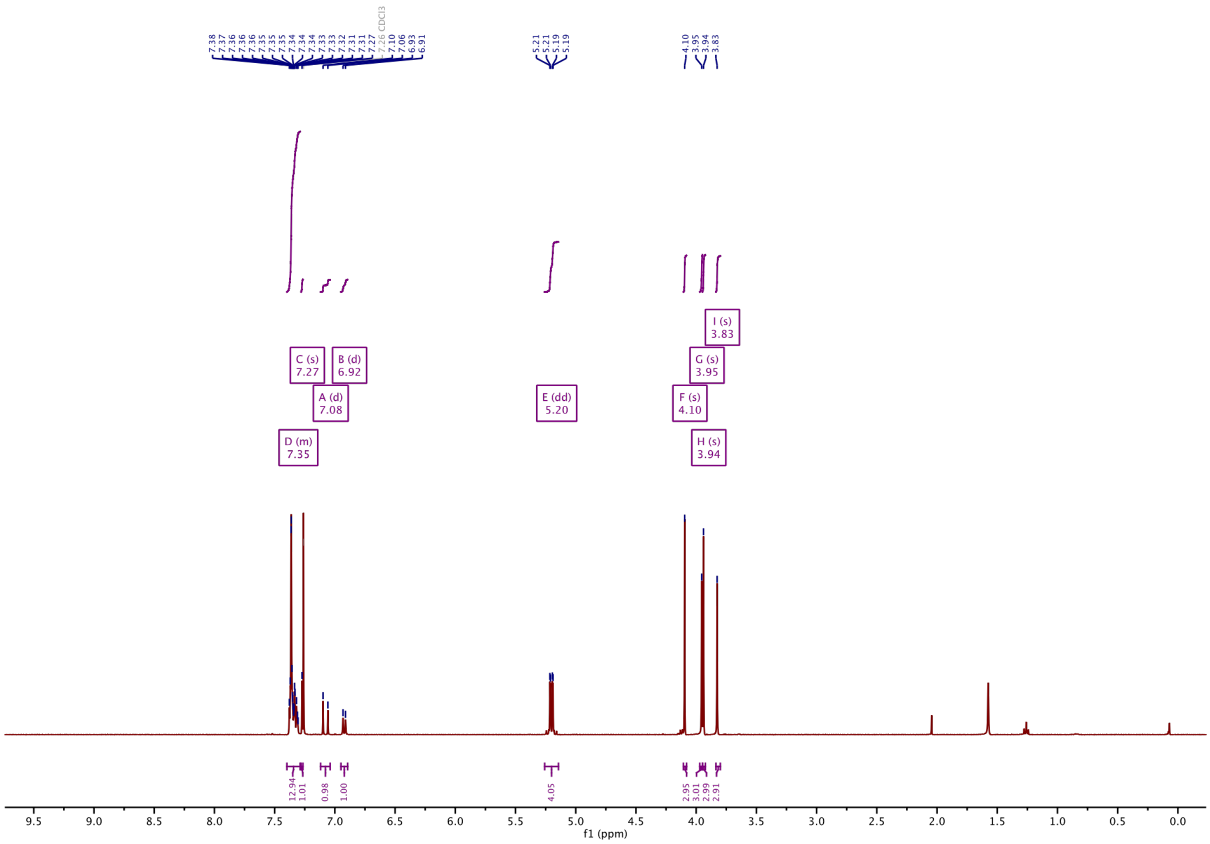

#### [13]C-NMR of Dibenzyl Protected SBTubA4P:

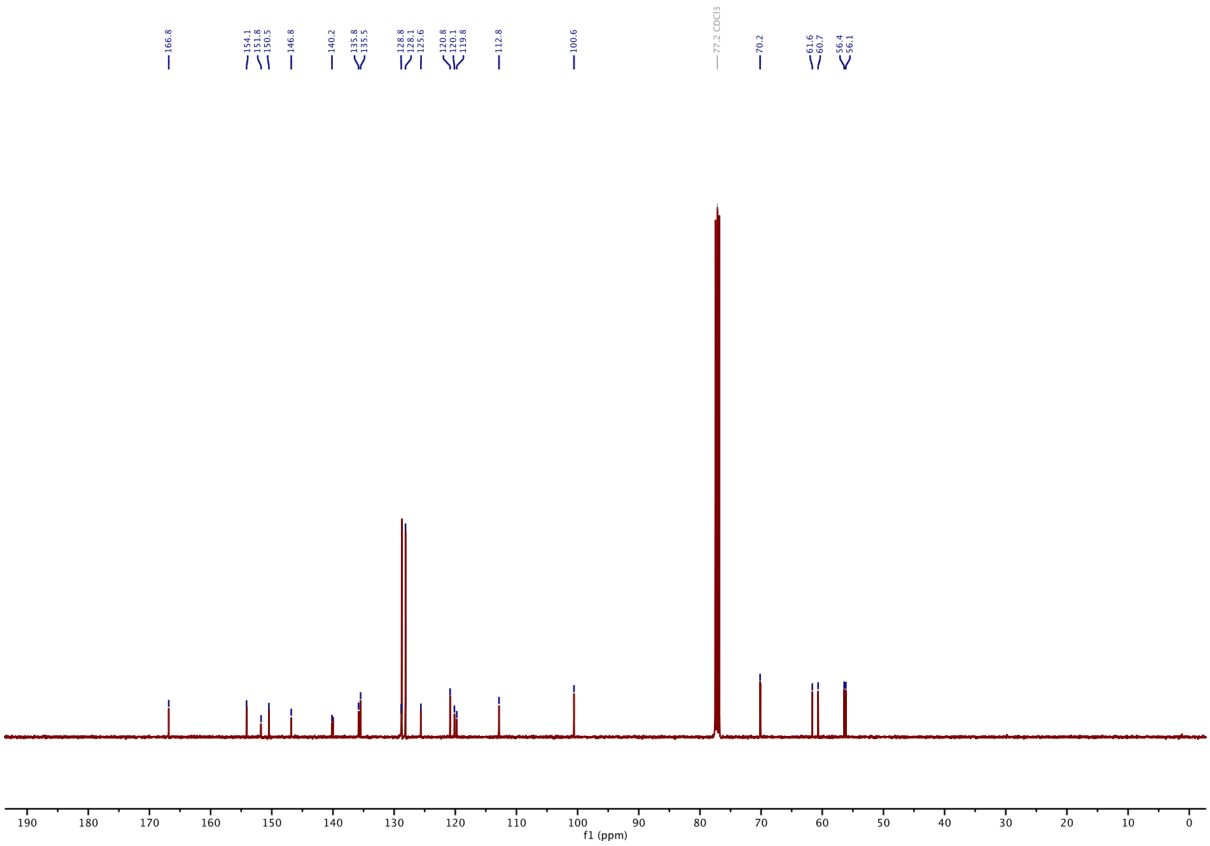

#### $^1$H-NMR of SBTubA4p:

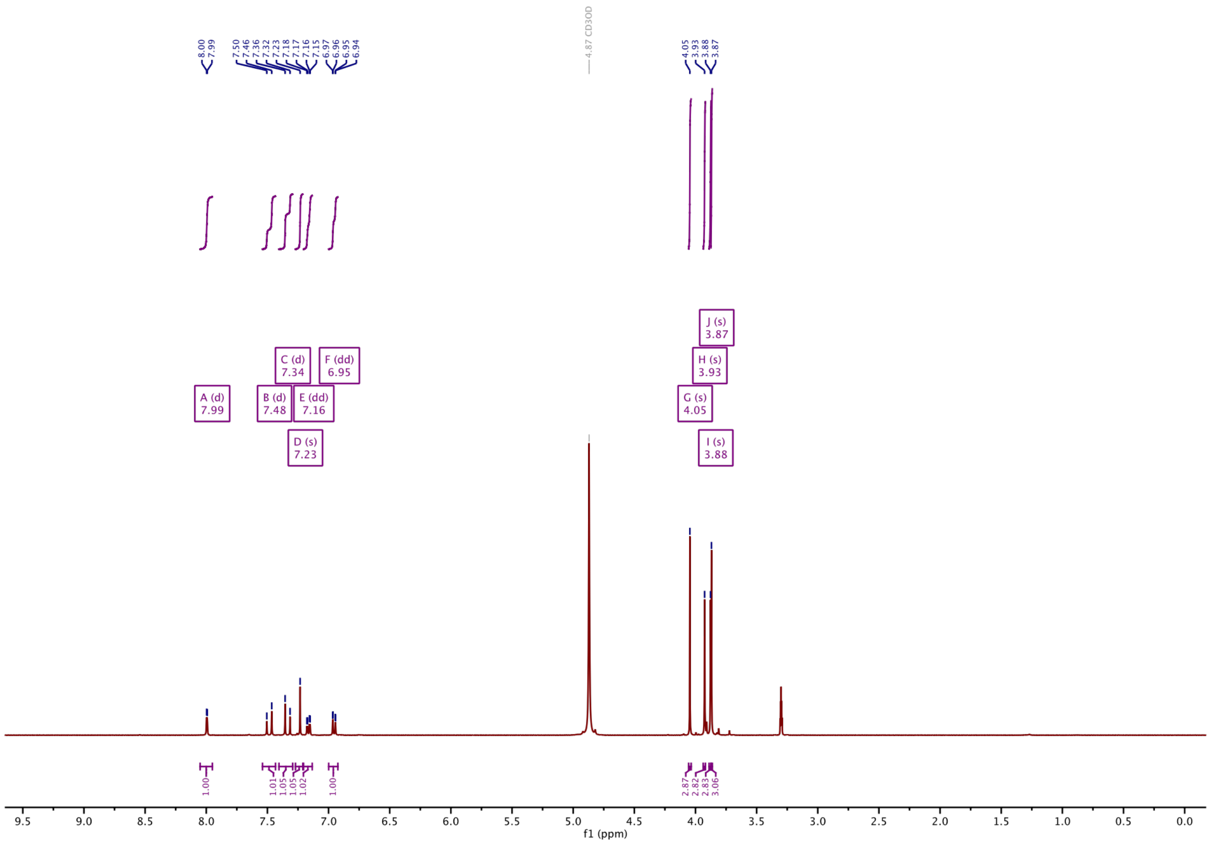

#### $^{13}$C-NMR of SBTubA4P:

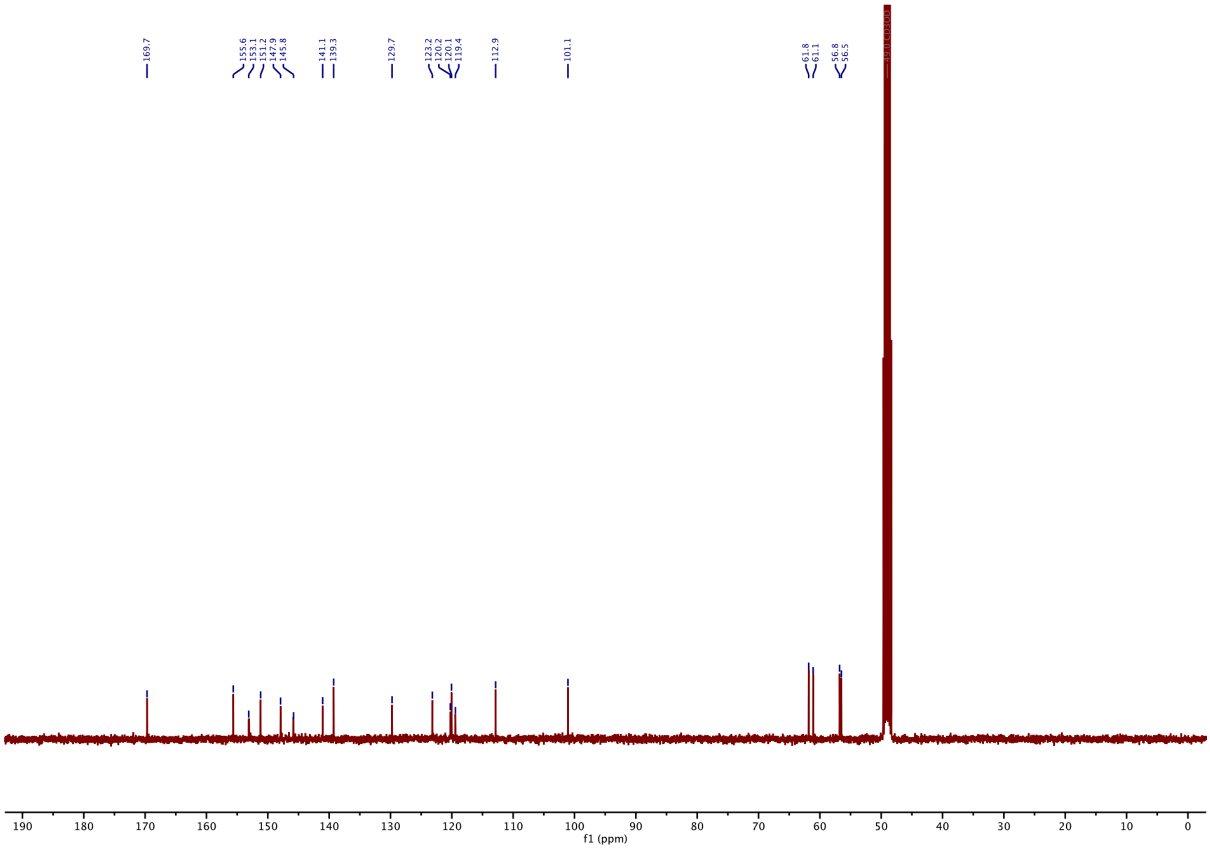

#### [1]H-NMR of S1:

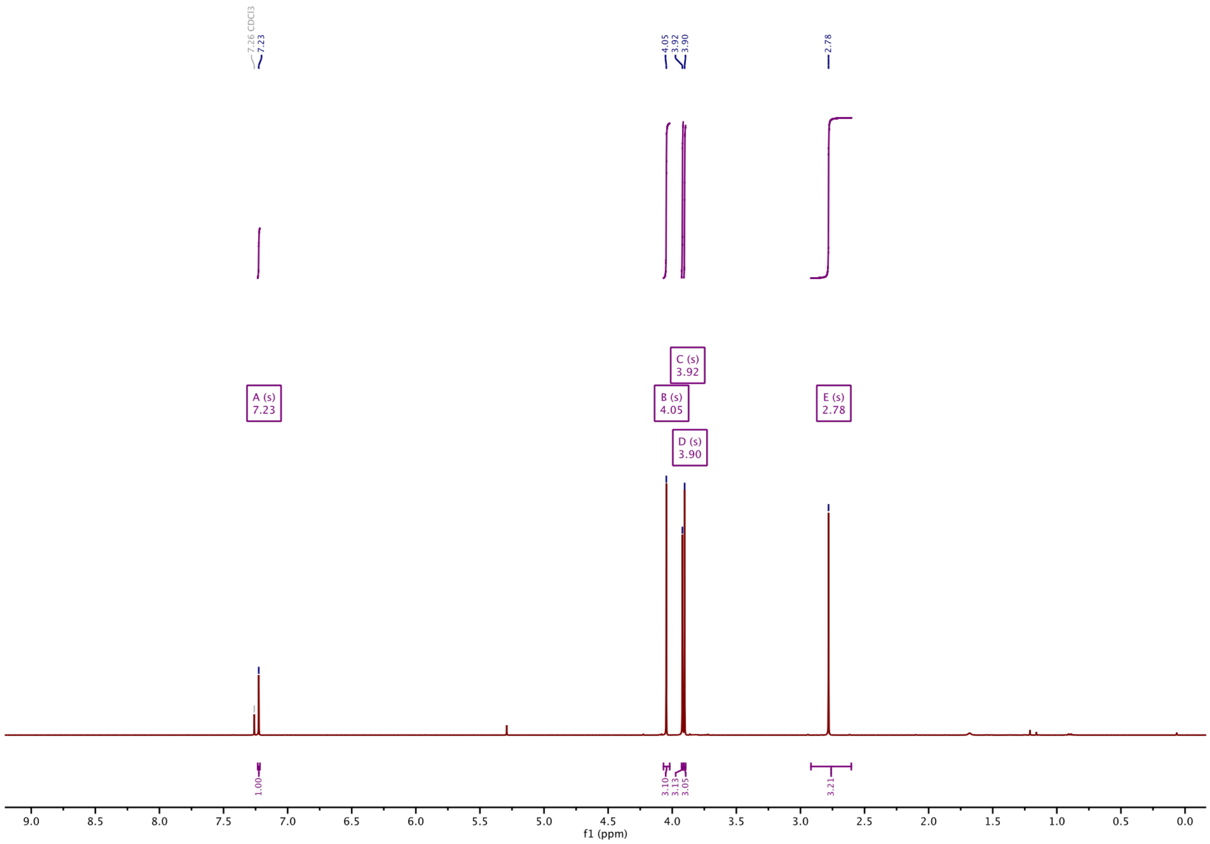

#### [13]C-NMR of S1:

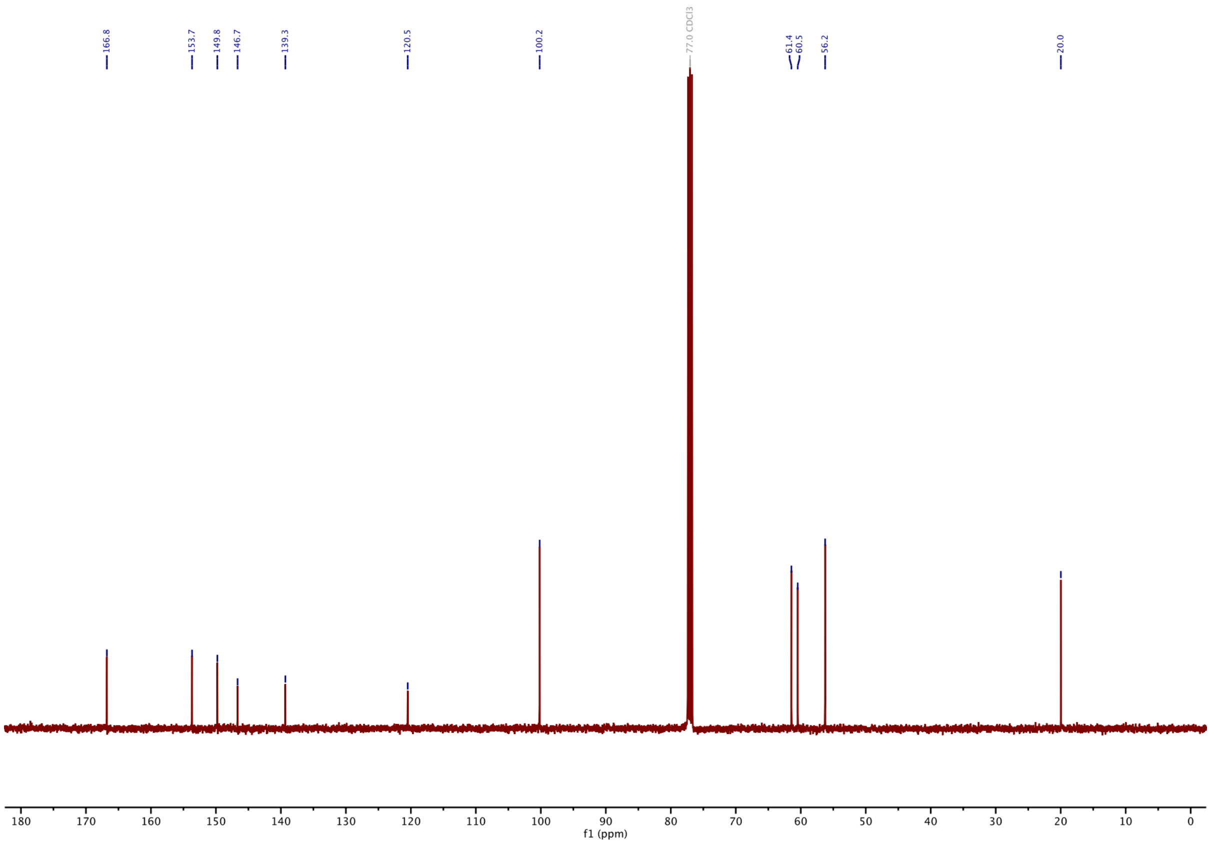

#### $^{1}$H-NMR of S2:

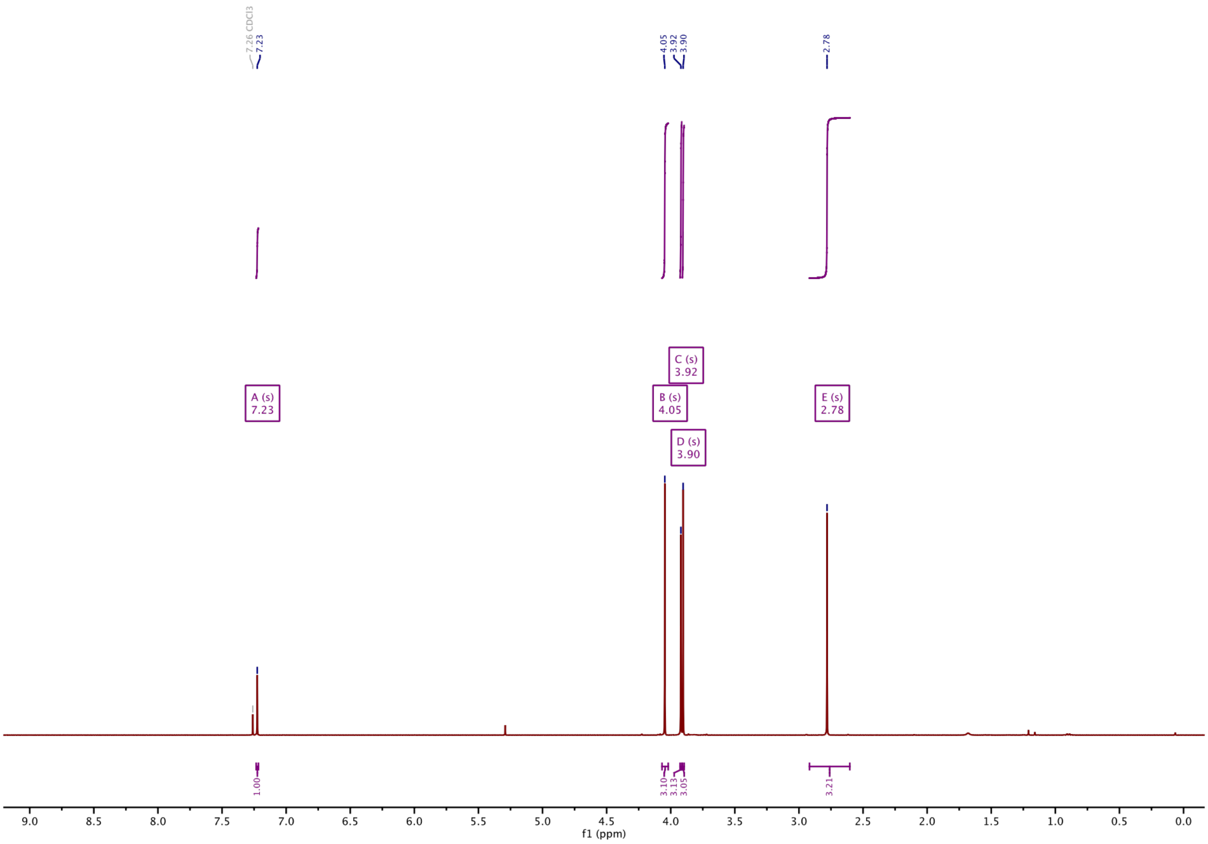

#### $^{13}$C-NMR of S2:

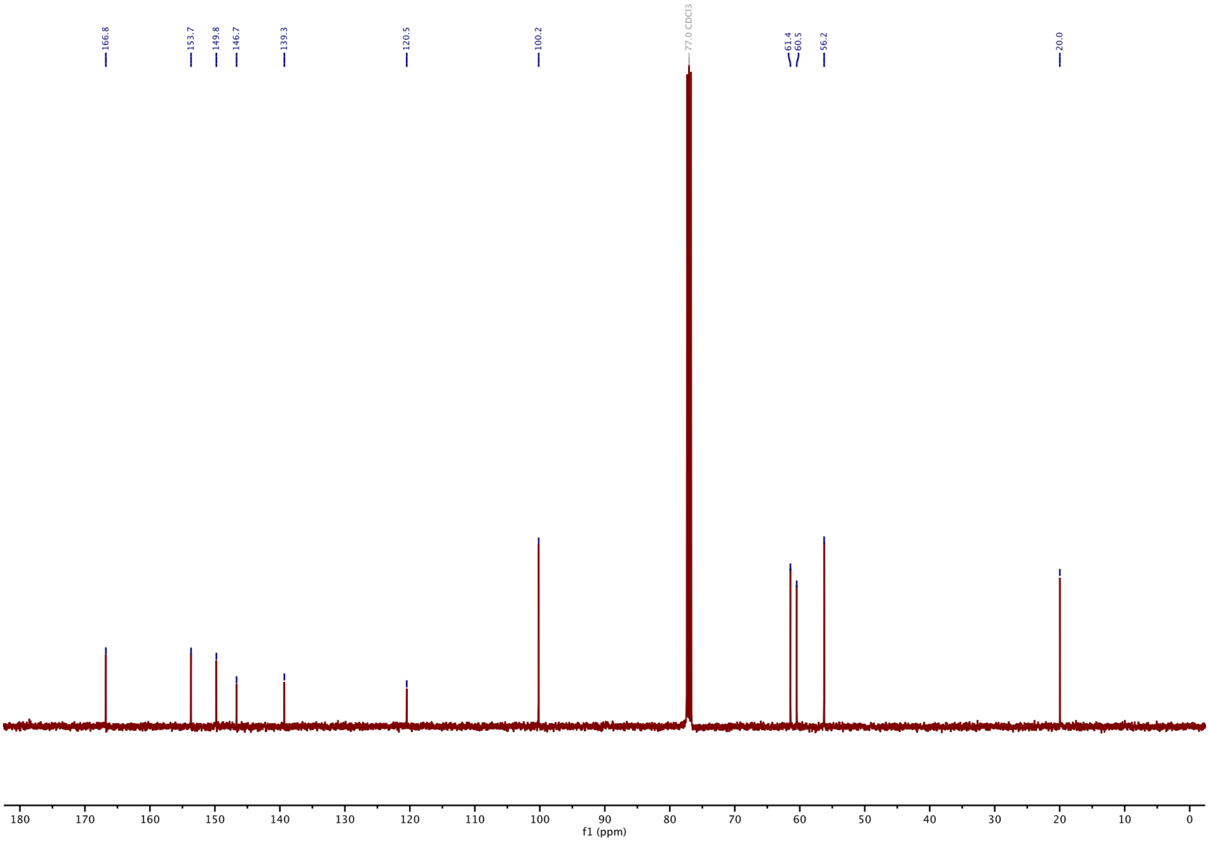

#### 1H-NMR of S3:

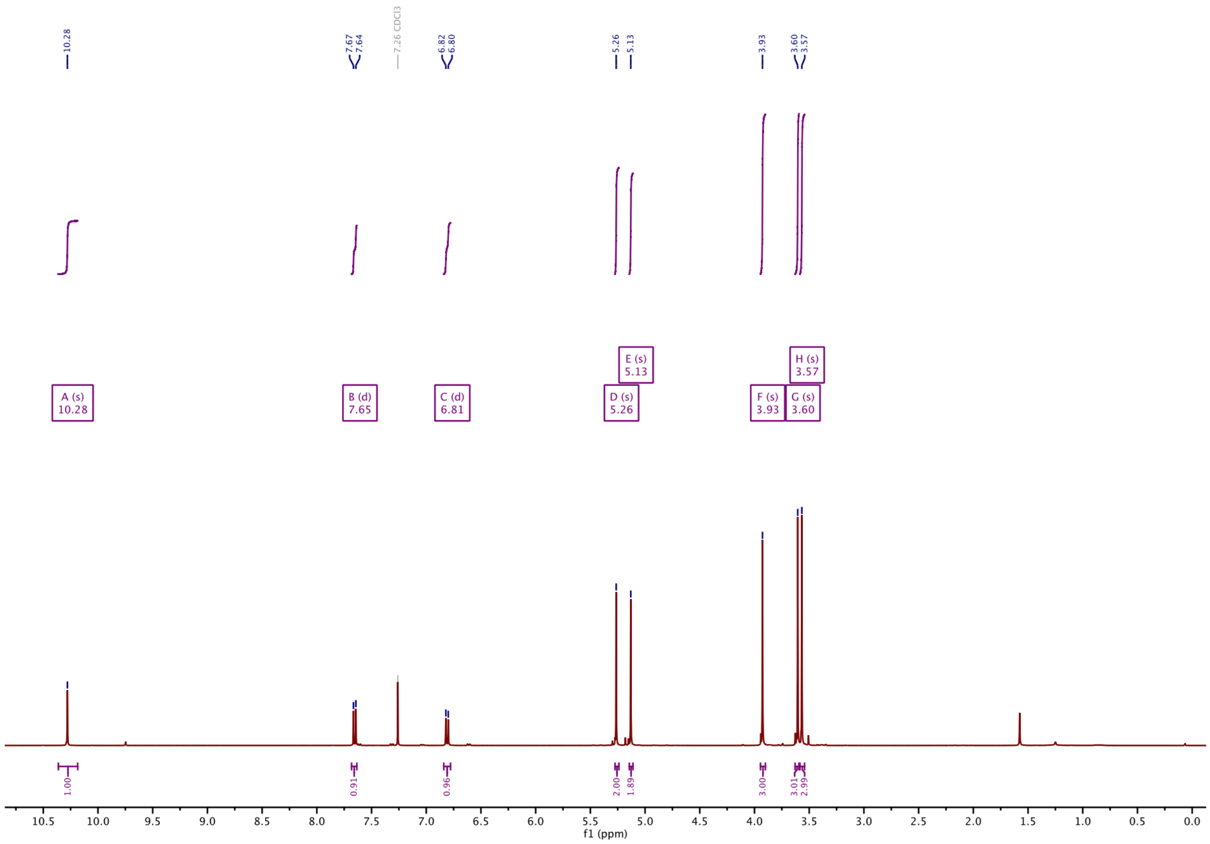

#### 13C-NMR of S3:

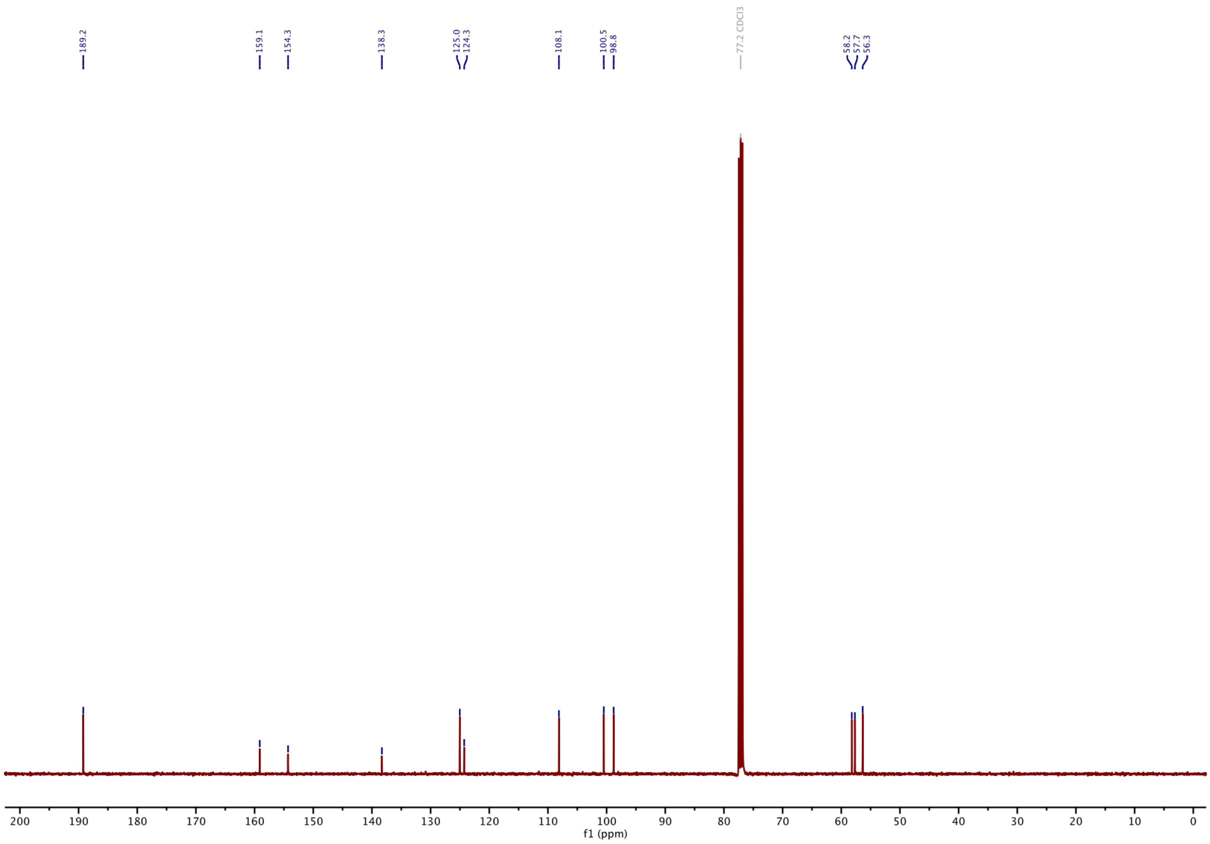

#### $^1$H-NMR of S4:

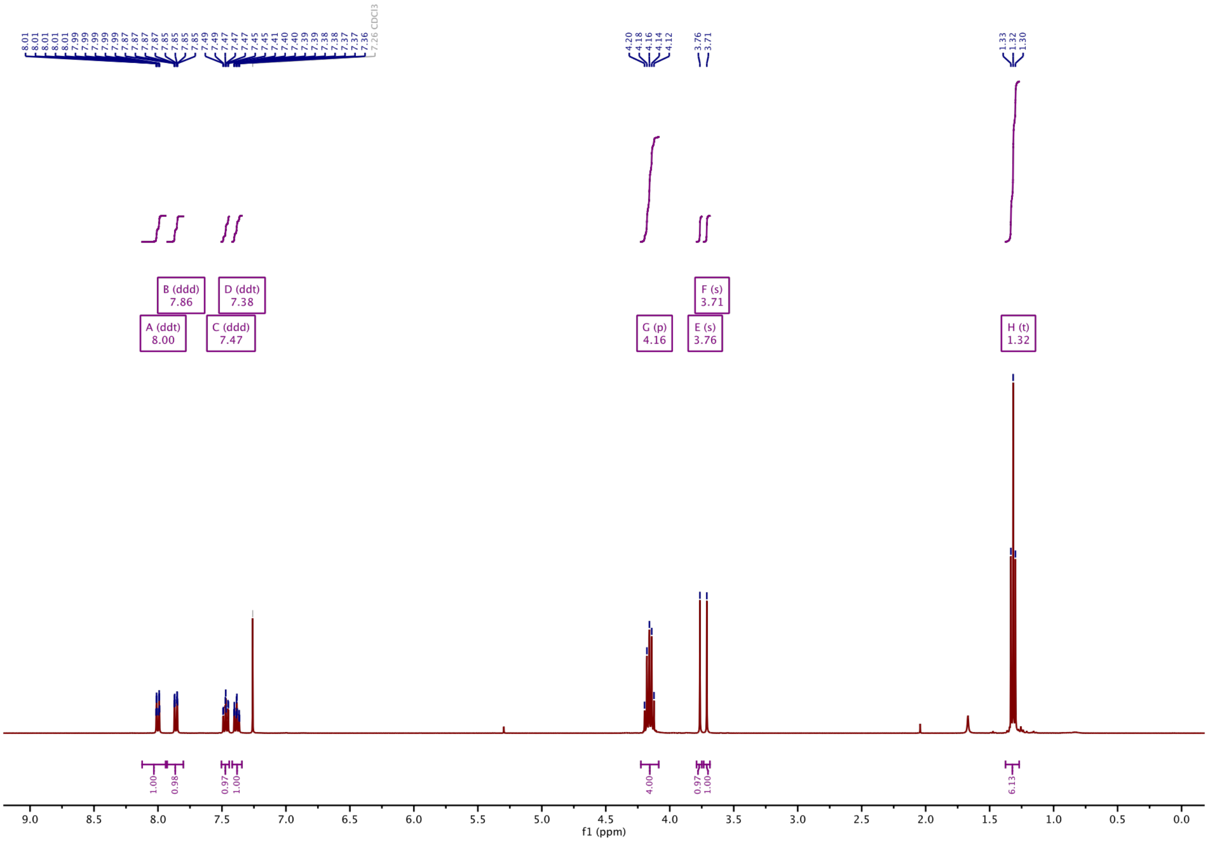

#### $^1$H-NMR of S5:

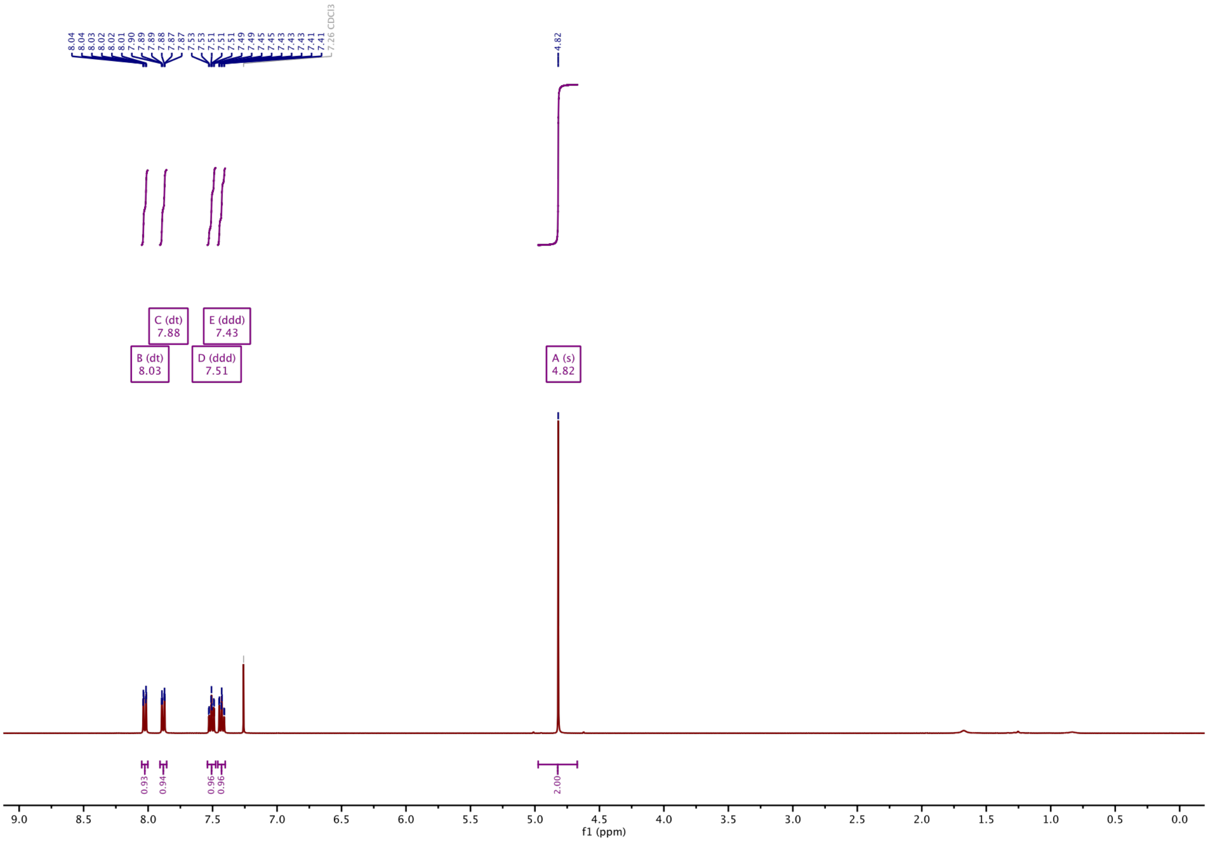

#### [1]H-NMR of S6:

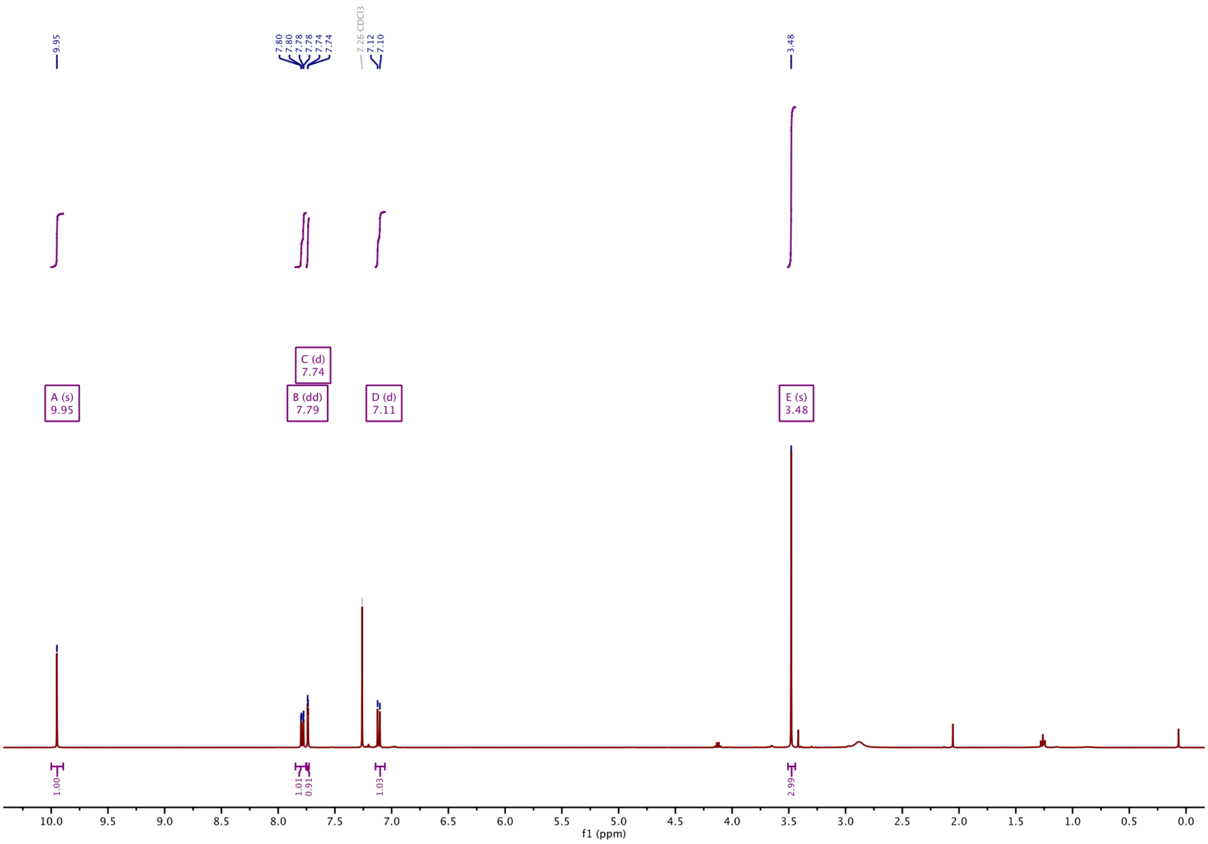

#### [1]H-NMR of S7 GAO461:

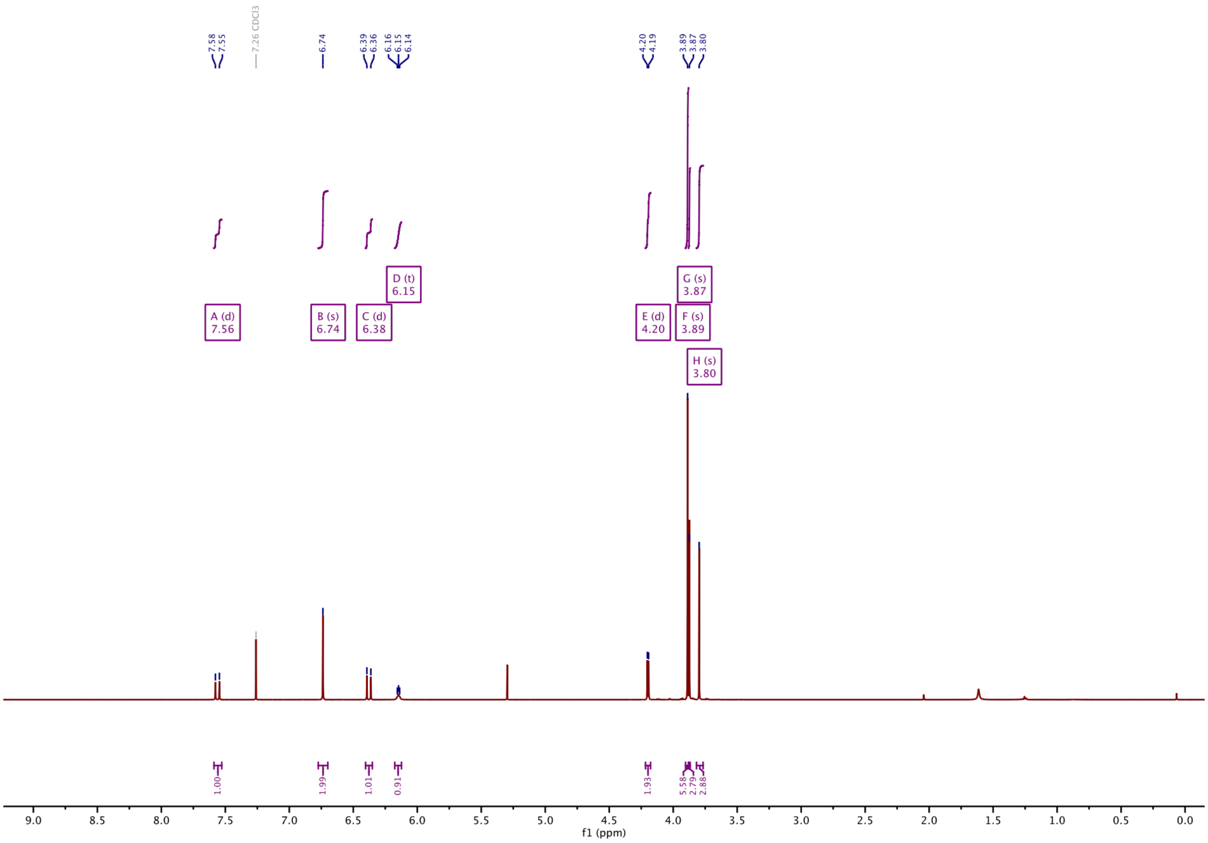

**$^{13}$C-NMR of S7 GAO461:**

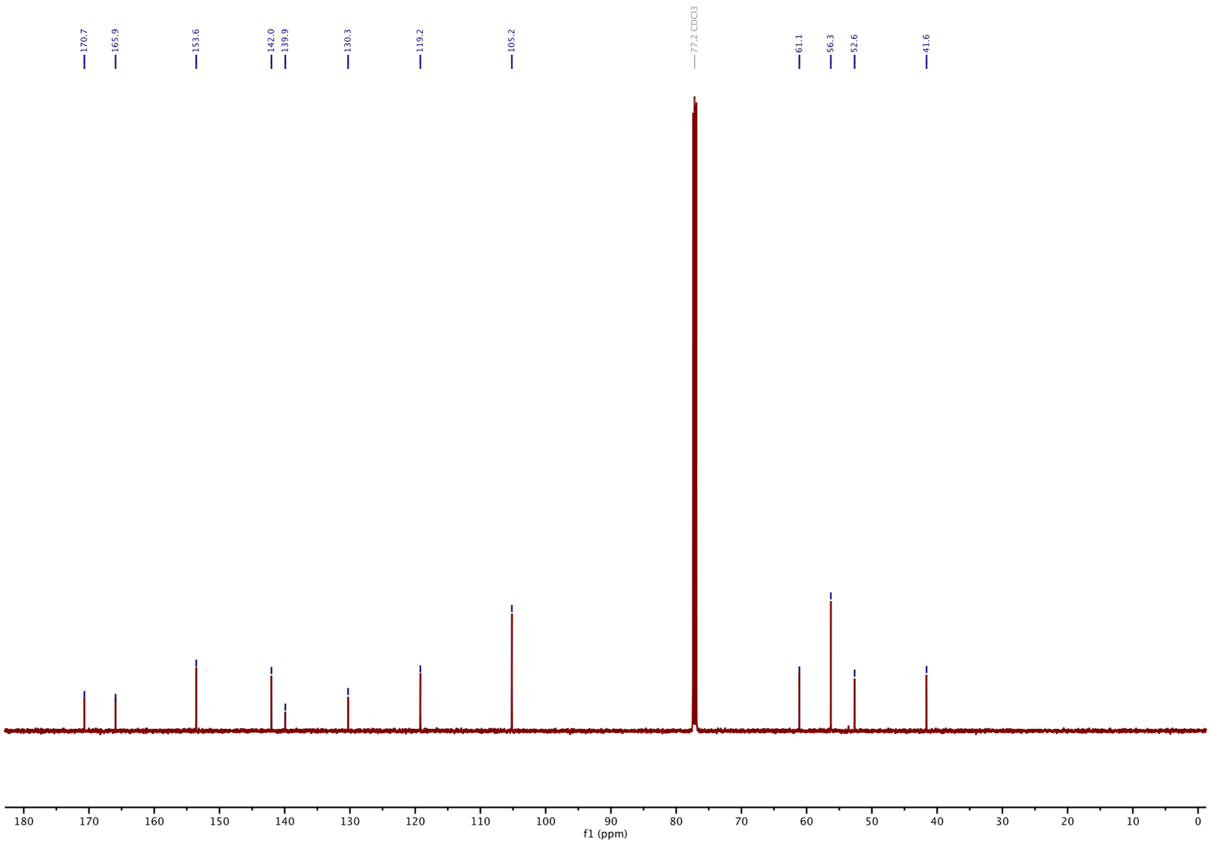

**$^{1}$H-NMR of S8 GAO274:**

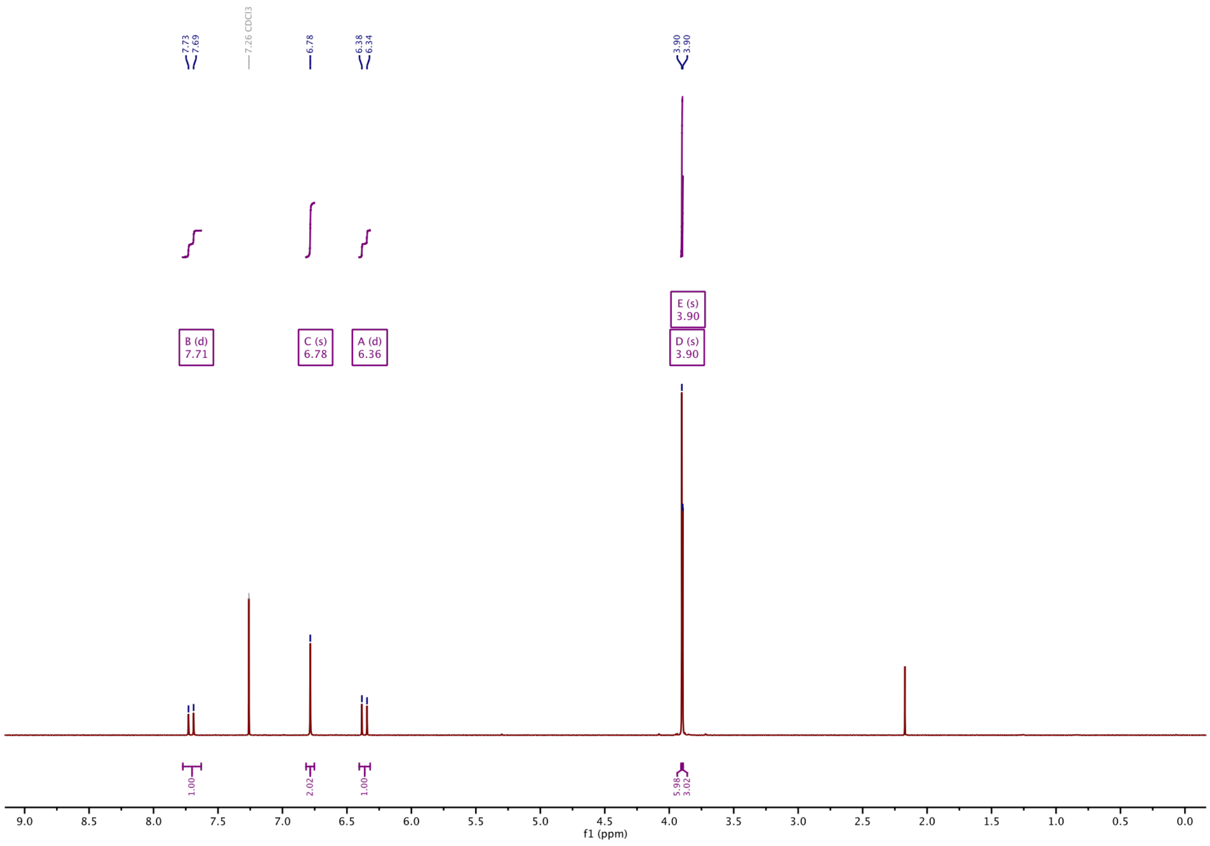

#### $^{13}$C-NMR of S8 GAO274:

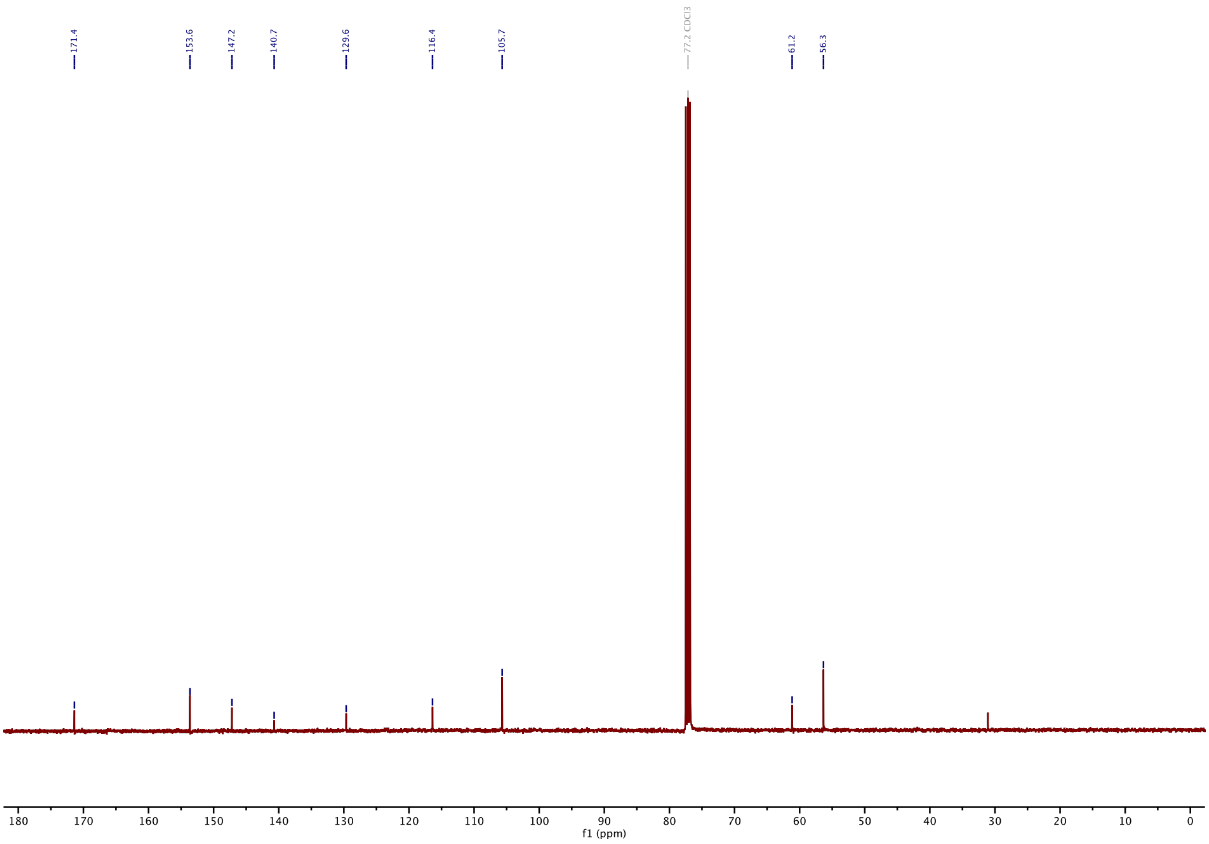

#### $^{1}$H-NMR of S9 GAO552:

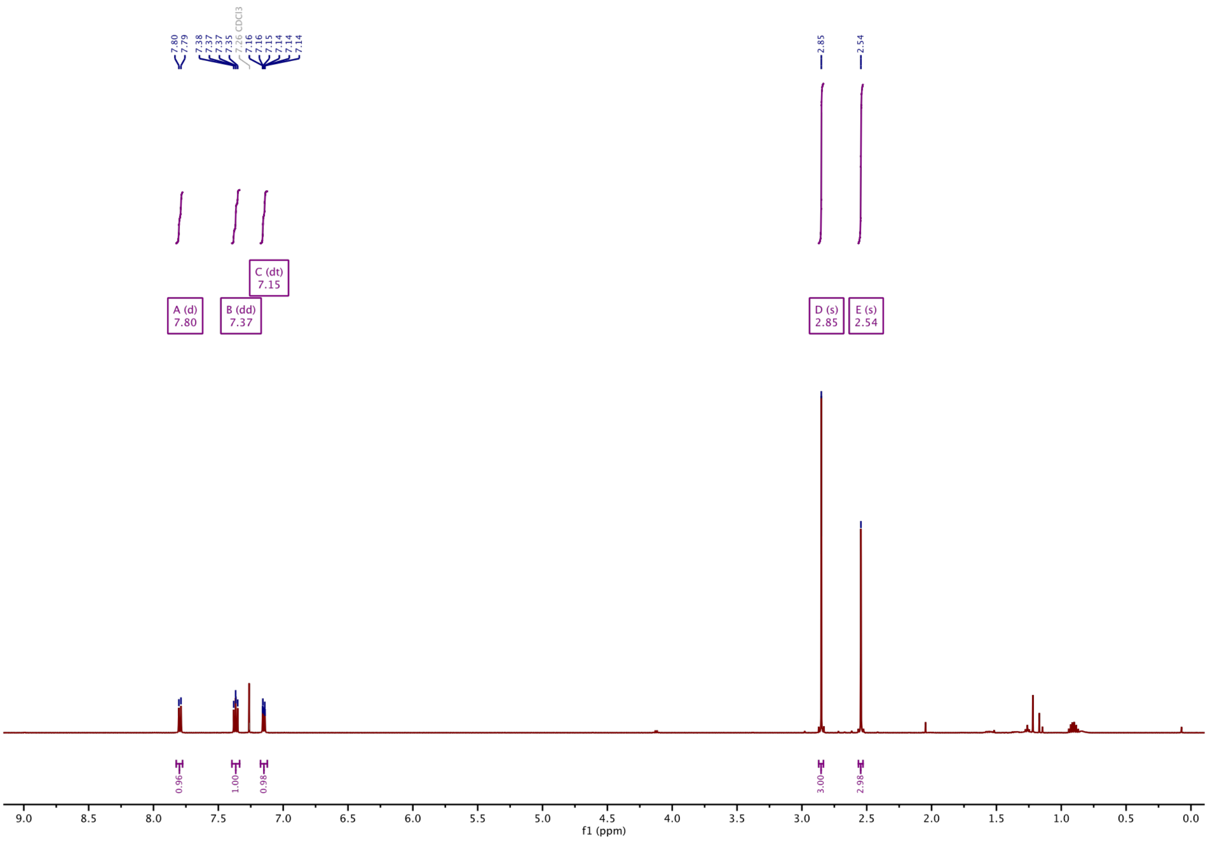

#### $^{13}$C-NMR of S9 GAO552:

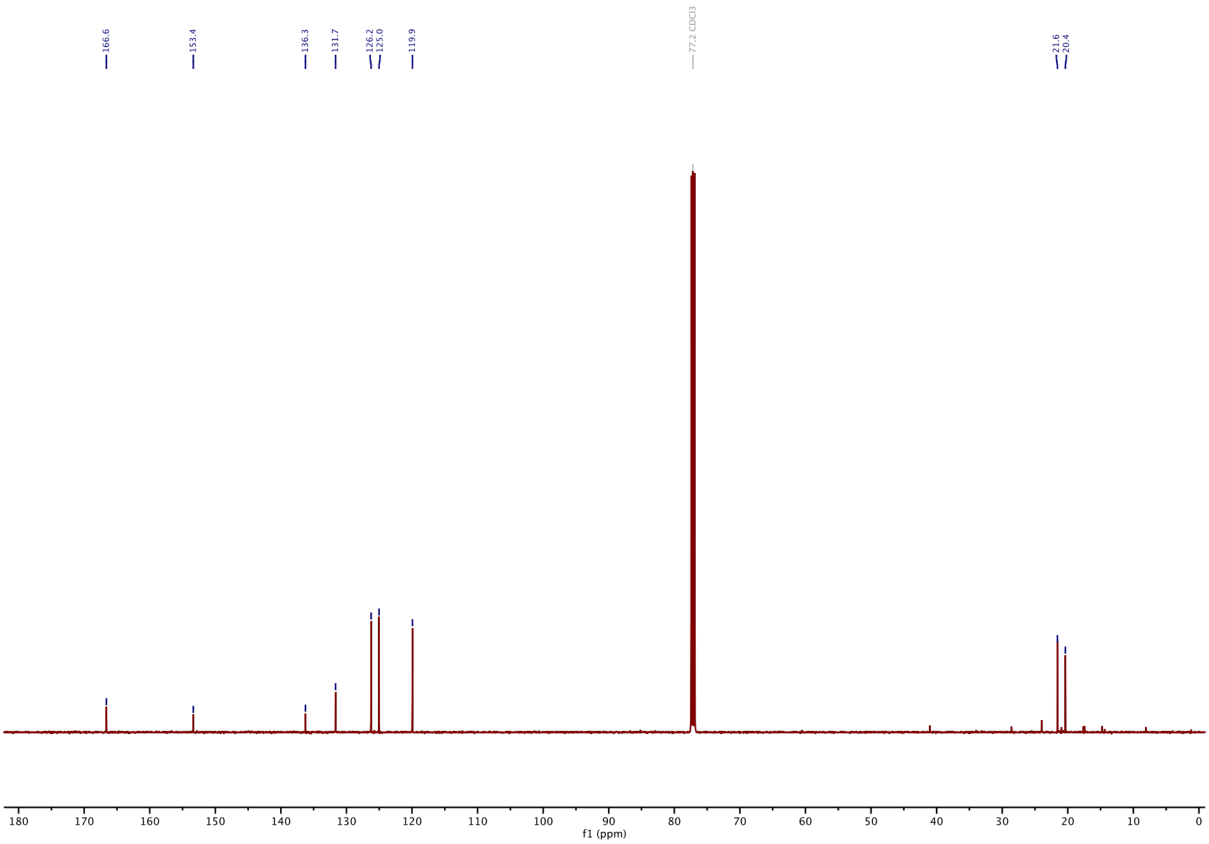

#### $^{1}$H-NMR of S10 GAO491:

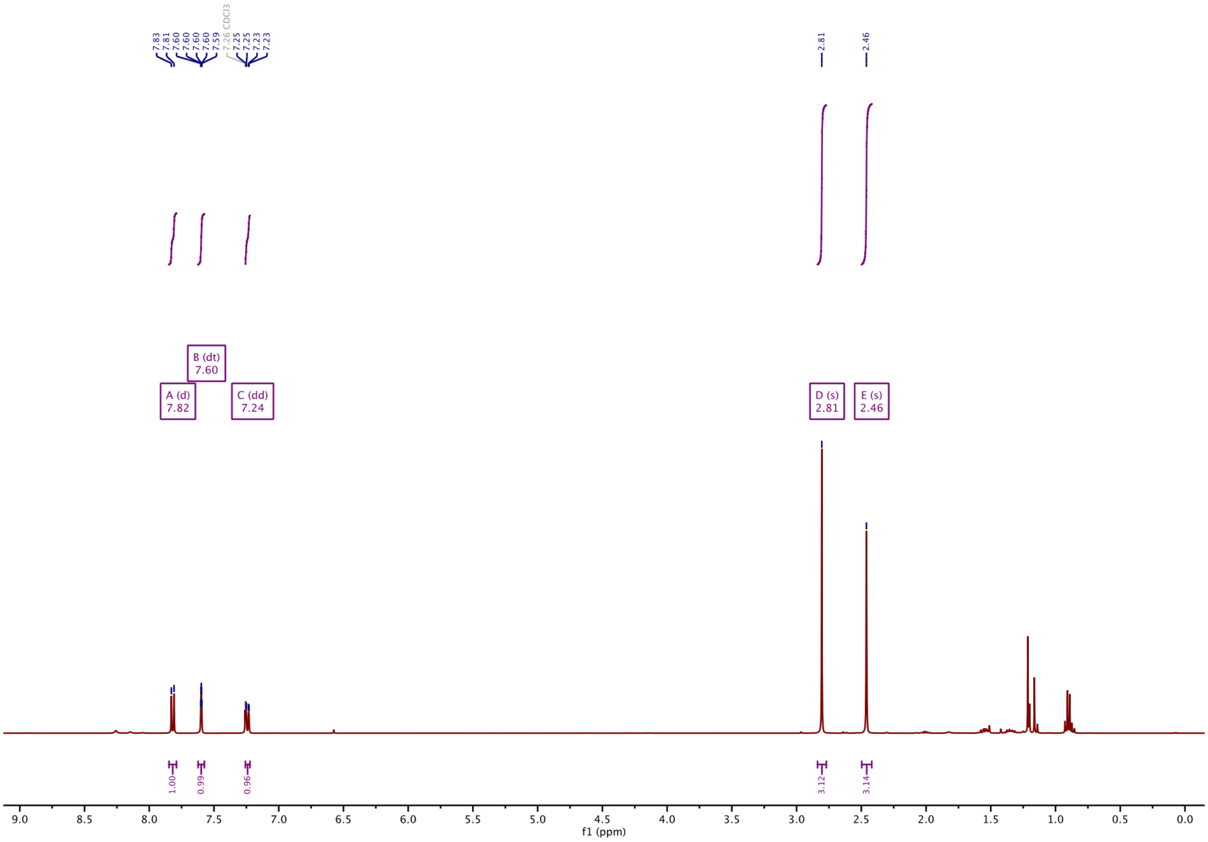

#### $^1$H-NMR of S11 GAO553:

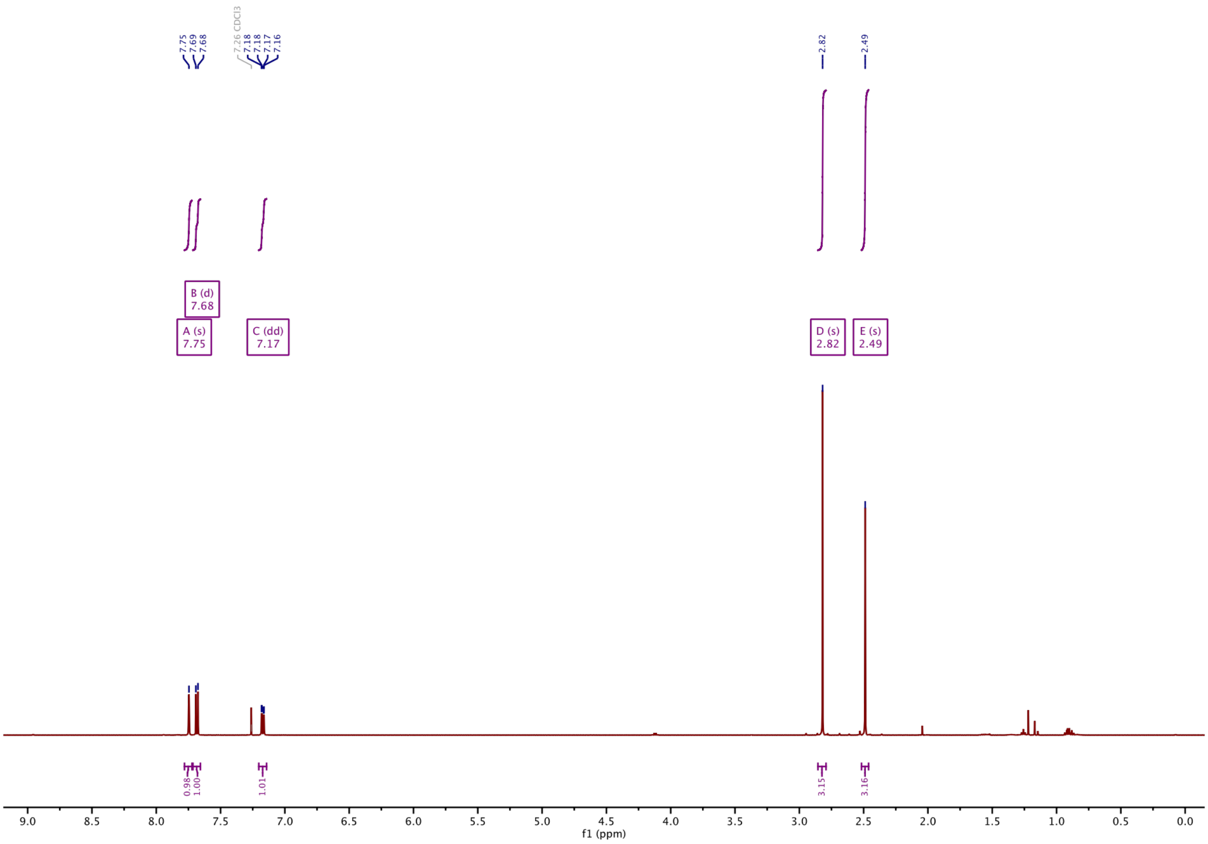

#### $^{13}$C-NMR of S11 GAO553:

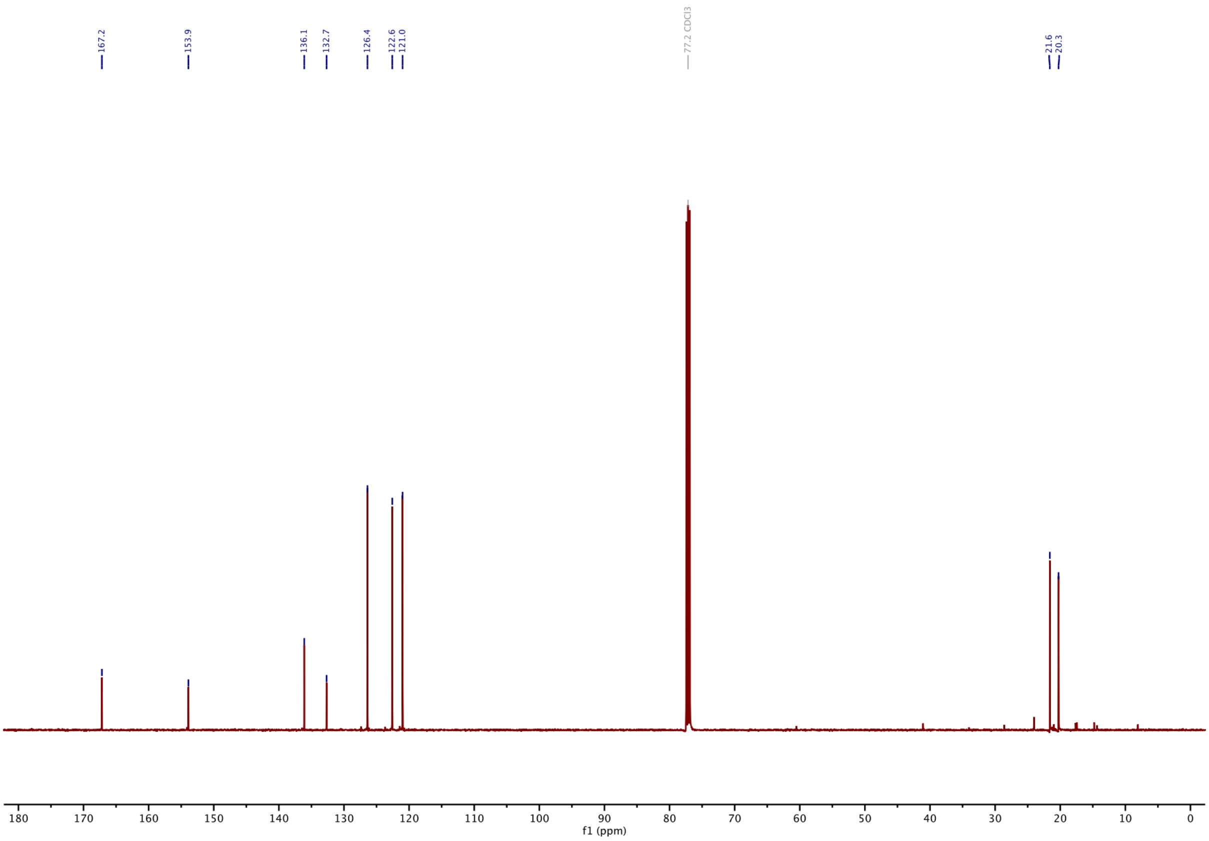

#### $^{1}$H-NMR of S12 GAO437:

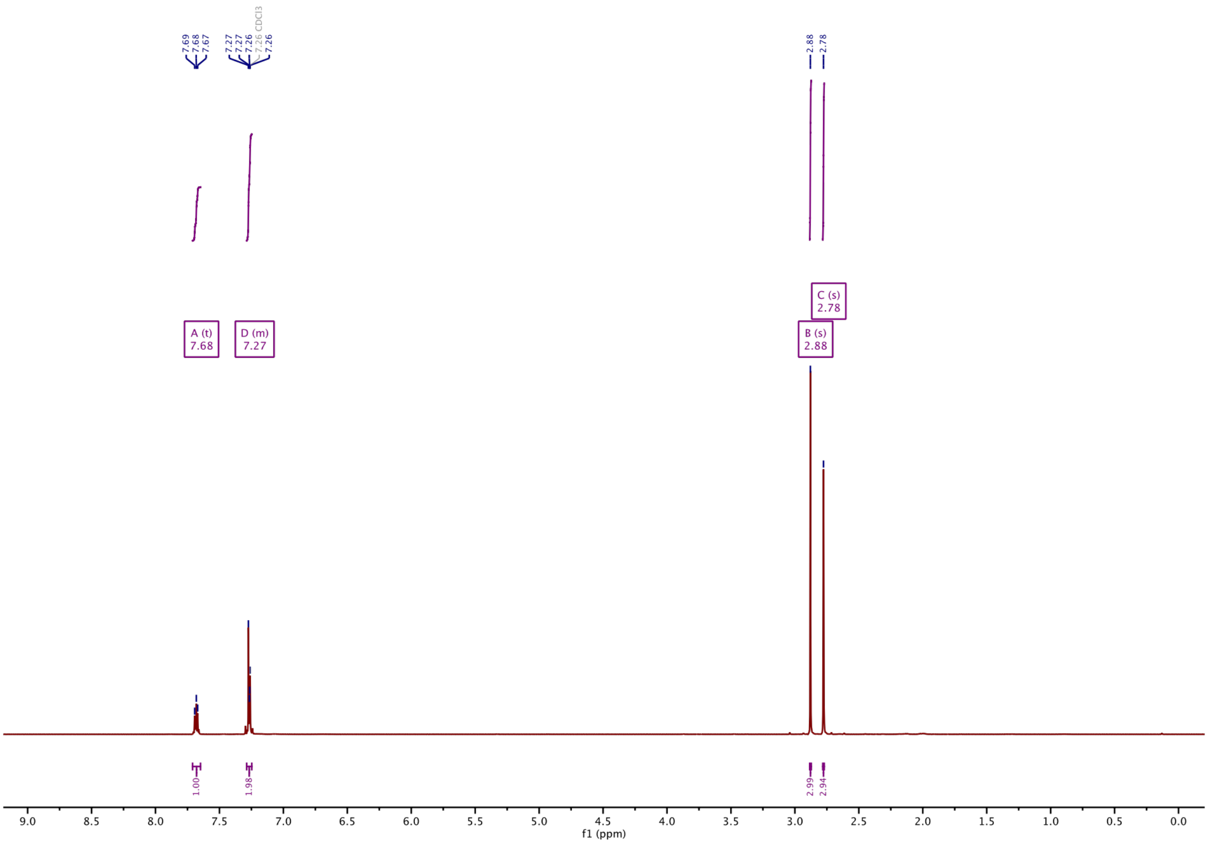

#### $^{13}$C-NMR of S12 GAO437:

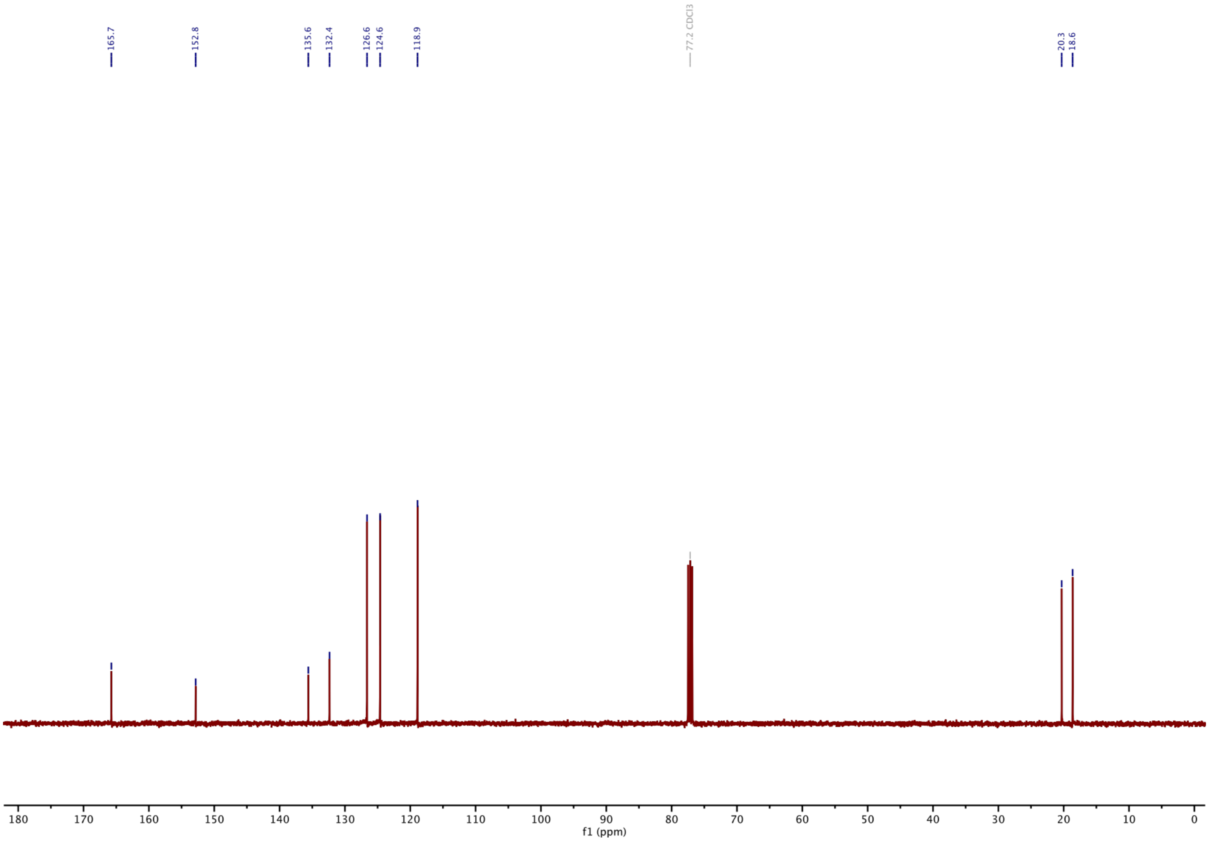

#### [1]H-NMR of S13 GAO490:

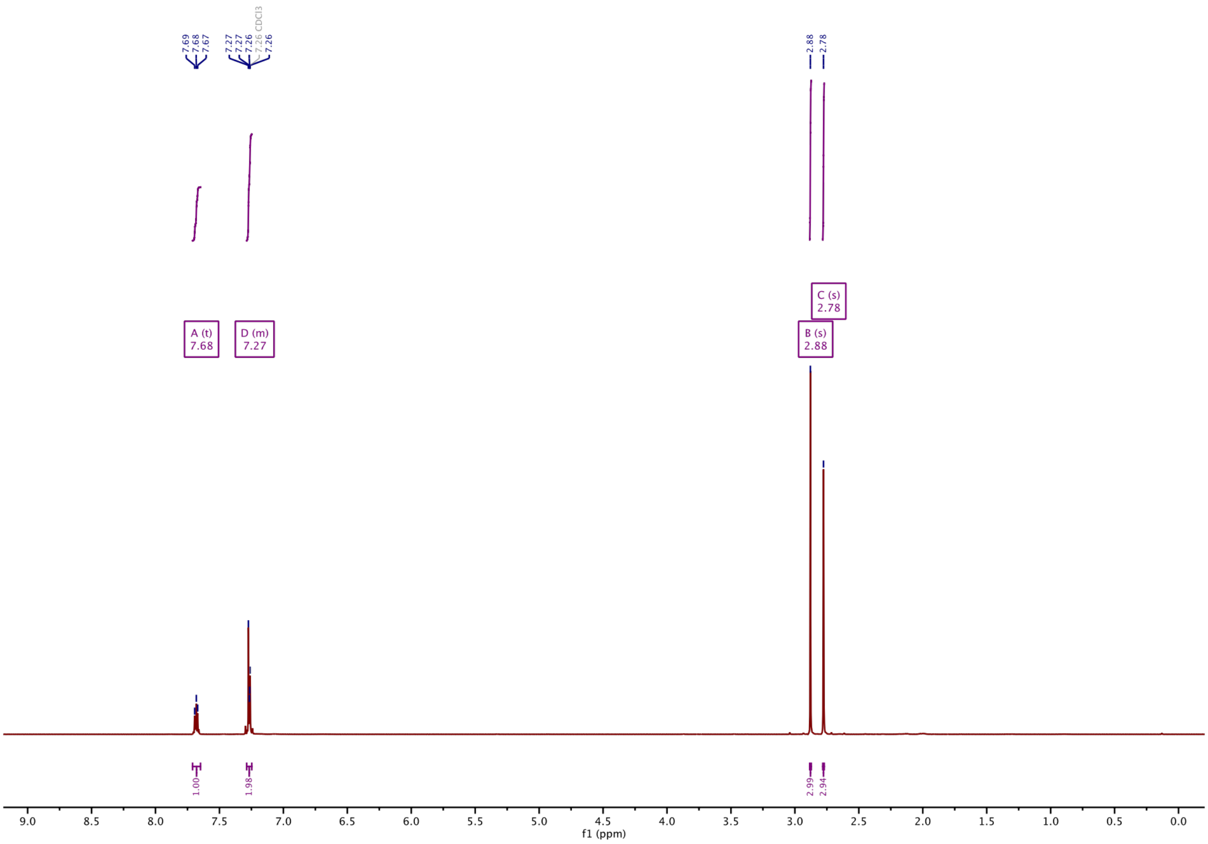

#### [13]C-NMR of S13 GAO490:

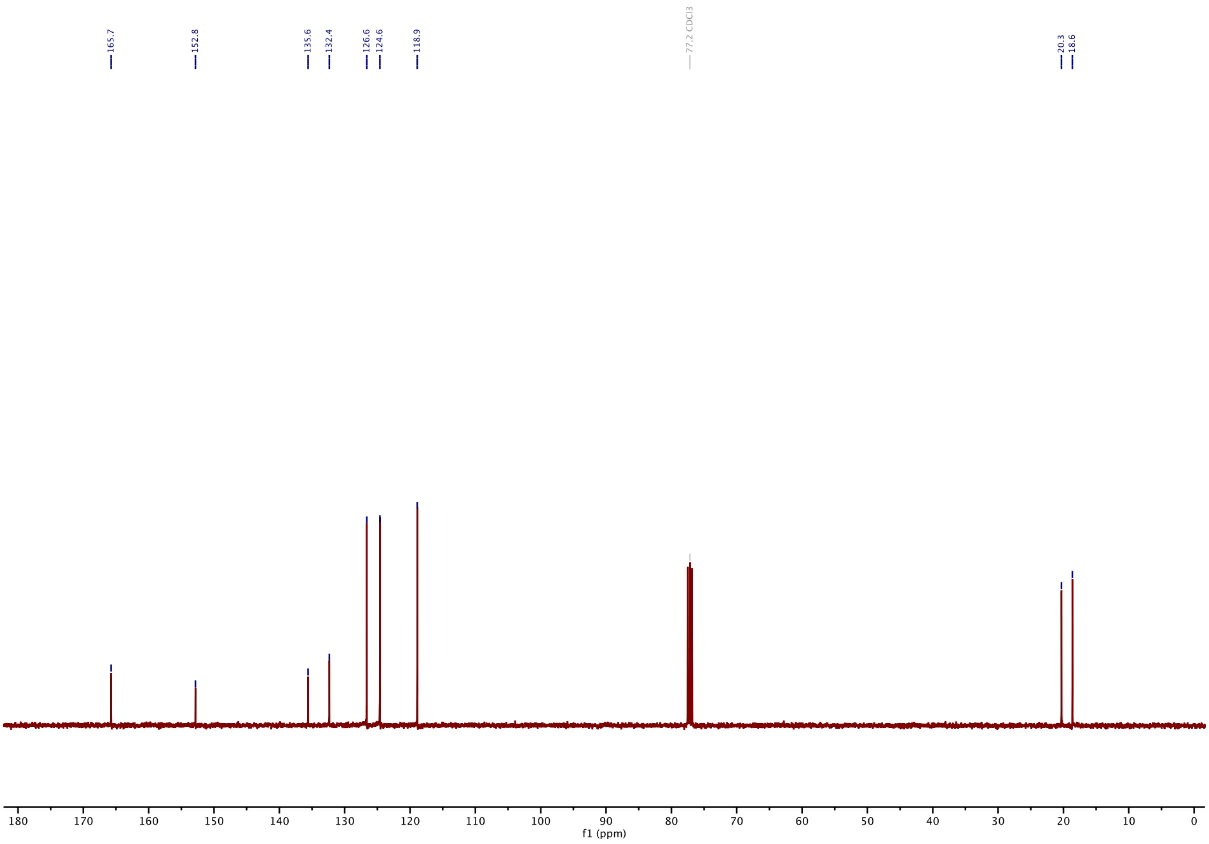

#### Part F: Bibliography

(30) Tickle, I. J.; Flensburg, C.; Keller, P.; Paciorek, W.; Sharff, A.; Smart, O.; Vonrhein, C.; Bricogne, G. *STARANISO Anisotropy & Bayesian Estimation Server*; Global Phasing Ltd: Cambridge, United Kingdom, 2018.

(31) Smart, O. S.; Womack, T. O.; Flensburg, C.; Keller, P.; Paciorek, W.; Sharff, A.; Vonrhein, C.; Bricogne, G. Exploiting Structure Similarity in Refinement: Automated NCS and Target-Structure Restraints in BUSTER. *Acta Crystallogr. D Biol. Crystallogr.* **2012**, *68* (4), 368–380. https://doi.org/10.1107/S0907444911056058.

(32) Adams, P. D.; Afonine, P. V.; Bunkoczi, G.; Chen, V. B.; Davis, I. W.; Echols, N.; Headd, J. J.; Hung, L.-W.; Kapral, G. J.; Grosse-Kunstleve, R. W.; McCoy, A. J.; Moriarty, N. W.; Oeffner, R.; Read, R. J.; Richardson, D. C.; Richardson, J. S.; Terwilliger, T. C.; Zwart, P. H. PHENIX: A Comprehensive Python-Based System for Macromolecular Structure Solution. *Acta Crystallogr. Sect. D* **2010**, *66* (2), 213–221. https://doi.org/10.1107/S0907444909052925.

(33) Emsley, P.; Cowtan, K. Coot: Model-Building Tools for Molecular Graphics. *Acta Crystallogr. Sect. D* **2004**, *60* (12 Part 1), 2126–2132. https://doi.org/10.1107/S0907444904019158.
